# GenErode: a bioinformatics pipeline to investigate genome erosion in endangered and extinct species

**DOI:** 10.1101/2022.03.04.482637

**Authors:** Verena E. Kutschera, Marcin Kierczak, Tom van der Valk, Johanna von Seth, Nicolas Dussex, Edana Lord, Marianne Dehasque, David W. G. Stanton, Payam Emami Khoonsari, Björn Nystedt, Love Dalén, David Díez-del-Molino

## Abstract

**Background:** Many wild species have suffered drastic population size declines over the past centuries, which have led to ‘genomic erosion’ processes characterized by reduced genetic diversity, increased inbreeding, and accumulation of harmful mutations. Yet, genomic erosion estimates of modern-day populations often lack concordance with dwindling population sizes and conservation status of threatened species. One way to directly quantify the genomic consequences of population declines is to compare genome-wide data from pre-decline museum samples and modern samples. However, doing so requires computational data processing and analysis tools specifically adapted to comparative analyses of degraded, ancient or historical, DNA data with modern DNA data as well as personnel trained to perform such analyses.

**Results:** Here, we present a highly flexible, scalable, and modular pipeline to compare patterns of genomic erosion using samples from disparate time periods. The GenErode pipeline uses state-of-the-art bioinformatics tools to simultaneously process whole-genome re-sequencing data from ancient/historical and modern samples, and to produce comparable estimates of several genomic erosion indices. No programming knowledge is required to run the pipeline and all bioinformatic steps are well-documented, making the pipeline accessible to users with different backgrounds. GenErode is written in Snakemake and Python3 and uses Conda and Singularity containers to achieve reproducibility on high-performance compute clusters. The source code is freely available on GitHub (https://github.com/NBISweden/GenErode).

**Conclusions:** GenErode is a user-friendly and reproducible pipeline that enables the standardization of genomic erosion indices from temporally sampled whole genome re-sequencing data.

## Background

A plethora of large scale projects are set to generate *de novo* assemblies for all eukaryotic species in the next few years (e.g. [1–3]), thus solving one of the traditional limitations of conservation genomic studies, namely the absence of reference genomes for non-model species. In all these projects, contributing to biodiversity conservation appears among the main objectives. They propose to do this by enabling genomic studies in endangered species, which has been sometimes presented as the solution to all sorts of conservation problems [4, 5]. Yet there are currently only a limited number of examples where genomic data has had concrete conservation impacts (e.g., [6–8]). While the reasons for this are several-fold, two of the most relevant ones are existing biases among genomic studies due to the lack of standard measures, and absence of validation of results using empirical data [9].

Human activities have caused populations of many wild species to suffer severe declines over the past few centuries [10]. These dwindling population sizes have led to an increased risk of extinction [11] as a consequence of genetic processes, such as the loss of genomic diversity, increased inbreeding, and accumulation of harmful mutations, processes together referred to as ‘genomic erosion’ [12–14]. Therefore, genomic approaches aimed at identifying the role of genomic erosion in shaping the fate of endangered populations are key to implementing more effective conservation efforts. Unfortunately, correlations between genomic erosion indices and current population sizes and/or the conservation status of endangered species are weak [15], which further complicates the inclusion of genetic parameters as a criterion to establish threat categories. This is because ancient bottlenecks or life-history traits typically overshadow the impact of the recent human-driven declines [16]. Consequently, interspecific comparisons of genome-wide diversity, inbreeding, and mutational load are poor predictors of population size and conservation status between modern-day endangered species. One powerful solution to this issue is to obtain genomic data from historical or ancient specimens that predate the recent demographic declines. Such temporal data can then be used to establish pre-decline baselines and enable direct quantification of the rate of change in genomic erosion indices that have resulted from recent population declines [15].

A second important limitation to bridging the gap between genomic studies of endangered species and applications in conservation is the need for advanced bioinformatics knowledge [9]. Handling and analyzing genomic data requires expertise in processing large-scale DNA datasets and programming skills that, sometimes, conservation biologists and wildlife managers are lacking. On top of this, the bioinformatic processing of the data and the interpretation of the analyses can be further complicated by the special characteristics of degraded historical and ancient DNA. Finally, reproducibility (obtaining the same results from analyzing the same data) is lacking across scientific disciplines [17]. However, genomic erosion indices can only be meaningful, comparable measures, if the results are reproducible. Therefore the increased access to user-friendly software and structured pipelines that generate reproducible results can be key factors for genomic data to be regularly applied to conservation [18].

With the aim to address these issues, we present GenErode, a highly flexible and modular pipeline focused on the user-friendly application of genomic data for conservation genomics. It performs analyses of wholegenome re-sequencing data, enabling the analysis of temporally sampled genomes with the goal to investigate patterns of genome erosion. GenErode employs state-of-the-art techniques to bioinformatically process genomic data and can handle both ancient/historical and modern samples simultaneously. It also generates estimates of commonly used genomic erosion indices (e.g., genome-wide diversity, inbreeding, and mutational load) with the objective of making them directly comparable between time periods.

## Implementation

### Description

In this pipeline, ancient/historical and modern whole-genome re-sequencing data are mapped to a reference genome assembly and processed according to the characteristics of each data type, aiming to make them comparable among different time periods (Figure 1). While it is optimized for the analysis of vertebrate re-sequencing data, all of the data processing tracks can be applied to a variety of organisms, including, for example, haploid insects. Written in Snakemake [19], this pipeline can be run on Linux systems such as high-performance computing (HPC) clusters.

**Figure 1.**
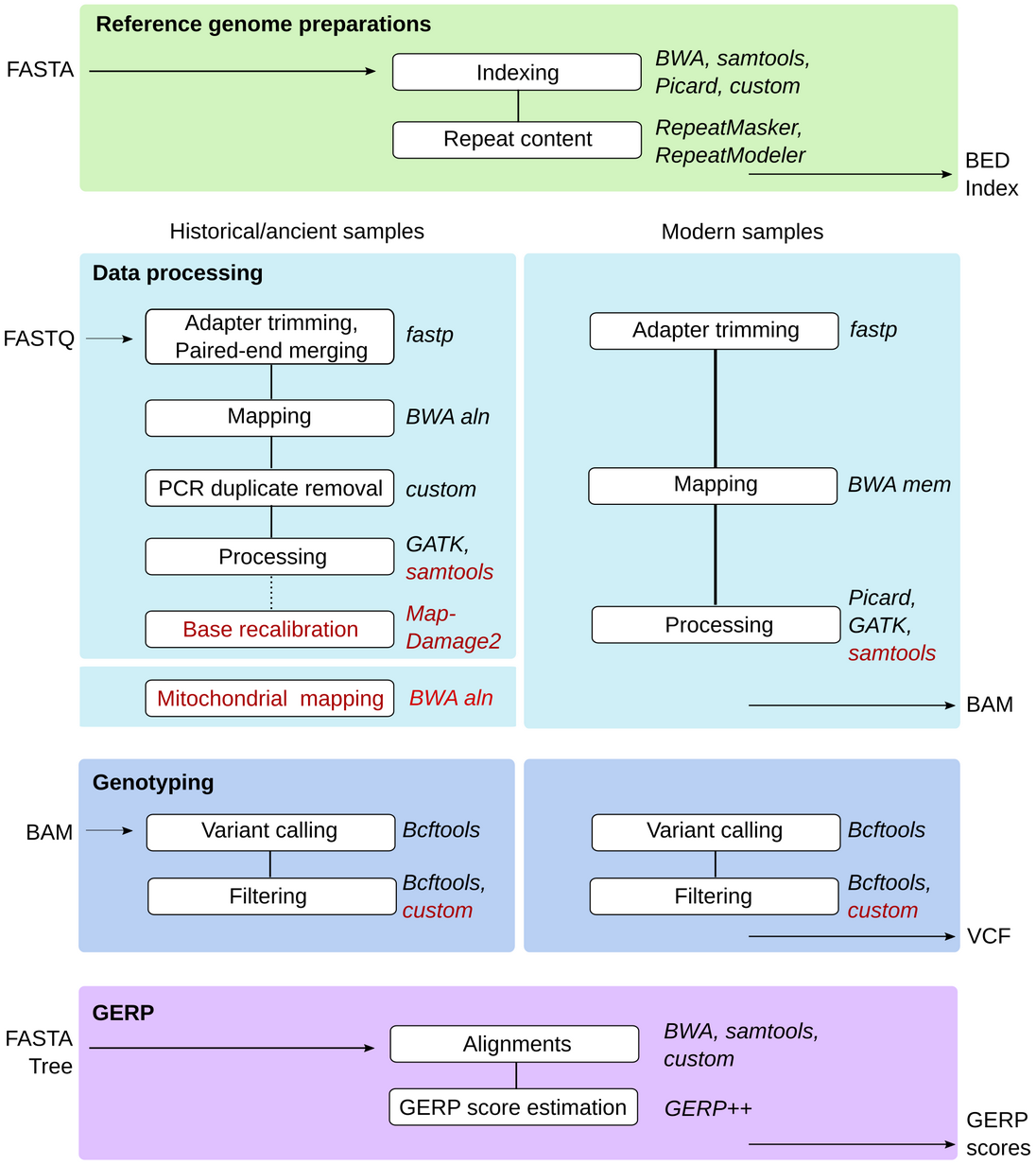
Overview of the GenErode pipeline data processing tracks. Input and output file formats, dependencies between steps, and main software used are shown. Optional steps are highlighted in red.

In short, GenErode takes paired-end FASTQ files as input, maps them to a reference genome assembly, performs variant calling, and runs several downstream analyses to estimate various genomic erosion indices. All data processing steps include quality checks and filtering. Optional steps include post-mortem damage base quality rescaling of historical/ancient samples, removal of CpG sites, subsampling a proportion of reads to achieve a similar average depth across samples, and mapping of historical/ancient data to human and other vertebrate mitochondrial genomes to assess the presence of non-endogenous reads in the data.

### Data processing track

Each data processing step is dependent on previous pipeline steps and is thus automatically run in sequence (see Figure 1).

#### 1. Reference genome assembly preparations

Prior to raw data processing, the reference genome assembly is prepared to generate files required for downstream analyses. This includes indexing with BWA [20] and samtools [21], generation of a FASTA dictionary using Picard (github.com/broadinstitute/picard) and of genome-coordinate BED files from the reference. At this stage, the pipeline also performs *de novo* identification and masking of repetitive regions of the genome using RepeatModeler [22] and RepeatMasker [23]. The identified repetitive regions will be excluded from all downstream steps of the data analysis track in order to avoid biases caused by mismapping.

#### 2. Data processing and mapping

Raw FASTQ files are trimmed with fastp [24], which automatically detects adapter sequences from the read data, under the assumption that only one adapter is present in the reads. Fastp also automatically enables poly-G trimming for Illumina NovaSeq or NextSeq samples by checking the flow cell identifier, so it handles read trimming for multiple Illumina platforms. For historical samples, fastp simultaneously adapter- and quality-trims reads and merges overlapping paired-end reads. By default, merged reads below a threshold of 30 bp are discarded. However, this threshold can be modified using the configuration file. Most ancient and historical DNA studies discard reads below a threshold of length between 30 and 35 bp, depending on the preservation of the samples (e.g. [25, 26]). The default merging settings are recommended in order to exclude modern-day contaminating sequences for which paired-end reads will not overlap since they are typically longer. Merged reads are then mapped to the reference genome using BWA aln with settings optimized for ancient/historical DNA (-l 16500 -n 0.01 -o 2) [27]. For modern samples, FASTQ files are adapter- and quality-trimmed with fastp and then mapped to the reference genome assembly with BWA mem [28] using default settings. For ancient/historical samples, multiple sequencing libraries are commonly generated per sample to avoid overrepresentation of specific PCR duplicates in the read data, using different index(es) for each sequencing library. The alignments of each index of a sample that were sequenced on different lanes are therefore merged to generate one BAM file per index. Next, PCR duplicates are identified and excluded from ancient/historical data using both read start and end mapping coordinates using a custom Python script. For modern samples, the same merging algorithm is run if applicable and PCR duplicates are identified and marked using Picard MarkDuplicates (broadinstitute.github.io/picard/). For both historical and modern data, the alignments of each sample from different indices are then merged to generate one BAM file per sample, and reads around indels are realigned with GATK IndelRealigner [29] to improve mapping accuracy. Basic mapping statistics are reported for each processing step using samtools and Qualimap [30], and are summarized using MultiQC [31]. Histograms depicting the depth per site as well as the genome-wide average depth and minimum and maximum depth thresholds for downstream analyses are also generated. The user can decide if genome-wide average depth should be calculated including or excluding sites with zero coverage. By default, depth thresholds are set to 1/3 and ten times the average genome-wide depth of each sample, which should be adjusted by the users according to their data characteristics. However, the pipeline uses an absolute minimum depth of three reads per site that cannot be changed by the user. Although the pipeline can be run with samples sequenced to lower depths, an average genome-wide depth of at least 6X per sample is recommended to have enough statistical power for detecting heterozygous sites. MultiQC reports and depth histograms based on BAM files after indel realignment are included into an automatically generated GenErode pipeline report.

There are two main optional steps to further process the BAM files before running any downstream data analyses: 1) base quality rescaling with MapDamage2 [32] for selected ancient/historical samples that have not been treated with the uracil-DNA glycosylase (UDG) enzyme and show post-mortem damage; and 2) subsampling of the selected BAM files from ancient/historical and/or modern samples using samtools to a target genome-wide depth to avoid biases introduced when comparing samples at different coverages. Additionally, a mitochondrial contamination check can be run for selected ancient/historical samples, in which the trimmed and merged reads are aligned to the mitogenomes of a set of species for which contamination may be present in the laboratory (i.e. from the laboratory reagents [33]). The user is asked to include a mitogenome FASTA file of the target species at this step. The GenErode pipeline produces BAM file statistics and a table listing the ratio of mapped reads to the mitochondrial genome of a potentially contaminating species and of the target species to help identify sequencing libraries with more reads mapping to the mitogenome from a different species. The results from this mitochondrial contamination check are not further used in the pipeline but the BAM files containing the sequences mapping to the mitochondrial genome of the target species are kept so that they can be used for downstream analyses outside of the pipeline. After each of these optional steps, basic mapping statistics are reported using samtools and Qualimap, and are summarized using MultiQC.

#### 3. Genotyping and variant filtering

This part of the pipeline is designed to perform variant calling on a per-sample basis. The rationale being that ancient, historical, and modern samples typically come from disparate time periods and locations, so using the information from some samples to predict variation in others can lead to undesired biases. Therefore, variants are called in each sample using bcftools [34] mpileup and call. Before proceeding to downstream analyses, the variant calls are subjected to several filtering steps. Methylated CpG sites are protected from UDG enzyme treatments and so post-mortem damage might remain in such sites [35, 36]. Ideally, a variant filter should therefore capture sites in ancient/historical samples that have a basepair change from CpG due to post-mortem damage. GenErode offers three optional methods to identify CpG sites: 1) identifying them in the reference genome assembly, 2) identifying them in selected samples once genotyped; and 3) combining both strategies. In all three cases, these CpG sites are automatically excluded from the final VCF files and downstream analyses. For datasets with only one or a few samples, we recommend identifying CpG sites in the reference genome assembly (option 1), which will remove damaged (or mutated) sites in the samples that are CpG in the reference. In larger datasets composed of modern and ancient/historical samples that are mapped to a more distantly related reference genome assembly (e.g. from a different species than the samples), it is recommended to identify CpG sites using all genotyped samples (option 2). Finally, we recommend a combination of both strategies (option 3) when a more stringent CpG filter is desired. After CpG site removal from VCF files, the pipeline moves on to other filtering steps. Only sites with mapping and base qualities of at least 30 are kept. The pipeline will exclude all variants that are located within 5 bp of indels (insertions or deletions), and will subsequently remove all indels. Also, all sites falling outside the depth thresholds specified by the user are removed (as described above). An allelic imbalance filter removes heterozygous sites in which less than 20% or more than 80% of reads support each allele to avoid erroneous genotypes caused by contamination or misalignments [37]. Variants falling within repetitive regions, earlier identified in the reference genome assembly, are also excluded in this step.

Finally, VCF files from all samples are merged and sites that are not biallelic as well as sites with more than 10% missing genotypes across all samples are removed. This missingness threshold can be adjusted by the user to suit each particular dataset, for example by relaxing it when many samples are to be analyzed at the same time. A BED file is created of all the genomic locations of sites remaining after the filtering and the merged VCF file is split into one VCF file of historical and one VCF file of modern samples.

### Data analysis track

All steps in the data analysis tracks are optional and can be run independently from each other once the data processing track is finished (see Figure 2).

**Figure 2.**
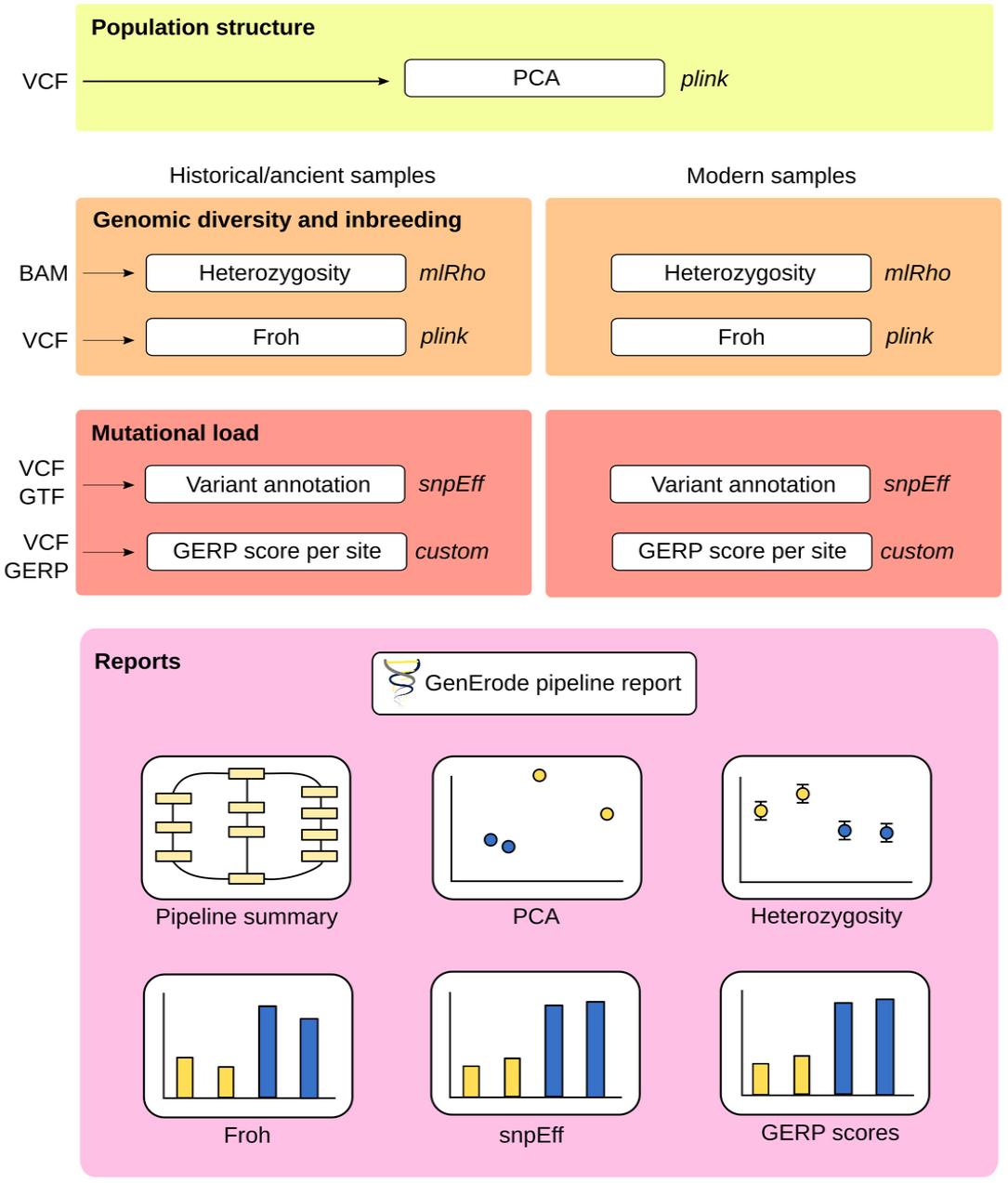
Overview of the GenErode pipeline data analysis tracks and final report. Input file formats and main software used are shown.

#### 4. Genome-wide diversity

GenErode uses mlRho [38] to estimate the maximum likelihood population mutation parameter (*θ* = 4*N_e_μ*), which under the infinite sites model approximates the per-site heterozygosity [39], from the final BAM file of each sample from step 2. All sites are filtered for mapping and base qualities as described above and for depth of coverage using the specified minimum and maximum thresholds. All sites within repetitive regions are excluded by default and CpG sites can be excluded if desired. GenErode also allows to estimate heterozygosity for autosomes and sex chromosomes separately if a list of known sex chromosomes (or scaffolds) is provided. BED files of repeat elements and/or CpG sites on autosomes and sex chromosomes are automatically generated using BEDtools [40]. A table and a figure summarizing *θ* estimates from mlRho is provided in the GenErode pipeline report.

#### 5. Inbreeding

Based on the VCF file of filtered genotypes from step 3, the GenErode pipeline uses the sliding window approach of ‘plink --homozyg’ [41] to identify runs of homozygosity (ROH). A wide array of plink settings such as ROH minimum size, maximum number of heterozygous sites per window, and maximum number of missing sites per window can be specified by the user. Long ROH (>= 2Mb) typically arise from recent mating with close relatives [42]. The fraction of the genome of each sample allocated in such ROH can be defined as the inbreeding coefficient (*F_ROH_*), which is estimated and visualized in a plot that is provided in the GenErode pipeline report.

#### 6. Population structure

The general population structure for the samples analyzed in the pipeline can be visualized with a Principal Component Analysis (PCA). Based on the VCF file of filtered genotypes from step 3, the pipeline generates PCAs for all samples combined and for historical and modern samples alone using plink. This analysis can be used to check genetic population structure, as well as to identify genetic outliers or biases among samples. All the PCA plots are made available in the GenErode pipeline report.

#### 7. Mutational load

GenErode offers two complementary methods to estimate mutational load, a proxy for genetic load [43], from the genomic data of the samples analyzed. For users intending to apply these two methods in endangered species for which the only remaining population is small or very small, we recommend running the GenErode pipeline using the reference genome assembly of a closely related species rather than the same species as the samples [43]. This is to avoid biases caused by a functional annotation that is based on an already genetically depauperate sample or from calling variants from data mapped to a reference genome assembly individual that is more genetically related to some of the samples under comparison (i.e., more related to the modern samples than to the historical ones).

The first method requires a genome annotation in GTF2 format based on which GenErode annotates individual variants from step 3 using snpEff [44]. snpEff allows to annotate all identified variants per sample and to estimate a simple prediction of their functional effects which can be compared among samples or groups of samples (i.e., historical vs modern). MultiQC reports summarizing snpEff results per dataset are provided in the GenErode pipeline report. Numbers of variants of different impact categories per sample (including loss-of-function (LOF) variants in the high impact category) as estimated by snpEff are visualized in a plot that is also available in the GenErode pipeline report.

The second method uses GERP scores [45] to infer the relative mutational load of each sample from the number of derived alleles in evolutionary constrained regions of the genome based on a panel of outgroup species [46]. This method does not require a functional annotation for the target species’ reference genome assembly, but it does require at least 30 genome assemblies from different species to act as outgroups [46], and a dated phylogeny from all the included species in NEWICK format and in billions of years (which can be obtained from, e.g. timetree.org). Our pipeline first splits each outgroup genome assembly into non-overlapping 35 bp sequences and maps them to the target species reference genome assembly to identify highly conserved regions of the genome [46]. Next, it estimates a per-site GERP score using GERP++ [47]. Finally, the ancestral and derived states of each variant from step 3 are inferred and used to estimate a relative mutational load. For each sample, heterozygous and homozygous sites are counted as contributing to one and two derived alleles, respectively. Only derived sites with a GERP score within a minimum and a maximum threshold set by the user are considered for relative load mutational calculations. Our approach is not suitable to identify fast evolving regions and positive GERP scores indicate evolutionary constraint, so a suitable minimum GERP score of at least zero should be chosen. A script to approximate a minimum GERP score based on a desired percentile of GERP scores from the genome from histogram bins is available in the GenErode GitHub repository (‘utilities/get_gerp_score_percentile.py’). A histogram of GERP scores across the genome is available in the GenErode pipeline report and can be used as guidance. At each of these sites, the GERP score is multiplied by the number of derived alleles per sample. The sum of these GERP scores is then normalized by the number of derived alleles at these sites (modified from [46]), to obtain a relative mutational load estimation for each sample. Finally, the pipeline generates a table and a figure with the relative mutational load estimates across samples that are included in the GenErode pipeline report.

### Test dataset

To help the user get familiarized with our pipeline we provide a test dataset based on the Sumatran rhinoceros (*Dicerorhinus sumatrensis*) re-sequencing data from von Seth et al. [43]. The test dataset is composed of re-sequencing data from three historical and three modern samples from the now-extinct Malay Peninsula population (Additional file 1, Table S1), which had been analyzed for patterns of genome erosion. These data correspond to those sequences that mapped to a single scaffold of 41 Mb size (‘Sc9M7eS_2_HRSCAF_41′) from the Sumatran rhinoceros genome assembly (GenBank nr. GCA_014189135.1), as well as to the mitochondrial genome (GenBank nr. NC_012684.1), which are both provided as references. A small proportion of sequences mapping elsewhere in the genome were included, too. We have also included three scaffolds from the White rhinoceros genome assembly (*Ceratotherium simum simum*; GenBank accession number GCF_000283155.1) for analyses that require a reference from a more distantly related species (‘NW_004454182.1′, ‘NW_004454248.1′, and ‘NW_004454260.1′). These scaffolds had been identified as being putatively orthologous to the Sumatran rhinoceros scaffold ‘Sc9M7eS_2_HRSCAF_41′. For mutational load analyses, gene predictions from the three White rhinoceros scaffolds were extracted in GTF format and scaffolds from the genome assemblies of 30 mammalian outgroup species (Additional file 1, table S2) were identified as putative orthologs to the Sumatran rhinoceros scaffold. A phylogeny of the White rhinoceros and the 30 outgroup species including divergence time estimates (in billions of years) was obtained from timetree.org in NEWICK format (Additional file 1, figure S1). A detailed description of the test dataset generation is provided in Additional file 1, Extended methods note 1. The test dataset is publicly available on the SciLifeLab Data Repository (DOI: 10.17044/scilifelab.19248172). The results presented in Figure 3 have been generated using the GenErode pipeline for this dataset on a high-performance compute (HPC) cluster using a slurm workload manager. The final reports generated by the GenErode pipeline are provided as Additional file 2 and Additional file 3, which include configuration files specifying all parameter choices.

**Figure 3.**
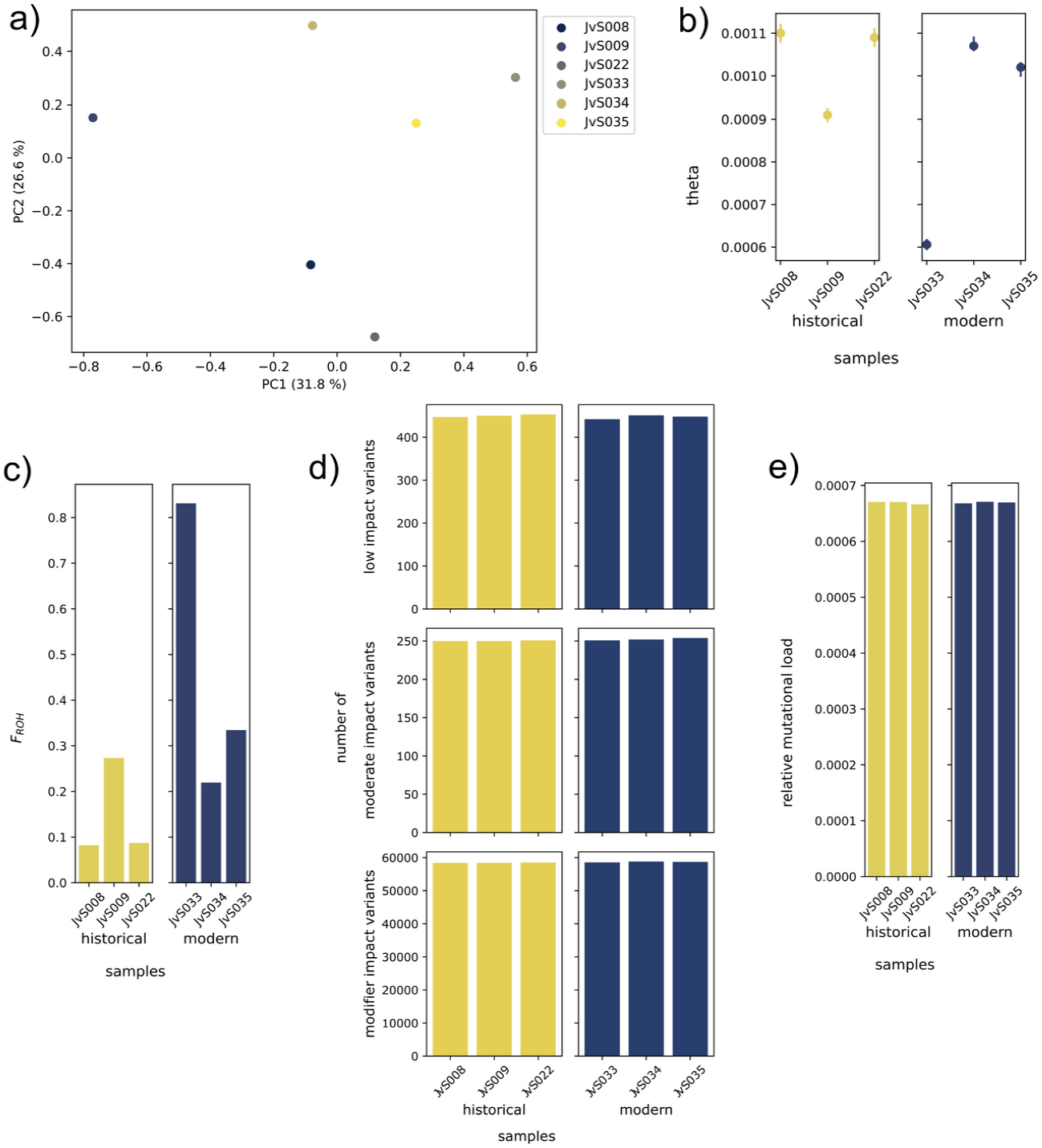
Results from the downstream analyses for historical and modern Sumatran rhinoceros samples from the test dataset. a) Principal component analysis (PCA) of historical and modern Sumatran rhinoceros samples using the Sumatran rhinoceros reference. Historical samples: JvS008, JvS009, JvS022. Modern samples: JvS033, JvS034, JvS035. See “PCA” in Additional file 3. b) Maximum likelihood estimates of genome-wide heterozygosity (*θ)* and 95% confidence intervals from mlRho in historical and modern Sumatran rhinoceros samples using the Sumatran rhinoceros reference. See ‘mlRho ‘mlRho’ in Additional file 4. c) Inbreeding coefficient estimated as the proportion of the genome in runs of homozygosity (*F_ROH_*) for ROH of length >= 2 Mb in historical and modern Sumatran rhinoceros samples. Parameter settings in the configuration file: homozyg-snp: 25; homozyg-kb: 100; homozyg-window-snp: 250; homozyg-window-het: 3; homozyg-window-missing: 15; homozyg-het: 750. See “ROH” in Additional file 3. d) Number of variants in each impact category in historical and modern Sumatran rhinoceros samples as estimated by snpEff and using the White rhinoceros reference with 253 protein-coding gene annotations. Note that none of the samples contained any high impact variants which are therefore not shown. See “snpEff” in Additional file 2. e) Relative mutational load estimated from GERP scores for historical and modern Sumatran rhinoceros samples using the White rhinoceros reference. GERP scores were estimated using 30 outgroup species from across the tree of mammals. Only derived alleles from sites with GERP scores above the 99th percentile were included (corresponding to GERP > 0.00048928). See “GERP” in Additional file 2.

## Results and discussion

### Requirements and pipeline configuration

For a species with a genome size of around 3 Gbp the GenErode pipeline requires a Linux system with at least 16 cores, each with a minimum of 6 GB of RAM (a total of 96 GB), and internet access so Snakemake can download the required containers on the fly. Depending on the number of samples to analyze and the sequencing effort of each sample the storage space required could be large. For example, in a test run with four Sumatran rhinoceros samples with an average genome-wide depth ranging from 9 to 18X and mapped to the Sumatran rhinoceros genome assembly, the pipeline directory size after running all data processing and downstream analyses (except the storage-space demanding GERP step) had a final size of ca. 750 GB. However, this was after GenErode had removed temporary intermediate files and the space required reached several TB at some points during this pipeline run, so storage space flexibility is recommended. The only software requirements to run the pipeline are Singularity (at least v3.6.1) and Conda (tested with Miniconda3), which have to be installed on the system. The GenErode pipeline is installed by cloning the GitHub repository into the directory in which it will be run. The user needs to create a Conda environment from the ‘environment.yml’ file provided in the GitHub repository, which contains Snakemake (version 6.12.1 [19]), Python3, and all required additional python modules. All custom python scripts used by the pipeline are included in the GitHub repository and all additional software and dependencies are automatically downloaded as Docker images and run as Singularity containers by Snakemake, so there is no need for any additional installations by the user. All output files of the pipeline will be automatically written to subdirectories within the GenErode directory, except for any files related to the reference genome assembly and the genome annotation, which will be written in the same directory as the reference genome assembly.

To configure a pipeline run, the user first needs to create a metadata file in which the sample names, sequencing runs, read group IDs, and paths to the raw sequencing files are specified. Examples are available in the ‘config’ directory of the GitHub repository and in the SciLifeLab Data Repository containing the test dataset (see below). This pipeline has been developed using the slurm workload manager [48] with the official Snakemake profile for slurm (github.com/Snakemake-Profiles/). The computational resources for the different jobs that are submitted to the cluster via Snakemake are specified in a cluster configuration file (‘config/cluster.yaml’) that needs to be edited according to the user’s system. Finally, to configure each run of the pipeline the user needs to edit the pipeline configuration file (‘config/config.yaml’) in which each step of the pipeline can be set to ‘True’ or ‘False’. Additionally, the user can specify any required parameters for the desired steps to run. For example, the adapter trimming step for historical samples requires the user to specify the minimum read length allowed after trimming and read merging (default 30 bp). When a step is set to ‘True’, all analyses within this step (‘rules’) are automatically run along with any upstream rules they depend on. For example, if the mapping step is set to ‘True’, adapter trimming and quality controls of raw and trimmed reads are also automatically run in case these previous rules have not been run before.

A complete guide on the requirements and configuration of the pipeline, including detailed explanations for each step and the parameters, is available in the wiki section of the pipeline’s GitHub page (github.com/NBISweden/GenErode/wiki/).

### Test dataset: genomic erosion in the Sumatran rhinoceros

We analyzed the test dataset of re-sequencing data from three historical and three modern Sumatran rhinoceros samples from the now-extinct Malay Peninsula population (Additional file 1, table S1; see “Implementation: Test dataset” and Additional file 1, Extended methods note 1). The GenErode pipeline was run once using the Sumatran rhinoceros scaffold ‘Sc9M7eS_2_HRSCAF_41′ as reference to obtain PCAs, *θ* and *F_ROH_* estimates, and a second time using the putatively orthologous White rhinoceros scaffolds ‘NW_004454182.1′, ‘NW_004454248.1′, and ‘NW_004454260.1′ as reference to infer PCAs and mutational load using snpEff annotations and GERP scores. The Sumatran rhinoceros scaffold consists of 40.8 Mbp of which 27.7% were masked by RepeatMasker. The White rhinoceros scaffolds have a combined length of 41.2 Mbp of which 27.1% were repeat masked. Three to 24 sequencing libraries were available per historical sample; each sample had been sequenced on two Illumina HiSeqX lanes. For each modern sample, one sequencing library was included. An average number of ca. 4.6 M paired-end reads were included into the test dataset per historical sample, and ca. 3.1 M paired-end reads were included per modern sample. The optional GenErode step to test for overrepresentation of mitochondrial reads in historical libraries revealed ratios of reads mapping to the included mitochondrial genomes (see Additional file 1, table S3) and the Sumatran rhinoceros mitochondrial genome ranging from 0.0 to 0.14 (mean=0.052, sd=0.037). This suggests that the presence of non-endogenous reads in the historical libraries of Sumatran rhinoceros is low or absent.

For the historical samples, after filtering per index and sample and PCR duplicate removal, an average of 2.7 M and 2.2 M reads mapped to the Sumatran and White rhinoceros references, respectively. Average genome-wide depth ranged from 4 to 6 for the Sumatran rhinoceros reference and 4 to 5 for the White rhinoceros reference (Additional file 1, figure S2A). For the modern samples, after filtering and PCR duplicate marking, an average of 3.0 M and 2.9 M paired-end reads mapped to the Sumatran and White rhinoceros references, respectively. This corresponds to average genome-wide depths ranging from 16 to 20 across all samples and references (Additional file 1, figure S2B). Prior to downstream data analysis steps, BAM files from modern samples were subsampled to an average depth of 6. In total, 397,642 and 395,780 CpG sites were identified in the Sumatran and White rhinoceros references, respectively. These sites were removed from individual VCF files followed by subsequent filtering using default settings of the pipeline. The filtered and merged multi-sample VCF files contained 7,484 SNPs for the Sumatran rhinoceros reference and 40,352 SNPs for the White rhinoceros reference. This difference can be explained with a large number of sites where the Sumatran rhinoceros samples are homozygous for the ALT allele, i.e. they are diverged from the White rhinoceros reference.

In a principal component analysis (PCA) of Sumatran rhinoceros samples using the VCF file based on the Sumatran rhinoceros reference, the first principal component (PC1) explained 31.8% of the variation (Figure 3A). The three modern samples clustered slightly closer together than the three historical samples. A PCA using the VCF file based on the White rhinoceros reference resulted in a highly similar plot (Additional file 1, figure S3). GenErode was next run to estimate various genomic erosion indices to compare them between historical and modern samples. Genome-wide heterozygosity was estimated as *θ* in mlRho using BAM files based on the Sumatran rhinoceros reference with default filtering settings and excluding CpG sites. Although heterozygosity for the test scaffold was lower than for the whole genome [43], historical and modern samples were similar (Figure 3B), consistent with genome-wide temporal comparisons from [41]. ROH were estimated using the VCF file based on the Sumatran rhinoceros reference (see “ROH” and “Configuration” in Additional file 3 for parameter choices). Larger proportions of the reference were in ROH >= 2 Mb in modern samples than in historical samples (Figure 3C), suggesting stronger effects of inbreeding in the modern population, consistent with von Seth et al. [43]. No variants of potentially high impact on the protein (which include LOF variants) were found in any of the samples in the snpEff analyses of the VCF files based on the White rhinoceros reference and 253 gene predictions. Numbers of variants in the other impact categories were similar across historical and modern samples (Figure 3D). Unlike genome-wide estimates [43], relative mutational load estimated from the top 1% of GERP scores based on the White rhinoceros reference and 30 outgroup species (Additional file 1, table S2, figure S1) were similar between historical and modern samples for our test dataset (Figure 3E; see Additional file 1, figure S4 for a histogram of GERP scores).

The expected runtime of all jobs run by GenErode for this test dataset on a HPC cluster with 16 cores (6 GB of RAM per core) can be up to ca. 15 core hours for data processing and downstream analyses using the Sumatran rhinoceros reference, and ca. 47 core hours for data processing and downstream analyses using the White rhinoceros reference.

### A bioinformatics pipeline for ancient/historical, and modern data

There are established bioinformatic pipelines tailored to processing and genotyping ancient DNA data (e.g. [49]) as well as mining metagenomic information from it (e.g. [50]). However, to our knowledge, GenErode is the first pipeline specifically designed to process ancient, historical and modern sequencing data from the same species with the aim to make them comparable. This is because GenErode has separately optimized tracks for processing of ancient/historical and modern data, enabling a better comparison between samples from different time periods. Moreover, in its simplest configuration, GenErode requires only a reference genome assembly and whole-genome re-sequencing data. It is therefore highly suitable for analyses of data from non-model species that often lack genomic resources, which is often the case for endangered taxa.

Our pipeline also outputs the results of a number of downstream analyses including estimates of genomic diversity indices, such as heterozygosity and inbreeding, as well as mutational load (including per-sample LOF estimates), that can be used to quantify genomic erosion through time. These genomic erosion indices have been proposed to be relevant in assessing the species risk of extinction when compared among time periods, such as historical versus modern samples (e.g., [15, 43, 51, 52]). It should be noted that GenErode offers only one estimation method for each data analysis track and genomic erosion index. While this can be sufficient to draw conclusions for some studies, for users that desire to conduct a more in-depth analysis of their data, the output files produced (i.e. BAM and VCF files) can be easily used as input for other types of analyses outside of the pipeline.

One important feature of GenErode is that it is a highly modular and flexible pipeline. It allows analyzing the data in a stepwise fashion with important options pre-set for the user but also plenty of customizable parameters and several optional steps so the user can adapt it to their particular dataset. Even though the genotyping and data analysis tracks are specifically designed for samples sequenced at a minimum average genome-wide depth of 6X, the data processing tracks (read processing, mapping, and BAM file filtering) can be used for samples of all depths. This means that the pipeline can also process low or very low coverage samples, either UDG treated or not, typical for ancient DNA projects or museum collections with highly degraded specimens, and at the same time provide tools for authentication such as post-mortem damage profiles.

Finally, alongside a comprehensive variety of output result files, including an automatically generated pipeline summary report, GenErode stores an organized set of parameter files and output logs for each step, providing the user with all information required to thoroughly document their analysis. Also, the majority of the pipelines’ analyses are run in separate, automatically executed Singularity containers, which enables the user to process and analyze sequencing data in a highly reproducible way [53]. Reproducibility still remains a challenge for most bioinformatic applications and analyses from published articles across the sciences [17], but is key to obtaining comparable genomic erosion measurements. This is necessary to integrate them into the criteria to establish threat categories in endangered species’ lists [8].

## Conclusions

We introduce GenErode, a bioinformatic pipeline to investigate genomic erosion that includes state-of-the-art tools for ancient/historical, and modern DNA data within an integrated, easily reproducible and accessible format. Two of the main challenges to close the gap between evolutionary research and conservation applications are the lack of standard methods for the measurement of genomic diversity, as well as the need for training in the bioinformatic methods required to analyze genomic data. GenErode aims to produce comparable estimates of genomic diversity indices from temporally sampled datasets that can be used to quantify genomic erosion through time. Additionally, setting up and running GenErode requires no programming knowledge and all the bioinformatic steps are well documented, making it easy to run for people with different backgrounds. Therefore, with this pipeline we aim to make the bioinformatic methods required to process ancient/historical, and modern genomic data to estimate diversity changes more streamlined and accessible to a wider conservation community, including museum curators as well as conservation researchers and practitioners.

## Availability and requirements

Project name: GenErode

Project home page: github.com/NBISweden/GenErode/

Archived version: 0.4.1

Operating system(s): Linux

Programming language: Snakemake, Python3

Other requirements: Miniconda or Anaconda, Singularity

License: GPL-3.0

Any restrictions to use by non-academics: None

GenErode is an open-source software and is available on GitHub (github.com/NBISweden/GenErode/), including extensive documentation. GenErode depends on Snakemake, the Conda package manager and Singularity. Conda can be downloaded as part of Anaconda or Miniconda (Python 3.7), and Singularity has to be installed. A Conda environment containing pipeline-wide dependencies (incl. Snakemake) is available on GitHub (‘environment.yml’). All other dependencies will be automatically handled by Snakemake using Singularity when running the pipeline.

## Availability of data and materials

The Sumatran rhinoceros test dataset can be downloaded from the SciLifeLab Data Repository (DOI: 10.17044/scilifelab.19248172) and is suitable for testing the GenErode pipeline. A minimum of 2 cores with at least 6 GB each is sufficient for each of the pipeline jobs to run with this test dataset, so a HPC system is not strictly mandatory. However, the execution of at least 16 jobs in parallel using the cluster configuration settings in ‘config/cluster.yaml’ from the GitHub repository is highly recommended to reduce the overall runtime.

For test runs of the pipeline with limited computational resources, a minimal test dataset is available in the ‘.test/data’ folder of the GitHub repository. This data was deposited there for continuous integration of the pipeline with GitHub actions. Note that the results from this dataset are not biologically meaningful due to the reduced size of the dataset.

## Additional files

**Additional file 1**: Supplementary Information, which contains Extended methods note 1, Tables S1-S3 and Figures S1-S4.

**Additional file 2**: GenErode pipeline report for analyses of the test dataset based on the White rhinoceros reference.

**Additional file 3**: GenErode pipeline report for analyses of the test dataset based on the Sumatran rhinoceros reference.

## Author contributions

D.D.d.M., V.E.K., N.D., J.v.S., B.N., and L.D. conceived the idea. D.D.d.M. performed conceptual work. V.E.K. designed and developed the pipeline. M.K. and T.v.V. developed parts of the pipeline. D.D.d.M., T.v.V., N.D., D.W.G.S., J.v.S., E.L., M.D., B.N., and L.D. provided critical input for the pipeline development. V.E.K., N.D., J.v.S., E.L., M.D., and P.E. tested the pipeline. V.E.K. generated the test dataset and analyzed it using the pipeline. D.D.d.M. and V.E.K. wrote the manuscript. All authors read and approved the final manuscript.

## Acknowledgements

We would like to express our thanks to Pontus Skoglund for providing us with his script ‘samremovedup.py’, Jonas Söderberg for the GenErode pipeline logo design, Camilo Chacón-Duque, Petter Larsson, Heloise Muller, Ioana Meleg, Jovanka Studerus for testing the pipeline, and Johan Nylander, Marcel Martin, Thijessen Naidoo, Winfried Kretzschmar, Fredrik Boulund, John Sundh, and Per Unneberg for helpful discussions on the pipeline development.

