## Additional file 1 for "GenErode: a bioinformatics pipeline to investigate genome erosion in endangered and extinct species"

### Additional file 1: Supplementary Information

**Table of contents**

Extended methods note 1 page 2

Supplementary tables page 5

Supplementary figures page 8

**<u>Extended methods note 1</u>**

**Detailed description of test dataset generation**

We provide a test dataset based on the Sumatran rhinoceros re-sequencing data from von Seth et al. [1] so that users have the possibility to get familiarized with the GenErode pipeline. The results presented in this article were obtained from analyzing this test dataset with the GenErode pipeline. The test dataset is available at the Scilifelab Data Repository (DOI: 10.17044/scilifelab.19248172) and all scripts used to generate the test dataset are available in the GenErode GitHub repository (github.com/NBISweden/GenErode/) in the directory 'docs/extras/test_dataset_generation' along with workflow descriptions.

<u>Reference genomes</u>

We extracted scaffold 'Sc9M7eS_2_HRSCAF_41' of size 40,842,778 bp from the Sumatran rhinoceros genome assembly (*Dicerorhinus sumatrensis harrissoni*; GenBank accession number GCA_014189135.1) to be used as reference genome for mlRho and ROH analyses. Some of the GenErode steps in the data analysis track require the reference genome of a closely related species to avoid biases from a functional annotation of an already genetically depauperate sample or from calling variants from data mapped to a reference genome individual that is closer related to some of the samples than to others in the dataset. We therefore identified scaffolds from the White rhinoceros genome assembly (*Ceratotherium simum simum*; GenBank accession number GCF_000283155.1) as being putatively orthologs to the Sumatran rhinoceros scaffold through a reciprocal blast and a mapping approach, to be used as reference genome for snpEff and GERP analyses.
For the mapping approach, the Sumatran rhinoceros scaffold 'Sc9M7eS_2_HRSCAF_41' was split into 35bp reads using the script 'fa2fq.py' from the GenErode pipeline. These reads were mapped to the White rhinoceros genome assembly using bwa mem version 0.7.17 [2] with default settings and samtools version 1.9 [3], keeping only uniquely mapped reads. Qualimap version 2.2.1 [4] was run to obtain BAM file statistics per scaffold. This approach identified three White rhinoceros scaffolds as putatively orthologous to the Sumatran rhinoceros scaffold: 'NW_004454182.1' with 15,206,520 mapped bases, 'NW_004454248.1' with 3,517,500 mapped bases, and 'NW_004454260.1' with 2,776,935 mapped bases. For comparison, the maximum number of mapped bases for any other scaffold was 48,580. The three scaffolds combined are 41,195,616 bp long. The same three scaffolds were identified with a reciprocal blast approach. First, protein sequences of genes located on the Sumatran rhinoceros scaffold 'Sc9M7eS_2_HRSCAF_41' were extracted from the genome-wide set of proteins available from the Sumatran rhinoceros genome annotation [5] and were blasted to the White rhinoceros genome assembly using tblastn (blast version 2.11.0+;[6]) with e-value 10e-5. The top hit per protein was identified and its sequence was extracted from the White rhinoceros genome

assembly in fasta format using custom code and BEDtools getfasta version 2.29.2 [7]. These White rhinoceros sequences were blasted to the genome-wide set of Sumatran rhinoceros protein sequences using blastx (e-value 10e-5). The top hit per White rhinoceros sequence was identified and only those top hits that had blasted to their original Sumatran rhinoceros protein sequence were kept. Finally, the numbers of reciprocal blast hits per scaffold were compared and the three White rhinoceros scaffolds ‘NW_004454182.1’, ‘NW_004454248.1’, and ‘NW_004454260.1’ were identified as the scaffolds with the largest numbers of reciprocal blast hits (114, 17, and 31 hits, respectively).
GenErode also requires a reference for the comparative mapping of historical samples to mitochondrial genomes (see Table S3), for which we provide the Sumatran rhinoceros mitochondrial genome ‘NC_012684.1’ in the test dataset.

Re-sequencing data

We extracted a subset of reads from re-sequencing data from six Sumatran rhinoceros from the now-extinct Malay Peninsula population to be included into the test dataset. The three historical and three modern samples (Table S1) had been sequenced and analyzed for patterns of genome erosion by von Seth et al. [1]. For two of the historical Sumatran rhinoceros samples, three sequencing libraries are available per sample that had been sequenced on two lanes each. For the third historical sample, 24 sequencing libraries are available that had been sequenced on two lanes (12 libraries per lane). Sequencing libraries of historical samples contain 21,775,075 to 98,421,390 paired-end reads. For each of the three modern Sumatran rhinoceros samples, one sequencing library is available, containing 173,233,214 to 221,534,836 paired-end reads. Reads were included into the test dataset that mapped to the Sumatran rhinoceros scaffold ‘Sc9M7eS_2_HRSCAF_41’, along with a small proportion of randomly selected reads that mapped to the Sumatran rhinoceros mitochondrial genome or elsewhere in the genome. Briefly, raw paired-end reads from each library were mapped to the full Sumatran rhinoceros genome assembly using bwa mem with default settings. The scaffold ‘Sc9M7eS_2_HRSCAF_41’ was extracted from the BAM files using samtools view and the BAM files were converted to fastq format using samtools collate and samtools fastq, removing secondary alignments. The same approach was applied to the Sumatran rhinoceros mitochondrial genome (GenBank accession number NC_012684.1) as reference. From the resulting fastq files of mitochondrial reads, 250, 1,000, and 5,000 reads (low coverage historical libraries, high coverage historical libraries, modern samples) were randomly extracted, respectively, using seqtk sample version 1.2-r101 (github.com/lh3/seqtk) and the seed ‘-s100’. Finally, 2,000, 10,000, and 42,000 reads (low coverage historical libraries, high coverage historical libraries, and modern samples) were randomly selected from the genome-wide, raw fastq files, respectively, using seqtk sample with the seed ‘-s999’. For each forward or reverse library, the fastq files for reads mapping to ‘Sc9M7eS_2_HRSCAF_41’, randomly selected reads mapping to the mitochondrial genome, and randomly selected reads from the entire genome were concatenated. In some libraries, randomly selected reads from the entire genome were duplicates of reads mapping to ‘Sc9M7eS_2_HRSCAF_41’. We identified the read names of unique reads in the concatenated fastq files using custom code and extracted them using seqtk subseq. For historical samples, this approach resulted in 86,373 to 974,743 paired-end reads per sequencing library, which sums up to a total number of 4,049,944, 4,888,241 and 4,878,523 paired-end reads per sample, respectively. We identified and extracted 2,735,683 to 3,570,154 paired-end reads per modern sample. These reads were included into the test dataset in fastq format.

<u>Mutational load analyses</u>

Gene predictions from the three White rhinoceros scaffolds were extracted from the genome-wide annotation and are included in the test dataset in GTF format to be used in snpEff analyses.

GERP++ analyses require the genome assemblies from at least 30 outgroup species [8] and a phylogeny with divergence time estimates to identify highly conserved regions in the reference. Scaffolds from the genome assemblies of 30 mammalian species (Table S2) were identified as putative orthologs to the Sumatran rhinoceros scaffold ‘Sc9M7eS_2_HRSCAF_41’ via reciprocal blast, using the same approach as described above. The scaffolds are included into the test dataset in fasta format to be used as outgroup sequences for the GERP analysis. A phylogeny of White rhinoceros and the 30 mammalian species with divergence time estimates (in billions of years) was obtained from timetree.org in NEWICK format (Figure S1). For compatibility with the GenErode pipeline, species names in the tree were changed to match their respective fasta file names and the edited NEWICK file was added to the test dataset. A script to approximate the 99th percentile of GERP scores to be used as minimum GERP is publicly available (github.com/NBISweden/GenErode/utilities/get_gerp_score_percentile.py).

#### Supplementary tables

**Table S1. Test dataset sample information.** Sumatran rhinoceros samples from the now-extinct Malay Peninsula peninsula population were subsampled to represent Sumatran rhinoceros scaffold ‘Sc9M7eS_2_HRSCAF_41’ and the mitochondrial genome to be used as test dataset.

| Sample ID (Test dataset) | Sample ID (von Seth et al. [1]) | Dataset | SRA identifier |
|---|---|---|---|
| JvS008 | SR08 | historical | ERS4044060 |
| JvS009 | SR09 | historical | ERS4044061 |
| JvS022 | SR22 | historical | ERS4044063 |
| JvS033 | KB6196 | modern | ERS4042484 |
| JvS034 | KB6197 | modern | ERS4042485 |
| JvS035 | KB6198 | modern | ERS4042486 |

**Table S2. GenBank accession numbers of genome assemblies from 30 mammalian species.** Putatively orthologous scaffolds to Sumatran rhinoceros scaffold 'Sc9M7eS_2_HRSCAF_41' were extracted from each genome assembly and included into the test dataset to be used in GERP analyses.

| **Species name** | **GenBank accession number** |
|---|---|
| *Ailurus fulgens* | GCA_002007465.1 |
| *Antilocapra americana* | GCA_007570785.1 |
| *Balaenoptera acutorostrata* | GCF_000493695.1 |
| *Bubalus bubalis* | GCF_003121395.1 |
| *Camelus dromedarius* | GCF_000803125.2 |
| *Canis lupus* | GCF_014441545.1 |
| *Catagonus wagneri* | GCA_004024745.2 |
| *Cervus elaphus* | GCA_002197005.1 |
| *Diceros bicornis* | GCA_013634535.1 |
| *Enhydra lutris* | GCF_002288905.1 |
| *Equus asinus* | GCA_016077325.1 |
| *Giraffa camelopardalis* | GCA_017591445.1 |
| *Hippopotamus amphibius* | GCA_004027065.2 |
| *Hyaena hyaena* | GCF_003009895.1 |
| *Leptonychotes weddellii* | GCF_000349705.1 |
| *Lipotes vexillifer* | GCF_000442215.1 |
| *Manis javanica* | GCF_014570535.1 |
| *Mesoplodon bidens* | GCA_004027085.1 |
| *Ovis aries* | GCF_002742125.1 |
| *Panthera leo* | GCA_008795835.1 |
| *Paradoxurus hermaphroditus* | GCA_004024585.1 |
| *Physeter catodon* | GCF_002837175.2 |
| *Procyon lotor* | GCA_015708975.1 |
| *Spilogale gracilis* | GCA_004023965.1 |
| *Suricata suricatta* | GCF_006229205.1 |
| *Tapirus indicus* | GCA_004024905.1 |
| *Tragulus javanicus* | GCA_004024965.2 |
| *Tursiops truncatus* | GCF_011762595.1 |
| *Ursus maritimus* | GCF_017311325.1 |
| *Zalophus californianus* | GCF_009762305.2 |

**Table S3. Mitochondrial genomes provided with the GenErode pipeline for comparative mapping analysis.** Available in the GitHub repository (github.com/NBISweden/GenErode/) along with the pipeline (‘data/mitogenomes’ folder).

| **Species** | **GenBank accession number** |
|---|---|
| *Gallus gallus* (chicken) | NC_001323.1 |
| *Bos taurus* (cow) | NC_006853.1 |
| *Homo sapiens* (human) | NC_012920.1 |
| *Mus musculus* (mouse) | NC_005089.1 |
| *Sus scrofa* (pig) | NC_000845.1 |

#### Supplementary figures

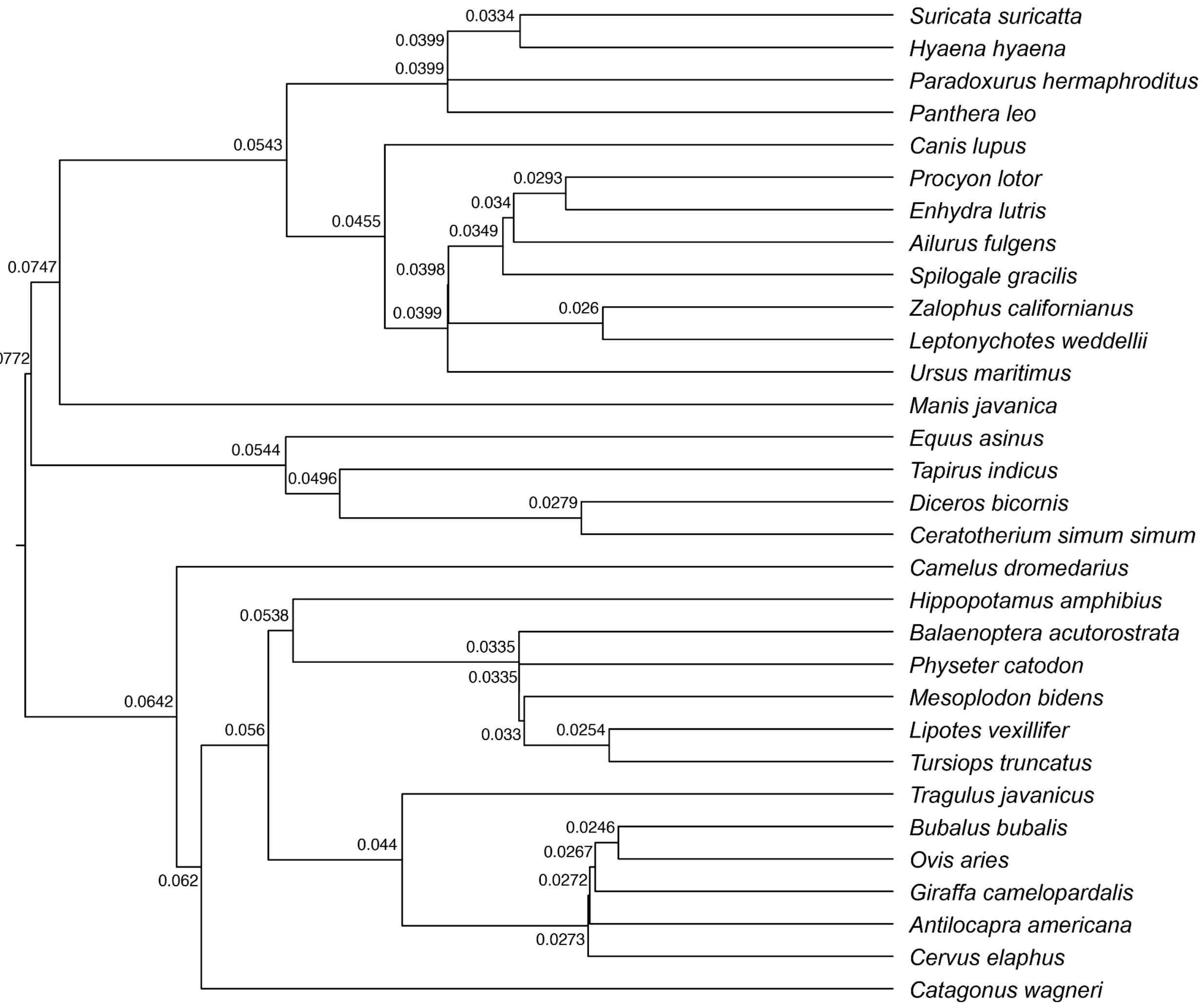


**Figure S1. Phylogenetic tree of the White rhinoceros (*Ceratotherium simum simum*) and 30 mammalian outgroup species (obtained from timetree.org).** Node labels represent divergence time estimates in billions of years. This tree is part of the test dataset in NEWICK format to be used in GERP analyses.

**A)**

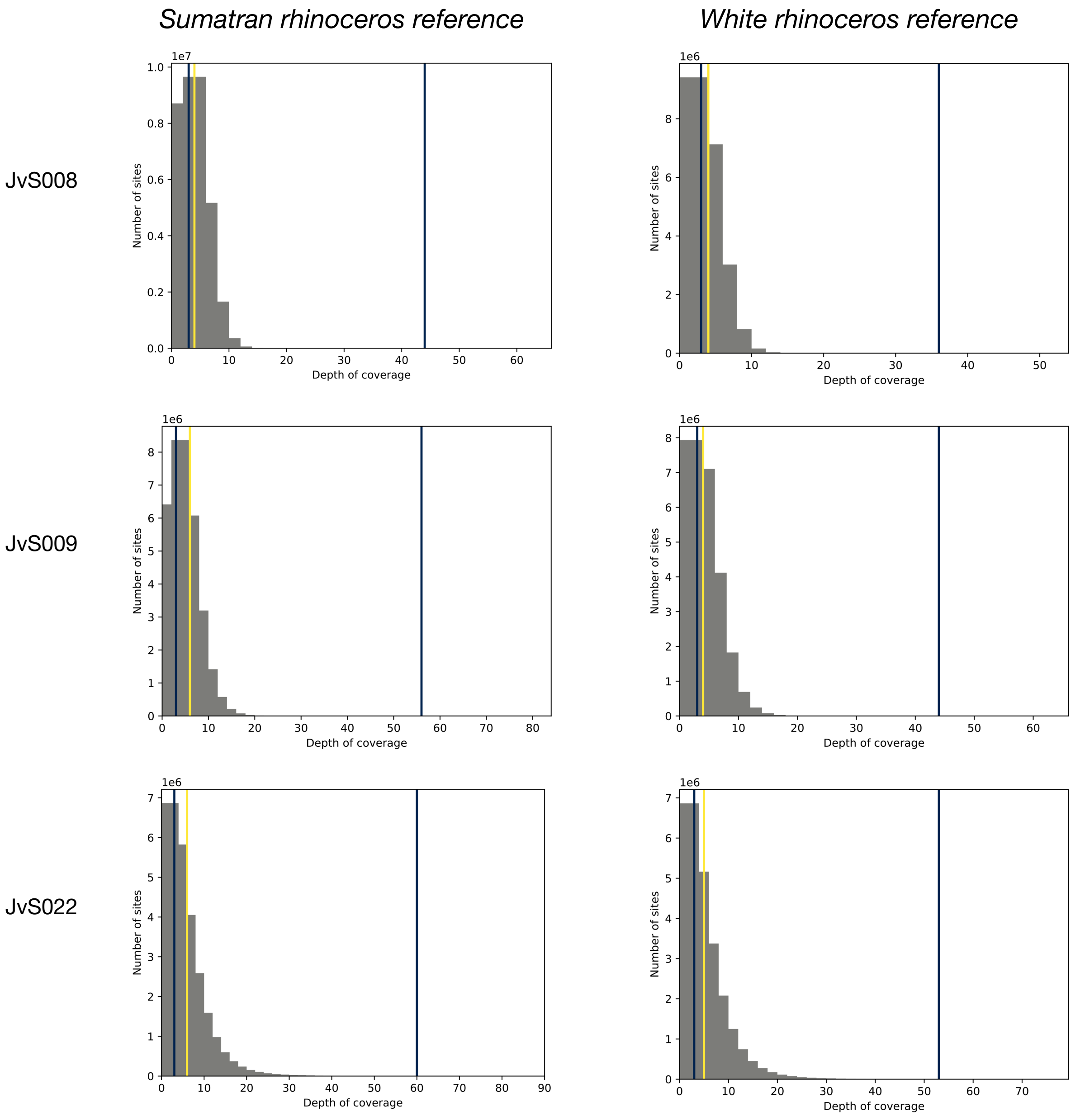


**Figure S2. Depth of coverage histograms from historical and modern Sumatran rhinoceros samples.** A) Samples mapped to the Sumatran rhinoceros scaffolds. B) Samples mapped to the White rhinoceros scaffolds. Yellow lines indicate the average reference-wide depth (calculated excluding sites with missing data), blue lines indicate the minimum and maximum depth thresholds used in this analysis (1/3 and 10X the average genome-wide depth).

**B)**

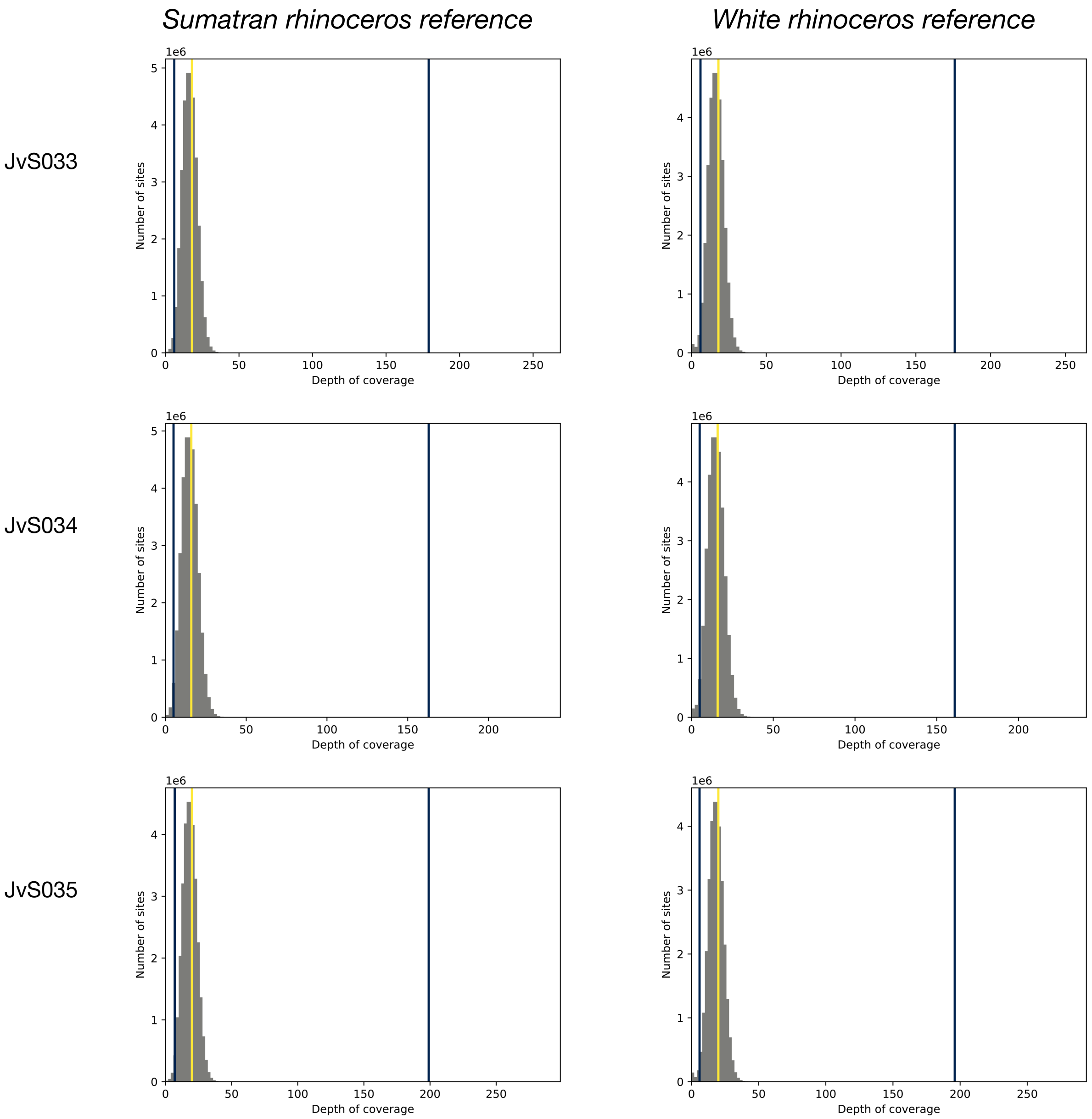


**Figure S2** *cont.*

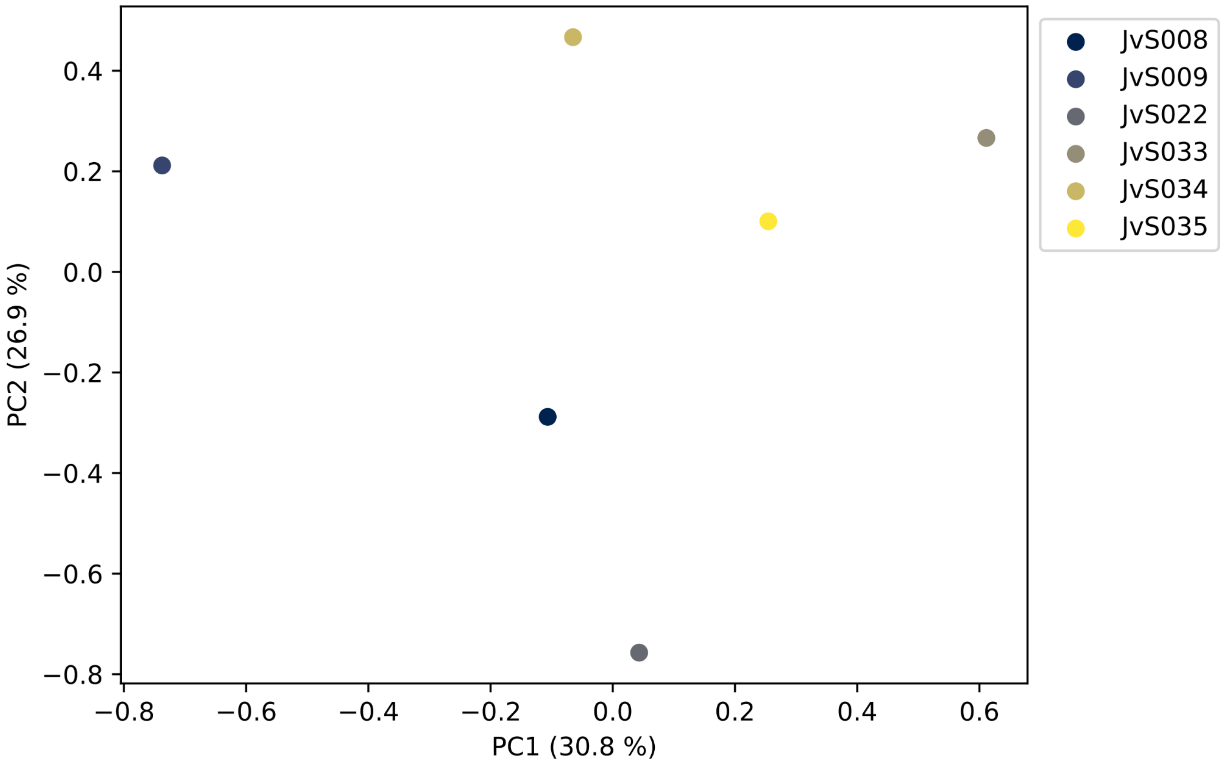


**Figure S3. Principal component analysis (PCA) of historical and modern Sumatran rhinoceros samples using the White rhinoceros reference**. Historical samples: JvS008, JvS009, JvS022. Modern samples: JvS033, JvS034, JvS035. See “PCA” in Additional file 2.

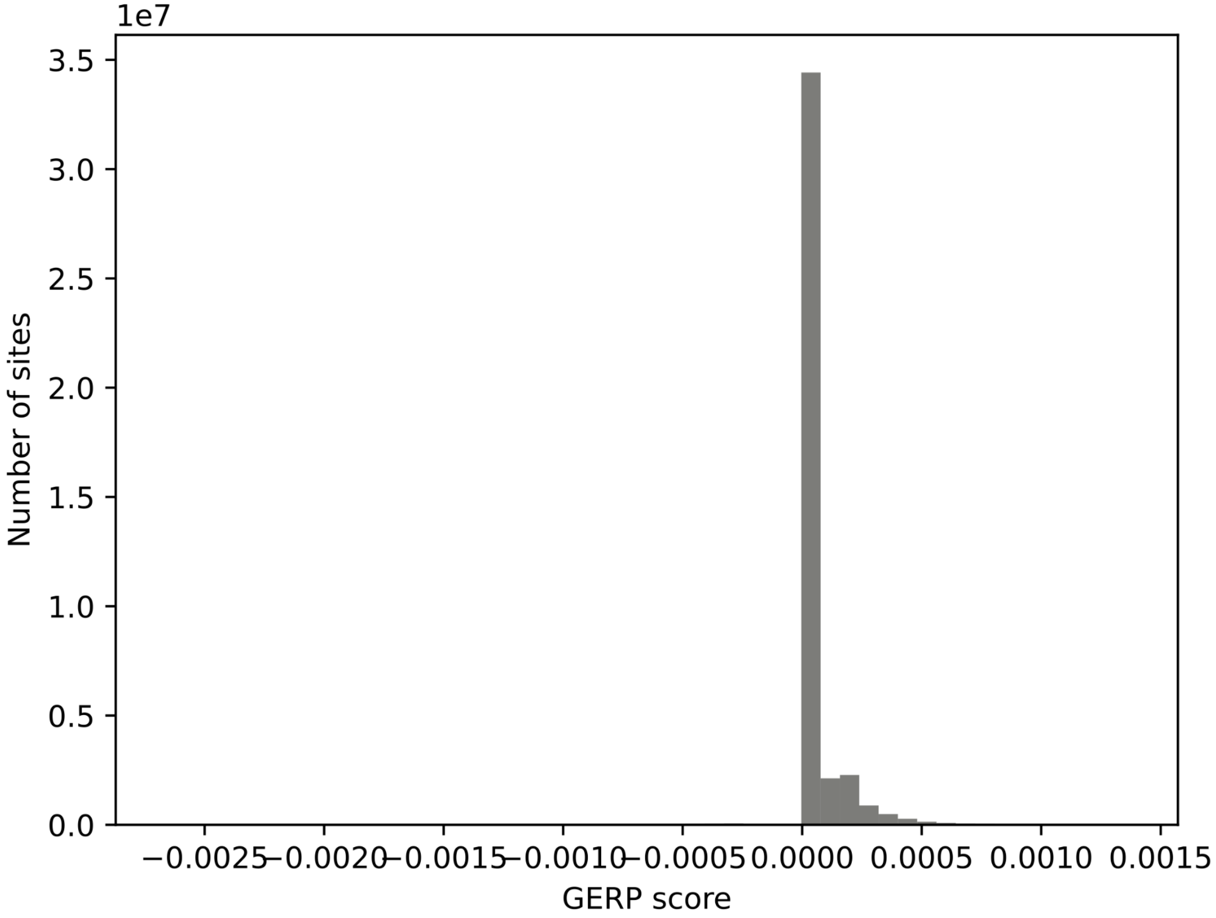


**Figure S4. Histogram of GERP scores estimated using the White rhinoceros scaffolds as reference and 30 mammalian outgroup species.**
