## Additional file 2 for "GenErode: a bioinformatics pipeline to investigate genome erosion in endangered and extinct species": Additional_file_2_v3.html

Snakemake Report

Loading Snakemake Report...

Please enable Javascript in your browser to see this report.

Loading 4.8 MB. For large reports, this can take a while.

Snakemake Report

- Wed Jan 19 11:35:08 2022 CET
- Snakemake 6.12.1

- GenErode pipeline report (current)
- Statistics
- Configuration

###### Results

- BAM file processing
- GERP
- PCA
- snpEff

### GenErode pipeline report

This is an automatic report from an execution of GenErode, created by Snakemake.

Each subsection in the Results section to the left contains a brief description of the output and links to some of the output files and the code used to generate them.
If you publish any research using results produced with GenErode, please cite our publication and refer to the source code.

Result sections are available for the following pipeline steps:

- BAM file processing
- mlRho
- PCA (plink)
- Runs of homozygosity (ROH; plink)
- snpEff
- GERP

Please set all steps to "True" in the config.yaml file that you would like to include into the report.

The rulegraph for GenErode is very large. If you wish to zoom into the image and inspect how the different rules of the pipeline are connected to each other, please save it as SVG or PNG (via the icon in the upper right corner of the rulegraph).

### BAM file processing

### GERP

### PCA

### snpEff

### Statistics

If the workflow has been executed in cluster/cloud, runtimes include the waiting time in the queue.

### Configuration

Configuration files

| File | Code |
| --- | --- |
|  | ``` 1 2 3 4 5 6 7 8 9 10 11 12 13 14 15 16 17 18 19 20 21 22 23 24 25 26 27 28 29 30 31 32 33 34 35 36 37 38 39 40 41 42 43 44 45 46 47 48 49 50 51 52 53 54 55 56 57 58 59 60 61 62 63 64 65 66 67 68 69 70 71 72 73 74 75 76 77 78 79 80 81 82 83 84 85 86 87 88 89 90 91 92 93 94 95 96 97 98 99 100 101 102 103 104 105 106 107 108 109 110 111 112 113 114 115 116 117 118 119 120 121 122 123 124 125 126 127 128 129 130 131 132 133 134 135 136 137 138 139 140 141 142 143 144 145 146 147 148 149 150 151 152 153 154 155 156 157 158 159 160 161 162 163 164 165 166 167 168 169 170 171 172 173 174 175 176 177 178 179 180 181 182 183 184 185 186 187 188 189 190 191 192 193 194 195 196 197 198 199 200 201 202 203 204 205 206 207 208 209 210 211 212 213 214 215 216 217 218 219 220 221 222 223 224 225 226 227 228 229 230 231 232 233 234 235 236 237 238 239 240 241 242 243 244 245 246 247 248 249 250 251 252 253 254 255 256 257 258 259 260 261 262 263 264 265 266 267 268 269 270 271 272 273 274 275 276 277 278 279 280 281 282 283 284 285 286 287 288 289 290 291 292 293 294 295 296 297 298 299 300 301 302 303 304 305 306 307 308 309 310 311 312 313 314 315 316 317 318 319 320 321 322 323 324 325 326 327 328 329 330 331 332 333 334 335 336 337 338 339 340 341 342 343 344 345 346 347 348 349 350 351 352 353 354 355 356 357 358 359 360 361 362 363 364 365 366 367 368 369 370 371 372 373 374 375 376 377 378 379 380 381 382 383 384 385 386 387 388 389 390 391 392 393 394 395 396 397 398 399 400 401 402 403 404 405 406 407 408 409 410 411 412 413 414 415 416 417 418 419 420 421 422 423 424 425 426 427 428 429 430 431 432 433 434 435 436 437 438 439 440 441 442 443 444 445 446 447 448 449 450 451 452 453 454 455 456 457 458 459 460 461 462 463 464 465 466 467 468 469 470 471 472 473 474 475 476 477 478 479 480 481 482 483 484 485 486 487 488 489 490 491 492 493 494 495 496 497 498 499 500 501 502 503 504 505 506 507 508 509 510 511 512 513 514 515 516 517 518 519 520 521 ``` \| ``` ################################################################# ################################################################# # Configuration settings for the GenErode pipeline v0.4.1 # # for ancient or historical samples, and modern samples # ################################################################# ################################################################# ################################################################# ################################################################# # Check the GitHub WIKI page for the full documentation: # # https://github.com/NBISweden/GenErode/wiki # ################################################################# ################################################################# ################################################################# ################################################################# # 1) Full path to reference genome assembly. # Reference genome has to be checked for short and concise FASTA # headers without special characters and has to be uncompressed. # The file name will be reused by the pipeline and can have the file # name extensions ".fasta" or ".fa". ref_path: "/proj/sllstore2017093/b2016342/b2016342_nobackup/lts/genome_erosion_pipeline/verena_testing/testdata/reference/GCF_000283155.1_CerSimSim1.0_genomic.Sc9M7eS_2_HRSCAF_41.fasta" ################################################################# ################################################################# # 2) Relative paths (from the main snakemake directory) to metadata # files with sample information. # Example files can be found in "config/" historical_samples: "config/rhino_3_historical_samples.txt" # leave empty ("") if not run for historical samples. modern_samples: "config/rhino_3_modern_samples.txt" # leave empty ("") if not run for modern samples. ################################################################# ################################################################# ################################################################# ################################################################# # 3) Pipeline steps to be run and related parameters. # If a step is set to True, all previous steps it depends on will # be loaded automatically. # Only one step should be set to True at a time, results and reports # should be double checked before continuing. ################################################################# ################################################################# # Rules for data processing (required for downstream analyses) # ################################################################# ##### # Repeat element de novo prediction and repeat masking of the # reference genome. # Generates BED files of repeats and repeat-masked regions for # the reference genome. # Output files will be placed into the same directory as the # reference genome FASTA file (as specified above). # That way, this step is run only once for a given reference genome. reference_repeat_identification: False ##### ##### # FastQC on raw reads, adapter trimming (incl. read merging # for historical samples) using fastp, FastQC on trimmed reads. # Adapter sequences are automatically detected. # Automatic detection of NovaSeq or NextSeq samples and activation of # poly-G tail trimming. fastq_processing: False # Minimum read length. # Historical samples (after trimming and read merging) hist_readlength: "30" # recommended setting: 30 bp # Modern samples (after trimming) mod_readlength: "30" ##### ##### # OPTIONAL: # Map historical reads to mitochondrial genomes from human and # several animal species that may be a contamination source. # Specify mitochondrial genome of target species ("species_mt"). # This step does not produce any files required for any of the # downstream steps and is therefore not included when running # any of the downstream steps. map_historical_to_mitogenomes: False # Full path including file name of mitochondrial genome FASTA of # the species under investigation, or that of a closely related species. # The FASTA file has to be uncompressed. The file name will be # reused by the pipeline and can have the file name extensions # ".fasta", ".fa" or ".fna". # Variable can be left empty ("") if not run. species_mt_path: ".test/data/references/rhino_NC_012684.fasta" # e.g. "/path/to/speciesname_accession_number.fasta" # Per default, only merged historical reads are mapped to the # different mitochondrial genomes. # If both merged and unmerged reads should be mapped, set the # following variable to True. map_unmerged_reads: False ##### ##### # Map historical and modern reads to reference genome assembly (specified above). mapping: False ##### ##### # BAM file processing: sort, merge samples from different lanes per PCR/index, # remove duplicates, merge BAM files per sample, realign indels, # calculate average genome-wide depth of coverage for depth filtering # of BAM and VCF files. bam_rmdup_realign_indels: False # Parameters related to depth filtering of BAM and VCF files. # After BAM file processing, the average genome-wide depth is calculated # per sample, from which minimum and maximum depth thresholds for quality # filtering are determined. # In the calculation of the average genome-wide depth of coverage, # sites with missing data (i.e. zero coverage) can be included or excluded. # Set to True if sites with missing data (zero coverage) should be # included in the average depth calculation. # Set to False if sites with missing data (zero coverage) should be # excluded from the average depth calculation. zerocoverage: False # Minimum depth threshold calculation per sample. # Will be applied to mlRho analysis and in VCF file filtering. # Factor by which the average genome-wide depth should be multiplied # to set a minimum depth threshold. For ultra low coverage samples, a # minimum hard threshold of 3X is applied that overrides this parameter. # A minimum depth of 6X should be aimed for. minDP: 0.33 # Maximum depth threshold calculation per sample. # Will be applied to mlRho analysis and in VCF file filtering. # Factor by which the average genome-wide depth should be multiplied # to set a maximum depth threshold. maxDP: 10 ##### ##### # CHECKPOINT: # Depth histograms with minimum and maximum depth thresholds and # average genome-wide depth as vertical lines and multiQC reports # are available in the GenErode pipeline report. # Carefully check the histograms and multiQC reports and adjust # the depth parameters above if necessary before moving on. ##### ##### # OPTIONAL: # Run mapDamage2 on historical samples specified in the list # "historical_rescaled_samplenames" below. # Will rescale base qualities at potentially damaged sites and # calculate statistics on ancient DNA damage patterns in # realigned bam files. historical_bam_mapDamage: False # List of historical samples on which mapDamage2 should be run. # Sample names without lane or index number in quotation marks, # separated by commas (e.g. ["VK01", "VK02", "VK03"]). # List has to be left empty ([]) if mapDamage2 is not run. # Keep the list of sample names for the entire pipeline run # if rescaled BAM files should be used in downstream analyses. historical_rescaled_samplenames: [] ##### ##### # OPTIONAL: # Subsample BAM files to a certain average genome-wide depth of # coverage to reduce biases in downstream analyses. bam_subsampling: False # Depth to which BAM files should be subsampled. # Check the GenErode report to screen the dataset for an appropriate # depth for subsampling. # Has to be lower than or equal to the average depth of the sample # with the lowest depth in the list below. # Keep the variable for the entire pipeline run so that the correct # files are used for downstream analyses. # Has to be set to False if not run. subsampling_depth: 6 # Combined list of modern and historical samples that should be # subsampled to a common depth. # Provide the sample names without lane or index number in # quotation marks, separated by commas (e.g. ["VK01", "VK02", "VK03"]). # List has to be left empty ([]) if subsampling is not run. # Keep the list of sample names for the entire pipeline run # if subsampled BAM files should be used in downstream analyses. subsampling_samplenames: ["JvS033", "JvS034", "JvS035"] ##### ##### # Call variants per sample from BAM files processed with mandatory # (and optional, if chosen) steps listed above. # Required for CpG filtering of BAM files for mlRho, ROH estimation, # PCA plotting, SNP annotation, and relative mutational load calculations # (from GERP scores). genotyping: False ##### ##### # OPTIONAL: # Identify CpG sites for removal from VCF files and from # downstream analyses and define samples to be CpG filtered. # Three different methods are available to identify CpG sites. # This step will generate several BED files containing genome # coordinates of CpG sites (file name ending with: "*.CpG_method.bed"), # all genome regions outside of CpG sites ("*.noCpG_method.bed"), # as well as intersected BED files of coordinates of CpG sites # and repeat elements ("*.CpG_method.repeats.bed") and regions # outside of CpG sites and repeat elements ("*.noCpG_method.repma.bed"). # Set CpG_identification and one of the three methods listed # below to True and specify samples in the list below in # which CpG sites should be identified and/or from which # CpG sites should be removed. CpG_identification: False # Method 1: # Identify CpG sites in single-individual VCF files of samples # listed below. BED files with CpG sites from all samples are # merged into one file for CpG site filtering. Only genotype # information is considered for CpG site identification. # Has to be kept at True for the rest of the pipeline, if chosen # method for CpG identification, so that the correct files are # used for downstream analyses. CpG_from_vcf: False # Method 2: # Identify CpG sites in the reference genome. # Ignores genotype information of samples mapped to the reference # genome. # Has to be kept at True for the rest of the pipeline, if chosen # method for CpG identification, so that the correct files are # used for downstream analyses. CpG_from_reference: True # Method 3: # Identify CpG sites in single-individual VCF files and in the # reference genome. BED files with CpG sites from all samples # and from the reference genome are merged into one file for # CpG site filtering. # Has to be kept at True for the rest of the pipeline, if chosen # method for CpG identification, so that the correct files are # used for downstream analyses. CpG_from_vcf_and_reference: False # Combined list of modern and historical samples in which CpG sites # should be identified if CpG_from_vcf or CpG_from_vcf_and_reference # is set to True, and from which CpG sites should be removed based # on any of the three available CpG identification methods chosen above. # Sample names without lane or index number in quotation marks, # separated by commas (e.g. ["VK01", "VK02", "VK03"]). # List has to be left empty ([]) if neither CpG identification nor # CpG site filtering is run. # Keep the list of sample names for the entire pipeline run # so that the correct files are used for downstream analyses. CpG_samplenames: ["JvS008", "JvS009", "JvS022", "JvS033", "JvS034", "JvS035"] ##### ################################################################# ################################################################# # Rules for BAM file processing for mlRho # ################################################################# ##### # OPTIONAL: # Generate BED files of autosomes and sex chromosomes for mlRho # analyses, in case these should be analyzed separately from each # other (see below for further options). # Includes intersecting of the new chromosome-specific BED files # with CpG- and repeat-masking BED files for downstream filtering. autosome_sexchromosome_bed_files: False # Relative path (from the main snakemake directory) to file listing # scaffolds/contigs linked to sex chromosomes (one scaffold/contig # name per line). # Leave empty ("") if identity of sex chromosomes unknown and/or # if mlRho should be run on all scaffolds/contigs of the genome. # Keep the path to the file when running the next step (mlRho) # separately for autosomes and sex chromosomes or only for autosomes. sexchromosomes: "" # for example, "config/chrX_candidate_scaffolds.txt" ##### ##### # Run mlRho 2.9 on filtered BAM files. # Automatically generates a PDF file with a plot of genome-wide # theta (with confidence intervals from mlRho) for each sample # (split up into autosomes and sex-chromosomes if specified below) # that can be retrieved from the GenErode pipeline report. # Default filters: quality, depth (based on parameters set above) # and repeat elements. # Requires sample lists and parameters for any desired optional # data filtering step (as described above). mlRho: False # There are three options how to run mlRho: # 1) If the identity of sex-chromosomal contigs/scaffolds is unknown # and/or mlRho should be run on all contigs/scaffolds, # set mlRho_autosomes_sexchromosomes to False and do not provide # a path to a text file with sex-chromosomal contigs/scaffolds # above when running mlRho. # # 2) If the identity of sex-chromosomal contigs/scaffolds is known, # mlRho analyses can be run for autosomes and sex chromosomes # separately from each other. # In that case, set mlRho_autosomes_sexchromosomes to True and # provide the path to the file with sex-chromosomal contigs/scaffolds # above when running mlRho. # # 3) If the identity of sex-chromosomal contigs/scaffolds is known, # sex-chromosomal contigs/scaffolds can be entirely excluded from # the analysis. # In that case, set mlRho_autosomes_sexchromosomes to False and # provide the path to the file with sex-chromosomal contigs/scaffolds # above when running mlRho. mlRho_autosomes_sexchromosomes: False ##### ################################################################# ################################################################# ################################################################# ################################################################# # Rules for VCF file processing for PCA plots, ROH estimation # # SNP annotation, and relative mutational load calculations # ################################################################# ##### # OPTIONAL: # Remove CpG sites from BCF files of of historical and modern samples # specified in the list "CpG_samplenames" under "CpG_identification". vcf_CpG_filtering: False ##### ##### # Filter BCF files of historical and modern samples for quality, # depth (based on parameters set above), indels, allelic imbalance, # and remove repeat regions that were identified in the reference genome. vcf_qual_repeat_filtering: False ##### ##### # Merge BCF files into a BCF file containing all samples and remove all # sites that are not biallelic and with missing data across all samples # up to a certain threshold as defined below. # Extract 1) all historical and 2) all modern samples from the merged and # filtered BCF file. # Create a BED file of sites that remain after filtering across all samples # to be used for downstream filtering of individual BCF files. merge_vcfs_per_dataset: False # Maximum allowed fraction of missing genotypes across all samples for a # site to be kept in the BCF and BED file, to ensure that the same sites # are compared between historical and modern samples. f_missing: 0.1 # default: 0.1 (i.e. maximum 10% missing genotypes per site) ##### ################################################################# ################################################################# ################################################################# ################################################################# # Rules for PCA, ROH estimation and SNP annotation # ################################################################# ##### # Plot PCAs for 1) all historical, 2) all modern, and 3) all modern # and historical samples combined based on the merged BCF files. # Automatically generates PDF files with plots of PC1 vs. PC2 and # PC1 vs. PC3 that can be retrieved from the GenErode pipeline report. # Requires sample lists and parameters for any desired optional # data filtering step (as described above). pca: True ##### ##### # Run plink 1.9 on merged BCF files to estimate runs of homozygosity. # Automatically generates PDF files with plots of F(ROH) larger than # 2 Mb in each sample that can be retrieved from the GenErode pipeline report. # Requires sample lists and parameters for any desired optional # data filtering step (as described above). ROH: False # Parameters: # Set a fixed combination of the following parameters for a given run. # See https://www.cog-genomics.org/plink/1.9/ibd#homozyg for more details. # Minimum SNP count. For example, 10, 25. Abbreviation in file name: homsnp. homozyg-snp: 25 # Minimum size of ROH in kilobases. Abbreviation in file name: homkb. homozyg-kb: 100 # Window size for ROH estimation. For example, 20, 50, 100, 1000. # Abbreviation in file name: homwinsnp. homozyg-window-snp: 250 # Maximum number of heterozygote sites per window. # For example, 1 for a stringent analysis, 3 for a relaxed setting. # Abbreviation in file name: homwinhet. homozyg-window-het: 3 # Maximum number of missing sites per window. # For example, 1 or 5 for a stringent analysis, # 10 for intermediate filtering and 15 for relaxed filtering. # Abbreviation in file name: homwinmis. homozyg-window-missing: 15 # Maximum number of heterozygote sites per ROH. # For example, 1 for a stringent analysis, # 3 as a relaxed setting. Disable this parameter by setting it to 999. # Abbreviation in file name: homhet. homozyg-het: 750 ##### ##### # Run snpEff v4.3.1 on per-sample BCF files to annotate SNPs # in protein-coding regions. # Automatically generates a PDF file with a plot of the numbers of # SNPs with high, moderate, low, and modifier effects for each sample # that can be retrieved from the GenErode pipeline report. # Sites from the BED file not passing the missingness and biallelic sites # filter across all samples are removed per default from the per-sample # BCF files. # Requires sample lists and parameters for any desired optional # data filtering step (as described above). snpEff: True # Full path to GTF file (GTF format version 2.2) with gene model predictions # for the reference genome specified at the top of this config file. # Read here about tools for file format conversions from GFF to GTF: # https://github.com/NBISweden/GAAS/blob/master/annotation/knowledge/gff_to_gtf.md # Output files from building the snpEff database will be placed into the # same directory as the GTF file. # That way, the database has to be built only once for a given GTF file. gtf_path: "/proj/sllstore2017093/b2016342/b2016342_nobackup/lts/genome_erosion_pipeline/verena_testing/testdata/reference/GCF_000283155.1_CerSimSim1.0_genomic.Sc9M7eS_2_HRSCAF_41.gtf" ################################################################# ################################################################# ################################################################# ################################################################# # Rules for GERP score and relative mutational load calculations# ################################################################# ##### # Estimate GERP scores for the reference genome of the target species, # based on at least 30 outgroup genomes and a dated phylogenetic tree, # and (optionally) calculate relative mutational load per sample from # GERP scores and counts of derived alleles per site and sample from # filtered per-sample BCF files (homozygous sites counted as 2 derived alleles # and heterozygous sites as 1 derived allele). Sites from the BED file not # passing the missingness and biallelic sites filter across all samples # are removed per default from the per-sample BCF files. # Relative mutational load is calculated per sample for all sites with # GERP scores within "min_gerp" and "max_gerp" (specified below) based on # the counts of derived alleles for these sites. # Automatically generates a PDF file with a histogram of GERP scores # that can be used to choose minimum and maximum GERP score thresholds, and # a PDF file with a plot of relative mutational load estimates per sample # that can be retrieved from the GenErode pipeline report. # Requires path to reference genome as provided at the top of this # config file. If relative mutational load should be calculated per sample, # metadata files with historical and/or modern samples are required. # If metadata files are provided, sample lists and parameters for any # desired optional data filtering step (as described above) are required. gerp: True # Full path to directory containing reference genomes of outgroup species in # FASTA format for GERP++ score estimation. # Files must be gzipped and FASTA file name extensions can be # "*.fa.gz", "*.fasta.gz" or "*.fna.gz". gerp_ref_path: "/proj/sllstore2017093/b2016342/b2016342_nobackup/lts/genome_erosion_pipeline/verena_testing/testdata/gerp/outgroup_Sc9M7eS_2_HRSCAF_41/all_scaffolds/" # Full path to phylogenetic tree of all species included in the analysis # (including the target species) in NEWICK format and including divergence # time estimates. # Divergence time estimates must be in billions of years for correct scaling # of GERP scores (see dated phylogenetic trees from www.timetree.org). # Species names in the tree have to be identical to the fasta file names # without ".fa.gz", ".fasta.gz" or ".fna.gz". tree: "/proj/sllstore2017093/b2016342/b2016342_nobackup/lts/genome_erosion_pipeline/verena_testing/testdata/scripts/testdataset_paper/Sc9M7eS_2_HRSCAF_41/gerp/gerp_30_outgroups_plus_whi_rhino_names.nwk" # Minimum and maximum GERP score for a site to be included into calculations # of relative mutational load. # Positive values indicate purifying selection. min_gerp: 0.00048928 max_gerp: 1000 ##### # NOTE: # The GERP step produces a large number of large intermediate files, # so several TB of storage space may required during a pipeline run. # The required storage space for intermediate files scales with the # numbers of outgroup genomes and samples. # The number of intermediate files scales additionally with the level of # fragmentation of the reference genome (i.e. with the number of contigs/ # scaffolds). ##### ##### ``` |

Loading...

×

Download original

###### Rule make\_no\_repeats\_bed

×

Rule properties

|  |  |
| --- | --- |
| Jobs | 1 |
| Input files | |
| - /proj/sllstore2017093/b2016342/b2016342\_nobackup/lts/genome\_erosion\_pipeline/verena\_testing/testdata/reference/GCF\_000283155.1\_CerSimSim1.0\_genomic.Sc9M7eS\_2\_HRSCAF\_41.bed - /proj/sllstore2017093/b2016342/b2016342\_nobackup/lts/genome\_erosion\_pipeline/verena\_testing/testdata/reference/GCF\_000283155.1\_CerSimSim1.0\_genomic.Sc9M7eS\_2\_HRSCAF\_41.repeats.sorted.bed | |
| Output files | |
| - /proj/sllstore2017093/b2016342/b2016342\_nobackup/lts/genome\_erosion\_pipeline/verena\_testing/testdata/reference/GCF\_000283155.1\_CerSimSim1.0\_genomic.Sc9M7eS\_2\_HRSCAF\_41.repma.bed - results/GCF\_000283155.1\_CerSimSim1.0\_genomic.Sc9M7eS\_2\_HRSCAF\_41.repma.bed | |
| Container image | |
| docker://quay.io/biocontainers/bedtools:2.29.2--hc088bd4\_0 | |
| Code | |
| ``` 1 2 3 ``` \| ``` bedtools subtract -a {input.ref_bed} -b {input.sorted_rep_bed} > {output.no_rep_bed} 2> {log} && cp {output.no_rep_bed} {output.no_rep_bed_dir} 2>> {log} ``` | |

###### Rule multiqc\_historical\_raw

×

Rule properties

|  |  |
| --- | --- |
| Jobs | 1 |
| Input files | |
| - data/raw\_reads\_symlinks/historical/stats/JvS008\_08\_L2\_R1\_fastqc.html - data/raw\_reads\_symlinks/historical/stats/JvS008\_08\_L2\_R2\_fastqc.html - data/raw\_reads\_symlinks/historical/stats/JvS008\_08\_L6\_R1\_fastqc.html - data/raw\_reads\_symlinks/historical/stats/JvS008\_08\_L6\_R2\_fastqc.html - data/raw\_reads\_symlinks/historical/stats/JvS008\_10\_L2\_R1\_fastqc.html - data/raw\_reads\_symlinks/historical/stats/JvS008\_10\_L2\_R2\_fastqc.html - data/raw\_reads\_symlinks/historical/stats/JvS008\_10\_L6\_R1\_fastqc.html - data/raw\_reads\_symlinks/historical/stats/JvS008\_10\_L6\_R2\_fastqc.html - data/raw\_reads\_symlinks/historical/stats/JvS008\_11\_L2\_R1\_fastqc.html - data/raw\_reads\_symlinks/historical/stats/JvS008\_11\_L2\_R2\_fastqc.html - data/raw\_reads\_symlinks/historical/stats/JvS008\_11\_L6\_R1\_fastqc.html - data/raw\_reads\_symlinks/historical/stats/JvS008\_11\_L6\_R2\_fastqc.html - data/raw\_reads\_symlinks/historical/stats/JvS009\_09\_L3\_R1\_fastqc.html - data/raw\_reads\_symlinks/historical/stats/JvS009\_09\_L3\_R2\_fastqc.html - data/raw\_reads\_symlinks/historical/stats/JvS009\_09\_L7\_R1\_fastqc.html - data/raw\_reads\_symlinks/historical/stats/JvS009\_09\_L7\_R2\_fastqc.html - data/raw\_reads\_symlinks/historical/stats/JvS009\_15\_L3\_R1\_fastqc.html - data/raw\_reads\_symlinks/historical/stats/JvS009\_15\_L3\_R2\_fastqc.html - data/raw\_reads\_symlinks/historical/stats/JvS009\_15\_L7\_R1\_fastqc.html - data/raw\_reads\_symlinks/historical/stats/JvS009\_15\_L7\_R2\_fastqc.html - data/raw\_reads\_symlinks/historical/stats/JvS009\_19\_L3\_R1\_fastqc.html - data/raw\_reads\_symlinks/historical/stats/JvS009\_19\_L3\_R2\_fastqc.html - data/raw\_reads\_symlinks/historical/stats/JvS009\_19\_L7\_R1\_fastqc.html - data/raw\_reads\_symlinks/historical/stats/JvS009\_19\_L7\_R2\_fastqc.html - data/raw\_reads\_symlinks/historical/stats/JvS022\_01\_L1\_R1\_fastqc.html - data/raw\_reads\_symlinks/historical/stats/JvS022\_01\_L1\_R2\_fastqc.html - data/raw\_reads\_symlinks/historical/stats/JvS022\_02\_L1\_R1\_fastqc.html - data/raw\_reads\_symlinks/historical/stats/JvS022\_02\_L1\_R2\_fastqc.html - data/raw\_reads\_symlinks/historical/stats/JvS022\_03\_L1\_R1\_fastqc.html - data/raw\_reads\_symlinks/historical/stats/JvS022\_03\_L1\_R2\_fastqc.html - data/raw\_reads\_symlinks/historical/stats/JvS022\_04\_L1\_R1\_fastqc.html - data/raw\_reads\_symlinks/historical/stats/JvS022\_04\_L1\_R2\_fastqc.html - data/raw\_reads\_symlinks/historical/stats/JvS022\_05\_L1\_R1\_fastqc.html - data/raw\_reads\_symlinks/historical/stats/JvS022\_05\_L1\_R2\_fastqc.html - data/raw\_reads\_symlinks/historical/stats/JvS022\_06\_L1\_R1\_fastqc.html - data/raw\_reads\_symlinks/historical/stats/JvS022\_06\_L1\_R2\_fastqc.html - data/raw\_reads\_symlinks/historical/stats/JvS022\_07\_L1\_R1\_fastqc.html - data/raw\_reads\_symlinks/historical/stats/JvS022\_07\_L1\_R2\_fastqc.html - data/raw\_reads\_symlinks/historical/stats/JvS022\_08\_L1\_R1\_fastqc.html - data/raw\_reads\_symlinks/historical/stats/JvS022\_08\_L1\_R2\_fastqc.html - data/raw\_reads\_symlinks/historical/stats/JvS022\_09\_L1\_R1\_fastqc.html - data/raw\_reads\_symlinks/historical/stats/JvS022\_09\_L1\_R2\_fastqc.html - data/raw\_reads\_symlinks/historical/stats/JvS022\_10\_L1\_R1\_fastqc.html - data/raw\_reads\_symlinks/historical/stats/JvS022\_10\_L1\_R2\_fastqc.html - data/raw\_reads\_symlinks/historical/stats/JvS022\_11\_L1\_R1\_fastqc.html - data/raw\_reads\_symlinks/historical/stats/JvS022\_11\_L1\_R2\_fastqc.html - data/raw\_reads\_symlinks/historical/stats/JvS022\_12\_L1\_R1\_fastqc.html - data/raw\_reads\_symlinks/historical/stats/JvS022\_12\_L1\_R2\_fastqc.html - data/raw\_reads\_symlinks/historical/stats/JvS022\_74\_L8\_R1\_fastqc.html - data/raw\_reads\_symlinks/historical/stats/JvS022\_74\_L8\_R2\_fastqc.html - data/raw\_reads\_symlinks/historical/stats/JvS022\_75\_L8\_R1\_fastqc.html - data/raw\_reads\_symlinks/historical/stats/JvS022\_75\_L8\_R2\_fastqc.html - data/raw\_reads\_symlinks/historical/stats/JvS022\_76\_L8\_R1\_fastqc.html - data/raw\_reads\_symlinks/historical/stats/JvS022\_76\_L8\_R2\_fastqc.html - data/raw\_reads\_symlinks/historical/stats/JvS022\_77\_L8\_R1\_fastqc.html - data/raw\_reads\_symlinks/historical/stats/JvS022\_77\_L8\_R2\_fastqc.html - data/raw\_reads\_symlinks/historical/stats/JvS022\_78\_L8\_R1\_fastqc.html - data/raw\_reads\_symlinks/historical/stats/JvS022\_78\_L8\_R2\_fastqc.html - data/raw\_reads\_symlinks/historical/stats/JvS022\_79\_L8\_R1\_fastqc.html - data/raw\_reads\_symlinks/historical/stats/JvS022\_79\_L8\_R2\_fastqc.html - data/raw\_reads\_symlinks/historical/stats/JvS022\_80\_L8\_R1\_fastqc.html - data/raw\_reads\_symlinks/historical/stats/JvS022\_80\_L8\_R2\_fastqc.html - data/raw\_reads\_symlinks/historical/stats/JvS022\_81\_L8\_R1\_fastqc.html - data/raw\_reads\_symlinks/historical/stats/JvS022\_81\_L8\_R2\_fastqc.html - data/raw\_reads\_symlinks/historical/stats/JvS022\_82\_L8\_R1\_fastqc.html - data/raw\_reads\_symlinks/historical/stats/JvS022\_82\_L8\_R2\_fastqc.html - data/raw\_reads\_symlinks/historical/stats/JvS022\_83\_L8\_R1\_fastqc.html - data/raw\_reads\_symlinks/historical/stats/JvS022\_83\_L8\_R2\_fastqc.html - data/raw\_reads\_symlinks/historical/stats/JvS022\_84\_L8\_R1\_fastqc.html - data/raw\_reads\_symlinks/historical/stats/JvS022\_84\_L8\_R2\_fastqc.html - data/raw\_reads\_symlinks/historical/stats/JvS022\_85\_L8\_R1\_fastqc.html - data/raw\_reads\_symlinks/historical/stats/JvS022\_85\_L8\_R2\_fastqc.html - data/raw\_reads\_symlinks/historical/stats/JvS008\_08\_L2\_R1\_fastqc.zip - data/raw\_reads\_symlinks/historical/stats/JvS008\_08\_L2\_R2\_fastqc.zip - data/raw\_reads\_symlinks/historical/stats/JvS008\_08\_L6\_R1\_fastqc.zip - data/raw\_reads\_symlinks/historical/stats/JvS008\_08\_L6\_R2\_fastqc.zip - data/raw\_reads\_symlinks/historical/stats/JvS008\_10\_L2\_R1\_fastqc.zip - data/raw\_reads\_symlinks/historical/stats/JvS008\_10\_L2\_R2\_fastqc.zip - data/raw\_reads\_symlinks/historical/stats/JvS008\_10\_L6\_R1\_fastqc.zip - data/raw\_reads\_symlinks/historical/stats/JvS008\_10\_L6\_R2\_fastqc.zip - data/raw\_reads\_symlinks/historical/stats/JvS008\_11\_L2\_R1\_fastqc.zip - data/raw\_reads\_symlinks/historical/stats/JvS008\_11\_L2\_R2\_fastqc.zip - data/raw\_reads\_symlinks/historical/stats/JvS008\_11\_L6\_R1\_fastqc.zip - data/raw\_reads\_symlinks/historical/stats/JvS008\_11\_L6\_R2\_fastqc.zip - data/raw\_reads\_symlinks/historical/stats/JvS009\_09\_L3\_R1\_fastqc.zip - data/raw\_reads\_symlinks/historical/stats/JvS009\_09\_L3\_R2\_fastqc.zip - data/raw\_reads\_symlinks/historical/stats/JvS009\_09\_L7\_R1\_fastqc.zip - data/raw\_reads\_symlinks/historical/stats/JvS009\_09\_L7\_R2\_fastqc.zip - data/raw\_reads\_symlinks/historical/stats/JvS009\_15\_L3\_R1\_fastqc.zip - data/raw\_reads\_symlinks/historical/stats/JvS009\_15\_L3\_R2\_fastqc.zip - data/raw\_reads\_symlinks/historical/stats/JvS009\_15\_L7\_R1\_fastqc.zip - data/raw\_reads\_symlinks/historical/stats/JvS009\_15\_L7\_R2\_fastqc.zip - data/raw\_reads\_symlinks/historical/stats/JvS009\_19\_L3\_R1\_fastqc.zip - data/raw\_reads\_symlinks/historical/stats/JvS009\_19\_L3\_R2\_fastqc.zip - data/raw\_reads\_symlinks/historical/stats/JvS009\_19\_L7\_R1\_fastqc.zip - data/raw\_reads\_symlinks/historical/stats/JvS009\_19\_L7\_R2\_fastqc.zip - data/raw\_reads\_symlinks/historical/stats/JvS022\_01\_L1\_R1\_fastqc.zip - data/raw\_reads\_symlinks/historical/stats/JvS022\_01\_L1\_R2\_fastqc.zip - data/raw\_reads\_symlinks/historical/stats/JvS022\_02\_L1\_R1\_fastqc.zip - data/raw\_reads\_symlinks/historical/stats/JvS022\_02\_L1\_R2\_fastqc.zip - data/raw\_reads\_symlinks/historical/stats/JvS022\_03\_L1\_R1\_fastqc.zip - data/raw\_reads\_symlinks/historical/stats/JvS022\_03\_L1\_R2\_fastqc.zip - data/raw\_reads\_symlinks/historical/stats/JvS022\_04\_L1\_R1\_fastqc.zip - data/raw\_reads\_symlinks/historical/stats/JvS022\_04\_L1\_R2\_fastqc.zip - data/raw\_reads\_symlinks/historical/stats/JvS022\_05\_L1\_R1\_fastqc.zip - data/raw\_reads\_symlinks/historical/stats/JvS022\_05\_L1\_R2\_fastqc.zip - data/raw\_reads\_symlinks/historical/stats/JvS022\_06\_L1\_R1\_fastqc.zip - data/raw\_reads\_symlinks/historical/stats/JvS022\_06\_L1\_R2\_fastqc.zip - data/raw\_reads\_symlinks/historical/stats/JvS022\_07\_L1\_R1\_fastqc.zip - data/raw\_reads\_symlinks/historical/stats/JvS022\_07\_L1\_R2\_fastqc.zip - data/raw\_reads\_symlinks/historical/stats/JvS022\_08\_L1\_R1\_fastqc.zip - data/raw\_reads\_symlinks/historical/stats/JvS022\_08\_L1\_R2\_fastqc.zip - data/raw\_reads\_symlinks/historical/stats/JvS022\_09\_L1\_R1\_fastqc.zip - data/raw\_reads\_symlinks/historical/stats/JvS022\_09\_L1\_R2\_fastqc.zip - data/raw\_reads\_symlinks/historical/stats/JvS022\_10\_L1\_R1\_fastqc.zip - data/raw\_reads\_symlinks/historical/stats/JvS022\_10\_L1\_R2\_fastqc.zip - data/raw\_reads\_symlinks/historical/stats/JvS022\_11\_L1\_R1\_fastqc.zip - data/raw\_reads\_symlinks/historical/stats/JvS022\_11\_L1\_R2\_fastqc.zip - data/raw\_reads\_symlinks/historical/stats/JvS022\_12\_L1\_R1\_fastqc.zip - data/raw\_reads\_symlinks/historical/stats/JvS022\_12\_L1\_R2\_fastqc.zip - data/raw\_reads\_symlinks/historical/stats/JvS022\_74\_L8\_R1\_fastqc.zip - data/raw\_reads\_symlinks/historical/stats/JvS022\_74\_L8\_R2\_fastqc.zip - data/raw\_reads\_symlinks/historical/stats/JvS022\_75\_L8\_R1\_fastqc.zip - data/raw\_reads\_symlinks/historical/stats/JvS022\_75\_L8\_R2\_fastqc.zip - data/raw\_reads\_symlinks/historical/stats/JvS022\_76\_L8\_R1\_fastqc.zip - data/raw\_reads\_symlinks/historical/stats/JvS022\_76\_L8\_R2\_fastqc.zip - data/raw\_reads\_symlinks/historical/stats/JvS022\_77\_L8\_R1\_fastqc.zip - data/raw\_reads\_symlinks/historical/stats/JvS022\_77\_L8\_R2\_fastqc.zip - data/raw\_reads\_symlinks/historical/stats/JvS022\_78\_L8\_R1\_fastqc.zip - data/raw\_reads\_symlinks/historical/stats/JvS022\_78\_L8\_R2\_fastqc.zip - data/raw\_reads\_symlinks/historical/stats/JvS022\_79\_L8\_R1\_fastqc.zip - data/raw\_reads\_symlinks/historical/stats/JvS022\_79\_L8\_R2\_fastqc.zip - data/raw\_reads\_symlinks/historical/stats/JvS022\_80\_L8\_R1\_fastqc.zip - data/raw\_reads\_symlinks/historical/stats/JvS022\_80\_L8\_R2\_fastqc.zip - data/raw\_reads\_symlinks/historical/stats/JvS022\_81\_L8\_R1\_fastqc.zip - data/raw\_reads\_symlinks/historical/stats/JvS022\_81\_L8\_R2\_fastqc.zip - data/raw\_reads\_symlinks/historical/stats/JvS022\_82\_L8\_R1\_fastqc.zip - data/raw\_reads\_symlinks/historical/stats/JvS022\_82\_L8\_R2\_fastqc.zip - data/raw\_reads\_symlinks/historical/stats/JvS022\_83\_L8\_R1\_fastqc.zip - data/raw\_reads\_symlinks/historical/stats/JvS022\_83\_L8\_R2\_fastqc.zip - data/raw\_reads\_symlinks/historical/stats/JvS022\_84\_L8\_R1\_fastqc.zip - data/raw\_reads\_symlinks/historical/stats/JvS022\_84\_L8\_R2\_fastqc.zip - data/raw\_reads\_symlinks/historical/stats/JvS022\_85\_L8\_R1\_fastqc.zip - data/raw\_reads\_symlinks/historical/stats/JvS022\_85\_L8\_R2\_fastqc.zip | |
| Output files | |
| - data/raw\_reads\_symlinks/historical/stats/multiqc/multiqc\_report.html | |
| Container image | |
| docker://quay.io/biocontainers/multiqc:1.9--pyh9f0ad1d\_0 | |
| Code | |
| ``` 1 2 ``` \| ``` multiqc -f {params.indir} -o {params.outdir} 2> {log} ``` | |

###### Rule fastqc\_historical\_raw

×

Rule properties

|  |  |
| --- | --- |
| Jobs | 72 |
| Input files | |
| - data/raw\_reads\_symlinks/historical/{sample}\_{index}\_{lane}\_R{nr}.fastq.gz | |
| Output files | |
| - data/raw\_reads\_symlinks/historical/stats/{sample,[A-Za-z0-9]+}\_{index}\_{lane}\_R{nr}\_fastqc.html - data/raw\_reads\_symlinks/historical/stats/{sample,[A-Za-z0-9]+}\_{index}\_{lane}\_R{nr}\_fastqc.zip - data/raw\_reads\_symlinks/historical/stats/{sample,[A-Za-z0-9]+}\_{index}\_{lane}\_R{nr}\_fastqc | |
| Container image | |
| docker://biocontainers/fastqc:v0.11.9\_cv7 | |
| Code | |
| ``` 1 2 ``` \| ``` fastqc -o {params.dir} -t {threads} --extract {input.fastq} 2> {log} ``` | |

###### Rule fastq\_historical\_symbolic\_links

×

Rule properties

|  |  |
| --- | --- |
| Jobs | 36 |
| Input files | |
| - config/rhino\_3\_historical\_samples.txt | |
| Output files | |
| - data/raw\_reads\_symlinks/historical/{sample,[A-Za-z0-9]+}\_{index}\_{lane}\_R1.fastq.gz - data/raw\_reads\_symlinks/historical/{sample,[A-Za-z0-9]+}\_{index}\_{lane}\_R2.fastq.gz | |

###### Rule multiqc\_historical\_trimmed

×

Rule properties

|  |  |
| --- | --- |
| Jobs | 1 |
| Input files | |
| - results/historical/trimming/stats/JvS008\_08\_L2\_trimmed\_merged\_fastqc.html - results/historical/trimming/stats/JvS008\_08\_L2\_trimmed\_merged\_fastqc.zip - results/historical/trimming/stats/JvS008\_08\_L6\_trimmed\_merged\_fastqc.html - results/historical/trimming/stats/JvS008\_08\_L6\_trimmed\_merged\_fastqc.zip - results/historical/trimming/stats/JvS008\_10\_L2\_trimmed\_merged\_fastqc.html - results/historical/trimming/stats/JvS008\_10\_L2\_trimmed\_merged\_fastqc.zip - results/historical/trimming/stats/JvS008\_10\_L6\_trimmed\_merged\_fastqc.html - results/historical/trimming/stats/JvS008\_10\_L6\_trimmed\_merged\_fastqc.zip - results/historical/trimming/stats/JvS008\_11\_L2\_trimmed\_merged\_fastqc.html - results/historical/trimming/stats/JvS008\_11\_L2\_trimmed\_merged\_fastqc.zip - results/historical/trimming/stats/JvS008\_11\_L6\_trimmed\_merged\_fastqc.html - results/historical/trimming/stats/JvS008\_11\_L6\_trimmed\_merged\_fastqc.zip - results/historical/trimming/stats/JvS009\_09\_L3\_trimmed\_merged\_fastqc.html - results/historical/trimming/stats/JvS009\_09\_L3\_trimmed\_merged\_fastqc.zip - results/historical/trimming/stats/JvS009\_09\_L7\_trimmed\_merged\_fastqc.html - results/historical/trimming/stats/JvS009\_09\_L7\_trimmed\_merged\_fastqc.zip - results/historical/trimming/stats/JvS009\_15\_L3\_trimmed\_merged\_fastqc.html - results/historical/trimming/stats/JvS009\_15\_L3\_trimmed\_merged\_fastqc.zip - results/historical/trimming/stats/JvS009\_15\_L7\_trimmed\_merged\_fastqc.html - results/historical/trimming/stats/JvS009\_15\_L7\_trimmed\_merged\_fastqc.zip - results/historical/trimming/stats/JvS009\_19\_L3\_trimmed\_merged\_fastqc.html - results/historical/trimming/stats/JvS009\_19\_L3\_trimmed\_merged\_fastqc.zip - results/historical/trimming/stats/JvS009\_19\_L7\_trimmed\_merged\_fastqc.html - results/historical/trimming/stats/JvS009\_19\_L7\_trimmed\_merged\_fastqc.zip - results/historical/trimming/stats/JvS022\_01\_L1\_trimmed\_merged\_fastqc.html - results/historical/trimming/stats/JvS022\_01\_L1\_trimmed\_merged\_fastqc.zip - results/historical/trimming/stats/JvS022\_02\_L1\_trimmed\_merged\_fastqc.html - results/historical/trimming/stats/JvS022\_02\_L1\_trimmed\_merged\_fastqc.zip - results/historical/trimming/stats/JvS022\_03\_L1\_trimmed\_merged\_fastqc.html - results/historical/trimming/stats/JvS022\_03\_L1\_trimmed\_merged\_fastqc.zip - results/historical/trimming/stats/JvS022\_04\_L1\_trimmed\_merged\_fastqc.html - results/historical/trimming/stats/JvS022\_04\_L1\_trimmed\_merged\_fastqc.zip - results/historical/trimming/stats/JvS022\_05\_L1\_trimmed\_merged\_fastqc.html - results/historical/trimming/stats/JvS022\_05\_L1\_trimmed\_merged\_fastqc.zip - results/historical/trimming/stats/JvS022\_06\_L1\_trimmed\_merged\_fastqc.html - results/historical/trimming/stats/JvS022\_06\_L1\_trimmed\_merged\_fastqc.zip - results/historical/trimming/stats/JvS022\_07\_L1\_trimmed\_merged\_fastqc.html - results/historical/trimming/stats/JvS022\_07\_L1\_trimmed\_merged\_fastqc.zip - results/historical/trimming/stats/JvS022\_08\_L1\_trimmed\_merged\_fastqc.html - results/historical/trimming/stats/JvS022\_08\_L1\_trimmed\_merged\_fastqc.zip - results/historical/trimming/stats/JvS022\_09\_L1\_trimmed\_merged\_fastqc.html - results/historical/trimming/stats/JvS022\_09\_L1\_trimmed\_merged\_fastqc.zip - results/historical/trimming/stats/JvS022\_10\_L1\_trimmed\_merged\_fastqc.html - results/historical/trimming/stats/JvS022\_10\_L1\_trimmed\_merged\_fastqc.zip - results/historical/trimming/stats/JvS022\_11\_L1\_trimmed\_merged\_fastqc.html - results/historical/trimming/stats/JvS022\_11\_L1\_trimmed\_merged\_fastqc.zip - results/historical/trimming/stats/JvS022\_12\_L1\_trimmed\_merged\_fastqc.html - results/historical/trimming/stats/JvS022\_12\_L1\_trimmed\_merged\_fastqc.zip - results/historical/trimming/stats/JvS022\_74\_L8\_trimmed\_merged\_fastqc.html - results/historical/trimming/stats/JvS022\_74\_L8\_trimmed\_merged\_fastqc.zip - results/historical/trimming/stats/JvS022\_75\_L8\_trimmed\_merged\_fastqc.html - results/historical/trimming/stats/JvS022\_75\_L8\_trimmed\_merged\_fastqc.zip - results/historical/trimming/stats/JvS022\_76\_L8\_trimmed\_merged\_fastqc.html - results/historical/trimming/stats/JvS022\_76\_L8\_trimmed\_merged\_fastqc.zip - results/historical/trimming/stats/JvS022\_77\_L8\_trimmed\_merged\_fastqc.html - results/historical/trimming/stats/JvS022\_77\_L8\_trimmed\_merged\_fastqc.zip - results/historical/trimming/stats/JvS022\_78\_L8\_trimmed\_merged\_fastqc.html - results/historical/trimming/stats/JvS022\_78\_L8\_trimmed\_merged\_fastqc.zip - results/historical/trimming/stats/JvS022\_79\_L8\_trimmed\_merged\_fastqc.html - results/historical/trimming/stats/JvS022\_79\_L8\_trimmed\_merged\_fastqc.zip - results/historical/trimming/stats/JvS022\_80\_L8\_trimmed\_merged\_fastqc.html - results/historical/trimming/stats/JvS022\_80\_L8\_trimmed\_merged\_fastqc.zip - results/historical/trimming/stats/JvS022\_81\_L8\_trimmed\_merged\_fastqc.html - results/historical/trimming/stats/JvS022\_81\_L8\_trimmed\_merged\_fastqc.zip - results/historical/trimming/stats/JvS022\_82\_L8\_trimmed\_merged\_fastqc.html - results/historical/trimming/stats/JvS022\_82\_L8\_trimmed\_merged\_fastqc.zip - results/historical/trimming/stats/JvS022\_83\_L8\_trimmed\_merged\_fastqc.html - results/historical/trimming/stats/JvS022\_83\_L8\_trimmed\_merged\_fastqc.zip - results/historical/trimming/stats/JvS022\_84\_L8\_trimmed\_merged\_fastqc.html - results/historical/trimming/stats/JvS022\_84\_L8\_trimmed\_merged\_fastqc.zip - results/historical/trimming/stats/JvS022\_85\_L8\_trimmed\_merged\_fastqc.html - results/historical/trimming/stats/JvS022\_85\_L8\_trimmed\_merged\_fastqc.zip - results/historical/trimming/stats/JvS008\_08\_L2\_fastp\_report.html - results/historical/trimming/stats/JvS008\_08\_L6\_fastp\_report.html - results/historical/trimming/stats/JvS008\_10\_L2\_fastp\_report.html - results/historical/trimming/stats/JvS008\_10\_L6\_fastp\_report.html - results/historical/trimming/stats/JvS008\_11\_L2\_fastp\_report.html - results/historical/trimming/stats/JvS008\_11\_L6\_fastp\_report.html - results/historical/trimming/stats/JvS009\_09\_L3\_fastp\_report.html - results/historical/trimming/stats/JvS009\_09\_L7\_fastp\_report.html - results/historical/trimming/stats/JvS009\_15\_L3\_fastp\_report.html - results/historical/trimming/stats/JvS009\_15\_L7\_fastp\_report.html - results/historical/trimming/stats/JvS009\_19\_L3\_fastp\_report.html - results/historical/trimming/stats/JvS009\_19\_L7\_fastp\_report.html - results/historical/trimming/stats/JvS022\_01\_L1\_fastp\_report.html - results/historical/trimming/stats/JvS022\_02\_L1\_fastp\_report.html - results/historical/trimming/stats/JvS022\_03\_L1\_fastp\_report.html - results/historical/trimming/stats/JvS022\_04\_L1\_fastp\_report.html - results/historical/trimming/stats/JvS022\_05\_L1\_fastp\_report.html - results/historical/trimming/stats/JvS022\_06\_L1\_fastp\_report.html - results/historical/trimming/stats/JvS022\_07\_L1\_fastp\_report.html - results/historical/trimming/stats/JvS022\_08\_L1\_fastp\_report.html - results/historical/trimming/stats/JvS022\_09\_L1\_fastp\_report.html - results/historical/trimming/stats/JvS022\_10\_L1\_fastp\_report.html - results/historical/trimming/stats/JvS022\_11\_L1\_fastp\_report.html - results/historical/trimming/stats/JvS022\_12\_L1\_fastp\_report.html - results/historical/trimming/stats/JvS022\_74\_L8\_fastp\_report.html - results/historical/trimming/stats/JvS022\_75\_L8\_fastp\_report.html - results/historical/trimming/stats/JvS022\_76\_L8\_fastp\_report.html - results/historical/trimming/stats/JvS022\_77\_L8\_fastp\_report.html - results/historical/trimming/stats/JvS022\_78\_L8\_fastp\_report.html - results/historical/trimming/stats/JvS022\_79\_L8\_fastp\_report.html - results/historical/trimming/stats/JvS022\_80\_L8\_fastp\_report.html - results/historical/trimming/stats/JvS022\_81\_L8\_fastp\_report.html - results/historical/trimming/stats/JvS022\_82\_L8\_fastp\_report.html - results/historical/trimming/stats/JvS022\_83\_L8\_fastp\_report.html - results/historical/trimming/stats/JvS022\_84\_L8\_fastp\_report.html - results/historical/trimming/stats/JvS022\_85\_L8\_fastp\_report.html - results/historical/trimming/stats/JvS008\_08\_L2\_R1\_unmerged\_fastqc.html - results/historical/trimming/stats/JvS008\_08\_L2\_R1\_unmerged\_fastqc.zip - results/historical/trimming/stats/JvS008\_08\_L2\_R2\_unmerged\_fastqc.html - results/historical/trimming/stats/JvS008\_08\_L2\_R2\_unmerged\_fastqc.zip - results/historical/trimming/stats/JvS008\_08\_L6\_R1\_unmerged\_fastqc.html - results/historical/trimming/stats/JvS008\_08\_L6\_R1\_unmerged\_fastqc.zip - results/historical/trimming/stats/JvS008\_08\_L6\_R2\_unmerged\_fastqc.html - results/historical/trimming/stats/JvS008\_08\_L6\_R2\_unmerged\_fastqc.zip - results/historical/trimming/stats/JvS008\_10\_L2\_R1\_unmerged\_fastqc.html - results/historical/trimming/stats/JvS008\_10\_L2\_R1\_unmerged\_fastqc.zip - results/historical/trimming/stats/JvS008\_10\_L2\_R2\_unmerged\_fastqc.html - results/historical/trimming/stats/JvS008\_10\_L2\_R2\_unmerged\_fastqc.zip - results/historical/trimming/stats/JvS008\_10\_L6\_R1\_unmerged\_fastqc.html - results/historical/trimming/stats/JvS008\_10\_L6\_R1\_unmerged\_fastqc.zip - results/historical/trimming/stats/JvS008\_10\_L6\_R2\_unmerged\_fastqc.html - results/historical/trimming/stats/JvS008\_10\_L6\_R2\_unmerged\_fastqc.zip - results/historical/trimming/stats/JvS008\_11\_L2\_R1\_unmerged\_fastqc.html - results/historical/trimming/stats/JvS008\_11\_L2\_R1\_unmerged\_fastqc.zip - results/historical/trimming/stats/JvS008\_11\_L2\_R2\_unmerged\_fastqc.html - results/historical/trimming/stats/JvS008\_11\_L2\_R2\_unmerged\_fastqc.zip - results/historical/trimming/stats/JvS008\_11\_L6\_R1\_unmerged\_fastqc.html - results/historical/trimming/stats/JvS008\_11\_L6\_R1\_unmerged\_fastqc.zip - results/historical/trimming/stats/JvS008\_11\_L6\_R2\_unmerged\_fastqc.html - results/historical/trimming/stats/JvS008\_11\_L6\_R2\_unmerged\_fastqc.zip - results/historical/trimming/stats/JvS009\_09\_L3\_R1\_unmerged\_fastqc.html - results/historical/trimming/stats/JvS009\_09\_L3\_R1\_unmerged\_fastqc.zip - results/historical/trimming/stats/JvS009\_09\_L3\_R2\_unmerged\_fastqc.html - results/historical/trimming/stats/JvS009\_09\_L3\_R2\_unmerged\_fastqc.zip - results/historical/trimming/stats/JvS009\_09\_L7\_R1\_unmerged\_fastqc.html - results/historical/trimming/stats/JvS009\_09\_L7\_R1\_unmerged\_fastqc.zip - results/historical/trimming/stats/JvS009\_09\_L7\_R2\_unmerged\_fastqc.html - results/historical/trimming/stats/JvS009\_09\_L7\_R2\_unmerged\_fastqc.zip - results/historical/trimming/stats/JvS009\_15\_L3\_R1\_unmerged\_fastqc.html - results/historical/trimming/stats/JvS009\_15\_L3\_R1\_unmerged\_fastqc.zip - results/historical/trimming/stats/JvS009\_15\_L3\_R2\_unmerged\_fastqc.html - results/historical/trimming/stats/JvS009\_15\_L3\_R2\_unmerged\_fastqc.zip - results/historical/trimming/stats/JvS009\_15\_L7\_R1\_unmerged\_fastqc.html - results/historical/trimming/stats/JvS009\_15\_L7\_R1\_unmerged\_fastqc.zip - results/historical/trimming/stats/JvS009\_15\_L7\_R2\_unmerged\_fastqc.html - results/historical/trimming/stats/JvS009\_15\_L7\_R2\_unmerged\_fastqc.zip - results/historical/trimming/stats/JvS009\_19\_L3\_R1\_unmerged\_fastqc.html - results/historical/trimming/stats/JvS009\_19\_L3\_R1\_unmerged\_fastqc.zip - results/historical/trimming/stats/JvS009\_19\_L3\_R2\_unmerged\_fastqc.html - results/historical/trimming/stats/JvS009\_19\_L3\_R2\_unmerged\_fastqc.zip - results/historical/trimming/stats/JvS009\_19\_L7\_R1\_unmerged\_fastqc.html - results/historical/trimming/stats/JvS009\_19\_L7\_R1\_unmerged\_fastqc.zip - results/historical/trimming/stats/JvS009\_19\_L7\_R2\_unmerged\_fastqc.html - results/historical/trimming/stats/JvS009\_19\_L7\_R2\_unmerged\_fastqc.zip - results/historical/trimming/stats/JvS022\_01\_L1\_R1\_unmerged\_fastqc.html - results/historical/trimming/stats/JvS022\_01\_L1\_R1\_unmerged\_fastqc.zip - results/historical/trimming/stats/JvS022\_01\_L1\_R2\_unmerged\_fastqc.html - results/historical/trimming/stats/JvS022\_01\_L1\_R2\_unmerged\_fastqc.zip - results/historical/trimming/stats/JvS022\_02\_L1\_R1\_unmerged\_fastqc.html - results/historical/trimming/stats/JvS022\_02\_L1\_R1\_unmerged\_fastqc.zip - results/historical/trimming/stats/JvS022\_02\_L1\_R2\_unmerged\_fastqc.html - results/historical/trimming/stats/JvS022\_02\_L1\_R2\_unmerged\_fastqc.zip - results/historical/trimming/stats/JvS022\_03\_L1\_R1\_unmerged\_fastqc.html - results/historical/trimming/stats/JvS022\_03\_L1\_R1\_unmerged\_fastqc.zip - results/historical/trimming/stats/JvS022\_03\_L1\_R2\_unmerged\_fastqc.html - results/historical/trimming/stats/JvS022\_03\_L1\_R2\_unmerged\_fastqc.zip - results/historical/trimming/stats/JvS022\_04\_L1\_R1\_unmerged\_fastqc.html - results/historical/trimming/stats/JvS022\_04\_L1\_R1\_unmerged\_fastqc.zip - results/historical/trimming/stats/JvS022\_04\_L1\_R2\_unmerged\_fastqc.html - results/historical/trimming/stats/JvS022\_04\_L1\_R2\_unmerged\_fastqc.zip - results/historical/trimming/stats/JvS022\_05\_L1\_R1\_unmerged\_fastqc.html - results/historical/trimming/stats/JvS022\_05\_L1\_R1\_unmerged\_fastqc.zip - results/historical/trimming/stats/JvS022\_05\_L1\_R2\_unmerged\_fastqc.html - results/historical/trimming/stats/JvS022\_05\_L1\_R2\_unmerged\_fastqc.zip - results/historical/trimming/stats/JvS022\_06\_L1\_R1\_unmerged\_fastqc.html - results/historical/trimming/stats/JvS022\_06\_L1\_R1\_unmerged\_fastqc.zip - results/historical/trimming/stats/JvS022\_06\_L1\_R2\_unmerged\_fastqc.html - results/historical/trimming/stats/JvS022\_06\_L1\_R2\_unmerged\_fastqc.zip - results/historical/trimming/stats/JvS022\_07\_L1\_R1\_unmerged\_fastqc.html - results/historical/trimming/stats/JvS022\_07\_L1\_R1\_unmerged\_fastqc.zip - results/historical/trimming/stats/JvS022\_07\_L1\_R2\_unmerged\_fastqc.html - results/historical/trimming/stats/JvS022\_07\_L1\_R2\_unmerged\_fastqc.zip - results/historical/trimming/stats/JvS022\_08\_L1\_R1\_unmerged\_fastqc.html - results/historical/trimming/stats/JvS022\_08\_L1\_R1\_unmerged\_fastqc.zip - results/historical/trimming/stats/JvS022\_08\_L1\_R2\_unmerged\_fastqc.html - results/historical/trimming/stats/JvS022\_08\_L1\_R2\_unmerged\_fastqc.zip - results/historical/trimming/stats/JvS022\_09\_L1\_R1\_unmerged\_fastqc.html - results/historical/trimming/stats/JvS022\_09\_L1\_R1\_unmerged\_fastqc.zip - results/historical/trimming/stats/JvS022\_09\_L1\_R2\_unmerged\_fastqc.html - results/historical/trimming/stats/JvS022\_09\_L1\_R2\_unmerged\_fastqc.zip - results/historical/trimming/stats/JvS022\_10\_L1\_R1\_unmerged\_fastqc.html - results/historical/trimming/stats/JvS022\_10\_L1\_R1\_unmerged\_fastqc.zip - results/historical/trimming/stats/JvS022\_10\_L1\_R2\_unmerged\_fastqc.html - results/historical/trimming/stats/JvS022\_10\_L1\_R2\_unmerged\_fastqc.zip - results/historical/trimming/stats/JvS022\_11\_L1\_R1\_unmerged\_fastqc.html - results/historical/trimming/stats/JvS022\_11\_L1\_R1\_unmerged\_fastqc.zip - results/historical/trimming/stats/JvS022\_11\_L1\_R2\_unmerged\_fastqc.html - results/historical/trimming/stats/JvS022\_11\_L1\_R2\_unmerged\_fastqc.zip - results/historical/trimming/stats/JvS022\_12\_L1\_R1\_unmerged\_fastqc.html - results/historical/trimming/stats/JvS022\_12\_L1\_R1\_unmerged\_fastqc.zip - results/historical/trimming/stats/JvS022\_12\_L1\_R2\_unmerged\_fastqc.html - results/historical/trimming/stats/JvS022\_12\_L1\_R2\_unmerged\_fastqc.zip - results/historical/trimming/stats/JvS022\_74\_L8\_R1\_unmerged\_fastqc.html - results/historical/trimming/stats/JvS022\_74\_L8\_R1\_unmerged\_fastqc.zip - results/historical/trimming/stats/JvS022\_74\_L8\_R2\_unmerged\_fastqc.html - results/historical/trimming/stats/JvS022\_74\_L8\_R2\_unmerged\_fastqc.zip - results/historical/trimming/stats/JvS022\_75\_L8\_R1\_unmerged\_fastqc.html - results/historical/trimming/stats/JvS022\_75\_L8\_R1\_unmerged\_fastqc.zip - results/historical/trimming/stats/JvS022\_75\_L8\_R2\_unmerged\_fastqc.html - results/historical/trimming/stats/JvS022\_75\_L8\_R2\_unmerged\_fastqc.zip - results/historical/trimming/stats/JvS022\_76\_L8\_R1\_unmerged\_fastqc.html - results/historical/trimming/stats/JvS022\_76\_L8\_R1\_unmerged\_fastqc.zip - results/historical/trimming/stats/JvS022\_76\_L8\_R2\_unmerged\_fastqc.html - results/historical/trimming/stats/JvS022\_76\_L8\_R2\_unmerged\_fastqc.zip - results/historical/trimming/stats/JvS022\_77\_L8\_R1\_unmerged\_fastqc.html - results/historical/trimming/stats/JvS022\_77\_L8\_R1\_unmerged\_fastqc.zip - results/historical/trimming/stats/JvS022\_77\_L8\_R2\_unmerged\_fastqc.html - results/historical/trimming/stats/JvS022\_77\_L8\_R2\_unmerged\_fastqc.zip - results/historical/trimming/stats/JvS022\_78\_L8\_R1\_unmerged\_fastqc.html - results/historical/trimming/stats/JvS022\_78\_L8\_R1\_unmerged\_fastqc.zip - results/historical/trimming/stats/JvS022\_78\_L8\_R2\_unmerged\_fastqc.html - results/historical/trimming/stats/JvS022\_78\_L8\_R2\_unmerged\_fastqc.zip - results/historical/trimming/stats/JvS022\_79\_L8\_R1\_unmerged\_fastqc.html - results/historical/trimming/stats/JvS022\_79\_L8\_R1\_unmerged\_fastqc.zip - results/historical/trimming/stats/JvS022\_79\_L8\_R2\_unmerged\_fastqc.html - results/historical/trimming/stats/JvS022\_79\_L8\_R2\_unmerged\_fastqc.zip - results/historical/trimming/stats/JvS022\_80\_L8\_R1\_unmerged\_fastqc.html - results/historical/trimming/stats/JvS022\_80\_L8\_R1\_unmerged\_fastqc.zip - results/historical/trimming/stats/JvS022\_80\_L8\_R2\_unmerged\_fastqc.html - results/historical/trimming/stats/JvS022\_80\_L8\_R2\_unmerged\_fastqc.zip - results/historical/trimming/stats/JvS022\_81\_L8\_R1\_unmerged\_fastqc.html - results/historical/trimming/stats/JvS022\_81\_L8\_R1\_unmerged\_fastqc.zip - results/historical/trimming/stats/JvS022\_81\_L8\_R2\_unmerged\_fastqc.html - results/historical/trimming/stats/JvS022\_81\_L8\_R2\_unmerged\_fastqc.zip - results/historical/trimming/stats/JvS022\_82\_L8\_R1\_unmerged\_fastqc.html - results/historical/trimming/stats/JvS022\_82\_L8\_R1\_unmerged\_fastqc.zip - results/historical/trimming/stats/JvS022\_82\_L8\_R2\_unmerged\_fastqc.html - results/historical/trimming/stats/JvS022\_82\_L8\_R2\_unmerged\_fastqc.zip - results/historical/trimming/stats/JvS022\_83\_L8\_R1\_unmerged\_fastqc.html - results/historical/trimming/stats/JvS022\_83\_L8\_R1\_unmerged\_fastqc.zip - results/historical/trimming/stats/JvS022\_83\_L8\_R2\_unmerged\_fastqc.html - results/historical/trimming/stats/JvS022\_83\_L8\_R2\_unmerged\_fastqc.zip - results/historical/trimming/stats/JvS022\_84\_L8\_R1\_unmerged\_fastqc.html - results/historical/trimming/stats/JvS022\_84\_L8\_R1\_unmerged\_fastqc.zip - results/historical/trimming/stats/JvS022\_84\_L8\_R2\_unmerged\_fastqc.html - results/historical/trimming/stats/JvS022\_84\_L8\_R2\_unmerged\_fastqc.zip - results/historical/trimming/stats/JvS022\_85\_L8\_R1\_unmerged\_fastqc.html - results/historical/trimming/stats/JvS022\_85\_L8\_R1\_unmerged\_fastqc.zip - results/historical/trimming/stats/JvS022\_85\_L8\_R2\_unmerged\_fastqc.html - results/historical/trimming/stats/JvS022\_85\_L8\_R2\_unmerged\_fastqc.zip | |
| Output files | |
| - results/historical/trimming/stats/multiqc/multiqc\_report.html | |
| Container image | |
| docker://quay.io/biocontainers/multiqc:1.9--pyh9f0ad1d\_0 | |
| Code | |
| ``` 1 2 ``` \| ``` multiqc -f {params.indir} -o {params.outdir} 2> {log} ``` | |

###### Rule fastqc\_historical\_merged

×

Rule properties

|  |  |
| --- | --- |
| Jobs | 36 |
| Input files | |
| - results/historical/trimming/{sample}\_{index}\_{lane}\_trimmed\_merged.fastq.gz | |
| Output files | |
| - results/historical/trimming/stats/{sample,[A-Za-z0-9]+}\_{index}\_{lane}\_trimmed\_merged\_fastqc.html - results/historical/trimming/stats/{sample,[A-Za-z0-9]+}\_{index}\_{lane}\_trimmed\_merged\_fastqc.zip - results/historical/trimming/stats/{sample,[A-Za-z0-9]+}\_{index}\_{lane}\_trimmed\_merged\_fastqc | |
| Container image | |
| docker://biocontainers/fastqc:v0.11.9\_cv7 | |
| Code | |
| ``` 1 2 ``` \| ``` fastqc -o {params.dir} -t {threads} --extract {input} 2> {log} ``` | |

###### Rule fastp\_historical

×

Rule properties

|  |  |
| --- | --- |
| Jobs | 36 |
| Input files | |
| - data/raw\_reads\_symlinks/historical/{sample}\_{index}\_{lane}\_R1.fastq.gz - data/raw\_reads\_symlinks/historical/{sample}\_{index}\_{lane}\_R2.fastq.gz | |
| Output files | |
| - results/historical/trimming/{sample,[A-Za-z0-9]+}\_{index}\_{lane}\_R1\_unmerged.fastq.gz - results/historical/trimming/{sample,[A-Za-z0-9]+}\_{index}\_{lane}\_R2\_unmerged.fastq.gz - results/historical/trimming/{sample,[A-Za-z0-9]+}\_{index}\_{lane}\_trimmed\_merged.fastq.gz - results/historical/trimming/stats/{sample,[A-Za-z0-9]+}\_{index}\_{lane}\_fastp\_report.html - results/modern/trimming/stats/{sample,[A-Za-z0-9]+}\_{index}\_{lane}\_fastp\_report.json | |
| Container image | |
| docker://quay.io/biocontainers/fastp:0.22.0--h2e03b76\_0 | |
| Code | |
| ``` 1 2 ``` \| ``` fastp -i {input.R1} -I {input.R2} -p -c --merge --merged_out={output.merged} -o {output.R1_un} -O {output.R2_un} -h {output.html} -j {output.json} -R '{params.report}' -w {threads} -l {params.readlength} 2> {log} ``` | |

###### Rule fastqc\_historical\_unmerged

×

Rule properties

|  |  |
| --- | --- |
| Jobs | 72 |
| Input files | |
| - results/historical/trimming/{sample}\_{index}\_{lane}\_R{nr}\_unmerged.fastq.gz | |
| Output files | |
| - results/historical/trimming/stats/{sample,[A-Za-z0-9]+}\_{index}\_{lane}\_R{nr}\_unmerged\_fastqc.html - results/historical/trimming/stats/{sample,[A-Za-z0-9]+}\_{index}\_{lane}\_R{nr}\_unmerged\_fastqc.zip - results/historical/trimming/stats/{sample,[A-Za-z0-9]+}\_{index}\_{lane}\_R{nr}\_unmerged\_fastqc | |
| Container image | |
| docker://biocontainers/fastqc:v0.11.9\_cv7 | |
| Code | |
| ``` 1 2 3 4 5 6 7 8 9 ``` \| ``` if [ -s {input} ] then fastqc -o {params.dir} -t {threads} --extract {input} 2> {log} else mkdir -p {output.dir} && touch {output.html} && touch {output.zip} && echo "No reads in {input} >> {log}" fi ``` | |

###### Rule multiqc\_modern\_raw

×

Rule properties

|  |  |
| --- | --- |
| Jobs | 1 |
| Input files | |
| - data/raw\_reads\_symlinks/modern/stats/JvS033\_01\_L7\_R1\_fastqc.html - data/raw\_reads\_symlinks/modern/stats/JvS033\_01\_L7\_R2\_fastqc.html - data/raw\_reads\_symlinks/modern/stats/JvS034\_02\_L8\_R1\_fastqc.html - data/raw\_reads\_symlinks/modern/stats/JvS034\_02\_L8\_R2\_fastqc.html - data/raw\_reads\_symlinks/modern/stats/JvS035\_14\_L7\_R1\_fastqc.html - data/raw\_reads\_symlinks/modern/stats/JvS035\_14\_L7\_R2\_fastqc.html - data/raw\_reads\_symlinks/modern/stats/JvS033\_01\_L7\_R1\_fastqc.zip - data/raw\_reads\_symlinks/modern/stats/JvS033\_01\_L7\_R2\_fastqc.zip - data/raw\_reads\_symlinks/modern/stats/JvS034\_02\_L8\_R1\_fastqc.zip - data/raw\_reads\_symlinks/modern/stats/JvS034\_02\_L8\_R2\_fastqc.zip - data/raw\_reads\_symlinks/modern/stats/JvS035\_14\_L7\_R1\_fastqc.zip - data/raw\_reads\_symlinks/modern/stats/JvS035\_14\_L7\_R2\_fastqc.zip | |
| Output files | |
| - data/raw\_reads\_symlinks/modern/stats/multiqc/multiqc\_report.html | |
| Container image | |
| docker://quay.io/biocontainers/multiqc:1.9--pyh9f0ad1d\_0 | |
| Code | |
| ``` 1 2 ``` \| ``` multiqc -f {params.indir} -o {params.outdir} 2> {log} ``` | |

###### Rule fastqc\_modern\_raw

×

Rule properties

|  |  |
| --- | --- |
| Jobs | 6 |
| Input files | |
| - data/raw\_reads\_symlinks/modern/{sample}\_{index}\_{lane}\_R{nr}.fastq.gz | |
| Output files | |
| - data/raw\_reads\_symlinks/modern/stats/{sample,[A-Za-z0-9]+}\_{index}\_{lane}\_R{nr}\_fastqc.html - data/raw\_reads\_symlinks/modern/stats/{sample,[A-Za-z0-9]+}\_{index}\_{lane}\_R{nr}\_fastqc.zip - data/raw\_reads\_symlinks/modern/stats/{sample,[A-Za-z0-9]+}\_{index}\_{lane}\_R{nr}\_fastqc | |
| Container image | |
| docker://biocontainers/fastqc:v0.11.9\_cv7 | |
| Code | |
| ``` 1 2 ``` \| ``` fastqc -o {params.dir} -t {threads} --extract {input.fastq} 2> {log} ``` | |

###### Rule fastq\_modern\_symbolic\_links

×

Rule properties

|  |  |
| --- | --- |
| Jobs | 3 |
| Input files | |
| - config/rhino\_3\_modern\_samples.txt | |
| Output files | |
| - data/raw\_reads\_symlinks/modern/{sample,[A-Za-z0-9]+}\_{index}\_{lane}\_R1.fastq.gz - data/raw\_reads\_symlinks/modern/{sample,[A-Za-z0-9]+}\_{index}\_{lane}\_R2.fastq.gz | |

###### Rule multiqc\_modern\_trimmed

×

Rule properties

|  |  |
| --- | --- |
| Jobs | 1 |
| Input files | |
| - results/modern/trimming/stats/JvS033\_01\_L7\_R1\_trimmed\_fastqc.html - results/modern/trimming/stats/JvS033\_01\_L7\_R1\_trimmed\_fastqc.zip - results/modern/trimming/stats/JvS033\_01\_L7\_R2\_trimmed\_fastqc.html - results/modern/trimming/stats/JvS033\_01\_L7\_R2\_trimmed\_fastqc.zip - results/modern/trimming/stats/JvS034\_02\_L8\_R1\_trimmed\_fastqc.html - results/modern/trimming/stats/JvS034\_02\_L8\_R1\_trimmed\_fastqc.zip - results/modern/trimming/stats/JvS034\_02\_L8\_R2\_trimmed\_fastqc.html - results/modern/trimming/stats/JvS034\_02\_L8\_R2\_trimmed\_fastqc.zip - results/modern/trimming/stats/JvS035\_14\_L7\_R1\_trimmed\_fastqc.html - results/modern/trimming/stats/JvS035\_14\_L7\_R1\_trimmed\_fastqc.zip - results/modern/trimming/stats/JvS035\_14\_L7\_R2\_trimmed\_fastqc.html - results/modern/trimming/stats/JvS035\_14\_L7\_R2\_trimmed\_fastqc.zip - results/modern/trimming/stats/JvS033\_01\_L7\_fastp\_report.html - results/modern/trimming/stats/JvS034\_02\_L8\_fastp\_report.html - results/modern/trimming/stats/JvS035\_14\_L7\_fastp\_report.html | |
| Output files | |
| - results/modern/trimming/stats/multiqc/multiqc\_report.html | |
| Container image | |
| docker://quay.io/biocontainers/multiqc:1.9--pyh9f0ad1d\_0 | |
| Code | |
| ``` 1 2 ``` \| ``` multiqc -f {params.indir} -o {params.outdir} 2> {log} ``` | |

###### Rule fastqc\_modern\_trimmed

×

Rule properties

|  |  |
| --- | --- |
| Jobs | 6 |
| Input files | |
| - results/modern/trimming/{sample}\_{index}\_{lane}\_R{nr}\_trimmed.fastq.gz | |
| Output files | |
| - results/modern/trimming/stats/{sample,[A-Za-z0-9]+}\_{index}\_{lane}\_R{nr}\_trimmed\_fastqc.html - results/modern/trimming/stats/{sample,[A-Za-z0-9]+}\_{index}\_{lane}\_R{nr}\_trimmed\_fastqc.zip - results/modern/trimming/stats/{sample,[A-Za-z0-9]+}\_{index}\_{lane}\_R{nr}\_trimmed\_fastqc | |
| Container image | |
| docker://biocontainers/fastqc:v0.11.9\_cv7 | |
| Code | |
| ``` 1 2 ``` \| ``` fastqc -o {params.dir} -t {threads} --extract {input} 2> {log} ``` | |

###### Rule fastp\_modern

×

Rule properties

|  |  |
| --- | --- |
| Jobs | 3 |
| Input files | |
| - data/raw\_reads\_symlinks/modern/{sample}\_{index}\_{lane}\_R1.fastq.gz - data/raw\_reads\_symlinks/modern/{sample}\_{index}\_{lane}\_R2.fastq.gz | |
| Output files | |
| - results/modern/trimming/{sample,[A-Za-z0-9]+}\_{index}\_{lane}\_R1\_trimmed.fastq.gz - results/modern/trimming/{sample,[A-Za-z0-9]+}\_{index}\_{lane}\_R2\_trimmed.fastq.gz - results/modern/trimming/stats/{sample,[A-Za-z0-9]+}\_{index}\_{lane}\_fastp\_report.html - results/modern/trimming/stats/{sample,[A-Za-z0-9]+}\_{index}\_{lane}\_fastp\_report.json | |
| Container image | |
| docker://quay.io/biocontainers/fastp:0.22.0--h2e03b76\_0 | |
| Code | |
| ``` 1 2 ``` \| ``` fastp -i {input.R1} -I {input.R2} -p -c -o {output.R1_trimmed} -O {output.R2_trimmed} -h {output.html} -j {output.json} -R '{params.report}' -w {threads} -l {params.readlength} 2> {log} ``` | |

###### Rule historical\_raw\_bam\_multiqc

×

Rule properties

|  |  |
| --- | --- |
| Jobs | 1 |
| Input files | |
| - results/historical/mapping/GCF\_000283155.1\_CerSimSim1.0\_genomic.Sc9M7eS\_2\_HRSCAF\_41/stats/bams\_sorted/JvS008\_08\_L2.sorted.bam.stats.txt - results/historical/mapping/GCF\_000283155.1\_CerSimSim1.0\_genomic.Sc9M7eS\_2\_HRSCAF\_41/stats/bams\_sorted/JvS008\_08\_L6.sorted.bam.stats.txt - results/historical/mapping/GCF\_000283155.1\_CerSimSim1.0\_genomic.Sc9M7eS\_2\_HRSCAF\_41/stats/bams\_sorted/JvS008\_10\_L2.sorted.bam.stats.txt - results/historical/mapping/GCF\_000283155.1\_CerSimSim1.0\_genomic.Sc9M7eS\_2\_HRSCAF\_41/stats/bams\_sorted/JvS008\_10\_L6.sorted.bam.stats.txt - results/historical/mapping/GCF\_000283155.1\_CerSimSim1.0\_genomic.Sc9M7eS\_2\_HRSCAF\_41/stats/bams\_sorted/JvS008\_11\_L2.sorted.bam.stats.txt - results/historical/mapping/GCF\_000283155.1\_CerSimSim1.0\_genomic.Sc9M7eS\_2\_HRSCAF\_41/stats/bams\_sorted/JvS008\_11\_L6.sorted.bam.stats.txt - results/historical/mapping/GCF\_000283155.1\_CerSimSim1.0\_genomic.Sc9M7eS\_2\_HRSCAF\_41/stats/bams\_sorted/JvS009\_09\_L3.sorted.bam.stats.txt - results/historical/mapping/GCF\_000283155.1\_CerSimSim1.0\_genomic.Sc9M7eS\_2\_HRSCAF\_41/stats/bams\_sorted/JvS009\_09\_L7.sorted.bam.stats.txt - results/historical/mapping/GCF\_000283155.1\_CerSimSim1.0\_genomic.Sc9M7eS\_2\_HRSCAF\_41/stats/bams\_sorted/JvS009\_15\_L3.sorted.bam.stats.txt - results/historical/mapping/GCF\_000283155.1\_CerSimSim1.0\_genomic.Sc9M7eS\_2\_HRSCAF\_41/stats/bams\_sorted/JvS009\_15\_L7.sorted.bam.stats.txt - results/historical/mapping/GCF\_000283155.1\_CerSimSim1.0\_genomic.Sc9M7eS\_2\_HRSCAF\_41/stats/bams\_sorted/JvS009\_19\_L3.sorted.bam.stats.txt - results/historical/mapping/GCF\_000283155.1\_CerSimSim1.0\_genomic.Sc9M7eS\_2\_HRSCAF\_41/stats/bams\_sorted/JvS009\_19\_L7.sorted.bam.stats.txt - results/historical/mapping/GCF\_000283155.1\_CerSimSim1.0\_genomic.Sc9M7eS\_2\_HRSCAF\_41/stats/bams\_sorted/JvS022\_01\_L1.sorted.bam.stats.txt - results/historical/mapping/GCF\_000283155.1\_CerSimSim1.0\_genomic.Sc9M7eS\_2\_HRSCAF\_41/stats/bams\_sorted/JvS022\_02\_L1.sorted.bam.stats.txt - results/historical/mapping/GCF\_000283155.1\_CerSimSim1.0\_genomic.Sc9M7eS\_2\_HRSCAF\_41/stats/bams\_sorted/JvS022\_03\_L1.sorted.bam.stats.txt - results/historical/mapping/GCF\_000283155.1\_CerSimSim1.0\_genomic.Sc9M7eS\_2\_HRSCAF\_41/stats/bams\_sorted/JvS022\_04\_L1.sorted.bam.stats.txt - results/historical/mapping/GCF\_000283155.1\_CerSimSim1.0\_genomic.Sc9M7eS\_2\_HRSCAF\_41/stats/bams\_sorted/JvS022\_05\_L1.sorted.bam.stats.txt - results/historical/mapping/GCF\_000283155.1\_CerSimSim1.0\_genomic.Sc9M7eS\_2\_HRSCAF\_41/stats/bams\_sorted/JvS022\_06\_L1.sorted.bam.stats.txt - results/historical/mapping/GCF\_000283155.1\_CerSimSim1.0\_genomic.Sc9M7eS\_2\_HRSCAF\_41/stats/bams\_sorted/JvS022\_07\_L1.sorted.bam.stats.txt - results/historical/mapping/GCF\_000283155.1\_CerSimSim1.0\_genomic.Sc9M7eS\_2\_HRSCAF\_41/stats/bams\_sorted/JvS022\_08\_L1.sorted.bam.stats.txt - results/historical/mapping/GCF\_000283155.1\_CerSimSim1.0\_genomic.Sc9M7eS\_2\_HRSCAF\_41/stats/bams\_sorted/JvS022\_09\_L1.sorted.bam.stats.txt - results/historical/mapping/GCF\_000283155.1\_CerSimSim1.0\_genomic.Sc9M7eS\_2\_HRSCAF\_41/stats/bams\_sorted/JvS022\_10\_L1.sorted.bam.stats.txt - results/historical/mapping/GCF\_000283155.1\_CerSimSim1.0\_genomic.Sc9M7eS\_2\_HRSCAF\_41/stats/bams\_sorted/JvS022\_11\_L1.sorted.bam.stats.txt - results/historical/mapping/GCF\_000283155.1\_CerSimSim1.0\_genomic.Sc9M7eS\_2\_HRSCAF\_41/stats/bams\_sorted/JvS022\_12\_L1.sorted.bam.stats.txt - results/historical/mapping/GCF\_000283155.1\_CerSimSim1.0\_genomic.Sc9M7eS\_2\_HRSCAF\_41/stats/bams\_sorted/JvS022\_74\_L8.sorted.bam.stats.txt - results/historical/mapping/GCF\_000283155.1\_CerSimSim1.0\_genomic.Sc9M7eS\_2\_HRSCAF\_41/stats/bams\_sorted/JvS022\_75\_L8.sorted.bam.stats.txt - results/historical/mapping/GCF\_000283155.1\_CerSimSim1.0\_genomic.Sc9M7eS\_2\_HRSCAF\_41/stats/bams\_sorted/JvS022\_76\_L8.sorted.bam.stats.txt - results/historical/mapping/GCF\_000283155.1\_CerSimSim1.0\_genomic.Sc9M7eS\_2\_HRSCAF\_41/stats/bams\_sorted/JvS022\_77\_L8.sorted.bam.stats.txt - results/historical/mapping/GCF\_000283155.1\_CerSimSim1.0\_genomic.Sc9M7eS\_2\_HRSCAF\_41/stats/bams\_sorted/JvS022\_78\_L8.sorted.bam.stats.txt - results/historical/mapping/GCF\_000283155.1\_CerSimSim1.0\_genomic.Sc9M7eS\_2\_HRSCAF\_41/stats/bams\_sorted/JvS022\_79\_L8.sorted.bam.stats.txt - results/historical/mapping/GCF\_000283155.1\_CerSimSim1.0\_genomic.Sc9M7eS\_2\_HRSCAF\_41/stats/bams\_sorted/JvS022\_80\_L8.sorted.bam.stats.txt - results/historical/mapping/GCF\_000283155.1\_CerSimSim1.0\_genomic.Sc9M7eS\_2\_HRSCAF\_41/stats/bams\_sorted/JvS022\_81\_L8.sorted.bam.stats.txt - results/historical/mapping/GCF\_000283155.1\_CerSimSim1.0\_genomic.Sc9M7eS\_2\_HRSCAF\_41/stats/bams\_sorted/JvS022\_82\_L8.sorted.bam.stats.txt - results/historical/mapping/GCF\_000283155.1\_CerSimSim1.0\_genomic.Sc9M7eS\_2\_HRSCAF\_41/stats/bams\_sorted/JvS022\_83\_L8.sorted.bam.stats.txt - results/historical/mapping/GCF\_000283155.1\_CerSimSim1.0\_genomic.Sc9M7eS\_2\_HRSCAF\_41/stats/bams\_sorted/JvS022\_84\_L8.sorted.bam.stats.txt - results/historical/mapping/GCF\_000283155.1\_CerSimSim1.0\_genomic.Sc9M7eS\_2\_HRSCAF\_41/stats/bams\_sorted/JvS022\_85\_L8.sorted.bam.stats.txt - results/historical/mapping/GCF\_000283155.1\_CerSimSim1.0\_genomic.Sc9M7eS\_2\_HRSCAF\_41/stats/bams\_sorted/JvS008\_08\_L2.sorted.bam.qualimap/qualimapReport.html - results/historical/mapping/GCF\_000283155.1\_CerSimSim1.0\_genomic.Sc9M7eS\_2\_HRSCAF\_41/stats/bams\_sorted/JvS008\_08\_L6.sorted.bam.qualimap/qualimapReport.html - results/historical/mapping/GCF\_000283155.1\_CerSimSim1.0\_genomic.Sc9M7eS\_2\_HRSCAF\_41/stats/bams\_sorted/JvS008\_10\_L2.sorted.bam.qualimap/qualimapReport.html - results/historical/mapping/GCF\_000283155.1\_CerSimSim1.0\_genomic.Sc9M7eS\_2\_HRSCAF\_41/stats/bams\_sorted/JvS008\_10\_L6.sorted.bam.qualimap/qualimapReport.html - results/historical/mapping/GCF\_000283155.1\_CerSimSim1.0\_genomic.Sc9M7eS\_2\_HRSCAF\_41/stats/bams\_sorted/JvS008\_11\_L2.sorted.bam.qualimap/qualimapReport.html - results/historical/mapping/GCF\_000283155.1\_CerSimSim1.0\_genomic.Sc9M7eS\_2\_HRSCAF\_41/stats/bams\_sorted/JvS008\_11\_L6.sorted.bam.qualimap/qualimapReport.html - results/historical/mapping/GCF\_000283155.1\_CerSimSim1.0\_genomic.Sc9M7eS\_2\_HRSCAF\_41/stats/bams\_sorted/JvS009\_09\_L3.sorted.bam.qualimap/qualimapReport.html - results/historical/mapping/GCF\_000283155.1\_CerSimSim1.0\_genomic.Sc9M7eS\_2\_HRSCAF\_41/stats/bams\_sorted/JvS009\_09\_L7.sorted.bam.qualimap/qualimapReport.html - results/historical/mapping/GCF\_000283155.1\_CerSimSim1.0\_genomic.Sc9M7eS\_2\_HRSCAF\_41/stats/bams\_sorted/JvS009\_15\_L3.sorted.bam.qualimap/qualimapReport.html - results/historical/mapping/GCF\_000283155.1\_CerSimSim1.0\_genomic.Sc9M7eS\_2\_HRSCAF\_41/stats/bams\_sorted/JvS009\_15\_L7.sorted.bam.qualimap/qualimapReport.html - results/historical/mapping/GCF\_000283155.1\_CerSimSim1.0\_genomic.Sc9M7eS\_2\_HRSCAF\_41/stats/bams\_sorted/JvS009\_19\_L3.sorted.bam.qualimap/qualimapReport.html - results/historical/mapping/GCF\_000283155.1\_CerSimSim1.0\_genomic.Sc9M7eS\_2\_HRSCAF\_41/stats/bams\_sorted/JvS009\_19\_L7.sorted.bam.qualimap/qualimapReport.html - results/historical/mapping/GCF\_000283155.1\_CerSimSim1.0\_genomic.Sc9M7eS\_2\_HRSCAF\_41/stats/bams\_sorted/JvS022\_01\_L1.sorted.bam.qualimap/qualimapReport.html - results/historical/mapping/GCF\_000283155.1\_CerSimSim1.0\_genomic.Sc9M7eS\_2\_HRSCAF\_41/stats/bams\_sorted/JvS022\_02\_L1.sorted.bam.qualimap/qualimapReport.html - results/historical/mapping/GCF\_000283155.1\_CerSimSim1.0\_genomic.Sc9M7eS\_2\_HRSCAF\_41/stats/bams\_sorted/JvS022\_03\_L1.sorted.bam.qualimap/qualimapReport.html - results/historical/mapping/GCF\_000283155.1\_CerSimSim1.0\_genomic.Sc9M7eS\_2\_HRSCAF\_41/stats/bams\_sorted/JvS022\_04\_L1.sorted.bam.qualimap/qualimapReport.html - results/historical/mapping/GCF\_000283155.1\_CerSimSim1.0\_genomic.Sc9M7eS\_2\_HRSCAF\_41/stats/bams\_sorted/JvS022\_05\_L1.sorted.bam.qualimap/qualimapReport.html - results/historical/mapping/GCF\_000283155.1\_CerSimSim1.0\_genomic.Sc9M7eS\_2\_HRSCAF\_41/stats/bams\_sorted/JvS022\_06\_L1.sorted.bam.qualimap/qualimapReport.html - results/historical/mapping/GCF\_000283155.1\_CerSimSim1.0\_genomic.Sc9M7eS\_2\_HRSCAF\_41/stats/bams\_sorted/JvS022\_07\_L1.sorted.bam.qualimap/qualimapReport.html - results/historical/mapping/GCF\_000283155.1\_CerSimSim1.0\_genomic.Sc9M7eS\_2\_HRSCAF\_41/stats/bams\_sorted/JvS022\_08\_L1.sorted.bam.qualimap/qualimapReport.html - results/historical/mapping/GCF\_000283155.1\_CerSimSim1.0\_genomic.Sc9M7eS\_2\_HRSCAF\_41/stats/bams\_sorted/JvS022\_09\_L1.sorted.bam.qualimap/qualimapReport.html - results/historical/mapping/GCF\_000283155.1\_CerSimSim1.0\_genomic.Sc9M7eS\_2\_HRSCAF\_41/stats/bams\_sorted/JvS022\_10\_L1.sorted.bam.qualimap/qualimapReport.html - results/historical/mapping/GCF\_000283155.1\_CerSimSim1.0\_genomic.Sc9M7eS\_2\_HRSCAF\_41/stats/bams\_sorted/JvS022\_11\_L1.sorted.bam.qualimap/qualimapReport.html - results/historical/mapping/GCF\_000283155.1\_CerSimSim1.0\_genomic.Sc9M7eS\_2\_HRSCAF\_41/stats/bams\_sorted/JvS022\_12\_L1.sorted.bam.qualimap/qualimapReport.html - results/historical/mapping/GCF\_000283155.1\_CerSimSim1.0\_genomic.Sc9M7eS\_2\_HRSCAF\_41/stats/bams\_sorted/JvS022\_74\_L8.sorted.bam.qualimap/qualimapReport.html - results/historical/mapping/GCF\_000283155.1\_CerSimSim1.0\_genomic.Sc9M7eS\_2\_HRSCAF\_41/stats/bams\_sorted/JvS022\_75\_L8.sorted.bam.qualimap/qualimapReport.html - results/historical/mapping/GCF\_000283155.1\_CerSimSim1.0\_genomic.Sc9M7eS\_2\_HRSCAF\_41/stats/bams\_sorted/JvS022\_76\_L8.sorted.bam.qualimap/qualimapReport.html - results/historical/mapping/GCF\_000283155.1\_CerSimSim1.0\_genomic.Sc9M7eS\_2\_HRSCAF\_41/stats/bams\_sorted/JvS022\_77\_L8.sorted.bam.qualimap/qualimapReport.html - results/historical/mapping/GCF\_000283155.1\_CerSimSim1.0\_genomic.Sc9M7eS\_2\_HRSCAF\_41/stats/bams\_sorted/JvS022\_78\_L8.sorted.bam.qualimap/qualimapReport.html - results/historical/mapping/GCF\_000283155.1\_CerSimSim1.0\_genomic.Sc9M7eS\_2\_HRSCAF\_41/stats/bams\_sorted/JvS022\_79\_L8.sorted.bam.qualimap/qualimapReport.html - results/historical/mapping/GCF\_000283155.1\_CerSimSim1.0\_genomic.Sc9M7eS\_2\_HRSCAF\_41/stats/bams\_sorted/JvS022\_80\_L8.sorted.bam.qualimap/qualimapReport.html - results/historical/mapping/GCF\_000283155.1\_CerSimSim1.0\_genomic.Sc9M7eS\_2\_HRSCAF\_41/stats/bams\_sorted/JvS022\_81\_L8.sorted.bam.qualimap/qualimapReport.html - results/historical/mapping/GCF\_000283155.1\_CerSimSim1.0\_genomic.Sc9M7eS\_2\_HRSCAF\_41/stats/bams\_sorted/JvS022\_82\_L8.sorted.bam.qualimap/qualimapReport.html - results/historical/mapping/GCF\_000283155.1\_CerSimSim1.0\_genomic.Sc9M7eS\_2\_HRSCAF\_41/stats/bams\_sorted/JvS022\_83\_L8.sorted.bam.qualimap/qualimapReport.html - results/historical/mapping/GCF\_000283155.1\_CerSimSim1.0\_genomic.Sc9M7eS\_2\_HRSCAF\_41/stats/bams\_sorted/JvS022\_84\_L8.sorted.bam.qualimap/qualimapReport.html - results/historical/mapping/GCF\_000283155.1\_CerSimSim1.0\_genomic.Sc9M7eS\_2\_HRSCAF\_41/stats/bams\_sorted/JvS022\_85\_L8.sorted.bam.qualimap/qualimapReport.html | |
| Output files | |
| - results/historical/mapping/GCF\_000283155.1\_CerSimSim1.0\_genomic.Sc9M7eS\_2\_HRSCAF\_41/stats/bams\_sorted/multiqc/multiqc\_report.html | |
| Container image | |
| docker://quay.io/biocontainers/multiqc:1.9--pyh9f0ad1d\_0 | |
| Code | |
| ``` 1 2 ``` \| ``` multiqc -f {params.indir} -o {params.outdir} 2> {log} ``` | |

###### Rule sorted\_bam\_stats

×

Rule properties

|  |  |
| --- | --- |
| Jobs | 39 |
| Input files | |
| - results/{dataset}/mapping/GCF\_000283155.1\_CerSimSim1.0\_genomic.Sc9M7eS\_2\_HRSCAF\_41/{sample}\_{index}\_{lane}.sorted.bam - results/{dataset}/mapping/GCF\_000283155.1\_CerSimSim1.0\_genomic.Sc9M7eS\_2\_HRSCAF\_41/{sample}\_{index}\_{lane}.sorted.bam.bai | |
| Output files | |
| - results/{dataset}/mapping/GCF\_000283155.1\_CerSimSim1.0\_genomic.Sc9M7eS\_2\_HRSCAF\_41/stats/bams\_sorted/{sample,[A-Za-z0-9]+}\_{index}\_{lane}.sorted.bam.stats.txt | |
| Container image | |
| docker://biocontainers/samtools:v1.9-4-deb\_cv1 | |
| Code | |
| ``` 1 2 ``` \| ``` samtools flagstat {input.bam} > {output.stats} 2> {log} ``` | |

###### Rule sai2bam

×

Rule properties

|  |  |
| --- | --- |
| Jobs | 36 |
| Input files | |
| - /proj/sllstore2017093/b2016342/b2016342\_nobackup/lts/genome\_erosion\_pipeline/verena\_testing/testdata/reference/GCF\_000283155.1\_CerSimSim1.0\_genomic.Sc9M7eS\_2\_HRSCAF\_41.fasta - /proj/sllstore2017093/b2016342/b2016342\_nobackup/lts/genome\_erosion\_pipeline/verena\_testing/testdata/reference/GCF\_000283155.1\_CerSimSim1.0\_genomic.Sc9M7eS\_2\_HRSCAF\_41.fasta.amb - /proj/sllstore2017093/b2016342/b2016342\_nobackup/lts/genome\_erosion\_pipeline/verena\_testing/testdata/reference/GCF\_000283155.1\_CerSimSim1.0\_genomic.Sc9M7eS\_2\_HRSCAF\_41.fasta.ann - /proj/sllstore2017093/b2016342/b2016342\_nobackup/lts/genome\_erosion\_pipeline/verena\_testing/testdata/reference/GCF\_000283155.1\_CerSimSim1.0\_genomic.Sc9M7eS\_2\_HRSCAF\_41.fasta.bwt - /proj/sllstore2017093/b2016342/b2016342\_nobackup/lts/genome\_erosion\_pipeline/verena\_testing/testdata/reference/GCF\_000283155.1\_CerSimSim1.0\_genomic.Sc9M7eS\_2\_HRSCAF\_41.fasta.pac - /proj/sllstore2017093/b2016342/b2016342\_nobackup/lts/genome\_erosion\_pipeline/verena\_testing/testdata/reference/GCF\_000283155.1\_CerSimSim1.0\_genomic.Sc9M7eS\_2\_HRSCAF\_41.fasta.sa - results/historical/trimming/{sample}\_{index}\_{lane}\_trimmed\_merged.fastq.gz - results/historical/mapping/GCF\_000283155.1\_CerSimSim1.0\_genomic.Sc9M7eS\_2\_HRSCAF\_41/{sample}\_{index}\_{lane}.sai - results/historical/mapping/GCF\_000283155.1\_CerSimSim1.0\_genomic.Sc9M7eS\_2\_HRSCAF\_41/{sample}\_{index}\_{lane}.readgroup.txt | |
| Output files | |
| - results/historical/mapping/GCF\_000283155.1\_CerSimSim1.0\_genomic.Sc9M7eS\_2\_HRSCAF\_41/{sample,[A-Za-z0-9]+}\_{index}\_{lane}.sorted.bam | |
| Container image | |
| docker://nbisweden/generode-bwa:latest | |
| Code | |
| ``` 1 2 ``` \| ``` bwa samse -r $(cat {input.rg}) {input.ref} {input.sai} {input.fastq_hist} \| samtools sort -@ {threads} - > {output.bam} 2> {log} ``` | |

###### Rule map\_historical

×

Rule properties

|  |  |
| --- | --- |
| Jobs | 36 |
| Input files | |
| - /proj/sllstore2017093/b2016342/b2016342\_nobackup/lts/genome\_erosion\_pipeline/verena\_testing/testdata/reference/GCF\_000283155.1\_CerSimSim1.0\_genomic.Sc9M7eS\_2\_HRSCAF\_41.fasta - /proj/sllstore2017093/b2016342/b2016342\_nobackup/lts/genome\_erosion\_pipeline/verena\_testing/testdata/reference/GCF\_000283155.1\_CerSimSim1.0\_genomic.Sc9M7eS\_2\_HRSCAF\_41.fasta.amb - /proj/sllstore2017093/b2016342/b2016342\_nobackup/lts/genome\_erosion\_pipeline/verena\_testing/testdata/reference/GCF\_000283155.1\_CerSimSim1.0\_genomic.Sc9M7eS\_2\_HRSCAF\_41.fasta.ann - /proj/sllstore2017093/b2016342/b2016342\_nobackup/lts/genome\_erosion\_pipeline/verena\_testing/testdata/reference/GCF\_000283155.1\_CerSimSim1.0\_genomic.Sc9M7eS\_2\_HRSCAF\_41.fasta.bwt - /proj/sllstore2017093/b2016342/b2016342\_nobackup/lts/genome\_erosion\_pipeline/verena\_testing/testdata/reference/GCF\_000283155.1\_CerSimSim1.0\_genomic.Sc9M7eS\_2\_HRSCAF\_41.fasta.pac - /proj/sllstore2017093/b2016342/b2016342\_nobackup/lts/genome\_erosion\_pipeline/verena\_testing/testdata/reference/GCF\_000283155.1\_CerSimSim1.0\_genomic.Sc9M7eS\_2\_HRSCAF\_41.fasta.sa - results/historical/trimming/{sample}\_{index}\_{lane}\_trimmed\_merged.fastq.gz | |
| Output files | |
| - results/historical/mapping/GCF\_000283155.1\_CerSimSim1.0\_genomic.Sc9M7eS\_2\_HRSCAF\_41/{sample,[A-Za-z0-9]+}\_{index}\_{lane}.sai | |
| Container image | |
| docker://biocontainers/bwa:v0.7.17-3-deb\_cv1 | |
| Code | |
| ``` 1 2 ``` \| ``` bwa aln -l 16500 -n 0.01 -o 2 -t {threads} {input.ref} {input.fastq_hist} > {output.sai} 2> {log} ``` | |

###### Rule readgroup\_ID\_historical

×

Rule properties

|  |  |
| --- | --- |
| Jobs | 36 |
| Input files | |
| - results/historical/mapping/GCF\_000283155.1\_CerSimSim1.0\_genomic.Sc9M7eS\_2\_HRSCAF\_41/{sample}\_{index}\_{lane}.sai | |
| Output files | |
| - results/historical/mapping/GCF\_000283155.1\_CerSimSim1.0\_genomic.Sc9M7eS\_2\_HRSCAF\_41/{sample,[A-Za-z0-9]+}\_{index}\_{lane}.readgroup.txt | |

###### Rule index\_sorted\_bams

×

Rule properties

|  |  |
| --- | --- |
| Jobs | 39 |
| Input files | |
| - results/{dataset}/mapping/GCF\_000283155.1\_CerSimSim1.0\_genomic.Sc9M7eS\_2\_HRSCAF\_41/{sample}\_{index}\_{lane}.sorted.bam | |
| Output files | |
| - results/{dataset}/mapping/GCF\_000283155.1\_CerSimSim1.0\_genomic.Sc9M7eS\_2\_HRSCAF\_41/{sample,[A-Za-z0-9]+}\_{index}\_{lane}.sorted.bam.bai | |
| Container image | |
| docker://biocontainers/samtools:v1.9-4-deb\_cv1 | |
| Code | |
| ``` 1 2 ``` \| ``` samtools index {input.bam} {output.index} 2> {log} ``` | |

###### Rule sorted\_bam\_qualimap

×

Rule properties

|  |  |
| --- | --- |
| Jobs | 39 |
| Input files | |
| - results/{dataset}/mapping/GCF\_000283155.1\_CerSimSim1.0\_genomic.Sc9M7eS\_2\_HRSCAF\_41/{sample}\_{index}\_{lane}.sorted.bam - results/{dataset}/mapping/GCF\_000283155.1\_CerSimSim1.0\_genomic.Sc9M7eS\_2\_HRSCAF\_41/{sample}\_{index}\_{lane}.sorted.bam.bai | |
| Output files | |
| - results/{dataset}/mapping/GCF\_000283155.1\_CerSimSim1.0\_genomic.Sc9M7eS\_2\_HRSCAF\_41/stats/bams\_sorted/{sample,[A-Za-z0-9]+}\_{index}\_{lane}.sorted.bam.qualimap/qualimapReport.html - results/{dataset}/mapping/GCF\_000283155.1\_CerSimSim1.0\_genomic.Sc9M7eS\_2\_HRSCAF\_41/stats/bams\_sorted/{sample,[A-Za-z0-9]+}\_{index}\_{lane}.sorted.bam.qualimap/genome\_results.txt - results/{dataset}/mapping/GCF\_000283155.1\_CerSimSim1.0\_genomic.Sc9M7eS\_2\_HRSCAF\_41/stats/bams\_sorted/{sample,[A-Za-z0-9]+}\_{index}\_{lane}.sorted.bam.qualimap | |
| Container image | |
| docker://quay.io/biocontainers/qualimap:2.2.2d--1 | |
| Code | |
| ``` 1 2 3 4 ``` \| ``` mem=$(((6 * {threads}) - 2)) unset DISPLAY qualimap bamqc -bam {input.bam} --java-mem-size=${{mem}}G -nt {threads} -outdir {params.outdir} -outformat html 2> {log} ``` | |

###### Rule modern\_raw\_bam\_multiqc

×

Rule properties

|  |  |
| --- | --- |
| Jobs | 1 |
| Input files | |
| - results/modern/mapping/GCF\_000283155.1\_CerSimSim1.0\_genomic.Sc9M7eS\_2\_HRSCAF\_41/stats/bams\_sorted/JvS033\_01\_L7.sorted.bam.stats.txt - results/modern/mapping/GCF\_000283155.1\_CerSimSim1.0\_genomic.Sc9M7eS\_2\_HRSCAF\_41/stats/bams\_sorted/JvS034\_02\_L8.sorted.bam.stats.txt - results/modern/mapping/GCF\_000283155.1\_CerSimSim1.0\_genomic.Sc9M7eS\_2\_HRSCAF\_41/stats/bams\_sorted/JvS035\_14\_L7.sorted.bam.stats.txt - results/modern/mapping/GCF\_000283155.1\_CerSimSim1.0\_genomic.Sc9M7eS\_2\_HRSCAF\_41/stats/bams\_sorted/JvS033\_01\_L7.sorted.bam.qualimap/qualimapReport.html - results/modern/mapping/GCF\_000283155.1\_CerSimSim1.0\_genomic.Sc9M7eS\_2\_HRSCAF\_41/stats/bams\_sorted/JvS034\_02\_L8.sorted.bam.qualimap/qualimapReport.html - results/modern/mapping/GCF\_000283155.1\_CerSimSim1.0\_genomic.Sc9M7eS\_2\_HRSCAF\_41/stats/bams\_sorted/JvS035\_14\_L7.sorted.bam.qualimap/qualimapReport.html | |
| Output files | |
| - results/modern/mapping/GCF\_000283155.1\_CerSimSim1.0\_genomic.Sc9M7eS\_2\_HRSCAF\_41/stats/bams\_sorted/multiqc/multiqc\_report.html | |
| Container image | |
| docker://quay.io/biocontainers/multiqc:1.9--pyh9f0ad1d\_0 | |
| Code | |
| ``` 1 2 ``` \| ``` multiqc -f {params.indir} -o {params.outdir} 2> {log} ``` | |

###### Rule map\_modern

×

Rule properties

|  |  |
| --- | --- |
| Jobs | 3 |
| Input files | |
| - /proj/sllstore2017093/b2016342/b2016342\_nobackup/lts/genome\_erosion\_pipeline/verena\_testing/testdata/reference/GCF\_000283155.1\_CerSimSim1.0\_genomic.Sc9M7eS\_2\_HRSCAF\_41.fasta - /proj/sllstore2017093/b2016342/b2016342\_nobackup/lts/genome\_erosion\_pipeline/verena\_testing/testdata/reference/GCF\_000283155.1\_CerSimSim1.0\_genomic.Sc9M7eS\_2\_HRSCAF\_41.fasta.amb - /proj/sllstore2017093/b2016342/b2016342\_nobackup/lts/genome\_erosion\_pipeline/verena\_testing/testdata/reference/GCF\_000283155.1\_CerSimSim1.0\_genomic.Sc9M7eS\_2\_HRSCAF\_41.fasta.ann - /proj/sllstore2017093/b2016342/b2016342\_nobackup/lts/genome\_erosion\_pipeline/verena\_testing/testdata/reference/GCF\_000283155.1\_CerSimSim1.0\_genomic.Sc9M7eS\_2\_HRSCAF\_41.fasta.bwt - /proj/sllstore2017093/b2016342/b2016342\_nobackup/lts/genome\_erosion\_pipeline/verena\_testing/testdata/reference/GCF\_000283155.1\_CerSimSim1.0\_genomic.Sc9M7eS\_2\_HRSCAF\_41.fasta.pac - /proj/sllstore2017093/b2016342/b2016342\_nobackup/lts/genome\_erosion\_pipeline/verena\_testing/testdata/reference/GCF\_000283155.1\_CerSimSim1.0\_genomic.Sc9M7eS\_2\_HRSCAF\_41.fasta.sa - results/modern/trimming/{sample}\_{index}\_{lane}\_R1\_trimmed.fastq.gz - results/modern/trimming/{sample}\_{index}\_{lane}\_R2\_trimmed.fastq.gz - results/modern/mapping/GCF\_000283155.1\_CerSimSim1.0\_genomic.Sc9M7eS\_2\_HRSCAF\_41/{sample}\_{index}\_{lane}.readgroup.txt | |
| Output files | |
| - results/modern/mapping/GCF\_000283155.1\_CerSimSim1.0\_genomic.Sc9M7eS\_2\_HRSCAF\_41/{sample,[A-Za-z0-9]+}\_{index}\_{lane}.sorted.bam | |
| Container image | |
| docker://nbisweden/generode-bwa:latest | |
| Code | |
| ``` 1 2 ``` \| ``` bwa mem -M -t {threads} -R $(cat {input.rg}) {input.ref} {input.fastq_mod_R1} {input.fastq_mod_R2} \| samtools sort -@ {threads} - > {output.bam} 2> {log} ``` | |

###### Rule readgroup\_ID\_modern

×

Rule properties

|  |  |
| --- | --- |
| Jobs | 3 |
| Input files | |
| - /proj/sllstore2017093/b2016342/b2016342\_nobackup/lts/genome\_erosion\_pipeline/verena\_testing/testdata/reference/GCF\_000283155.1\_CerSimSim1.0\_genomic.Sc9M7eS\_2\_HRSCAF\_41.fasta - /proj/sllstore2017093/b2016342/b2016342\_nobackup/lts/genome\_erosion\_pipeline/verena\_testing/testdata/reference/GCF\_000283155.1\_CerSimSim1.0\_genomic.Sc9M7eS\_2\_HRSCAF\_41.fasta.amb - /proj/sllstore2017093/b2016342/b2016342\_nobackup/lts/genome\_erosion\_pipeline/verena\_testing/testdata/reference/GCF\_000283155.1\_CerSimSim1.0\_genomic.Sc9M7eS\_2\_HRSCAF\_41.fasta.ann - /proj/sllstore2017093/b2016342/b2016342\_nobackup/lts/genome\_erosion\_pipeline/verena\_testing/testdata/reference/GCF\_000283155.1\_CerSimSim1.0\_genomic.Sc9M7eS\_2\_HRSCAF\_41.fasta.bwt - /proj/sllstore2017093/b2016342/b2016342\_nobackup/lts/genome\_erosion\_pipeline/verena\_testing/testdata/reference/GCF\_000283155.1\_CerSimSim1.0\_genomic.Sc9M7eS\_2\_HRSCAF\_41.fasta.pac - /proj/sllstore2017093/b2016342/b2016342\_nobackup/lts/genome\_erosion\_pipeline/verena\_testing/testdata/reference/GCF\_000283155.1\_CerSimSim1.0\_genomic.Sc9M7eS\_2\_HRSCAF\_41.fasta.sa - results/modern/trimming/{sample}\_{index}\_{lane}\_R1\_trimmed.fastq.gz - results/modern/trimming/{sample}\_{index}\_{lane}\_R2\_trimmed.fastq.gz | |
| Output files | |
| - results/modern/mapping/GCF\_000283155.1\_CerSimSim1.0\_genomic.Sc9M7eS\_2\_HRSCAF\_41/{sample,[A-Za-z0-9]+}\_{index}\_{lane}.readgroup.txt | |

###### Rule historical\_merged\_index\_bam\_multiqc

×

Rule properties

|  |  |
| --- | --- |
| Jobs | 1 |
| Input files | |
| - results/historical/mapping/GCF\_000283155.1\_CerSimSim1.0\_genomic.Sc9M7eS\_2\_HRSCAF\_41/stats/bams\_merged\_index/JvS008\_08.merged.bam.stats.txt - results/historical/mapping/GCF\_000283155.1\_CerSimSim1.0\_genomic.Sc9M7eS\_2\_HRSCAF\_41/stats/bams\_merged\_index/JvS008\_10.merged.bam.stats.txt - results/historical/mapping/GCF\_000283155.1\_CerSimSim1.0\_genomic.Sc9M7eS\_2\_HRSCAF\_41/stats/bams\_merged\_index/JvS008\_11.merged.bam.stats.txt - results/historical/mapping/GCF\_000283155.1\_CerSimSim1.0\_genomic.Sc9M7eS\_2\_HRSCAF\_41/stats/bams\_merged\_index/JvS009\_09.merged.bam.stats.txt - results/historical/mapping/GCF\_000283155.1\_CerSimSim1.0\_genomic.Sc9M7eS\_2\_HRSCAF\_41/stats/bams\_merged\_index/JvS009\_15.merged.bam.stats.txt - results/historical/mapping/GCF\_000283155.1\_CerSimSim1.0\_genomic.Sc9M7eS\_2\_HRSCAF\_41/stats/bams\_merged\_index/JvS009\_19.merged.bam.stats.txt - results/historical/mapping/GCF\_000283155.1\_CerSimSim1.0\_genomic.Sc9M7eS\_2\_HRSCAF\_41/stats/bams\_merged\_index/JvS022\_01.merged.bam.stats.txt - results/historical/mapping/GCF\_000283155.1\_CerSimSim1.0\_genomic.Sc9M7eS\_2\_HRSCAF\_41/stats/bams\_merged\_index/JvS022\_02.merged.bam.stats.txt - results/historical/mapping/GCF\_000283155.1\_CerSimSim1.0\_genomic.Sc9M7eS\_2\_HRSCAF\_41/stats/bams\_merged\_index/JvS022\_03.merged.bam.stats.txt - results/historical/mapping/GCF\_000283155.1\_CerSimSim1.0\_genomic.Sc9M7eS\_2\_HRSCAF\_41/stats/bams\_merged\_index/JvS022\_04.merged.bam.stats.txt - results/historical/mapping/GCF\_000283155.1\_CerSimSim1.0\_genomic.Sc9M7eS\_2\_HRSCAF\_41/stats/bams\_merged\_index/JvS022\_05.merged.bam.stats.txt - results/historical/mapping/GCF\_000283155.1\_CerSimSim1.0\_genomic.Sc9M7eS\_2\_HRSCAF\_41/stats/bams\_merged\_index/JvS022\_06.merged.bam.stats.txt - results/historical/mapping/GCF\_000283155.1\_CerSimSim1.0\_genomic.Sc9M7eS\_2\_HRSCAF\_41/stats/bams\_merged\_index/JvS022\_07.merged.bam.stats.txt - results/historical/mapping/GCF\_000283155.1\_CerSimSim1.0\_genomic.Sc9M7eS\_2\_HRSCAF\_41/stats/bams\_merged\_index/JvS022\_08.merged.bam.stats.txt - results/historical/mapping/GCF\_000283155.1\_CerSimSim1.0\_genomic.Sc9M7eS\_2\_HRSCAF\_41/stats/bams\_merged\_index/JvS022\_09.merged.bam.stats.txt - results/historical/mapping/GCF\_000283155.1\_CerSimSim1.0\_genomic.Sc9M7eS\_2\_HRSCAF\_41/stats/bams\_merged\_index/JvS022\_10.merged.bam.stats.txt - results/historical/mapping/GCF\_000283155.1\_CerSimSim1.0\_genomic.Sc9M7eS\_2\_HRSCAF\_41/stats/bams\_merged\_index/JvS022\_11.merged.bam.stats.txt - results/historical/mapping/GCF\_000283155.1\_CerSimSim1.0\_genomic.Sc9M7eS\_2\_HRSCAF\_41/stats/bams\_merged\_index/JvS022\_12.merged.bam.stats.txt - results/historical/mapping/GCF\_000283155.1\_CerSimSim1.0\_genomic.Sc9M7eS\_2\_HRSCAF\_41/stats/bams\_merged\_index/JvS022\_74.merged.bam.stats.txt - results/historical/mapping/GCF\_000283155.1\_CerSimSim1.0\_genomic.Sc9M7eS\_2\_HRSCAF\_41/stats/bams\_merged\_index/JvS022\_75.merged.bam.stats.txt - results/historical/mapping/GCF\_000283155.1\_CerSimSim1.0\_genomic.Sc9M7eS\_2\_HRSCAF\_41/stats/bams\_merged\_index/JvS022\_76.merged.bam.stats.txt - results/historical/mapping/GCF\_000283155.1\_CerSimSim1.0\_genomic.Sc9M7eS\_2\_HRSCAF\_41/stats/bams\_merged\_index/JvS022\_77.merged.bam.stats.txt - results/historical/mapping/GCF\_000283155.1\_CerSimSim1.0\_genomic.Sc9M7eS\_2\_HRSCAF\_41/stats/bams\_merged\_index/JvS022\_78.merged.bam.stats.txt - results/historical/mapping/GCF\_000283155.1\_CerSimSim1.0\_genomic.Sc9M7eS\_2\_HRSCAF\_41/stats/bams\_merged\_index/JvS022\_79.merged.bam.stats.txt - results/historical/mapping/GCF\_000283155.1\_CerSimSim1.0\_genomic.Sc9M7eS\_2\_HRSCAF\_41/stats/bams\_merged\_index/JvS022\_80.merged.bam.stats.txt - results/historical/mapping/GCF\_000283155.1\_CerSimSim1.0\_genomic.Sc9M7eS\_2\_HRSCAF\_41/stats/bams\_merged\_index/JvS022\_81.merged.bam.stats.txt - results/historical/mapping/GCF\_000283155.1\_CerSimSim1.0\_genomic.Sc9M7eS\_2\_HRSCAF\_41/stats/bams\_merged\_index/JvS022\_82.merged.bam.stats.txt - results/historical/mapping/GCF\_000283155.1\_CerSimSim1.0\_genomic.Sc9M7eS\_2\_HRSCAF\_41/stats/bams\_merged\_index/JvS022\_83.merged.bam.stats.txt - results/historical/mapping/GCF\_000283155.1\_CerSimSim1.0\_genomic.Sc9M7eS\_2\_HRSCAF\_41/stats/bams\_merged\_index/JvS022\_84.merged.bam.stats.txt - results/historical/mapping/GCF\_000283155.1\_CerSimSim1.0\_genomic.Sc9M7eS\_2\_HRSCAF\_41/stats/bams\_merged\_index/JvS022\_85.merged.bam.stats.txt - results/historical/mapping/GCF\_000283155.1\_CerSimSim1.0\_genomic.Sc9M7eS\_2\_HRSCAF\_41/stats/bams\_merged\_index/JvS008\_08.merged.bam.qualimap/qualimapReport.html - results/historical/mapping/GCF\_000283155.1\_CerSimSim1.0\_genomic.Sc9M7eS\_2\_HRSCAF\_41/stats/bams\_merged\_index/JvS008\_10.merged.bam.qualimap/qualimapReport.html - results/historical/mapping/GCF\_000283155.1\_CerSimSim1.0\_genomic.Sc9M7eS\_2\_HRSCAF\_41/stats/bams\_merged\_index/JvS008\_11.merged.bam.qualimap/qualimapReport.html - results/historical/mapping/GCF\_000283155.1\_CerSimSim1.0\_genomic.Sc9M7eS\_2\_HRSCAF\_41/stats/bams\_merged\_index/JvS009\_09.merged.bam.qualimap/qualimapReport.html - results/historical/mapping/GCF\_000283155.1\_CerSimSim1.0\_genomic.Sc9M7eS\_2\_HRSCAF\_41/stats/bams\_merged\_index/JvS009\_15.merged.bam.qualimap/qualimapReport.html - results/historical/mapping/GCF\_000283155.1\_CerSimSim1.0\_genomic.Sc9M7eS\_2\_HRSCAF\_41/stats/bams\_merged\_index/JvS009\_19.merged.bam.qualimap/qualimapReport.html - results/historical/mapping/GCF\_000283155.1\_CerSimSim1.0\_genomic.Sc9M7eS\_2\_HRSCAF\_41/stats/bams\_merged\_index/JvS022\_01.merged.bam.qualimap/qualimapReport.html - results/historical/mapping/GCF\_000283155.1\_CerSimSim1.0\_genomic.Sc9M7eS\_2\_HRSCAF\_41/stats/bams\_merged\_index/JvS022\_02.merged.bam.qualimap/qualimapReport.html - results/historical/mapping/GCF\_000283155.1\_CerSimSim1.0\_genomic.Sc9M7eS\_2\_HRSCAF\_41/stats/bams\_merged\_index/JvS022\_03.merged.bam.qualimap/qualimapReport.html - results/historical/mapping/GCF\_000283155.1\_CerSimSim1.0\_genomic.Sc9M7eS\_2\_HRSCAF\_41/stats/bams\_merged\_index/JvS022\_04.merged.bam.qualimap/qualimapReport.html - results/historical/mapping/GCF\_000283155.1\_CerSimSim1.0\_genomic.Sc9M7eS\_2\_HRSCAF\_41/stats/bams\_merged\_index/JvS022\_05.merged.bam.qualimap/qualimapReport.html - results/historical/mapping/GCF\_000283155.1\_CerSimSim1.0\_genomic.Sc9M7eS\_2\_HRSCAF\_41/stats/bams\_merged\_index/JvS022\_06.merged.bam.qualimap/qualimapReport.html - results/historical/mapping/GCF\_000283155.1\_CerSimSim1.0\_genomic.Sc9M7eS\_2\_HRSCAF\_41/stats/bams\_merged\_index/JvS022\_07.merged.bam.qualimap/qualimapReport.html - results/historical/mapping/GCF\_000283155.1\_CerSimSim1.0\_genomic.Sc9M7eS\_2\_HRSCAF\_41/stats/bams\_merged\_index/JvS022\_08.merged.bam.qualimap/qualimapReport.html - results/historical/mapping/GCF\_000283155.1\_CerSimSim1.0\_genomic.Sc9M7eS\_2\_HRSCAF\_41/stats/bams\_merged\_index/JvS022\_09.merged.bam.qualimap/qualimapReport.html - results/historical/mapping/GCF\_000283155.1\_CerSimSim1.0\_genomic.Sc9M7eS\_2\_HRSCAF\_41/stats/bams\_merged\_index/JvS022\_10.merged.bam.qualimap/qualimapReport.html - results/historical/mapping/GCF\_000283155.1\_CerSimSim1.0\_genomic.Sc9M7eS\_2\_HRSCAF\_41/stats/bams\_merged\_index/JvS022\_11.merged.bam.qualimap/qualimapReport.html - results/historical/mapping/GCF\_000283155.1\_CerSimSim1.0\_genomic.Sc9M7eS\_2\_HRSCAF\_41/stats/bams\_merged\_index/JvS022\_12.merged.bam.qualimap/qualimapReport.html - results/historical/mapping/GCF\_000283155.1\_CerSimSim1.0\_genomic.Sc9M7eS\_2\_HRSCAF\_41/stats/bams\_merged\_index/JvS022\_74.merged.bam.qualimap/qualimapReport.html - results/historical/mapping/GCF\_000283155.1\_CerSimSim1.0\_genomic.Sc9M7eS\_2\_HRSCAF\_41/stats/bams\_merged\_index/JvS022\_75.merged.bam.qualimap/qualimapReport.html - results/historical/mapping/GCF\_000283155.1\_CerSimSim1.0\_genomic.Sc9M7eS\_2\_HRSCAF\_41/stats/bams\_merged\_index/JvS022\_76.merged.bam.qualimap/qualimapReport.html - results/historical/mapping/GCF\_000283155.1\_CerSimSim1.0\_genomic.Sc9M7eS\_2\_HRSCAF\_41/stats/bams\_merged\_index/JvS022\_77.merged.bam.qualimap/qualimapReport.html - results/historical/mapping/GCF\_000283155.1\_CerSimSim1.0\_genomic.Sc9M7eS\_2\_HRSCAF\_41/stats/bams\_merged\_index/JvS022\_78.merged.bam.qualimap/qualimapReport.html - results/historical/mapping/GCF\_000283155.1\_CerSimSim1.0\_genomic.Sc9M7eS\_2\_HRSCAF\_41/stats/bams\_merged\_index/JvS022\_79.merged.bam.qualimap/qualimapReport.html - results/historical/mapping/GCF\_000283155.1\_CerSimSim1.0\_genomic.Sc9M7eS\_2\_HRSCAF\_41/stats/bams\_merged\_index/JvS022\_80.merged.bam.qualimap/qualimapReport.html - results/historical/mapping/GCF\_000283155.1\_CerSimSim1.0\_genomic.Sc9M7eS\_2\_HRSCAF\_41/stats/bams\_merged\_index/JvS022\_81.merged.bam.qualimap/qualimapReport.html - results/historical/mapping/GCF\_000283155.1\_CerSimSim1.0\_genomic.Sc9M7eS\_2\_HRSCAF\_41/stats/bams\_merged\_index/JvS022\_82.merged.bam.qualimap/qualimapReport.html - results/historical/mapping/GCF\_000283155.1\_CerSimSim1.0\_genomic.Sc9M7eS\_2\_HRSCAF\_41/stats/bams\_merged\_index/JvS022\_83.merged.bam.qualimap/qualimapReport.html - results/historical/mapping/GCF\_000283155.1\_CerSimSim1.0\_genomic.Sc9M7eS\_2\_HRSCAF\_41/stats/bams\_merged\_index/JvS022\_84.merged.bam.qualimap/qualimapReport.html - results/historical/mapping/GCF\_000283155.1\_CerSimSim1.0\_genomic.Sc9M7eS\_2\_HRSCAF\_41/stats/bams\_merged\_index/JvS022\_85.merged.bam.qualimap/qualimapReport.html | |
| Output files | |
| - results/historical/mapping/GCF\_000283155.1\_CerSimSim1.0\_genomic.Sc9M7eS\_2\_HRSCAF\_41/stats/bams\_merged\_index/multiqc/multiqc\_report.html | |
| Container image | |
| docker://quay.io/biocontainers/multiqc:1.9--pyh9f0ad1d\_0 | |
| Code | |
| ``` 1 2 ``` \| ``` multiqc -f {params.indir} -o {params.outdir} 2> {log} ``` | |

###### Rule merged\_index\_bam\_stats

×

Rule properties

|  |  |
| --- | --- |
| Jobs | 33 |
| Input files | |
| - results/{dataset}/mapping/GCF\_000283155.1\_CerSimSim1.0\_genomic.Sc9M7eS\_2\_HRSCAF\_41/{sample}\_{index}.merged.bam - results/{dataset}/mapping/GCF\_000283155.1\_CerSimSim1.0\_genomic.Sc9M7eS\_2\_HRSCAF\_41/{sample}\_{index}.merged.bam.bai | |
| Output files | |
| - results/{dataset}/mapping/GCF\_000283155.1\_CerSimSim1.0\_genomic.Sc9M7eS\_2\_HRSCAF\_41/stats/bams\_merged\_index/{sample,[A-Za-z0-9]+}\_{index}.merged.bam.stats.txt | |
| Container image | |
| docker://biocontainers/samtools:v1.9-4-deb\_cv1 | |
| Code | |
| ``` 1 2 ``` \| ``` samtools flagstat {input.bam} > {output.stats} 2> {log} ``` | |

###### Rule merge\_historical\_bams\_per\_index

×

Rule properties

|  |  |
| --- | --- |
| Jobs | 30 |
| Input files | |
| Output files | |
| - results/historical/mapping/GCF\_000283155.1\_CerSimSim1.0\_genomic.Sc9M7eS\_2\_HRSCAF\_41/{sample,[A-Za-z0-9]+}\_{index}.merged.bam | |
| Container image | |
| docker://biocontainers/samtools:v1.9-4-deb\_cv1 | |
| Code | |
| ``` 1 2 3 4 5 6 7 8 9 ``` \| ``` files=`echo {input} \| awk '{{print NF}}'` if [ $files -gt 1 ] # check if there are at least 2 files for merging. If there is only one file, copy the sorted bam file. then samtools merge {output.merged} {input} 2> {log} else cp {input} {output.merged} && touch {output.merged} 2> {log} echo "Only one file present for merging. Copying the input bam file." >> {log} fi ``` | |

###### Rule index\_merged\_index\_bams

×

Rule properties

|  |  |
| --- | --- |
| Jobs | 33 |
| Input files | |
| - results/{dataset}/mapping/GCF\_000283155.1\_CerSimSim1.0\_genomic.Sc9M7eS\_2\_HRSCAF\_41/{sample}\_{index}.merged.bam | |
| Output files | |
| - results/{dataset}/mapping/GCF\_000283155.1\_CerSimSim1.0\_genomic.Sc9M7eS\_2\_HRSCAF\_41/{sample,[A-Za-z0-9]+}\_{index}.merged.bam.bai | |
| Container image | |
| docker://biocontainers/samtools:v1.9-4-deb\_cv1 | |
| Code | |
| ``` 1 2 ``` \| ``` samtools index {input.bam} {output.index} 2> {log} ``` | |

###### Rule merged\_index\_bam\_qualimap

×

Rule properties

|  |  |
| --- | --- |
| Jobs | 33 |
| Input files | |
| - results/{dataset}/mapping/GCF\_000283155.1\_CerSimSim1.0\_genomic.Sc9M7eS\_2\_HRSCAF\_41/{sample}\_{index}.merged.bam - results/{dataset}/mapping/GCF\_000283155.1\_CerSimSim1.0\_genomic.Sc9M7eS\_2\_HRSCAF\_41/{sample}\_{index}.merged.bam.bai | |
| Output files | |
| - results/{dataset}/mapping/GCF\_000283155.1\_CerSimSim1.0\_genomic.Sc9M7eS\_2\_HRSCAF\_41/stats/bams\_merged\_index/{sample,[A-Za-z0-9]+}\_{index}.merged.bam.qualimap/qualimapReport.html - results/{dataset}/mapping/GCF\_000283155.1\_CerSimSim1.0\_genomic.Sc9M7eS\_2\_HRSCAF\_41/stats/bams\_merged\_index/{sample,[A-Za-z0-9]+}\_{index}.merged.bam.qualimap/genome\_results.txt - results/{dataset}/mapping/GCF\_000283155.1\_CerSimSim1.0\_genomic.Sc9M7eS\_2\_HRSCAF\_41/stats/bams\_merged\_index/{sample,[A-Za-z0-9]+}\_{index}.merged.bam.qualimap | |
| Container image | |
| docker://quay.io/biocontainers/qualimap:2.2.2d--1 | |
| Code | |
| ``` 1 2 3 4 ``` \| ``` mem=$(((6 * {threads}) - 2)) unset DISPLAY qualimap bamqc -bam {input.bam} --java-mem-size=${{mem}}G -nt {threads} -outdir {params.outdir} -outformat html 2> {log} ``` | |

###### Rule historical\_rmdup\_bam\_multiqc

×

Rule properties

|  |  |
| --- | --- |
| Jobs | 1 |
| Input files | |
| - results/historical/mapping/GCF\_000283155.1\_CerSimSim1.0\_genomic.Sc9M7eS\_2\_HRSCAF\_41/stats/bams\_rmdup/JvS008\_08.merged.rmdup.bam.stats.txt - results/historical/mapping/GCF\_000283155.1\_CerSimSim1.0\_genomic.Sc9M7eS\_2\_HRSCAF\_41/stats/bams\_rmdup/JvS008\_10.merged.rmdup.bam.stats.txt - results/historical/mapping/GCF\_000283155.1\_CerSimSim1.0\_genomic.Sc9M7eS\_2\_HRSCAF\_41/stats/bams\_rmdup/JvS008\_11.merged.rmdup.bam.stats.txt - results/historical/mapping/GCF\_000283155.1\_CerSimSim1.0\_genomic.Sc9M7eS\_2\_HRSCAF\_41/stats/bams\_rmdup/JvS009\_09.merged.rmdup.bam.stats.txt - results/historical/mapping/GCF\_000283155.1\_CerSimSim1.0\_genomic.Sc9M7eS\_2\_HRSCAF\_41/stats/bams\_rmdup/JvS009\_15.merged.rmdup.bam.stats.txt - results/historical/mapping/GCF\_000283155.1\_CerSimSim1.0\_genomic.Sc9M7eS\_2\_HRSCAF\_41/stats/bams\_rmdup/JvS009\_19.merged.rmdup.bam.stats.txt - results/historical/mapping/GCF\_000283155.1\_CerSimSim1.0\_genomic.Sc9M7eS\_2\_HRSCAF\_41/stats/bams\_rmdup/JvS022\_01.merged.rmdup.bam.stats.txt - results/historical/mapping/GCF\_000283155.1\_CerSimSim1.0\_genomic.Sc9M7eS\_2\_HRSCAF\_41/stats/bams\_rmdup/JvS022\_02.merged.rmdup.bam.stats.txt - results/historical/mapping/GCF\_000283155.1\_CerSimSim1.0\_genomic.Sc9M7eS\_2\_HRSCAF\_41/stats/bams\_rmdup/JvS022\_03.merged.rmdup.bam.stats.txt - results/historical/mapping/GCF\_000283155.1\_CerSimSim1.0\_genomic.Sc9M7eS\_2\_HRSCAF\_41/stats/bams\_rmdup/JvS022\_04.merged.rmdup.bam.stats.txt - results/historical/mapping/GCF\_000283155.1\_CerSimSim1.0\_genomic.Sc9M7eS\_2\_HRSCAF\_41/stats/bams\_rmdup/JvS022\_05.merged.rmdup.bam.stats.txt - results/historical/mapping/GCF\_000283155.1\_CerSimSim1.0\_genomic.Sc9M7eS\_2\_HRSCAF\_41/stats/bams\_rmdup/JvS022\_06.merged.rmdup.bam.stats.txt - results/historical/mapping/GCF\_000283155.1\_CerSimSim1.0\_genomic.Sc9M7eS\_2\_HRSCAF\_41/stats/bams\_rmdup/JvS022\_07.merged.rmdup.bam.stats.txt - results/historical/mapping/GCF\_000283155.1\_CerSimSim1.0\_genomic.Sc9M7eS\_2\_HRSCAF\_41/stats/bams\_rmdup/JvS022\_08.merged.rmdup.bam.stats.txt - results/historical/mapping/GCF\_000283155.1\_CerSimSim1.0\_genomic.Sc9M7eS\_2\_HRSCAF\_41/stats/bams\_rmdup/JvS022\_09.merged.rmdup.bam.stats.txt - results/historical/mapping/GCF\_000283155.1\_CerSimSim1.0\_genomic.Sc9M7eS\_2\_HRSCAF\_41/stats/bams\_rmdup/JvS022\_10.merged.rmdup.bam.stats.txt - results/historical/mapping/GCF\_000283155.1\_CerSimSim1.0\_genomic.Sc9M7eS\_2\_HRSCAF\_41/stats/bams\_rmdup/JvS022\_11.merged.rmdup.bam.stats.txt - results/historical/mapping/GCF\_000283155.1\_CerSimSim1.0\_genomic.Sc9M7eS\_2\_HRSCAF\_41/stats/bams\_rmdup/JvS022\_12.merged.rmdup.bam.stats.txt - results/historical/mapping/GCF\_000283155.1\_CerSimSim1.0\_genomic.Sc9M7eS\_2\_HRSCAF\_41/stats/bams\_rmdup/JvS022\_74.merged.rmdup.bam.stats.txt - results/historical/mapping/GCF\_000283155.1\_CerSimSim1.0\_genomic.Sc9M7eS\_2\_HRSCAF\_41/stats/bams\_rmdup/JvS022\_75.merged.rmdup.bam.stats.txt - results/historical/mapping/GCF\_000283155.1\_CerSimSim1.0\_genomic.Sc9M7eS\_2\_HRSCAF\_41/stats/bams\_rmdup/JvS022\_76.merged.rmdup.bam.stats.txt - results/historical/mapping/GCF\_000283155.1\_CerSimSim1.0\_genomic.Sc9M7eS\_2\_HRSCAF\_41/stats/bams\_rmdup/JvS022\_77.merged.rmdup.bam.stats.txt - results/historical/mapping/GCF\_000283155.1\_CerSimSim1.0\_genomic.Sc9M7eS\_2\_HRSCAF\_41/stats/bams\_rmdup/JvS022\_78.merged.rmdup.bam.stats.txt - results/historical/mapping/GCF\_000283155.1\_CerSimSim1.0\_genomic.Sc9M7eS\_2\_HRSCAF\_41/stats/bams\_rmdup/JvS022\_79.merged.rmdup.bam.stats.txt - results/historical/mapping/GCF\_000283155.1\_CerSimSim1.0\_genomic.Sc9M7eS\_2\_HRSCAF\_41/stats/bams\_rmdup/JvS022\_80.merged.rmdup.bam.stats.txt - results/historical/mapping/GCF\_000283155.1\_CerSimSim1.0\_genomic.Sc9M7eS\_2\_HRSCAF\_41/stats/bams\_rmdup/JvS022\_81.merged.rmdup.bam.stats.txt - results/historical/mapping/GCF\_000283155.1\_CerSimSim1.0\_genomic.Sc9M7eS\_2\_HRSCAF\_41/stats/bams\_rmdup/JvS022\_82.merged.rmdup.bam.stats.txt - results/historical/mapping/GCF\_000283155.1\_CerSimSim1.0\_genomic.Sc9M7eS\_2\_HRSCAF\_41/stats/bams\_rmdup/JvS022\_83.merged.rmdup.bam.stats.txt - results/historical/mapping/GCF\_000283155.1\_CerSimSim1.0\_genomic.Sc9M7eS\_2\_HRSCAF\_41/stats/bams\_rmdup/JvS022\_84.merged.rmdup.bam.stats.txt - results/historical/mapping/GCF\_000283155.1\_CerSimSim1.0\_genomic.Sc9M7eS\_2\_HRSCAF\_41/stats/bams\_rmdup/JvS022\_85.merged.rmdup.bam.stats.txt - results/historical/mapping/GCF\_000283155.1\_CerSimSim1.0\_genomic.Sc9M7eS\_2\_HRSCAF\_41/stats/bams\_rmdup/JvS008\_08.merged.rmdup.bam.qualimap/qualimapReport.html - results/historical/mapping/GCF\_000283155.1\_CerSimSim1.0\_genomic.Sc9M7eS\_2\_HRSCAF\_41/stats/bams\_rmdup/JvS008\_10.merged.rmdup.bam.qualimap/qualimapReport.html - results/historical/mapping/GCF\_000283155.1\_CerSimSim1.0\_genomic.Sc9M7eS\_2\_HRSCAF\_41/stats/bams\_rmdup/JvS008\_11.merged.rmdup.bam.qualimap/qualimapReport.html - results/historical/mapping/GCF\_000283155.1\_CerSimSim1.0\_genomic.Sc9M7eS\_2\_HRSCAF\_41/stats/bams\_rmdup/JvS009\_09.merged.rmdup.bam.qualimap/qualimapReport.html - results/historical/mapping/GCF\_000283155.1\_CerSimSim1.0\_genomic.Sc9M7eS\_2\_HRSCAF\_41/stats/bams\_rmdup/JvS009\_15.merged.rmdup.bam.qualimap/qualimapReport.html - results/historical/mapping/GCF\_000283155.1\_CerSimSim1.0\_genomic.Sc9M7eS\_2\_HRSCAF\_41/stats/bams\_rmdup/JvS009\_19.merged.rmdup.bam.qualimap/qualimapReport.html - results/historical/mapping/GCF\_000283155.1\_CerSimSim1.0\_genomic.Sc9M7eS\_2\_HRSCAF\_41/stats/bams\_rmdup/JvS022\_01.merged.rmdup.bam.qualimap/qualimapReport.html - results/historical/mapping/GCF\_000283155.1\_CerSimSim1.0\_genomic.Sc9M7eS\_2\_HRSCAF\_41/stats/bams\_rmdup/JvS022\_02.merged.rmdup.bam.qualimap/qualimapReport.html - results/historical/mapping/GCF\_000283155.1\_CerSimSim1.0\_genomic.Sc9M7eS\_2\_HRSCAF\_41/stats/bams\_rmdup/JvS022\_03.merged.rmdup.bam.qualimap/qualimapReport.html - results/historical/mapping/GCF\_000283155.1\_CerSimSim1.0\_genomic.Sc9M7eS\_2\_HRSCAF\_41/stats/bams\_rmdup/JvS022\_04.merged.rmdup.bam.qualimap/qualimapReport.html - results/historical/mapping/GCF\_000283155.1\_CerSimSim1.0\_genomic.Sc9M7eS\_2\_HRSCAF\_41/stats/bams\_rmdup/JvS022\_05.merged.rmdup.bam.qualimap/qualimapReport.html - results/historical/mapping/GCF\_000283155.1\_CerSimSim1.0\_genomic.Sc9M7eS\_2\_HRSCAF\_41/stats/bams\_rmdup/JvS022\_06.merged.rmdup.bam.qualimap/qualimapReport.html - results/historical/mapping/GCF\_000283155.1\_CerSimSim1.0\_genomic.Sc9M7eS\_2\_HRSCAF\_41/stats/bams\_rmdup/JvS022\_07.merged.rmdup.bam.qualimap/qualimapReport.html - results/historical/mapping/GCF\_000283155.1\_CerSimSim1.0\_genomic.Sc9M7eS\_2\_HRSCAF\_41/stats/bams\_rmdup/JvS022\_08.merged.rmdup.bam.qualimap/qualimapReport.html - results/historical/mapping/GCF\_000283155.1\_CerSimSim1.0\_genomic.Sc9M7eS\_2\_HRSCAF\_41/stats/bams\_rmdup/JvS022\_09.merged.rmdup.bam.qualimap/qualimapReport.html - results/historical/mapping/GCF\_000283155.1\_CerSimSim1.0\_genomic.Sc9M7eS\_2\_HRSCAF\_41/stats/bams\_rmdup/JvS022\_10.merged.rmdup.bam.qualimap/qualimapReport.html - results/historical/mapping/GCF\_000283155.1\_CerSimSim1.0\_genomic.Sc9M7eS\_2\_HRSCAF\_41/stats/bams\_rmdup/JvS022\_11.merged.rmdup.bam.qualimap/qualimapReport.html - results/historical/mapping/GCF\_000283155.1\_CerSimSim1.0\_genomic.Sc9M7eS\_2\_HRSCAF\_41/stats/bams\_rmdup/JvS022\_12.merged.rmdup.bam.qualimap/qualimapReport.html - results/historical/mapping/GCF\_000283155.1\_CerSimSim1.0\_genomic.Sc9M7eS\_2\_HRSCAF\_41/stats/bams\_rmdup/JvS022\_74.merged.rmdup.bam.qualimap/qualimapReport.html - results/historical/mapping/GCF\_000283155.1\_CerSimSim1.0\_genomic.Sc9M7eS\_2\_HRSCAF\_41/stats/bams\_rmdup/JvS022\_75.merged.rmdup.bam.qualimap/qualimapReport.html - results/historical/mapping/GCF\_000283155.1\_CerSimSim1.0\_genomic.Sc9M7eS\_2\_HRSCAF\_41/stats/bams\_rmdup/JvS022\_76.merged.rmdup.bam.qualimap/qualimapReport.html - results/historical/mapping/GCF\_000283155.1\_CerSimSim1.0\_genomic.Sc9M7eS\_2\_HRSCAF\_41/stats/bams\_rmdup/JvS022\_77.merged.rmdup.bam.qualimap/qualimapReport.html - results/historical/mapping/GCF\_000283155.1\_CerSimSim1.0\_genomic.Sc9M7eS\_2\_HRSCAF\_41/stats/bams\_rmdup/JvS022\_78.merged.rmdup.bam.qualimap/qualimapReport.html - results/historical/mapping/GCF\_000283155.1\_CerSimSim1.0\_genomic.Sc9M7eS\_2\_HRSCAF\_41/stats/bams\_rmdup/JvS022\_79.merged.rmdup.bam.qualimap/qualimapReport.html - results/historical/mapping/GCF\_000283155.1\_CerSimSim1.0\_genomic.Sc9M7eS\_2\_HRSCAF\_41/stats/bams\_rmdup/JvS022\_80.merged.rmdup.bam.qualimap/qualimapReport.html - results/historical/mapping/GCF\_000283155.1\_CerSimSim1.0\_genomic.Sc9M7eS\_2\_HRSCAF\_41/stats/bams\_rmdup/JvS022\_81.merged.rmdup.bam.qualimap/qualimapReport.html - results/historical/mapping/GCF\_000283155.1\_CerSimSim1.0\_genomic.Sc9M7eS\_2\_HRSCAF\_41/stats/bams\_rmdup/JvS022\_82.merged.rmdup.bam.qualimap/qualimapReport.html - results/historical/mapping/GCF\_000283155.1\_CerSimSim1.0\_genomic.Sc9M7eS\_2\_HRSCAF\_41/stats/bams\_rmdup/JvS022\_83.merged.rmdup.bam.qualimap/qualimapReport.html - results/historical/mapping/GCF\_000283155.1\_CerSimSim1.0\_genomic.Sc9M7eS\_2\_HRSCAF\_41/stats/bams\_rmdup/JvS022\_84.merged.rmdup.bam.qualimap/qualimapReport.html - results/historical/mapping/GCF\_000283155.1\_CerSimSim1.0\_genomic.Sc9M7eS\_2\_HRSCAF\_41/stats/bams\_rmdup/JvS022\_85.merged.rmdup.bam.qualimap/qualimapReport.html | |
| Output files | |
| - results/historical/mapping/GCF\_000283155.1\_CerSimSim1.0\_genomic.Sc9M7eS\_2\_HRSCAF\_41/stats/bams\_rmdup/multiqc/multiqc\_report.html | |
| Container image | |
| docker://quay.io/biocontainers/multiqc:1.9--pyh9f0ad1d\_0 | |
| Code | |
| ``` 1 2 ``` \| ``` multiqc -f {params.indir} -o {params.outdir} 2> {log} ``` | |

###### Rule rmdup\_bam\_stats

×

Rule properties

|  |  |
| --- | --- |
| Jobs | 33 |
| Input files | |
| - results/{dataset}/mapping/GCF\_000283155.1\_CerSimSim1.0\_genomic.Sc9M7eS\_2\_HRSCAF\_41/{sample}\_{index}.merged.rmdup.bam - results/{dataset}/mapping/GCF\_000283155.1\_CerSimSim1.0\_genomic.Sc9M7eS\_2\_HRSCAF\_41/{sample}\_{index}.merged.rmdup.bam.bai | |
| Output files | |
| - results/{dataset}/mapping/GCF\_000283155.1\_CerSimSim1.0\_genomic.Sc9M7eS\_2\_HRSCAF\_41/stats/bams\_rmdup/{sample,[A-Za-z0-9]+}\_{index}.merged.rmdup.bam.stats.txt | |
| Container image | |
| docker://biocontainers/samtools:v1.9-4-deb\_cv1 | |
| Code | |
| ``` 1 2 ``` \| ``` samtools flagstat {input.bam} > {output.stats} 2> {log} ``` | |

###### Rule rmdup\_historical\_bams

×

Rule properties

|  |  |
| --- | --- |
| Jobs | 30 |
| Input files | |
| - results/historical/mapping/GCF\_000283155.1\_CerSimSim1.0\_genomic.Sc9M7eS\_2\_HRSCAF\_41/{sample}\_{index}.merged.bam - results/historical/mapping/GCF\_000283155.1\_CerSimSim1.0\_genomic.Sc9M7eS\_2\_HRSCAF\_41/{sample}\_{index}.merged.bam.bai | |
| Output files | |
| - results/historical/mapping/GCF\_000283155.1\_CerSimSim1.0\_genomic.Sc9M7eS\_2\_HRSCAF\_41/{sample,[A-Za-z0-9]+}\_{index}.merged.rmdup.bam | |
| Container image | |
| docker://biocontainers/samtools:v1.9-4-deb\_cv1 | |
| Code | |
| ``` 1 2 ``` \| ``` samtools view -@ {threads} -h {input.merged} \| python3 workflow/scripts/samremovedup.py \| samtools view -b -o {output.rmdup} 2> {log} ``` | |

###### Rule index\_rmdup\_bams

×

Rule properties

|  |  |
| --- | --- |
| Jobs | 33 |
| Input files | |
| - results/{dataset}/mapping/GCF\_000283155.1\_CerSimSim1.0\_genomic.Sc9M7eS\_2\_HRSCAF\_41/{sample}\_{index}.merged.rmdup.bam | |
| Output files | |
| - results/{dataset}/mapping/GCF\_000283155.1\_CerSimSim1.0\_genomic.Sc9M7eS\_2\_HRSCAF\_41/{sample,[A-Za-z0-9]+}\_{index}.merged.rmdup.bam.bai | |
| Container image | |
| docker://biocontainers/samtools:v1.9-4-deb\_cv1 | |
| Code | |
| ``` 1 2 ``` \| ``` samtools index {input.bam} {output.index} 2> {log} ``` | |

###### Rule rmdup\_bam\_qualimap

×

Rule properties

|  |  |
| --- | --- |
| Jobs | 33 |
| Input files | |
| - results/{dataset}/mapping/GCF\_000283155.1\_CerSimSim1.0\_genomic.Sc9M7eS\_2\_HRSCAF\_41/{sample}\_{index}.merged.rmdup.bam - results/{dataset}/mapping/GCF\_000283155.1\_CerSimSim1.0\_genomic.Sc9M7eS\_2\_HRSCAF\_41/{sample}\_{index}.merged.rmdup.bam.bai | |
| Output files | |
| - results/{dataset}/mapping/GCF\_000283155.1\_CerSimSim1.0\_genomic.Sc9M7eS\_2\_HRSCAF\_41/stats/bams\_rmdup/{sample,[A-Za-z0-9]+}\_{index}.merged.rmdup.bam.qualimap/qualimapReport.html - results/{dataset}/mapping/GCF\_000283155.1\_CerSimSim1.0\_genomic.Sc9M7eS\_2\_HRSCAF\_41/stats/bams\_rmdup/{sample,[A-Za-z0-9]+}\_{index}.merged.rmdup.bam.qualimap/genome\_results.txt - results/{dataset}/mapping/GCF\_000283155.1\_CerSimSim1.0\_genomic.Sc9M7eS\_2\_HRSCAF\_41/stats/bams\_rmdup/{sample,[A-Za-z0-9]+}\_{index}.merged.rmdup.bam.qualimap | |
| Container image | |
| docker://quay.io/biocontainers/qualimap:2.2.2d--1 | |
| Code | |
| ``` 1 2 3 4 ``` \| ``` mem=$(((6 * {threads}) - 2)) unset DISPLAY qualimap bamqc -bam {input.bam} --java-mem-size=${{mem}}G -nt {threads} -outdir {params.outdir} -outformat html 2> {log} ``` | |

###### Rule historical\_merged\_sample\_bam\_multiqc

×

Rule properties

|  |  |
| --- | --- |
| Jobs | 1 |
| Input files | |
| - results/historical/mapping/GCF\_000283155.1\_CerSimSim1.0\_genomic.Sc9M7eS\_2\_HRSCAF\_41/stats/bams\_merged\_sample/JvS008.merged.rmdup.merged.bam.stats.txt - results/historical/mapping/GCF\_000283155.1\_CerSimSim1.0\_genomic.Sc9M7eS\_2\_HRSCAF\_41/stats/bams\_merged\_sample/JvS009.merged.rmdup.merged.bam.stats.txt - results/historical/mapping/GCF\_000283155.1\_CerSimSim1.0\_genomic.Sc9M7eS\_2\_HRSCAF\_41/stats/bams\_merged\_sample/JvS022.merged.rmdup.merged.bam.stats.txt - results/historical/mapping/GCF\_000283155.1\_CerSimSim1.0\_genomic.Sc9M7eS\_2\_HRSCAF\_41/stats/bams\_merged\_sample/JvS008.merged.rmdup.merged.bam.qualimap/qualimapReport.html - results/historical/mapping/GCF\_000283155.1\_CerSimSim1.0\_genomic.Sc9M7eS\_2\_HRSCAF\_41/stats/bams\_merged\_sample/JvS009.merged.rmdup.merged.bam.qualimap/qualimapReport.html - results/historical/mapping/GCF\_000283155.1\_CerSimSim1.0\_genomic.Sc9M7eS\_2\_HRSCAF\_41/stats/bams\_merged\_sample/JvS022.merged.rmdup.merged.bam.qualimap/qualimapReport.html | |
| Output files | |
| - results/historical/mapping/GCF\_000283155.1\_CerSimSim1.0\_genomic.Sc9M7eS\_2\_HRSCAF\_41/stats/bams\_merged\_sample/multiqc/multiqc\_report.html | |
| Container image | |
| docker://quay.io/biocontainers/multiqc:1.9--pyh9f0ad1d\_0 | |
| Code | |
| ``` 1 2 ``` \| ``` multiqc -f {params.indir} -o {params.outdir} 2> {log} ``` | |

###### Rule merged\_sample\_bam\_stats

×

Rule properties

|  |  |
| --- | --- |
| Jobs | 6 |
| Input files | |
| - results/{dataset}/mapping/GCF\_000283155.1\_CerSimSim1.0\_genomic.Sc9M7eS\_2\_HRSCAF\_41/{sample}.merged.rmdup.merged.bam - results/{dataset}/mapping/GCF\_000283155.1\_CerSimSim1.0\_genomic.Sc9M7eS\_2\_HRSCAF\_41/{sample}.merged.rmdup.merged.bam.bai | |
| Output files | |
| - results/{dataset}/mapping/GCF\_000283155.1\_CerSimSim1.0\_genomic.Sc9M7eS\_2\_HRSCAF\_41/stats/bams\_merged\_sample/{sample,[A-Za-z0-9]+}.merged.rmdup.merged.bam.stats.txt | |
| Container image | |
| docker://biocontainers/samtools:v1.9-4-deb\_cv1 | |
| Code | |
| ``` 1 2 ``` \| ``` samtools flagstat {input.bam} > {output.stats} 2> {log} ``` | |

###### Rule merge\_historical\_bams\_per\_sample

×

Rule properties

|  |  |
| --- | --- |
| Jobs | 3 |
| Input files | |
| Output files | |
| - results/historical/mapping/GCF\_000283155.1\_CerSimSim1.0\_genomic.Sc9M7eS\_2\_HRSCAF\_41/{sample,[A-Za-z0-9]+}.merged.rmdup.merged.bam | |
| Container image | |
| docker://biocontainers/samtools:v1.9-4-deb\_cv1 | |
| Code | |
| ``` 1 2 3 4 5 6 7 8 9 ``` \| ``` files=`echo {input} \| awk '{{print NF}}'` if [ $files -gt 1 ] # check if there are at least 2 files for merging. If there is only one file, copy the sorted bam file. then samtools merge {output.merged} {input} 2> {log} else cp {input} {output.merged} && touch {output.merged} 2> {log} echo "Only one file present for merging. Copying the input bam file." >> {log} fi ``` | |

###### Rule index\_merged\_sample\_bams

×

Rule properties

|  |  |
| --- | --- |
| Jobs | 6 |
| Input files | |
| - results/{dataset}/mapping/GCF\_000283155.1\_CerSimSim1.0\_genomic.Sc9M7eS\_2\_HRSCAF\_41/{sample}.merged.rmdup.merged.bam | |
| Output files | |
| - results/{dataset}/mapping/GCF\_000283155.1\_CerSimSim1.0\_genomic.Sc9M7eS\_2\_HRSCAF\_41/{sample,[A-Za-z0-9]+}.merged.rmdup.merged.bam.bai | |
| Container image | |
| docker://biocontainers/samtools:v1.9-4-deb\_cv1 | |
| Code | |
| ``` 1 2 ``` \| ``` samtools index {input.bam} {output.index} 2> {log} ``` | |

###### Rule merged\_sample\_bam\_qualimap

×

Rule properties

|  |  |
| --- | --- |
| Jobs | 6 |
| Input files | |
| - results/{dataset}/mapping/GCF\_000283155.1\_CerSimSim1.0\_genomic.Sc9M7eS\_2\_HRSCAF\_41/{sample}.merged.rmdup.merged.bam - results/{dataset}/mapping/GCF\_000283155.1\_CerSimSim1.0\_genomic.Sc9M7eS\_2\_HRSCAF\_41/{sample}.merged.rmdup.merged.bam.bai | |
| Output files | |
| - results/{dataset}/mapping/GCF\_000283155.1\_CerSimSim1.0\_genomic.Sc9M7eS\_2\_HRSCAF\_41/stats/bams\_merged\_sample/{sample,[A-Za-z0-9]+}.merged.rmdup.merged.bam.qualimap/qualimapReport.html - results/{dataset}/mapping/GCF\_000283155.1\_CerSimSim1.0\_genomic.Sc9M7eS\_2\_HRSCAF\_41/stats/bams\_merged\_sample/{sample,[A-Za-z0-9]+}.merged.rmdup.merged.bam.qualimap/genome\_results.txt - results/{dataset}/mapping/GCF\_000283155.1\_CerSimSim1.0\_genomic.Sc9M7eS\_2\_HRSCAF\_41/stats/bams\_merged\_sample/{sample,[A-Za-z0-9]+}.merged.rmdup.merged.bam.qualimap | |
| Container image | |
| docker://quay.io/biocontainers/qualimap:2.2.2d--1 | |
| Code | |
| ``` 1 2 3 4 ``` \| ``` mem=$(((6 * {threads}) - 2)) unset DISPLAY qualimap bamqc -bam {input.bam} --java-mem-size=${{mem}}G -nt {threads} -outdir {params.outdir} -outformat html 2> {log} ``` | |

###### Rule historical\_realigned\_bam\_multiqc

×

Rule properties

|  |  |
| --- | --- |
| Jobs | 1 |
| Input files | |
| - results/historical/mapping/GCF\_000283155.1\_CerSimSim1.0\_genomic.Sc9M7eS\_2\_HRSCAF\_41/stats/bams\_indels\_realigned/JvS008.merged.rmdup.merged.realn.repma.Q30.bam.dp.hist.pdf - results/historical/mapping/GCF\_000283155.1\_CerSimSim1.0\_genomic.Sc9M7eS\_2\_HRSCAF\_41/stats/bams\_indels\_realigned/JvS009.merged.rmdup.merged.realn.repma.Q30.bam.dp.hist.pdf - results/historical/mapping/GCF\_000283155.1\_CerSimSim1.0\_genomic.Sc9M7eS\_2\_HRSCAF\_41/stats/bams\_indels\_realigned/JvS022.merged.rmdup.merged.realn.repma.Q30.bam.dp.hist.pdf - results/historical/mapping/GCF\_000283155.1\_CerSimSim1.0\_genomic.Sc9M7eS\_2\_HRSCAF\_41/stats/bams\_indels\_realigned/JvS008.merged.rmdup.merged.realn.bam.stats.txt - results/historical/mapping/GCF\_000283155.1\_CerSimSim1.0\_genomic.Sc9M7eS\_2\_HRSCAF\_41/stats/bams\_indels\_realigned/JvS009.merged.rmdup.merged.realn.bam.stats.txt - results/historical/mapping/GCF\_000283155.1\_CerSimSim1.0\_genomic.Sc9M7eS\_2\_HRSCAF\_41/stats/bams\_indels\_realigned/JvS022.merged.rmdup.merged.realn.bam.stats.txt - results/historical/mapping/GCF\_000283155.1\_CerSimSim1.0\_genomic.Sc9M7eS\_2\_HRSCAF\_41/stats/bams\_indels\_realigned/JvS008.merged.rmdup.merged.realn.bam.qualimap/qualimapReport.html - results/historical/mapping/GCF\_000283155.1\_CerSimSim1.0\_genomic.Sc9M7eS\_2\_HRSCAF\_41/stats/bams\_indels\_realigned/JvS009.merged.rmdup.merged.realn.bam.qualimap/qualimapReport.html - results/historical/mapping/GCF\_000283155.1\_CerSimSim1.0\_genomic.Sc9M7eS\_2\_HRSCAF\_41/stats/bams\_indels\_realigned/JvS022.merged.rmdup.merged.realn.bam.qualimap/qualimapReport.html - results/historical/mapping/GCF\_000283155.1\_CerSimSim1.0\_genomic.Sc9M7eS\_2\_HRSCAF\_41/stats/bams\_indels\_realigned/fastqc/JvS008.merged.rmdup.merged.realn\_fastqc.html - results/historical/mapping/GCF\_000283155.1\_CerSimSim1.0\_genomic.Sc9M7eS\_2\_HRSCAF\_41/stats/bams\_indels\_realigned/fastqc/JvS009.merged.rmdup.merged.realn\_fastqc.html - results/historical/mapping/GCF\_000283155.1\_CerSimSim1.0\_genomic.Sc9M7eS\_2\_HRSCAF\_41/stats/bams\_indels\_realigned/fastqc/JvS022.merged.rmdup.merged.realn\_fastqc.html - results/historical/mapping/GCF\_000283155.1\_CerSimSim1.0\_genomic.Sc9M7eS\_2\_HRSCAF\_41/stats/bams\_indels\_realigned/fastqc/JvS008.merged.rmdup.merged.realn\_fastqc.zip - results/historical/mapping/GCF\_000283155.1\_CerSimSim1.0\_genomic.Sc9M7eS\_2\_HRSCAF\_41/stats/bams\_indels\_realigned/fastqc/JvS009.merged.rmdup.merged.realn\_fastqc.zip - results/historical/mapping/GCF\_000283155.1\_CerSimSim1.0\_genomic.Sc9M7eS\_2\_HRSCAF\_41/stats/bams\_indels\_realigned/fastqc/JvS022.merged.rmdup.merged.realn\_fastqc.zip - results/historical/mapping/GCF\_000283155.1\_CerSimSim1.0\_genomic.Sc9M7eS\_2\_HRSCAF\_41/stats/bams\_indels\_realigned/JvS008.merged.rmdup.merged.realn.repma.Q30.bam.dp.hist.pdf - results/historical/mapping/GCF\_000283155.1\_CerSimSim1.0\_genomic.Sc9M7eS\_2\_HRSCAF\_41/stats/bams\_indels\_realigned/JvS009.merged.rmdup.merged.realn.repma.Q30.bam.dp.hist.pdf - results/historical/mapping/GCF\_000283155.1\_CerSimSim1.0\_genomic.Sc9M7eS\_2\_HRSCAF\_41/stats/bams\_indels\_realigned/JvS022.merged.rmdup.merged.realn.repma.Q30.bam.dp.hist.pdf | |
| Output files | |
| - results/historical/mapping/GCF\_000283155.1\_CerSimSim1.0\_genomic.Sc9M7eS\_2\_HRSCAF\_41/stats/bams\_indels\_realigned/multiqc/multiqc\_report.html | |
| Container image | |
| docker://quay.io/biocontainers/multiqc:1.9--pyh9f0ad1d\_0 | |
| Code | |
| ``` 1 2 ``` \| ``` multiqc -f {params.indir} -o {params.outdir} 2> {log} ``` | |

###### Rule plot\_dp\_hist

×

Rule properties

|  |  |
| --- | --- |
| Jobs | 6 |
| Input files | |
| - results/{dataset}/mapping/GCF\_000283155.1\_CerSimSim1.0\_genomic.Sc9M7eS\_2\_HRSCAF\_41/stats/bams\_indels\_realigned/{sample}.merged.rmdup.merged.realn.repma.Q30.bam.dpstats.txt - results/{dataset}/mapping/GCF\_000283155.1\_CerSimSim1.0\_genomic.Sc9M7eS\_2\_HRSCAF\_41/stats/bams\_indels\_realigned/{sample}.merged.rmdup.merged.realn.repma.Q30.bam.dp | |
| Output files | |
| - results/{dataset}/mapping/GCF\_000283155.1\_CerSimSim1.0\_genomic.Sc9M7eS\_2\_HRSCAF\_41/stats/bams\_indels\_realigned/{sample,[A-Za-z0-9]+}.merged.rmdup.merged.realn.repma.Q30.bam.dp.hist.pdf | |

###### Rule realigned\_bam\_depth

×

Rule properties

|  |  |
| --- | --- |
| Jobs | 6 |
| Input files | |
| - results/{dataset}/mapping/GCF\_000283155.1\_CerSimSim1.0\_genomic.Sc9M7eS\_2\_HRSCAF\_41/{sample}.merged.rmdup.merged.realn.bam - /proj/sllstore2017093/b2016342/b2016342\_nobackup/lts/genome\_erosion\_pipeline/verena\_testing/testdata/reference/GCF\_000283155.1\_CerSimSim1.0\_genomic.Sc9M7eS\_2\_HRSCAF\_41.repma.bed | |
| Output files | |
| - results/{dataset}/mapping/GCF\_000283155.1\_CerSimSim1.0\_genomic.Sc9M7eS\_2\_HRSCAF\_41/stats/bams\_indels\_realigned/{sample,[A-Za-z0-9]+}.merged.rmdup.merged.realn.repma.Q30.bam.dp - results/{dataset}/mapping/GCF\_000283155.1\_CerSimSim1.0\_genomic.Sc9M7eS\_2\_HRSCAF\_41/stats/bams\_indels\_realigned/{sample,[A-Za-z0-9]+}.merged.rmdup.merged.realn.repma.Q30.bam.dpstats.txt | |
| Container image | |
| docker://biocontainers/samtools:v1.9-4-deb\_cv1 | |
| Code | |
| ``` 1 2 3 4 5 6 7 8 9 10 ``` \| ``` if [ {params.cov} = "True" ] # include sites with missing data / zero coverage then samtools depth -a -Q 30 -q 30 -b {input.no_rep_bed} {input.bam} > {output.tmp} 2> {log} && awk '{{sum+=$3}} END {{ print sum/NR }}' {output.tmp} \| awk -v min={params.minDP} -v max={params.maxDP} '{{ printf "%.0f %.0f %.0f", $1, $1*min, $1*max }}' > {output.dp} 2>> {log} elif [ {params.cov} = "False" ] # exclude sites with missing data / zero coverage then samtools depth -Q 30 -q 30 -b {input.no_rep_bed} {input.bam} > {output.tmp} 2> {log} && awk '{{sum+=$3}} END {{ print sum/NR }}' {output.tmp} \| awk -v min={params.minDP} -v max={params.maxDP} '{{ printf "%.0f %.0f %.0f", $1, $1*min, $1*max }}' > {output.dp} 2>> {log} fi ``` | |

###### Rule indel\_realigner

×

Rule properties

|  |  |
| --- | --- |
| Jobs | 6 |
| Input files | |
| - /proj/sllstore2017093/b2016342/b2016342\_nobackup/lts/genome\_erosion\_pipeline/verena\_testing/testdata/reference/GCF\_000283155.1\_CerSimSim1.0\_genomic.Sc9M7eS\_2\_HRSCAF\_41.fasta - /proj/sllstore2017093/b2016342/b2016342\_nobackup/lts/genome\_erosion\_pipeline/verena\_testing/testdata/reference/GCF\_000283155.1\_CerSimSim1.0\_genomic.Sc9M7eS\_2\_HRSCAF\_41.dict - /proj/sllstore2017093/b2016342/b2016342\_nobackup/lts/genome\_erosion\_pipeline/verena\_testing/testdata/reference/GCF\_000283155.1\_CerSimSim1.0\_genomic.Sc9M7eS\_2\_HRSCAF\_41.fasta.fai - results/{dataset}/mapping/GCF\_000283155.1\_CerSimSim1.0\_genomic.Sc9M7eS\_2\_HRSCAF\_41/{sample}.merged.rmdup.merged.bam - results/{dataset}/mapping/GCF\_000283155.1\_CerSimSim1.0\_genomic.Sc9M7eS\_2\_HRSCAF\_41/{sample}.merged.rmdup.merged.bam.bai - results/{dataset}/mapping/GCF\_000283155.1\_CerSimSim1.0\_genomic.Sc9M7eS\_2\_HRSCAF\_41/{sample}.merged.rmdup.merged.realn\_targets.list | |
| Output files | |
| - results/{dataset}/mapping/GCF\_000283155.1\_CerSimSim1.0\_genomic.Sc9M7eS\_2\_HRSCAF\_41/{sample,[A-Za-z0-9]+}.merged.rmdup.merged.realn.bam | |
| Container image | |
| docker://broadinstitute/gatk3:3.7-0 | |
| Code | |
| ``` 1 2 ``` \| ``` java -jar /usr/GenomeAnalysisTK.jar -T IndelRealigner -R {input.ref} -I {input.bam} -targetIntervals {input.target_list} -o {output.realigned} 2> {log} ``` | |

###### Rule indel\_realigner\_targets

×

Rule properties

|  |  |
| --- | --- |
| Jobs | 6 |
| Input files | |
| - /proj/sllstore2017093/b2016342/b2016342\_nobackup/lts/genome\_erosion\_pipeline/verena\_testing/testdata/reference/GCF\_000283155.1\_CerSimSim1.0\_genomic.Sc9M7eS\_2\_HRSCAF\_41.fasta - /proj/sllstore2017093/b2016342/b2016342\_nobackup/lts/genome\_erosion\_pipeline/verena\_testing/testdata/reference/GCF\_000283155.1\_CerSimSim1.0\_genomic.Sc9M7eS\_2\_HRSCAF\_41.dict - /proj/sllstore2017093/b2016342/b2016342\_nobackup/lts/genome\_erosion\_pipeline/verena\_testing/testdata/reference/GCF\_000283155.1\_CerSimSim1.0\_genomic.Sc9M7eS\_2\_HRSCAF\_41.fasta.fai - results/{dataset}/mapping/GCF\_000283155.1\_CerSimSim1.0\_genomic.Sc9M7eS\_2\_HRSCAF\_41/{sample}.merged.rmdup.merged.bam - results/{dataset}/mapping/GCF\_000283155.1\_CerSimSim1.0\_genomic.Sc9M7eS\_2\_HRSCAF\_41/{sample}.merged.rmdup.merged.bam.bai | |
| Output files | |
| - results/{dataset}/mapping/GCF\_000283155.1\_CerSimSim1.0\_genomic.Sc9M7eS\_2\_HRSCAF\_41/{sample,[A-Za-z0-9]+}.merged.rmdup.merged.realn\_targets.list | |
| Container image | |
| docker://broadinstitute/gatk3:3.7-0 | |
| Code | |
| ``` 1 2 ``` \| ``` java -jar /usr/GenomeAnalysisTK.jar -T RealignerTargetCreator -R {input.ref} -I {input.bam} -o {output.target_list} -nt {threads} 2> {log} ``` | |

###### Rule realigned\_bam\_stats

×

Rule properties

|  |  |
| --- | --- |
| Jobs | 6 |
| Input files | |
| - results/{dataset}/mapping/GCF\_000283155.1\_CerSimSim1.0\_genomic.Sc9M7eS\_2\_HRSCAF\_41/{sample}.merged.rmdup.merged.realn.bam - results/{dataset}/mapping/GCF\_000283155.1\_CerSimSim1.0\_genomic.Sc9M7eS\_2\_HRSCAF\_41/{sample}.merged.rmdup.merged.realn.bam.bai | |
| Output files | |
| - results/{dataset}/mapping/GCF\_000283155.1\_CerSimSim1.0\_genomic.Sc9M7eS\_2\_HRSCAF\_41/stats/bams\_indels\_realigned/{sample,[A-Za-z0-9]+}.merged.rmdup.merged.realn.bam.stats.txt | |
| Container image | |
| docker://biocontainers/samtools:v1.9-4-deb\_cv1 | |
| Code | |
| ``` 1 2 ``` \| ``` samtools flagstat {input.bam} > {output.stats} 2> {log} ``` | |

###### Rule index\_realigned\_bams

×

Rule properties

|  |  |
| --- | --- |
| Jobs | 6 |
| Input files | |
| - results/{dataset}/mapping/GCF\_000283155.1\_CerSimSim1.0\_genomic.Sc9M7eS\_2\_HRSCAF\_41/{sample}.merged.rmdup.merged.bam | |
| Output files | |
| - results/{dataset}/mapping/GCF\_000283155.1\_CerSimSim1.0\_genomic.Sc9M7eS\_2\_HRSCAF\_41/{sample,[A-Za-z0-9]+}.merged.rmdup.merged.realn.bam.bai | |
| Container image | |
| docker://biocontainers/samtools:v1.9-4-deb\_cv1 | |
| Code | |
| ``` 1 2 ``` \| ``` samtools index {input.bam} {output.index} 2> {log} ``` | |

###### Rule realigned\_bam\_qualimap

×

Rule properties

|  |  |
| --- | --- |
| Jobs | 6 |
| Input files | |
| - results/{dataset}/mapping/GCF\_000283155.1\_CerSimSim1.0\_genomic.Sc9M7eS\_2\_HRSCAF\_41/{sample}.merged.rmdup.merged.realn.bam - results/{dataset}/mapping/GCF\_000283155.1\_CerSimSim1.0\_genomic.Sc9M7eS\_2\_HRSCAF\_41/{sample}.merged.rmdup.merged.realn.bam.bai | |
| Output files | |
| - results/{dataset}/mapping/GCF\_000283155.1\_CerSimSim1.0\_genomic.Sc9M7eS\_2\_HRSCAF\_41/stats/bams\_indels\_realigned/{sample,[A-Za-z0-9]+}.merged.rmdup.merged.realn.bam.qualimap/qualimapReport.html - results/{dataset}/mapping/GCF\_000283155.1\_CerSimSim1.0\_genomic.Sc9M7eS\_2\_HRSCAF\_41/stats/bams\_indels\_realigned/{sample,[A-Za-z0-9]+}.merged.rmdup.merged.realn.bam.qualimap/genome\_results.txt - results/{dataset}/mapping/GCF\_000283155.1\_CerSimSim1.0\_genomic.Sc9M7eS\_2\_HRSCAF\_41/stats/bams\_indels\_realigned/{sample,[A-Za-z0-9]+}.merged.rmdup.merged.realn.bam.qualimap | |
| Container image | |
| docker://quay.io/biocontainers/qualimap:2.2.2d--1 | |
| Code | |
| ``` 1 2 3 4 ``` \| ``` mem=$(((6 * {threads}) - 2)) unset DISPLAY qualimap bamqc -bam {input.bam} --java-mem-size=${{mem}}G -nt {threads} -outdir {params.outdir} -outformat html 2> {log} ``` | |

###### Rule realigned\_bam\_fastqc

×

Rule properties

|  |  |
| --- | --- |
| Jobs | 6 |
| Input files | |
| - results/{dataset}/mapping/GCF\_000283155.1\_CerSimSim1.0\_genomic.Sc9M7eS\_2\_HRSCAF\_41/{sample}.merged.rmdup.merged.realn.bam | |
| Output files | |
| - results/{dataset}/mapping/GCF\_000283155.1\_CerSimSim1.0\_genomic.Sc9M7eS\_2\_HRSCAF\_41/stats/bams\_indels\_realigned/fastqc/{sample,[A-Za-z0-9]+}.merged.rmdup.merged.realn\_fastqc.html - results/{dataset}/mapping/GCF\_000283155.1\_CerSimSim1.0\_genomic.Sc9M7eS\_2\_HRSCAF\_41/stats/bams\_indels\_realigned/fastqc/{sample,[A-Za-z0-9]+}.merged.rmdup.merged.realn\_fastqc.zip - results/{dataset}/mapping/GCF\_000283155.1\_CerSimSim1.0\_genomic.Sc9M7eS\_2\_HRSCAF\_41/stats/bams\_indels\_realigned/fastqc/{sample,[A-Za-z0-9]+}.merged.rmdup.merged.realn\_fastqc | |
| Container image | |
| docker://biocontainers/fastqc:v0.11.9\_cv7 | |
| Code | |
| ``` 1 2 ``` \| ``` fastqc -o {params.dir} -t {threads} --extract {input.bam} 2> {log} ``` | |

###### Rule modern\_merged\_index\_bam\_multiqc

×

Rule properties

|  |  |
| --- | --- |
| Jobs | 1 |
| Input files | |
| - results/modern/mapping/GCF\_000283155.1\_CerSimSim1.0\_genomic.Sc9M7eS\_2\_HRSCAF\_41/stats/bams\_merged\_index/JvS033\_01.merged.bam.stats.txt - results/modern/mapping/GCF\_000283155.1\_CerSimSim1.0\_genomic.Sc9M7eS\_2\_HRSCAF\_41/stats/bams\_merged\_index/JvS034\_02.merged.bam.stats.txt - results/modern/mapping/GCF\_000283155.1\_CerSimSim1.0\_genomic.Sc9M7eS\_2\_HRSCAF\_41/stats/bams\_merged\_index/JvS035\_14.merged.bam.stats.txt - results/modern/mapping/GCF\_000283155.1\_CerSimSim1.0\_genomic.Sc9M7eS\_2\_HRSCAF\_41/stats/bams\_merged\_index/JvS033\_01.merged.bam.qualimap/qualimapReport.html - results/modern/mapping/GCF\_000283155.1\_CerSimSim1.0\_genomic.Sc9M7eS\_2\_HRSCAF\_41/stats/bams\_merged\_index/JvS034\_02.merged.bam.qualimap/qualimapReport.html - results/modern/mapping/GCF\_000283155.1\_CerSimSim1.0\_genomic.Sc9M7eS\_2\_HRSCAF\_41/stats/bams\_merged\_index/JvS035\_14.merged.bam.qualimap/qualimapReport.html | |
| Output files | |
| - results/modern/mapping/GCF\_000283155.1\_CerSimSim1.0\_genomic.Sc9M7eS\_2\_HRSCAF\_41/stats/bams\_merged\_index/multiqc/multiqc\_report.html | |
| Container image | |
| docker://quay.io/biocontainers/multiqc:1.9--pyh9f0ad1d\_0 | |
| Code | |
| ``` 1 2 ``` \| ``` multiqc -f {params.indir} -o {params.outdir} 2> {log} ``` | |

###### Rule merge\_modern\_bams\_per\_index

×

Rule properties

|  |  |
| --- | --- |
| Jobs | 3 |
| Input files | |
| Output files | |
| - results/modern/mapping/GCF\_000283155.1\_CerSimSim1.0\_genomic.Sc9M7eS\_2\_HRSCAF\_41/{sample,[A-Za-z0-9]+}\_{index}.merged.bam | |
| Container image | |
| docker://biocontainers/samtools:v1.9-4-deb\_cv1 | |
| Code | |
| ``` 1 2 3 4 5 6 7 8 9 ``` \| ``` files=`echo {input} \| awk '{{print NF}}'` if [ $files -gt 1 ] # check if there are at least 2 files for merging. If there is only one file, copy the sorted bam file. then samtools merge {output.merged} {input} 2> {log} else cp {input} {output.merged} && touch {output.merged} 2> {log} echo "Only one file present for merging. Copying the input bam file." >> {log} fi ``` | |

###### Rule modern\_rmdup\_bam\_multiqc

×

Rule properties

|  |  |
| --- | --- |
| Jobs | 1 |
| Input files | |
| - results/modern/mapping/GCF\_000283155.1\_CerSimSim1.0\_genomic.Sc9M7eS\_2\_HRSCAF\_41/stats/bams\_rmdup/JvS033\_01.merged.rmdup.bam.stats.txt - results/modern/mapping/GCF\_000283155.1\_CerSimSim1.0\_genomic.Sc9M7eS\_2\_HRSCAF\_41/stats/bams\_rmdup/JvS034\_02.merged.rmdup.bam.stats.txt - results/modern/mapping/GCF\_000283155.1\_CerSimSim1.0\_genomic.Sc9M7eS\_2\_HRSCAF\_41/stats/bams\_rmdup/JvS035\_14.merged.rmdup.bam.stats.txt - results/modern/mapping/GCF\_000283155.1\_CerSimSim1.0\_genomic.Sc9M7eS\_2\_HRSCAF\_41/stats/bams\_rmdup/JvS033\_01.merged.rmdup.bam.qualimap/qualimapReport.html - results/modern/mapping/GCF\_000283155.1\_CerSimSim1.0\_genomic.Sc9M7eS\_2\_HRSCAF\_41/stats/bams\_rmdup/JvS034\_02.merged.rmdup.bam.qualimap/qualimapReport.html - results/modern/mapping/GCF\_000283155.1\_CerSimSim1.0\_genomic.Sc9M7eS\_2\_HRSCAF\_41/stats/bams\_rmdup/JvS035\_14.merged.rmdup.bam.qualimap/qualimapReport.html | |
| Output files | |
| - results/modern/mapping/GCF\_000283155.1\_CerSimSim1.0\_genomic.Sc9M7eS\_2\_HRSCAF\_41/stats/bams\_rmdup/multiqc/multiqc\_report.html | |
| Container image | |
| docker://quay.io/biocontainers/multiqc:1.9--pyh9f0ad1d\_0 | |
| Code | |
| ``` 1 2 ``` \| ``` multiqc -f {params.indir} -o {params.outdir} 2> {log} ``` | |

###### Rule rmdup\_modern\_bams

×

Rule properties

|  |  |
| --- | --- |
| Jobs | 3 |
| Input files | |
| - results/modern/mapping/GCF\_000283155.1\_CerSimSim1.0\_genomic.Sc9M7eS\_2\_HRSCAF\_41/{sample}\_{index}.merged.bam - results/modern/mapping/GCF\_000283155.1\_CerSimSim1.0\_genomic.Sc9M7eS\_2\_HRSCAF\_41/{sample}\_{index}.merged.bam.bai | |
| Output files | |
| - results/modern/mapping/GCF\_000283155.1\_CerSimSim1.0\_genomic.Sc9M7eS\_2\_HRSCAF\_41/{sample,[A-Za-z0-9]+}\_{index}.merged.rmdup.bam - results/modern/mapping/GCF\_000283155.1\_CerSimSim1.0\_genomic.Sc9M7eS\_2\_HRSCAF\_41/{sample,[A-Za-z0-9]+}\_{index}.merged.rmdup\_metrics.txt | |
| Container image | |
| docker://quay.io/biocontainers/picard:2.26.6--hdfd78af\_0 | |
| Code | |
| ``` 1 2 3 ``` \| ``` mem=$(((6 * {threads}) - 2)) picard MarkDuplicates -Xmx${{mem}}g INPUT={input.merged} OUTPUT={output.rmdup} METRICS_FILE={output.metrix} 2> {log} ``` | |

###### Rule modern\_merged\_sample\_bam\_multiqc

×

Rule properties

|  |  |
| --- | --- |
| Jobs | 1 |
| Input files | |
| - results/modern/mapping/GCF\_000283155.1\_CerSimSim1.0\_genomic.Sc9M7eS\_2\_HRSCAF\_41/stats/bams\_merged\_sample/JvS033.merged.rmdup.merged.bam.stats.txt - results/modern/mapping/GCF\_000283155.1\_CerSimSim1.0\_genomic.Sc9M7eS\_2\_HRSCAF\_41/stats/bams\_merged\_sample/JvS034.merged.rmdup.merged.bam.stats.txt - results/modern/mapping/GCF\_000283155.1\_CerSimSim1.0\_genomic.Sc9M7eS\_2\_HRSCAF\_41/stats/bams\_merged\_sample/JvS035.merged.rmdup.merged.bam.stats.txt - results/modern/mapping/GCF\_000283155.1\_CerSimSim1.0\_genomic.Sc9M7eS\_2\_HRSCAF\_41/stats/bams\_merged\_sample/JvS033.merged.rmdup.merged.bam.qualimap/qualimapReport.html - results/modern/mapping/GCF\_000283155.1\_CerSimSim1.0\_genomic.Sc9M7eS\_2\_HRSCAF\_41/stats/bams\_merged\_sample/JvS034.merged.rmdup.merged.bam.qualimap/qualimapReport.html - results/modern/mapping/GCF\_000283155.1\_CerSimSim1.0\_genomic.Sc9M7eS\_2\_HRSCAF\_41/stats/bams\_merged\_sample/JvS035.merged.rmdup.merged.bam.qualimap/qualimapReport.html | |
| Output files | |
| - results/modern/mapping/GCF\_000283155.1\_CerSimSim1.0\_genomic.Sc9M7eS\_2\_HRSCAF\_41/stats/bams\_merged\_sample/multiqc/multiqc\_report.html | |
| Container image | |
| docker://quay.io/biocontainers/multiqc:1.9--pyh9f0ad1d\_0 | |
| Code | |
| ``` 1 2 ``` \| ``` multiqc -f {params.indir} -o {params.outdir} 2> {log} ``` | |

###### Rule merge\_modern\_bams\_per\_sample

×

Rule properties

|  |  |
| --- | --- |
| Jobs | 3 |
| Input files | |
| Output files | |
| - results/modern/mapping/GCF\_000283155.1\_CerSimSim1.0\_genomic.Sc9M7eS\_2\_HRSCAF\_41/{sample,[A-Za-z0-9]+}.merged.rmdup.merged.bam | |
| Container image | |
| docker://biocontainers/samtools:v1.9-4-deb\_cv1 | |
| Code | |
| ``` 1 2 3 4 5 6 7 8 9 ``` \| ``` files=`echo {input} \| awk '{{print NF}}'` if [ $files -gt 1 ] # check if there are at least 2 files for merging. If there is only one file, copy the sorted bam file. then samtools merge {output.merged} {input} 2> {log} else cp {input} {output.merged} && touch {output.merged} 2> {log} echo "Only one file present for merging. Copying the input bam file." >> {log} fi ``` | |

###### Rule modern\_realigned\_bam\_multiqc

×

Rule properties

|  |  |
| --- | --- |
| Jobs | 1 |
| Input files | |
| - results/modern/mapping/GCF\_000283155.1\_CerSimSim1.0\_genomic.Sc9M7eS\_2\_HRSCAF\_41/stats/bams\_indels\_realigned/JvS033.merged.rmdup.merged.realn.repma.Q30.bam.dp.hist.pdf - results/modern/mapping/GCF\_000283155.1\_CerSimSim1.0\_genomic.Sc9M7eS\_2\_HRSCAF\_41/stats/bams\_indels\_realigned/JvS034.merged.rmdup.merged.realn.repma.Q30.bam.dp.hist.pdf - results/modern/mapping/GCF\_000283155.1\_CerSimSim1.0\_genomic.Sc9M7eS\_2\_HRSCAF\_41/stats/bams\_indels\_realigned/JvS035.merged.rmdup.merged.realn.repma.Q30.bam.dp.hist.pdf - results/modern/mapping/GCF\_000283155.1\_CerSimSim1.0\_genomic.Sc9M7eS\_2\_HRSCAF\_41/stats/bams\_indels\_realigned/JvS033.merged.rmdup.merged.realn.bam.stats.txt - results/modern/mapping/GCF\_000283155.1\_CerSimSim1.0\_genomic.Sc9M7eS\_2\_HRSCAF\_41/stats/bams\_indels\_realigned/JvS034.merged.rmdup.merged.realn.bam.stats.txt - results/modern/mapping/GCF\_000283155.1\_CerSimSim1.0\_genomic.Sc9M7eS\_2\_HRSCAF\_41/stats/bams\_indels\_realigned/JvS035.merged.rmdup.merged.realn.bam.stats.txt - results/modern/mapping/GCF\_000283155.1\_CerSimSim1.0\_genomic.Sc9M7eS\_2\_HRSCAF\_41/stats/bams\_indels\_realigned/JvS033.merged.rmdup.merged.realn.bam.qualimap/qualimapReport.html - results/modern/mapping/GCF\_000283155.1\_CerSimSim1.0\_genomic.Sc9M7eS\_2\_HRSCAF\_41/stats/bams\_indels\_realigned/JvS034.merged.rmdup.merged.realn.bam.qualimap/qualimapReport.html - results/modern/mapping/GCF\_000283155.1\_CerSimSim1.0\_genomic.Sc9M7eS\_2\_HRSCAF\_41/stats/bams\_indels\_realigned/JvS035.merged.rmdup.merged.realn.bam.qualimap/qualimapReport.html - results/modern/mapping/GCF\_000283155.1\_CerSimSim1.0\_genomic.Sc9M7eS\_2\_HRSCAF\_41/stats/bams\_indels\_realigned/fastqc/JvS033.merged.rmdup.merged.realn\_fastqc.html - results/modern/mapping/GCF\_000283155.1\_CerSimSim1.0\_genomic.Sc9M7eS\_2\_HRSCAF\_41/stats/bams\_indels\_realigned/fastqc/JvS034.merged.rmdup.merged.realn\_fastqc.html - results/modern/mapping/GCF\_000283155.1\_CerSimSim1.0\_genomic.Sc9M7eS\_2\_HRSCAF\_41/stats/bams\_indels\_realigned/fastqc/JvS035.merged.rmdup.merged.realn\_fastqc.html - results/modern/mapping/GCF\_000283155.1\_CerSimSim1.0\_genomic.Sc9M7eS\_2\_HRSCAF\_41/stats/bams\_indels\_realigned/fastqc/JvS033.merged.rmdup.merged.realn\_fastqc.zip - results/modern/mapping/GCF\_000283155.1\_CerSimSim1.0\_genomic.Sc9M7eS\_2\_HRSCAF\_41/stats/bams\_indels\_realigned/fastqc/JvS034.merged.rmdup.merged.realn\_fastqc.zip - results/modern/mapping/GCF\_000283155.1\_CerSimSim1.0\_genomic.Sc9M7eS\_2\_HRSCAF\_41/stats/bams\_indels\_realigned/fastqc/JvS035.merged.rmdup.merged.realn\_fastqc.zip - results/modern/mapping/GCF\_000283155.1\_CerSimSim1.0\_genomic.Sc9M7eS\_2\_HRSCAF\_41/stats/bams\_indels\_realigned/JvS033.merged.rmdup.merged.realn.repma.Q30.bam.dp.hist.pdf - results/modern/mapping/GCF\_000283155.1\_CerSimSim1.0\_genomic.Sc9M7eS\_2\_HRSCAF\_41/stats/bams\_indels\_realigned/JvS034.merged.rmdup.merged.realn.repma.Q30.bam.dp.hist.pdf - results/modern/mapping/GCF\_000283155.1\_CerSimSim1.0\_genomic.Sc9M7eS\_2\_HRSCAF\_41/stats/bams\_indels\_realigned/JvS035.merged.rmdup.merged.realn.repma.Q30.bam.dp.hist.pdf | |
| Output files | |
| - results/modern/mapping/GCF\_000283155.1\_CerSimSim1.0\_genomic.Sc9M7eS\_2\_HRSCAF\_41/stats/bams\_indels\_realigned/multiqc/multiqc\_report.html | |
| Container image | |
| docker://quay.io/biocontainers/multiqc:1.9--pyh9f0ad1d\_0 | |
| Code | |
| ``` 1 2 ``` \| ``` multiqc -f {params.indir} -o {params.outdir} 2> {log} ``` | |

###### Rule modern\_subsampled\_bam\_multiqc

×

Rule properties

|  |  |
| --- | --- |
| Jobs | 1 |
| Input files | |
| - results/modern/mapping/GCF\_000283155.1\_CerSimSim1.0\_genomic.Sc9M7eS\_2\_HRSCAF\_41/stats/bams\_subsampled/JvS034.merged.rmdup.merged.realn.mapped\_q30.subs\_dp6.bam.stats.txt - results/modern/mapping/GCF\_000283155.1\_CerSimSim1.0\_genomic.Sc9M7eS\_2\_HRSCAF\_41/stats/bams\_subsampled/JvS033.merged.rmdup.merged.realn.mapped\_q30.subs\_dp6.bam.stats.txt - results/modern/mapping/GCF\_000283155.1\_CerSimSim1.0\_genomic.Sc9M7eS\_2\_HRSCAF\_41/stats/bams\_subsampled/JvS035.merged.rmdup.merged.realn.mapped\_q30.subs\_dp6.bam.stats.txt - results/modern/mapping/GCF\_000283155.1\_CerSimSim1.0\_genomic.Sc9M7eS\_2\_HRSCAF\_41/stats/bams\_subsampled/JvS034.merged.rmdup.merged.realn.mapped\_q30.subs\_dp6.bam.qualimap/qualimapReport.html - results/modern/mapping/GCF\_000283155.1\_CerSimSim1.0\_genomic.Sc9M7eS\_2\_HRSCAF\_41/stats/bams\_subsampled/JvS033.merged.rmdup.merged.realn.mapped\_q30.subs\_dp6.bam.qualimap/qualimapReport.html - results/modern/mapping/GCF\_000283155.1\_CerSimSim1.0\_genomic.Sc9M7eS\_2\_HRSCAF\_41/stats/bams\_subsampled/JvS035.merged.rmdup.merged.realn.mapped\_q30.subs\_dp6.bam.qualimap/qualimapReport.html - results/modern/mapping/GCF\_000283155.1\_CerSimSim1.0\_genomic.Sc9M7eS\_2\_HRSCAF\_41/stats/bams\_subsampled/JvS034.merged.rmdup.merged.realn.mapped\_q30.subs\_dp6.repma.Q30.bam.dpstats.txt - results/modern/mapping/GCF\_000283155.1\_CerSimSim1.0\_genomic.Sc9M7eS\_2\_HRSCAF\_41/stats/bams\_subsampled/JvS033.merged.rmdup.merged.realn.mapped\_q30.subs\_dp6.repma.Q30.bam.dpstats.txt - results/modern/mapping/GCF\_000283155.1\_CerSimSim1.0\_genomic.Sc9M7eS\_2\_HRSCAF\_41/stats/bams\_subsampled/JvS035.merged.rmdup.merged.realn.mapped\_q30.subs\_dp6.repma.Q30.bam.dpstats.txt | |
| Output files | |
| - results/modern/mapping/GCF\_000283155.1\_CerSimSim1.0\_genomic.Sc9M7eS\_2\_HRSCAF\_41/stats/bams\_subsampled/multiqc/multiqc\_report.html | |
| Container image | |
| docker://quay.io/biocontainers/multiqc:1.9--pyh9f0ad1d\_0 | |
| Code | |
| ``` 1 2 ``` \| ``` multiqc -f {params.indir} -o {params.outdir} 2> {log} ``` | |

###### Rule subsampled\_bam\_stats

×

Rule properties

|  |  |
| --- | --- |
| Jobs | 3 |
| Input files | |
| - results/{dataset}/mapping/GCF\_000283155.1\_CerSimSim1.0\_genomic.Sc9M7eS\_2\_HRSCAF\_41/{sample}.merged.rmdup.merged.{processed}.mapped\_q30.subs\_dp{DP}.bam - results/{dataset}/mapping/GCF\_000283155.1\_CerSimSim1.0\_genomic.Sc9M7eS\_2\_HRSCAF\_41/{sample}.merged.rmdup.merged.{processed}.mapped\_q30.subs\_dp{DP}.bam.bai | |
| Output files | |
| - results/{dataset}/mapping/GCF\_000283155.1\_CerSimSim1.0\_genomic.Sc9M7eS\_2\_HRSCAF\_41/stats/bams\_subsampled/{sample,[A-Za-z0-9]+}.merged.rmdup.merged.{processed}.mapped\_q30.subs\_dp{DP,[+-]?(\d+(\.\d\*)?\|\.\d+)([eE][+-]?\d+)?}.bam.stats.txt | |
| Container image | |
| docker://biocontainers/samtools:v1.9-4-deb\_cv1 | |
| Code | |
| ``` 1 2 ``` \| ``` samtools flagstat {input.bam} > {output.stats} 2> {log} ``` | |

###### Rule subsample\_bams

×

Rule properties

|  |  |
| --- | --- |
| Jobs | 3 |
| Input files | |
| - results/{dataset}/mapping/GCF\_000283155.1\_CerSimSim1.0\_genomic.Sc9M7eS\_2\_HRSCAF\_41/{sample}.merged.rmdup.merged.{processed}.mapped\_q30.bam - results/{dataset}/mapping/GCF\_000283155.1\_CerSimSim1.0\_genomic.Sc9M7eS\_2\_HRSCAF\_41/stats/bams\_indels\_realigned/{sample}.merged.rmdup.merged.realn.repma.Q30.bam.dpstats.txt | |
| Output files | |
| - results/{dataset}/mapping/GCF\_000283155.1\_CerSimSim1.0\_genomic.Sc9M7eS\_2\_HRSCAF\_41/{sample,[A-Za-z0-9]+}.merged.rmdup.merged.{processed}.mapped\_q30.subs\_dp{DP,[+-]?(\d+(\.\d\*)?\|\.\d+)([eE][+-]?\d+)?}.bam | |
| Container image | |
| docker://biocontainers/samtools:v1.9-4-deb\_cv1 | |
| Code | |
| ``` 1 2 3 4 5 6 7 8 9 10 11 ``` \| ``` depth=`head -n 1 {input.dp} \| cut -d' ' -f 1` frac=`awk -v s={params.DP} -v d=$depth "BEGIN {{print s/d}}"` if [ `awk 'BEGIN {{print ('$frac' <= 1.0)}}'` = 1 ] # awk will return 1 if the statement is true, and 0 if it is false then samtools view -h -b -s $frac -@ {threads} -o {output.subsam} {input.bam} 2> {log} else echo "!!! Warning [genome erosion workflow]: The sample {input.bam} has a lower average depth than the target depth for subsampling. Remove the sample from the subsampling list in the config file or choose a lower target depth. !!!" >> {log} fi ``` | |

###### Rule filter\_bam\_mapped\_mq

×

Rule properties

|  |  |
| --- | --- |
| Jobs | 3 |
| Input files | |
| - results/{dataset}/mapping/GCF\_000283155.1\_CerSimSim1.0\_genomic.Sc9M7eS\_2\_HRSCAF\_41/{sample}.merged.rmdup.merged.{processed}.bam | |
| Output files | |
| - results/{dataset}/mapping/GCF\_000283155.1\_CerSimSim1.0\_genomic.Sc9M7eS\_2\_HRSCAF\_41/{sample,[A-Za-z0-9]+}.merged.rmdup.merged.{processed}.mapped\_q30.bam | |
| Container image | |
| docker://biocontainers/samtools:v1.9-4-deb\_cv1 | |
| Code | |
| ``` 1 2 ``` \| ``` samtools view -h -b -F 4 -q 30 -@ {threads} -o {output.filtered} {input.bam} 2> {log} ``` | |

###### Rule index\_subsampled\_bams

×

Rule properties

|  |  |
| --- | --- |
| Jobs | 3 |
| Input files | |
| - results/{dataset}/mapping/GCF\_000283155.1\_CerSimSim1.0\_genomic.Sc9M7eS\_2\_HRSCAF\_41/{sample}.merged.rmdup.merged.{processed}.mapped\_q30.subs\_dp{DP}.bam | |
| Output files | |
| - results/{dataset}/mapping/GCF\_000283155.1\_CerSimSim1.0\_genomic.Sc9M7eS\_2\_HRSCAF\_41/{sample,[A-Za-z0-9]+}.merged.rmdup.merged.{processed}.mapped\_q30.subs\_dp{DP,[+-]?(\d+(\.\d\*)?\|\.\d+)([eE][+-]?\d+)?}.bam.bai | |
| Container image | |
| docker://biocontainers/samtools:v1.9-4-deb\_cv1 | |
| Code | |
| ``` 1 2 ``` \| ``` samtools index {input.bam} {output.index} 2> {log} ``` | |

###### Rule subsampled\_bam\_qualimap

×

Rule properties

|  |  |
| --- | --- |
| Jobs | 3 |
| Input files | |
| - results/{dataset}/mapping/GCF\_000283155.1\_CerSimSim1.0\_genomic.Sc9M7eS\_2\_HRSCAF\_41/{sample}.merged.rmdup.merged.{processed}.mapped\_q30.subs\_dp{DP}.bam - results/{dataset}/mapping/GCF\_000283155.1\_CerSimSim1.0\_genomic.Sc9M7eS\_2\_HRSCAF\_41/{sample}.merged.rmdup.merged.{processed}.mapped\_q30.subs\_dp{DP}.bam.bai | |
| Output files | |
| - results/{dataset}/mapping/GCF\_000283155.1\_CerSimSim1.0\_genomic.Sc9M7eS\_2\_HRSCAF\_41/stats/bams\_subsampled/{sample,[A-Za-z0-9]+}.merged.rmdup.merged.{processed}.mapped\_q30.subs\_dp{DP,[+-]?(\d+(\.\d\*)?\|\.\d+)([eE][+-]?\d+)?}.bam.qualimap/qualimapReport.html - results/{dataset}/mapping/GCF\_000283155.1\_CerSimSim1.0\_genomic.Sc9M7eS\_2\_HRSCAF\_41/stats/bams\_subsampled/{sample,[A-Za-z0-9]+}.merged.rmdup.merged.{processed}.mapped\_q30.subs\_dp{DP,[+-]?(\d+(\.\d\*)?\|\.\d+)([eE][+-]?\d+)?}.bam.qualimap/genome\_results.txt - results/{dataset}/mapping/GCF\_000283155.1\_CerSimSim1.0\_genomic.Sc9M7eS\_2\_HRSCAF\_41/stats/bams\_subsampled/{sample,[A-Za-z0-9]+}.merged.rmdup.merged.{processed}.mapped\_q30.subs\_dp{DP,[+-]?(\d+(\.\d\*)?\|\.\d+)([eE][+-]?\d+)?}.bam.qualimap | |
| Container image | |
| docker://quay.io/biocontainers/qualimap:2.2.2d--1 | |
| Code | |
| ``` 1 2 3 4 ``` \| ``` mem=$(((6 * {threads}) - 2)) unset DISPLAY qualimap bamqc -bam {input.bam} --java-mem-size=${{mem}}G -nt {threads} -outdir {params.outdir} -outformat html 2> {log} ``` | |

###### Rule subsampled\_bam\_depth

×

Rule properties

|  |  |
| --- | --- |
| Jobs | 3 |
| Input files | |
| - results/{dataset}/mapping/GCF\_000283155.1\_CerSimSim1.0\_genomic.Sc9M7eS\_2\_HRSCAF\_41/{sample}.merged.rmdup.merged.{processed}.mapped\_q30.subs\_dp{DP}.bam - /proj/sllstore2017093/b2016342/b2016342\_nobackup/lts/genome\_erosion\_pipeline/verena\_testing/testdata/reference/GCF\_000283155.1\_CerSimSim1.0\_genomic.Sc9M7eS\_2\_HRSCAF\_41.repma.bed | |
| Output files | |
| - results/{dataset}/mapping/GCF\_000283155.1\_CerSimSim1.0\_genomic.Sc9M7eS\_2\_HRSCAF\_41/stats/bams\_subsampled/{sample,[A-Za-z0-9]+}.merged.rmdup.merged.{processed}.mapped\_q30.subs\_dp{DP,[+-]?(\d+(\.\d\*)?\|\.\d+)([eE][+-]?\d+)?}.repma.Q30.bam.dp - results/{dataset}/mapping/GCF\_000283155.1\_CerSimSim1.0\_genomic.Sc9M7eS\_2\_HRSCAF\_41/stats/bams\_subsampled/{sample,[A-Za-z0-9]+}.merged.rmdup.merged.{processed}.mapped\_q30.subs\_dp{DP,[+-]?(\d+(\.\d\*)?\|\.\d+)([eE][+-]?\d+)?}.repma.Q30.bam.dpstats.txt | |
| Container image | |
| docker://biocontainers/samtools:v1.9-4-deb\_cv1 | |
| Code | |
| ``` 1 2 3 4 5 6 7 8 9 10 ``` \| ``` if [ {params.cov} = "True" ] # include sites with missing data / zero coverage then samtools depth -a -Q 30 -q 30 -b {input.no_rep_bed} {input.bam} > {output.tmp} 2> {log} && awk '{{sum+=$3}} END {{ print sum/NR }}' {output.tmp} \| awk -v min={params.minDP} -v max={params.maxDP} '{{ printf "%.0f %.0f %.0f", $1, $1*min, $1*max }}' > {output.dp} 2>> {log} elif [ {params.cov} = "False" ] # exclude sites with missing data / zero coverage then samtools depth -Q 30 -q 30 -b {input.no_rep_bed} {input.bam} > {output.tmp} 2> {log} && awk '{{sum+=$3}} END {{ print sum/NR }}' {output.tmp} \| awk -v min={params.minDP} -v max={params.maxDP} '{{ printf "%.0f %.0f %.0f", $1, $1*min, $1*max }}' > {output.dp} 2>> {log} fi ``` | |

###### Rule historical\_sorted\_vcf\_multiqc

×

Rule properties

|  |  |
| --- | --- |
| Jobs | 1 |
| Input files | |
| - results/historical/vcf/GCF\_000283155.1\_CerSimSim1.0\_genomic.Sc9M7eS\_2\_HRSCAF\_41/stats/vcf\_sorted/JvS009.merged.rmdup.merged.realn.Q30.sorted.vcf.stats.txt - results/historical/vcf/GCF\_000283155.1\_CerSimSim1.0\_genomic.Sc9M7eS\_2\_HRSCAF\_41/stats/vcf\_sorted/JvS022.merged.rmdup.merged.realn.Q30.sorted.vcf.stats.txt - results/historical/vcf/GCF\_000283155.1\_CerSimSim1.0\_genomic.Sc9M7eS\_2\_HRSCAF\_41/stats/vcf\_sorted/JvS008.merged.rmdup.merged.realn.Q30.sorted.vcf.stats.txt | |
| Output files | |
| - results/historical/vcf/GCF\_000283155.1\_CerSimSim1.0\_genomic.Sc9M7eS\_2\_HRSCAF\_41/stats/vcf\_sorted/multiqc/multiqc\_report.html | |
| Container image | |
| docker://quay.io/biocontainers/multiqc:1.9--pyh9f0ad1d\_0 | |
| Code | |
| ``` 1 2 ``` \| ``` multiqc -f {params.indir} -o {params.outdir} 2> {log} ``` | |

###### Rule sorted\_vcf\_stats

×

Rule properties

|  |  |
| --- | --- |
| Jobs | 6 |
| Input files | |
| - results/{dataset}/vcf/GCF\_000283155.1\_CerSimSim1.0\_genomic.Sc9M7eS\_2\_HRSCAF\_41/{sample}.merged.rmdup.merged.{processed}.Q30.sorted.bcf - results/{dataset}/vcf/GCF\_000283155.1\_CerSimSim1.0\_genomic.Sc9M7eS\_2\_HRSCAF\_41/{sample}.merged.rmdup.merged.{processed}.Q30.sorted.bcf.csi | |
| Output files | |
| - results/{dataset}/vcf/GCF\_000283155.1\_CerSimSim1.0\_genomic.Sc9M7eS\_2\_HRSCAF\_41/stats/vcf\_sorted/{sample,[A-Za-z0-9]+}.merged.rmdup.merged.{processed}.Q30.sorted.vcf.stats.txt | |
| Container image | |
| docker://quay.io/biocontainers/bcftools:1.9--h68d8f2e\_9 | |
| Code | |
| ``` 1 2 ``` \| ``` bcftools stats {input.sort} > {output.stats} 2> {log} ``` | |

###### Rule sort\_vcfs

×

Rule properties

|  |  |
| --- | --- |
| Jobs | 6 |
| Input files | |
| - results/{dataset}/vcf/GCF\_000283155.1\_CerSimSim1.0\_genomic.Sc9M7eS\_2\_HRSCAF\_41/{sample}.merged.rmdup.merged.{processed}.Q30.bcf | |
| Output files | |
| - results/{dataset}/vcf/GCF\_000283155.1\_CerSimSim1.0\_genomic.Sc9M7eS\_2\_HRSCAF\_41/{sample,[A-Za-z0-9]+}.merged.rmdup.merged.{processed}.Q30.sorted.bcf | |
| Container image | |
| docker://quay.io/biocontainers/bcftools:1.9--h68d8f2e\_9 | |
| Code | |
| ``` 1 2 ``` \| ``` bcftools sort -O b -T {params.tmpdir} -o {output.sort} {input.bcf} 2> {log} ``` | |

###### Rule variant\_calling

×

Rule properties

|  |  |
| --- | --- |
| Jobs | 6 |
| Input files | |
| - /proj/sllstore2017093/b2016342/b2016342\_nobackup/lts/genome\_erosion\_pipeline/verena\_testing/testdata/reference/GCF\_000283155.1\_CerSimSim1.0\_genomic.Sc9M7eS\_2\_HRSCAF\_41.fasta - results/{dataset}/mapping/GCF\_000283155.1\_CerSimSim1.0\_genomic.Sc9M7eS\_2\_HRSCAF\_41/{sample}.merged.rmdup.merged.{processed}.bam | |
| Output files | |
| - results/{dataset}/vcf/GCF\_000283155.1\_CerSimSim1.0\_genomic.Sc9M7eS\_2\_HRSCAF\_41/{sample,[A-Za-z0-9]+}.merged.rmdup.merged.{processed}.Q30.bcf | |
| Container image | |
| docker://quay.io/biocontainers/bcftools:1.9--h68d8f2e\_9 | |
| Code | |
| ``` 1 2 ``` \| ``` bcftools mpileup -Ou -Q 30 -q 30 -B -f {input.ref} {input.bam} \| bcftools call -c -M -O b --threads {threads} -o {output.bcf} 2> {log} ``` | |

###### Rule index\_sorted\_vcfs

×

Rule properties

|  |  |
| --- | --- |
| Jobs | 6 |
| Input files | |
| - results/{dataset}/vcf/GCF\_000283155.1\_CerSimSim1.0\_genomic.Sc9M7eS\_2\_HRSCAF\_41/{sample}.merged.rmdup.merged.{processed}.Q30.sorted.bcf | |
| Output files | |
| - results/{dataset}/vcf/GCF\_000283155.1\_CerSimSim1.0\_genomic.Sc9M7eS\_2\_HRSCAF\_41/{sample,[A-Za-z0-9]+}.merged.rmdup.merged.{processed}.Q30.sorted.bcf.csi | |
| Container image | |
| docker://quay.io/biocontainers/bcftools:1.9--h68d8f2e\_9 | |
| Code | |
| ``` 1 2 ``` \| ``` bcftools index -o {output.index} {input.sort} 2> {log} ``` | |

###### Rule modern\_sorted\_vcf\_multiqc

×

Rule properties

|  |  |
| --- | --- |
| Jobs | 1 |
| Input files | |
| - results/modern/vcf/GCF\_000283155.1\_CerSimSim1.0\_genomic.Sc9M7eS\_2\_HRSCAF\_41/stats/vcf\_sorted/JvS034.merged.rmdup.merged.realn.mapped\_q30.subs\_dp6.Q30.sorted.vcf.stats.txt - results/modern/vcf/GCF\_000283155.1\_CerSimSim1.0\_genomic.Sc9M7eS\_2\_HRSCAF\_41/stats/vcf\_sorted/JvS033.merged.rmdup.merged.realn.mapped\_q30.subs\_dp6.Q30.sorted.vcf.stats.txt - results/modern/vcf/GCF\_000283155.1\_CerSimSim1.0\_genomic.Sc9M7eS\_2\_HRSCAF\_41/stats/vcf\_sorted/JvS035.merged.rmdup.merged.realn.mapped\_q30.subs\_dp6.Q30.sorted.vcf.stats.txt | |
| Output files | |
| - results/modern/vcf/GCF\_000283155.1\_CerSimSim1.0\_genomic.Sc9M7eS\_2\_HRSCAF\_41/stats/vcf\_sorted/multiqc/multiqc\_report.html | |
| Container image | |
| docker://quay.io/biocontainers/multiqc:1.9--pyh9f0ad1d\_0 | |
| Code | |
| ``` 1 2 ``` \| ``` multiqc -f {params.indir} -o {params.outdir} 2> {log} ``` | |

###### Rule make\_CpG\_reference\_bed

×

Rule properties

|  |  |
| --- | --- |
| Jobs | 1 |
| Input files | |
| - /proj/sllstore2017093/b2016342/b2016342\_nobackup/lts/genome\_erosion\_pipeline/verena\_testing/testdata/reference/GCF\_000283155.1\_CerSimSim1.0\_genomic.Sc9M7eS\_2\_HRSCAF\_41.upper.fasta | |
| Output files | |
| - results/GCF\_000283155.1\_CerSimSim1.0\_genomic.Sc9M7eS\_2\_HRSCAF\_41.CpG\_ref.bed | |
| Code | |
| ``` 1 2 3 ``` \| ``` python workflow/scripts/find_CpG_ref_sites.py {input.ref} {params.refdirbed} 2> {log} && cp {params.refdirbed}.bed {output.bed} 2>> {log} ``` | |

###### Rule make\_noCpG\_bed

×

Rule properties

|  |  |
| --- | --- |
| Jobs | 1 |
| Input files | |
| - /proj/sllstore2017093/b2016342/b2016342\_nobackup/lts/genome\_erosion\_pipeline/verena\_testing/testdata/reference/GCF\_000283155.1\_CerSimSim1.0\_genomic.Sc9M7eS\_2\_HRSCAF\_41.bed - results/GCF\_000283155.1\_CerSimSim1.0\_genomic.Sc9M7eS\_2\_HRSCAF\_41.{CpG\_method}.bed | |
| Output files | |
| - results/GCF\_000283155.1\_CerSimSim1.0\_genomic.Sc9M7eS\_2\_HRSCAF\_41.no{CpG\_method,CpG\_[vcfre]{3,6}}.bed | |
| Container image | |
| docker://quay.io/biocontainers/bedtools:2.29.2--hc088bd4\_0 | |
| Code | |
| ``` 1 2 ``` \| ``` bedtools subtract -a {input.ref_bed} -b {input.CpG_bed} > {output.no_CpG_bed} 2> {log} ``` | |

###### Rule make\_noCpG\_repma\_bed

×

Rule properties

|  |  |
| --- | --- |
| Jobs | 1 |
| Input files | |
| - /proj/sllstore2017093/b2016342/b2016342\_nobackup/lts/genome\_erosion\_pipeline/verena\_testing/testdata/reference/GCF\_000283155.1\_CerSimSim1.0\_genomic.Sc9M7eS\_2\_HRSCAF\_41.bed - results/GCF\_000283155.1\_CerSimSim1.0\_genomic.Sc9M7eS\_2\_HRSCAF\_41.merged.{CpG\_method}.repeats.bed | |
| Output files | |
| - results/GCF\_000283155.1\_CerSimSim1.0\_genomic.Sc9M7eS\_2\_HRSCAF\_41.no{CpG\_method,CpG\_[vcfre]{3,6}}.repma.bed | |
| Container image | |
| docker://quay.io/biocontainers/bedtools:2.29.2--hc088bd4\_0 | |
| Code | |
| ``` 1 2 ``` \| ``` bedtools subtract -a {input.ref_bed} -b {input.merged_bed} > {output.no_CpG_repma_bed} 2> {log} ``` | |

###### Rule merge\_CpG\_repeats\_beds

×

Rule properties

|  |  |
| --- | --- |
| Jobs | 1 |
| Input files | |
| - results/GCF\_000283155.1\_CerSimSim1.0\_genomic.Sc9M7eS\_2\_HRSCAF\_41.{CpG\_method}.bed - /proj/sllstore2017093/b2016342/b2016342\_nobackup/lts/genome\_erosion\_pipeline/verena\_testing/testdata/reference/GCF\_000283155.1\_CerSimSim1.0\_genomic.Sc9M7eS\_2\_HRSCAF\_41.repeats.sorted.bed | |
| Output files | |
| - results/GCF\_000283155.1\_CerSimSim1.0\_genomic.Sc9M7eS\_2\_HRSCAF\_41.concatenated.{CpG\_method,CpG\_[vcfre]{3,6}}.repeats.bed - results/GCF\_000283155.1\_CerSimSim1.0\_genomic.Sc9M7eS\_2\_HRSCAF\_41.merged.{CpG\_method,CpG\_[vcfre]{3,6}}.repeats.bed | |
| Container image | |
| docker://quay.io/biocontainers/bedtools:2.29.2--hc088bd4\_0 | |
| Code | |
| ``` 1 2 3 ``` \| ``` cat {input[0]} {input[1]} \| sort -k1,1 -k2,2n > {output.tmp} 2> {log} && bedtools merge -i {output.tmp} > {output.merged} 2>> {log} ``` | |

###### Rule sort\_CpG\_repeats\_beds

×

Rule properties

|  |  |
| --- | --- |
| Jobs | 1 |
| Input files | |
| - results/GCF\_000283155.1\_CerSimSim1.0\_genomic.Sc9M7eS\_2\_HRSCAF\_41.merged.{CpG\_method}.repeats.bed - /proj/sllstore2017093/b2016342/b2016342\_nobackup/lts/genome\_erosion\_pipeline/verena\_testing/testdata/reference/GCF\_000283155.1\_CerSimSim1.0\_genomic.Sc9M7eS\_2\_HRSCAF\_41.genome | |
| Output files | |
| - results/GCF\_000283155.1\_CerSimSim1.0\_genomic.Sc9M7eS\_2\_HRSCAF\_41.{CpG\_method,CpG\_[vcfre]{3,6}}.repeats.bed | |
| Container image | |
| docker://quay.io/biocontainers/bedtools:2.29.2--hc088bd4\_0 | |
| Code | |
| ``` 1 2 ``` \| ``` bedtools sort -g {input.genomefile} -i {input.merged_bed} > {output.sorted_bed} 2> {log} ``` | |

###### Rule historical\_CpG\_filtered\_vcf\_multiqc

×

Rule properties

|  |  |
| --- | --- |
| Jobs | 1 |
| Input files | |
| Output files | |
| - results/historical/vcf/GCF\_000283155.1\_CerSimSim1.0\_genomic.Sc9M7eS\_2\_HRSCAF\_41/stats/vcf\_CpG\_filtered/multiqc/multiqc\_report.html | |
| Container image | |
| docker://quay.io/biocontainers/multiqc:1.9--pyh9f0ad1d\_0 | |
| Code | |
| ``` 1 2 ``` \| ``` multiqc -f {params.indir} -o {params.outdir} 2> {log} ``` | |

###### Rule CpG\_filtered\_vcf\_stats

×

Rule properties

|  |  |
| --- | --- |
| Jobs | 6 |
| Input files | |
| - results/{dataset}/vcf/GCF\_000283155.1\_CerSimSim1.0\_genomic.Sc9M7eS\_2\_HRSCAF\_41/{sample}.merged.rmdup.merged.{processed}.Q30.sorted.no{CpG\_method}.bcf - results/{dataset}/vcf/GCF\_000283155.1\_CerSimSim1.0\_genomic.Sc9M7eS\_2\_HRSCAF\_41/{sample}.merged.rmdup.merged.{processed}.Q30.sorted.no{CpG\_method}.bcf.csi | |
| Output files | |
| - results/{dataset}/vcf/GCF\_000283155.1\_CerSimSim1.0\_genomic.Sc9M7eS\_2\_HRSCAF\_41/stats/vcf\_CpG\_filtered/{sample,[A-Za-z0-9]+}.merged.rmdup.merged.{processed}.Q30.sorted.no{CpG\_method,CpG\_[vcfre]{3,6}}.bcf.stats.txt | |
| Container image | |
| docker://quay.io/biocontainers/bcftools:1.9--h68d8f2e\_9 | |
| Code | |
| ``` 1 2 ``` \| ``` bcftools stats {input.bcf} > {output.stats} 2> {log} ``` | |

###### Rule CpG\_vcf2bcf

×

Rule properties

|  |  |
| --- | --- |
| Jobs | 6 |
| Input files | |
| - results/{dataset}/vcf/GCF\_000283155.1\_CerSimSim1.0\_genomic.Sc9M7eS\_2\_HRSCAF\_41/{sample}.merged.rmdup.merged.{processed}.Q30.sorted.no{CpG\_method}.vcf | |
| Output files | |
| - results/{dataset}/vcf/GCF\_000283155.1\_CerSimSim1.0\_genomic.Sc9M7eS\_2\_HRSCAF\_41/{sample,[A-Za-z0-9]+}.merged.rmdup.merged.{processed}.Q30.sorted.no{CpG\_method,CpG\_[vcfre]{3,6}}.bcf | |
| Container image | |
| docker://quay.io/biocontainers/bcftools:1.9--h68d8f2e\_9 | |
| Code | |
| ``` 1 2 ``` \| ``` bcftools convert -O b -o {output.bcf} {input.filtered} 2> {log} ``` | |

###### Rule remove\_CpG\_vcf

×

Rule properties

|  |  |
| --- | --- |
| Jobs | 6 |
| Input files | |
| - results/{dataset}/vcf/GCF\_000283155.1\_CerSimSim1.0\_genomic.Sc9M7eS\_2\_HRSCAF\_41/{sample}.merged.rmdup.merged.{processed}.Q30.sorted.CpG\_rm.vcf.gz - results/GCF\_000283155.1\_CerSimSim1.0\_genomic.Sc9M7eS\_2\_HRSCAF\_41.no{CpG\_method}.bed - /proj/sllstore2017093/b2016342/b2016342\_nobackup/lts/genome\_erosion\_pipeline/verena\_testing/testdata/reference/GCF\_000283155.1\_CerSimSim1.0\_genomic.Sc9M7eS\_2\_HRSCAF\_41.genome | |
| Output files | |
| - results/{dataset}/vcf/GCF\_000283155.1\_CerSimSim1.0\_genomic.Sc9M7eS\_2\_HRSCAF\_41/{sample,[A-Za-z0-9]+}.merged.rmdup.merged.{processed}.Q30.sorted.no{CpG\_method,CpG\_[vcfre]{3,6}}.vcf | |
| Container image | |
| docker://quay.io/biocontainers/bedtools:2.29.2--hc088bd4\_0 | |
| Code | |
| ``` 1 2 ``` \| ``` bedtools intersect -a {input.vcf} -b {input.bed} -header -sorted -g {input.genomefile} > {output.filtered} 2> {log} ``` | |

###### Rule sorted\_bcf2vcf\_CpG\_removal

×

Rule properties

|  |  |
| --- | --- |
| Jobs | 6 |
| Input files | |
| - results/{dataset}/vcf/GCF\_000283155.1\_CerSimSim1.0\_genomic.Sc9M7eS\_2\_HRSCAF\_41/{sample}.merged.rmdup.merged.{processed}.Q30.sorted.bcf | |
| Output files | |
| - results/{dataset}/vcf/GCF\_000283155.1\_CerSimSim1.0\_genomic.Sc9M7eS\_2\_HRSCAF\_41/{sample,[A-Za-z0-9]+}.merged.rmdup.merged.{processed}.Q30.sorted.CpG\_rm.vcf.gz | |
| Container image | |
| docker://quay.io/biocontainers/bcftools:1.9--h68d8f2e\_9 | |
| Code | |
| ``` 1 2 ``` \| ``` bcftools convert -O z -o {output.vcf} {input.bcf} 2> {log} ``` | |

###### Rule index\_CpG\_bcf

×

Rule properties

|  |  |
| --- | --- |
| Jobs | 6 |
| Input files | |
| - results/{dataset}/vcf/GCF\_000283155.1\_CerSimSim1.0\_genomic.Sc9M7eS\_2\_HRSCAF\_41/{sample}.merged.rmdup.merged.{processed}.Q30.sorted.no{CpG\_method}.bcf | |
| Output files | |
| - results/{dataset}/vcf/GCF\_000283155.1\_CerSimSim1.0\_genomic.Sc9M7eS\_2\_HRSCAF\_41/{sample,[A-Za-z0-9]+}.merged.rmdup.merged.{processed}.Q30.sorted.no{CpG\_method,CpG\_[vcfre]{3,6}}.bcf.csi | |
| Container image | |
| docker://quay.io/biocontainers/bcftools:1.9--h68d8f2e\_9 | |
| Code | |
| ``` 1 2 ``` \| ``` bcftools index -o {output.index} {input.bcf} 2> {log} ``` | |

###### Rule modern\_CpG\_filtered\_vcf\_multiqc

×

Rule properties

|  |  |
| --- | --- |
| Jobs | 1 |
| Input files | |
| Output files | |
| - results/modern/vcf/GCF\_000283155.1\_CerSimSim1.0\_genomic.Sc9M7eS\_2\_HRSCAF\_41/stats/vcf\_CpG\_filtered/multiqc/multiqc\_report.html | |
| Container image | |
| docker://quay.io/biocontainers/multiqc:1.9--pyh9f0ad1d\_0 | |
| Code | |
| ``` 1 2 ``` \| ``` multiqc -f {params.indir} -o {params.outdir} 2> {log} ``` | |

###### Rule historical\_quality\_filtered\_vcf\_multiqc

×

Rule properties

|  |  |
| --- | --- |
| Jobs | 1 |
| Input files | |
| Output files | |
| - results/historical/vcf/GCF\_000283155.1\_CerSimSim1.0\_genomic.Sc9M7eS\_2\_HRSCAF\_41/stats/vcf\_qual\_filtered/multiqc/multiqc\_report.html | |
| Container image | |
| docker://quay.io/biocontainers/multiqc:1.9--pyh9f0ad1d\_0 | |
| Code | |
| ``` 1 2 ``` \| ``` multiqc -f {params.indir} -o {params.outdir} 2> {log} ``` | |

###### Rule filtered\_vcf\_stats

×

Rule properties

|  |  |
| --- | --- |
| Jobs | 6 |
| Input files | |
| - results/{dataset}/vcf/GCF\_000283155.1\_CerSimSim1.0\_genomic.Sc9M7eS\_2\_HRSCAF\_41/{sample}.merged.rmdup.merged.{processed}.snps5.noIndel.QUAL30.dp.AB.bcf - results/{dataset}/vcf/GCF\_000283155.1\_CerSimSim1.0\_genomic.Sc9M7eS\_2\_HRSCAF\_41/{sample}.merged.rmdup.merged.{processed}.snps5.noIndel.QUAL30.dp.AB.bcf.csi | |
| Output files | |
| - results/{dataset}/vcf/GCF\_000283155.1\_CerSimSim1.0\_genomic.Sc9M7eS\_2\_HRSCAF\_41/stats/vcf\_qual\_filtered/{sample,[A-Za-z0-9]+}.merged.rmdup.merged.{processed}.snps5.noIndel.QUAL30.dp.AB.bcf.stats.txt | |
| Container image | |
| docker://quay.io/biocontainers/bcftools:1.9--h68d8f2e\_9 | |
| Code | |
| ``` 1 2 ``` \| ``` bcftools stats {input.bcf} > {output.stats} 2> {log} ``` | |

###### Rule filter\_vcfs\_allelic\_balance

×

Rule properties

|  |  |
| --- | --- |
| Jobs | 6 |
| Input files | |
| - results/{dataset}/vcf/GCF\_000283155.1\_CerSimSim1.0\_genomic.Sc9M7eS\_2\_HRSCAF\_41/{sample}.merged.rmdup.merged.{processed}.snps5.noIndel.QUAL30.dp.bcf | |
| Output files | |
| - results/{dataset}/vcf/GCF\_000283155.1\_CerSimSim1.0\_genomic.Sc9M7eS\_2\_HRSCAF\_41/{sample,[A-Za-z0-9]+}.merged.rmdup.merged.{processed}.snps5.noIndel.QUAL30.dp.AB.bcf | |
| Container image | |
| docker://quay.io/biocontainers/bcftools:1.9--h68d8f2e\_9 | |
| Code | |
| ``` 1 2 ``` \| ``` bcftools view -e 'GT="0/1" & (DP4[2]+DP4[3])/(DP4[0]+DP4[1]+DP4[2]+DP4[3]) < 0.2' {input.bcf} \| bcftools view -e 'GT="0/1" & (DP4[2]+DP4[3])/(DP4[0]+DP4[1]+DP4[2]+DP4[3]) > 0.8' -Ob > {output.filtered} 2> {log} ``` | |

###### Rule filter\_vcfs\_qual\_dp

×

Rule properties

|  |  |
| --- | --- |
| Jobs | 6 |
| Input files | |
| - results/{dataset}/vcf/GCF\_000283155.1\_CerSimSim1.0\_genomic.Sc9M7eS\_2\_HRSCAF\_41/{sample}.merged.rmdup.merged.{processed}.snps5.bcf | |
| Output files | |
| - results/{dataset}/vcf/GCF\_000283155.1\_CerSimSim1.0\_genomic.Sc9M7eS\_2\_HRSCAF\_41/{sample,[A-Za-z0-9]+}.merged.rmdup.merged.{processed}.snps5.noIndel.QUAL30.dp.bcf | |
| Container image | |
| docker://quay.io/biocontainers/bcftools:1.9--h68d8f2e\_9 | |
| Code | |
| ``` 1 2 3 4 5 6 7 8 9 10 11 ``` \| ``` minDP=`head -n 1 {input.dp} \| cut -d' ' -f 2` maxDP=`head -n 1 {input.dp} \| cut -d' ' -f 3` # check minimum depth threshold if awk "BEGIN{{exit ! ($minDP < 3)}}" then minDP=3 fi bcftools filter -i "(DP4[0]+DP4[1]+DP4[2]+DP4[3])>$minDP & (DP4[0]+DP4[1]+DP4[2]+DP4[3])<$maxDP & QUAL>=30 & INDEL=0" -O b --threads {threads} -o {output.filtered} {input.bcf} 2> {log} ``` | |

###### Rule remove\_snps\_near\_indels

×

Rule properties

|  |  |
| --- | --- |
| Jobs | 6 |
| Input files | |
| - results/{dataset}/vcf/GCF\_000283155.1\_CerSimSim1.0\_genomic.Sc9M7eS\_2\_HRSCAF\_41/{sample}.merged.rmdup.merged.{processed}.bcf | |
| Output files | |
| - results/{dataset}/vcf/GCF\_000283155.1\_CerSimSim1.0\_genomic.Sc9M7eS\_2\_HRSCAF\_41/{sample,[A-Za-z0-9]+}.merged.rmdup.merged.{processed}.snps5.bcf | |
| Container image | |
| docker://quay.io/biocontainers/bcftools:1.9--h68d8f2e\_9 | |
| Code | |
| ``` 1 2 ``` \| ``` bcftools filter -g 5 -O b --threads {threads} -o {output.snps} {input.bcf} 2> {log} ``` | |

###### Rule index\_filtered\_vcfs

×

Rule properties

|  |  |
| --- | --- |
| Jobs | 6 |
| Input files | |
| - results/{dataset}/vcf/GCF\_000283155.1\_CerSimSim1.0\_genomic.Sc9M7eS\_2\_HRSCAF\_41/{sample}.merged.rmdup.merged.{processed}.snps5.noIndel.QUAL30.dp.AB.bcf | |
| Output files | |
| - results/{dataset}/vcf/GCF\_000283155.1\_CerSimSim1.0\_genomic.Sc9M7eS\_2\_HRSCAF\_41/{sample,[A-Za-z0-9]+}.merged.rmdup.merged.{processed}.snps5.noIndel.QUAL30.dp.AB.bcf.csi | |
| Container image | |
| docker://quay.io/biocontainers/bcftools:1.9--h68d8f2e\_9 | |
| Code | |
| ``` 1 2 ``` \| ``` bcftools index -o {output.index} {input.bcf} 2> {log} ``` | |

###### Rule historical\_repmasked\_vcf\_multiqc

×

Rule properties

|  |  |
| --- | --- |
| Jobs | 1 |
| Input files | |
| Output files | |
| - results/historical/vcf/GCF\_000283155.1\_CerSimSim1.0\_genomic.Sc9M7eS\_2\_HRSCAF\_41/stats/vcf\_repmasked/multiqc/multiqc\_report.html | |
| Container image | |
| docker://quay.io/biocontainers/multiqc:1.9--pyh9f0ad1d\_0 | |
| Code | |
| ``` 1 2 ``` \| ``` multiqc -f {params.indir} -o {params.outdir} 2> {log} ``` | |

###### Rule repmasked\_vcf\_stats

×

Rule properties

|  |  |
| --- | --- |
| Jobs | 6 |
| Input files | |
| - results/{dataset}/vcf/GCF\_000283155.1\_CerSimSim1.0\_genomic.Sc9M7eS\_2\_HRSCAF\_41/{sample}.merged.rmdup.merged.{processed}.snps5.noIndel.QUAL30.dp.AB.repma.bcf - results/{dataset}/vcf/GCF\_000283155.1\_CerSimSim1.0\_genomic.Sc9M7eS\_2\_HRSCAF\_41/{sample}.merged.rmdup.merged.{processed}.snps5.noIndel.QUAL30.dp.AB.repma.bcf.csi | |
| Output files | |
| - results/{dataset}/vcf/GCF\_000283155.1\_CerSimSim1.0\_genomic.Sc9M7eS\_2\_HRSCAF\_41/stats/vcf\_repmasked/{sample,[A-Za-z0-9]+}.merged.rmdup.merged.{processed}.snps5.noIndel.QUAL30.dp.AB.repma.bcf.stats.txt | |
| Container image | |
| docker://quay.io/biocontainers/bcftools:1.9--h68d8f2e\_9 | |
| Code | |
| ``` 1 2 ``` \| ``` bcftools stats {input.bcf} > {output.stats} 2> {log} ``` | |

###### Rule filtered\_vcf2bcf

×

Rule properties

|  |  |
| --- | --- |
| Jobs | 6 |
| Input files | |
| - results/{dataset}/vcf/GCF\_000283155.1\_CerSimSim1.0\_genomic.Sc9M7eS\_2\_HRSCAF\_41/{sample}.merged.rmdup.merged.{processed}.snps5.noIndel.QUAL30.dp.AB.repma.vcf | |
| Output files | |
| - results/{dataset}/vcf/GCF\_000283155.1\_CerSimSim1.0\_genomic.Sc9M7eS\_2\_HRSCAF\_41/{sample,[A-Za-z0-9]+}.merged.rmdup.merged.{processed}.snps5.noIndel.QUAL30.dp.AB.repma.bcf | |
| Container image | |
| docker://quay.io/biocontainers/bcftools:1.9--h68d8f2e\_9 | |
| Code | |
| ``` 1 2 ``` \| ``` bcftools convert -O b -o {output.bcf} {input.filtered} 2> {log} ``` | |

###### Rule remove\_repeats\_vcf

×

Rule properties

|  |  |
| --- | --- |
| Jobs | 6 |
| Input files | |
| - results/{dataset}/vcf/GCF\_000283155.1\_CerSimSim1.0\_genomic.Sc9M7eS\_2\_HRSCAF\_41/{sample}.merged.rmdup.merged.{processed}.snps5.noIndel.QUAL30.dp.AB.vcf.gz - results/GCF\_000283155.1\_CerSimSim1.0\_genomic.Sc9M7eS\_2\_HRSCAF\_41.repma.bed - /proj/sllstore2017093/b2016342/b2016342\_nobackup/lts/genome\_erosion\_pipeline/verena\_testing/testdata/reference/GCF\_000283155.1\_CerSimSim1.0\_genomic.Sc9M7eS\_2\_HRSCAF\_41.genome | |
| Output files | |
| - results/{dataset}/vcf/GCF\_000283155.1\_CerSimSim1.0\_genomic.Sc9M7eS\_2\_HRSCAF\_41/{sample,[A-Za-z0-9]+}.merged.rmdup.merged.{processed}.snps5.noIndel.QUAL30.dp.AB.repma.vcf | |
| Container image | |
| docker://quay.io/biocontainers/bedtools:2.29.2--hc088bd4\_0 | |
| Code | |
| ``` 1 2 ``` \| ``` bedtools intersect -a {input.vcf} -b {input.bed} -header -sorted -g {input.genomefile} > {output.filtered} 2> {log} ``` | |

###### Rule filtered\_bcf2vcf

×

Rule properties

|  |  |
| --- | --- |
| Jobs | 6 |
| Input files | |
| - results/{dataset}/vcf/GCF\_000283155.1\_CerSimSim1.0\_genomic.Sc9M7eS\_2\_HRSCAF\_41/{sample}.merged.rmdup.merged.{processed}.snps5.noIndel.QUAL30.dp.AB.bcf | |
| Output files | |
| - results/{dataset}/vcf/GCF\_000283155.1\_CerSimSim1.0\_genomic.Sc9M7eS\_2\_HRSCAF\_41/{sample,[A-Za-z0-9]+}.merged.rmdup.merged.{processed}.snps5.noIndel.QUAL30.dp.AB.vcf.gz | |
| Container image | |
| docker://quay.io/biocontainers/bcftools:1.9--h68d8f2e\_9 | |
| Code | |
| ``` 1 2 ``` \| ``` bcftools convert -O z -o {output.vcf} {input.bcf} 2> {log} ``` | |

###### Rule index\_repmasked\_vcfs

×

Rule properties

|  |  |
| --- | --- |
| Jobs | 6 |
| Input files | |
| - results/{dataset}/vcf/GCF\_000283155.1\_CerSimSim1.0\_genomic.Sc9M7eS\_2\_HRSCAF\_41/{sample}.merged.rmdup.merged.{processed}.snps5.noIndel.QUAL30.dp.AB.repma.bcf | |
| Output files | |
| - results/{dataset}/vcf/GCF\_000283155.1\_CerSimSim1.0\_genomic.Sc9M7eS\_2\_HRSCAF\_41/{sample,[A-Za-z0-9]+}.merged.rmdup.merged.{processed}.snps5.noIndel.QUAL30.dp.AB.repma.bcf.csi | |
| Container image | |
| docker://quay.io/biocontainers/bcftools:1.9--h68d8f2e\_9 | |
| Code | |
| ``` 1 2 ``` \| ``` bcftools index -o {output.index} {input.bcf} 2> {log} ``` | |

###### Rule modern\_quality\_filtered\_vcf\_multiqc

×

Rule properties

|  |  |
| --- | --- |
| Jobs | 1 |
| Input files | |
| Output files | |
| - results/modern/vcf/GCF\_000283155.1\_CerSimSim1.0\_genomic.Sc9M7eS\_2\_HRSCAF\_41/stats/vcf\_qual\_filtered/multiqc/multiqc\_report.html | |
| Container image | |
| docker://quay.io/biocontainers/multiqc:1.9--pyh9f0ad1d\_0 | |
| Code | |
| ``` 1 2 ``` \| ``` multiqc -f {params.indir} -o {params.outdir} 2> {log} ``` | |

###### Rule modern\_repmasked\_vcf\_multiqc

×

Rule properties

|  |  |
| --- | --- |
| Jobs | 1 |
| Input files | |
| Output files | |
| - results/modern/vcf/GCF\_000283155.1\_CerSimSim1.0\_genomic.Sc9M7eS\_2\_HRSCAF\_41/stats/vcf\_repmasked/multiqc/multiqc\_report.html | |
| Container image | |
| docker://quay.io/biocontainers/multiqc:1.9--pyh9f0ad1d\_0 | |
| Code | |
| ``` 1 2 ``` \| ``` multiqc -f {params.indir} -o {params.outdir} 2> {log} ``` | |

###### Rule missingness\_filtered\_vcf\_multiqc

×

Rule properties

|  |  |
| --- | --- |
| Jobs | 1 |
| Input files | |
| Output files | |
| - results/all/vcf/GCF\_000283155.1\_CerSimSim1.0\_genomic.Sc9M7eS\_2\_HRSCAF\_41/stats/vcf\_merged\_missing/multiqc/multiqc\_report.html | |
| Container image | |
| docker://quay.io/biocontainers/multiqc:1.9--pyh9f0ad1d\_0 | |
| Code | |
| ``` 1 2 ``` \| ``` multiqc -f {params.indir} -o {params.outdir} 2> {log} ``` | |

###### Rule missingness\_filtered\_vcf\_stats

×

Rule properties

|  |  |
| --- | --- |
| Jobs | 3 |
| Input files | |
| - results/{dataset}/vcf/GCF\_000283155.1\_CerSimSim1.0\_genomic.Sc9M7eS\_2\_HRSCAF\_41.{dataset}.merged.biallelic.fmissing{fmiss}.vcf.gz - results/{dataset}/vcf/GCF\_000283155.1\_CerSimSim1.0\_genomic.Sc9M7eS\_2\_HRSCAF\_41.{dataset}.merged.biallelic.fmissing{fmiss}.vcf.gz.csi | |
| Output files | |
| - results/{dataset}/vcf/GCF\_000283155.1\_CerSimSim1.0\_genomic.Sc9M7eS\_2\_HRSCAF\_41/stats/vcf\_merged\_missing/GCF\_000283155.1\_CerSimSim1.0\_genomic.Sc9M7eS\_2\_HRSCAF\_41.{dataset}.merged.biallelic.fmissing{fmiss}.vcf.stats.txt | |
| Container image | |
| docker://quay.io/biocontainers/bcftools:1.9--h68d8f2e\_9 | |
| Code | |
| ``` 1 2 ``` \| ``` bcftools stats {input.merged} > {output.stats} 2> {log} ``` | |

###### Rule filter\_vcf\_missing

×

Rule properties

|  |  |
| --- | --- |
| Jobs | 1 |
| Input files | |
| - results/all/vcf/GCF\_000283155.1\_CerSimSim1.0\_genomic.Sc9M7eS\_2\_HRSCAF\_41.all.merged.biallelic.bcf - results/all/vcf/GCF\_000283155.1\_CerSimSim1.0\_genomic.Sc9M7eS\_2\_HRSCAF\_41.all.merged.biallelic.bcf.csi - results/all/vcf/GCF\_000283155.1\_CerSimSim1.0\_genomic.Sc9M7eS\_2\_HRSCAF\_41/stats/vcf\_merged\_biallelic/GCF\_000283155.1\_CerSimSim1.0\_genomic.Sc9M7eS\_2\_HRSCAF\_41.all.merged.biallelic.vcf.stats.txt - results/all/vcf/GCF\_000283155.1\_CerSimSim1.0\_genomic.Sc9M7eS\_2\_HRSCAF\_41/stats/vcf\_merged\_biallelic/multiqc/multiqc\_report.html | |
| Output files | |
| - results/all/vcf/GCF\_000283155.1\_CerSimSim1.0\_genomic.Sc9M7eS\_2\_HRSCAF\_41.all.merged.biallelic.fmissing{fmiss}.vcf.gz - results/all/vcf/GCF\_000283155.1\_CerSimSim1.0\_genomic.Sc9M7eS\_2\_HRSCAF\_41.all.merged.biallelic.fmissing{fmiss}.vcf.gz.csi | |
| Container image | |
| docker://quay.io/biocontainers/bcftools:1.9--h68d8f2e\_9 | |
| Code | |
| ``` 1 2 3 ``` \| ``` bcftools view -i 'F_MISSING < {params.fmiss}' -Oz -o {output.vcf} {input.bcf} 2> {log} && bcftools index -f {output.vcf} 2>> {log} ``` | |

###### Rule filter\_vcf\_biallelic

×

Rule properties

|  |  |
| --- | --- |
| Jobs | 1 |
| Input files | |
| - results/all/vcf/GCF\_000283155.1\_CerSimSim1.0\_genomic.Sc9M7eS\_2\_HRSCAF\_41.all.merged.snps.bcf - results/all/vcf/GCF\_000283155.1\_CerSimSim1.0\_genomic.Sc9M7eS\_2\_HRSCAF\_41.all.merged.snps.bcf.csi - results/all/vcf/GCF\_000283155.1\_CerSimSim1.0\_genomic.Sc9M7eS\_2\_HRSCAF\_41/stats/vcf\_merged/GCF\_000283155.1\_CerSimSim1.0\_genomic.Sc9M7eS\_2\_HRSCAF\_41.all.merged.snps.bcf.stats.txt - results/all/vcf/GCF\_000283155.1\_CerSimSim1.0\_genomic.Sc9M7eS\_2\_HRSCAF\_41/stats/vcf\_merged/multiqc/multiqc\_report.html | |
| Output files | |
| - results/all/vcf/GCF\_000283155.1\_CerSimSim1.0\_genomic.Sc9M7eS\_2\_HRSCAF\_41.all.merged.biallelic.bcf - results/all/vcf/GCF\_000283155.1\_CerSimSim1.0\_genomic.Sc9M7eS\_2\_HRSCAF\_41.all.merged.biallelic.bcf.csi | |
| Container image | |
| docker://quay.io/biocontainers/bcftools:1.9--h68d8f2e\_9 | |
| Code | |
| ``` 1 2 3 ``` \| ``` bcftools view -m2 -M2 -v snps -Ob -o {output.bcf} {input.bcf} 2> {log} && bcftools index -f {output.bcf} 2>> {log} ``` | |

###### Rule merge\_all\_vcfs

×

Rule properties

|  |  |
| --- | --- |
| Jobs | 1 |
| Input files | |
| Output files | |
| - results/all/vcf/GCF\_000283155.1\_CerSimSim1.0\_genomic.Sc9M7eS\_2\_HRSCAF\_41.all.merged.snps.bcf | |
| Container image | |
| docker://quay.io/biocontainers/bcftools:1.9--h68d8f2e\_9 | |
| Code | |
| ``` 1 2 3 4 5 6 7 8 9 ``` \| ``` files=`echo {input.bcf} \| awk '{{print NF}}'` if [ $files -gt 1 ] # check if there are at least 2 files for merging. If there is only one file, copy the bcf file. then bcftools merge -m snps -O b -o {output.merged} {input.bcf} 2> {log} else cp {input.bcf} {output.merged} && touch {output.merged} 2> {log} echo "Only one file present for merging. Copying the input bcf file." >> {log} fi ``` | |

###### Rule index\_merged\_vcf

×

Rule properties

|  |  |
| --- | --- |
| Jobs | 1 |
| Input files | |
| - results/all/vcf/GCF\_000283155.1\_CerSimSim1.0\_genomic.Sc9M7eS\_2\_HRSCAF\_41.all.merged.snps.bcf | |
| Output files | |
| - results/all/vcf/GCF\_000283155.1\_CerSimSim1.0\_genomic.Sc9M7eS\_2\_HRSCAF\_41.all.merged.snps.bcf.csi | |
| Container image | |
| docker://quay.io/biocontainers/bcftools:1.9--h68d8f2e\_9 | |
| Code | |
| ``` 1 2 ``` \| ``` bcftools index -o {output.index} {input.bcf} 2> {log} ``` | |

###### Rule merged\_vcf\_stats

×

Rule properties

|  |  |
| --- | --- |
| Jobs | 1 |
| Input files | |
| - results/all/vcf/GCF\_000283155.1\_CerSimSim1.0\_genomic.Sc9M7eS\_2\_HRSCAF\_41.all.merged.snps.bcf - results/all/vcf/GCF\_000283155.1\_CerSimSim1.0\_genomic.Sc9M7eS\_2\_HRSCAF\_41.all.merged.snps.bcf.csi | |
| Output files | |
| - results/all/vcf/GCF\_000283155.1\_CerSimSim1.0\_genomic.Sc9M7eS\_2\_HRSCAF\_41/stats/vcf\_merged/GCF\_000283155.1\_CerSimSim1.0\_genomic.Sc9M7eS\_2\_HRSCAF\_41.all.merged.snps.bcf.stats.txt | |
| Container image | |
| docker://quay.io/biocontainers/bcftools:1.9--h68d8f2e\_9 | |
| Code | |
| ``` 1 2 ``` \| ``` bcftools stats {input.merged} > {output.stats} 2> {log} ``` | |

###### Rule merged\_vcf\_multiqc

×

Rule properties

|  |  |
| --- | --- |
| Jobs | 1 |
| Input files | |
| - results/all/vcf/GCF\_000283155.1\_CerSimSim1.0\_genomic.Sc9M7eS\_2\_HRSCAF\_41/stats/vcf\_merged/GCF\_000283155.1\_CerSimSim1.0\_genomic.Sc9M7eS\_2\_HRSCAF\_41.all.merged.snps.bcf.stats.txt | |
| Output files | |
| - results/all/vcf/GCF\_000283155.1\_CerSimSim1.0\_genomic.Sc9M7eS\_2\_HRSCAF\_41/stats/vcf\_merged/multiqc/multiqc\_report.html | |
| Container image | |
| docker://quay.io/biocontainers/multiqc:1.9--pyh9f0ad1d\_0 | |
| Code | |
| ``` 1 2 ``` \| ``` multiqc -f {params.indir} -o {params.outdir} 2> {log} ``` | |

###### Rule biallelic\_filtered\_vcf\_stats

×

Rule properties

|  |  |
| --- | --- |
| Jobs | 1 |
| Input files | |
| - results/all/vcf/GCF\_000283155.1\_CerSimSim1.0\_genomic.Sc9M7eS\_2\_HRSCAF\_41.all.merged.biallelic.bcf - results/all/vcf/GCF\_000283155.1\_CerSimSim1.0\_genomic.Sc9M7eS\_2\_HRSCAF\_41.all.merged.biallelic.bcf.csi | |
| Output files | |
| - results/all/vcf/GCF\_000283155.1\_CerSimSim1.0\_genomic.Sc9M7eS\_2\_HRSCAF\_41/stats/vcf\_merged\_biallelic/GCF\_000283155.1\_CerSimSim1.0\_genomic.Sc9M7eS\_2\_HRSCAF\_41.all.merged.biallelic.vcf.stats.txt | |
| Container image | |
| docker://quay.io/biocontainers/bcftools:1.9--h68d8f2e\_9 | |
| Code | |
| ``` 1 2 ``` \| ``` bcftools stats {input.bcf} > {output.stats} 2> {log} ``` | |

###### Rule biallelic\_filtered\_vcf\_multiqc

×

Rule properties

|  |  |
| --- | --- |
| Jobs | 1 |
| Input files | |
| - results/all/vcf/GCF\_000283155.1\_CerSimSim1.0\_genomic.Sc9M7eS\_2\_HRSCAF\_41/stats/vcf\_merged\_biallelic/GCF\_000283155.1\_CerSimSim1.0\_genomic.Sc9M7eS\_2\_HRSCAF\_41.all.merged.biallelic.vcf.stats.txt | |
| Output files | |
| - results/all/vcf/GCF\_000283155.1\_CerSimSim1.0\_genomic.Sc9M7eS\_2\_HRSCAF\_41/stats/vcf\_merged\_biallelic/multiqc/multiqc\_report.html | |
| Container image | |
| docker://quay.io/biocontainers/multiqc:1.9--pyh9f0ad1d\_0 | |
| Code | |
| ``` 1 2 ``` \| ``` multiqc -f {params.indir} -o {params.outdir} 2> {log} ``` | |

###### Rule extract\_historical\_samples

×

Rule properties

|  |  |
| --- | --- |
| Jobs | 1 |
| Input files | |
| - results/all/vcf/GCF\_000283155.1\_CerSimSim1.0\_genomic.Sc9M7eS\_2\_HRSCAF\_41.all.merged.biallelic.fmissing{fmiss}.vcf.gz - results/all/vcf/GCF\_000283155.1\_CerSimSim1.0\_genomic.Sc9M7eS\_2\_HRSCAF\_41.all.merged.biallelic.fmissing{fmiss}.vcf.gz.csi - results/all/vcf/GCF\_000283155.1\_CerSimSim1.0\_genomic.Sc9M7eS\_2\_HRSCAF\_41/stats/vcf\_merged\_biallelic/GCF\_000283155.1\_CerSimSim1.0\_genomic.Sc9M7eS\_2\_HRSCAF\_41.all.merged.biallelic.vcf.stats.txt - results/all/vcf/GCF\_000283155.1\_CerSimSim1.0\_genomic.Sc9M7eS\_2\_HRSCAF\_41.all.merged.biallelic.fmissing{fmiss}.bed | |
| Output files | |
| - results/historical/vcf/GCF\_000283155.1\_CerSimSim1.0\_genomic.Sc9M7eS\_2\_HRSCAF\_41.historical.merged.biallelic.fmissing{fmiss}.vcf.gz - results/historical/vcf/GCF\_000283155.1\_CerSimSim1.0\_genomic.Sc9M7eS\_2\_HRSCAF\_41.historical.merged.biallelic.fmissing{fmiss}.vcf.gz.csi | |
| Container image | |
| docker://quay.io/biocontainers/bcftools:1.9--h68d8f2e\_9 | |
| Code | |
| ``` 1 2 3 4 5 6 7 8 9 10 11 12 13 14 ``` \| ``` samples_edited=`echo {params.samples} \| sed 's/ /,/g'` samples_len=`echo {params.samples} \| wc -w` # count the number of historical samples all_samples_len=`echo {params.all_samples} \| wc -w` # count the number of all samples if [ $samples_len != $all_samples_len ] then bcftools view -Oz -s $samples_edited -o {output.vcf} {input.vcf} 2> {log} && bcftools index -f {output.vcf} 2>> {log} else cp {input.vcf} {output.vcf} && touch {output.vcf} 2> {log} && bcftools index -f {output.vcf} 2>> {log} echo "Only historical samples present. Copying the input vcf file." >> {log} fi ``` | |

###### Rule filtered\_vcf2bed

×

Rule properties

|  |  |
| --- | --- |
| Jobs | 1 |
| Input files | |
| - results/all/vcf/GCF\_000283155.1\_CerSimSim1.0\_genomic.Sc9M7eS\_2\_HRSCAF\_41.all.merged.biallelic.fmissing{fmiss}.vcf.gz | |
| Output files | |
| - results/all/vcf/GCF\_000283155.1\_CerSimSim1.0\_genomic.Sc9M7eS\_2\_HRSCAF\_41.all.merged.biallelic.fmissing{fmiss}.bed | |
| Container image | |
| docker://quay.io/biocontainers/bedtools:2.29.2--hc088bd4\_0 | |
| Code | |
| ``` 1 2 ``` \| ``` gzip -cd {input.vcf} \| grep -v "^#" \| awk -F'	' '{{print $1, $2-1, $2}}' OFS='	' > {output.bed} 2> {log} ``` | |

###### Rule extract\_modern\_samples

×

Rule properties

|  |  |
| --- | --- |
| Jobs | 1 |
| Input files | |
| - results/all/vcf/GCF\_000283155.1\_CerSimSim1.0\_genomic.Sc9M7eS\_2\_HRSCAF\_41.all.merged.biallelic.fmissing{fmiss}.vcf.gz - results/all/vcf/GCF\_000283155.1\_CerSimSim1.0\_genomic.Sc9M7eS\_2\_HRSCAF\_41.all.merged.biallelic.fmissing{fmiss}.vcf.gz.csi - results/all/vcf/GCF\_000283155.1\_CerSimSim1.0\_genomic.Sc9M7eS\_2\_HRSCAF\_41/stats/vcf\_merged\_biallelic/GCF\_000283155.1\_CerSimSim1.0\_genomic.Sc9M7eS\_2\_HRSCAF\_41.all.merged.biallelic.vcf.stats.txt - results/all/vcf/GCF\_000283155.1\_CerSimSim1.0\_genomic.Sc9M7eS\_2\_HRSCAF\_41.all.merged.biallelic.fmissing{fmiss}.bed | |
| Output files | |
| - results/modern/vcf/GCF\_000283155.1\_CerSimSim1.0\_genomic.Sc9M7eS\_2\_HRSCAF\_41.modern.merged.biallelic.fmissing{fmiss}.vcf.gz - results/modern/vcf/GCF\_000283155.1\_CerSimSim1.0\_genomic.Sc9M7eS\_2\_HRSCAF\_41.modern.merged.biallelic.fmissing{fmiss}.vcf.gz.csi | |
| Container image | |
| docker://quay.io/biocontainers/bcftools:1.9--h68d8f2e\_9 | |
| Code | |
| ``` 1 2 3 4 5 6 7 8 9 10 11 12 13 14 ``` \| ``` samples_edited=`echo {params.samples} \| sed 's/ /,/g'` samples_len=`echo {params.samples} \| wc -w` # count the number of historical samples all_samples_len=`echo {params.all_samples} \| wc -w` # count the number of all samples if [ $samples_len != $all_samples_len ] then bcftools view -Oz -s $samples_edited -o {output.vcf} {input.vcf} 2> {log} && bcftools index -f {output.vcf} 2>> {log} else cp {input.vcf} {output.vcf} && touch {output.vcf} 2> {log} && bcftools index -f {output.vcf} 2>> {log} echo "Only modern samples present. Copying the input vcf file." >> {log} fi ``` | |

###### Rule plot\_pc1\_pc2

×

Rule properties

|  |  |
| --- | --- |
| Jobs | 3 |
| Input files | |
| - results/{dataset}/pca/GCF\_000283155.1\_CerSimSim1.0\_genomic.Sc9M7eS\_2\_HRSCAF\_41.{dataset}.merged.biallelic.fmissing{fmiss}.eigenvec - results/{dataset}/pca/GCF\_000283155.1\_CerSimSim1.0\_genomic.Sc9M7eS\_2\_HRSCAF\_41.{dataset}.merged.biallelic.fmissing{fmiss}.eigenval | |
| Output files | |
| - results/{dataset}/pca/GCF\_000283155.1\_CerSimSim1.0\_genomic.Sc9M7eS\_2\_HRSCAF\_41.{dataset}.merged.biallelic.fmissing{fmiss}.pc1\_pc2.pdf | |

###### Rule plink\_eigenvec

×

Rule properties

|  |  |
| --- | --- |
| Jobs | 3 |
| Input files | |
| - results/{dataset}/pca/GCF\_000283155.1\_CerSimSim1.0\_genomic.Sc9M7eS\_2\_HRSCAF\_41.{dataset}.merged.biallelic.fmissing{fmiss}.bed - results/{dataset}/pca/GCF\_000283155.1\_CerSimSim1.0\_genomic.Sc9M7eS\_2\_HRSCAF\_41.{dataset}.merged.biallelic.fmissing{fmiss}.bim - results/{dataset}/pca/GCF\_000283155.1\_CerSimSim1.0\_genomic.Sc9M7eS\_2\_HRSCAF\_41.{dataset}.merged.biallelic.fmissing{fmiss}.fam - results/{dataset}/pca/GCF\_000283155.1\_CerSimSim1.0\_genomic.Sc9M7eS\_2\_HRSCAF\_41.{dataset}.merged.biallelic.fmissing{fmiss}.nosex | |
| Output files | |
| - results/{dataset}/pca/GCF\_000283155.1\_CerSimSim1.0\_genomic.Sc9M7eS\_2\_HRSCAF\_41.{dataset}.merged.biallelic.fmissing{fmiss}.eigenvec - results/{dataset}/pca/GCF\_000283155.1\_CerSimSim1.0\_genomic.Sc9M7eS\_2\_HRSCAF\_41.{dataset}.merged.biallelic.fmissing{fmiss}.eigenval | |
| Container image | |
| docker://quay.io/biocontainers/plink:1.90b6.12--heea4ae3\_0 | |
| Code | |
| ``` 1 2 3 4 5 6 7 8 9 ``` \| ``` samples=`cat {input.fam} \| wc -l` if [ "$samples" -gt 1 ] then plink --bfile {params.bfile} --allow-extra-chr --pca --out {params.bfile} 2> {log} else touch {output.eigenvec} && touch {output.eigenval} 2> {log} echo "Not enough samples to calculate a PCA." >> {log} fi ``` | |

###### Rule vcf2plink\_pca

×

Rule properties

|  |  |
| --- | --- |
| Jobs | 3 |
| Input files | |
| - results/{dataset}/vcf/GCF\_000283155.1\_CerSimSim1.0\_genomic.Sc9M7eS\_2\_HRSCAF\_41.{dataset}.merged.biallelic.fmissing{fmiss}.vcf.gz - results/{dataset}/vcf/GCF\_000283155.1\_CerSimSim1.0\_genomic.Sc9M7eS\_2\_HRSCAF\_41.{dataset}.merged.biallelic.fmissing{fmiss}.vcf.gz.csi | |
| Output files | |
| - results/{dataset}/pca/GCF\_000283155.1\_CerSimSim1.0\_genomic.Sc9M7eS\_2\_HRSCAF\_41.{dataset}.merged.biallelic.fmissing{fmiss}.bed - results/{dataset}/pca/GCF\_000283155.1\_CerSimSim1.0\_genomic.Sc9M7eS\_2\_HRSCAF\_41.{dataset}.merged.biallelic.fmissing{fmiss}.bim - results/{dataset}/pca/GCF\_000283155.1\_CerSimSim1.0\_genomic.Sc9M7eS\_2\_HRSCAF\_41.{dataset}.merged.biallelic.fmissing{fmiss}.fam - results/{dataset}/pca/GCF\_000283155.1\_CerSimSim1.0\_genomic.Sc9M7eS\_2\_HRSCAF\_41.{dataset}.merged.biallelic.fmissing{fmiss}.nosex | |
| Container image | |
| docker://quay.io/biocontainers/plink:1.90b6.12--heea4ae3\_0 | |
| Code | |
| ``` 1 2 ``` \| ``` plink --vcf {input.vcf} --make-bed --allow-extra-chr --out {params.bfile} 2> {log} ``` | |

###### Rule plot\_pc1\_pc3

×

Rule properties

|  |  |
| --- | --- |
| Jobs | 3 |
| Input files | |
| - results/{dataset}/pca/GCF\_000283155.1\_CerSimSim1.0\_genomic.Sc9M7eS\_2\_HRSCAF\_41.{dataset}.merged.biallelic.fmissing{fmiss}.eigenvec - results/{dataset}/pca/GCF\_000283155.1\_CerSimSim1.0\_genomic.Sc9M7eS\_2\_HRSCAF\_41.{dataset}.merged.biallelic.fmissing{fmiss}.eigenval | |
| Output files | |
| - results/{dataset}/pca/GCF\_000283155.1\_CerSimSim1.0\_genomic.Sc9M7eS\_2\_HRSCAF\_41.{dataset}.merged.biallelic.fmissing{fmiss}.pc1\_pc3.pdf | |

###### Rule snpEff\_variant\_impact\_plot

×

Rule properties

|  |  |
| --- | --- |
| Jobs | 1 |
| Input files | |
| - results/{dataset}/snpEff/GCF\_000283155.1\_CerSimSim1.0\_genomic.Sc9M7eS\_2\_HRSCAF\_41.{dataset}.fmissing{fmiss}.snpEff\_variant\_impact\_table.txt | |
| Output files | |
| - results/{dataset}/snpEff/GCF\_000283155.1\_CerSimSim1.0\_genomic.Sc9M7eS\_2\_HRSCAF\_41.{dataset}.fmissing{fmiss}.snpEff\_variant\_impact\_plot.pdf | |

###### Rule snpEff\_variant\_impact\_table

×

Rule properties

|  |  |
| --- | --- |
| Jobs | 1 |
| Input files | |
| Output files | |
| - results/{dataset}/snpEff/GCF\_000283155.1\_CerSimSim1.0\_genomic.Sc9M7eS\_2\_HRSCAF\_41.{dataset}.fmissing{fmiss}.snpEff\_variant\_impact\_table.txt | |

###### Rule annotate\_vcf

×

Rule properties

|  |  |
| --- | --- |
| Jobs | 6 |
| Input files | |
| - results/{dataset}/snpEff/GCF\_000283155.1\_CerSimSim1.0\_genomic.Sc9M7eS\_2\_HRSCAF\_41/{sample}.merged.rmdup.merged.{processed}.snps5.noIndel.QUAL30.dp.AB.repma.biallelic.fmissing{fmiss}.vcf - /proj/sllstore2017093/b2016342/b2016342\_nobackup/lts/genome\_erosion\_pipeline/verena\_testing/testdata/reference/snpEff/data/GCF\_000283155.1\_CerSimSim1.0\_genomic.Sc9M7eS\_2\_HRSCAF\_41/snpEffectPredictor.bin - /proj/sllstore2017093/b2016342/b2016342\_nobackup/lts/genome\_erosion\_pipeline/verena\_testing/testdata/reference/snpEff/data/GCF\_000283155.1\_CerSimSim1.0\_genomic.Sc9M7eS\_2\_HRSCAF\_41/snpEff.config | |
| Output files | |
| - results/{dataset}/snpEff/GCF\_000283155.1\_CerSimSim1.0\_genomic.Sc9M7eS\_2\_HRSCAF\_41/{sample,[A-Za-z0-9]+}.merged.rmdup.merged.{processed}.snps5.noIndel.QUAL30.dp.AB.repma.biallelic.fmissing{fmiss}.ann.vcf - results/{dataset}/snpEff/GCF\_000283155.1\_CerSimSim1.0\_genomic.Sc9M7eS\_2\_HRSCAF\_41/{sample,[A-Za-z0-9]+}.merged.rmdup.merged.{processed}.snps5.noIndel.QUAL30.dp.AB.repma.biallelic.fmissing{fmiss}\_stats.csv - results/{dataset}/snpEff/GCF\_000283155.1\_CerSimSim1.0\_genomic.Sc9M7eS\_2\_HRSCAF\_41/{sample,[A-Za-z0-9]+}.merged.rmdup.merged.{processed}.snps5.noIndel.QUAL30.dp.AB.repma.biallelic.fmissing{fmiss}\_stats.html | |
| Container image | |
| docker://quay.io/biocontainers/snpeff:4.3.1t--3 | |
| Code | |
| ``` 1 2 ``` \| ``` snpEff -c {params.abs_config} -dataDir {params.abs_data_dir} -s {output.html} -csvStats {output.csv} -treatAllAsProteinCoding -v -d -lof {params.ref_name} {input.vcf} > {output.ann} 2> {log} ``` | |

###### Rule filter\_biallelic\_missing\_vcf\_snpEff

×

Rule properties

|  |  |
| --- | --- |
| Jobs | 6 |
| Input files | |
| - results/{dataset}/snpEff/GCF\_000283155.1\_CerSimSim1.0\_genomic.Sc9M7eS\_2\_HRSCAF\_41/{sample}.merged.rmdup.merged.{processed}.snps5.noIndel.QUAL30.dp.AB.repma.vcf.gz - results/all/vcf/GCF\_000283155.1\_CerSimSim1.0\_genomic.Sc9M7eS\_2\_HRSCAF\_41.all.merged.biallelic.fmissing{fmiss}.bed - /proj/sllstore2017093/b2016342/b2016342\_nobackup/lts/genome\_erosion\_pipeline/verena\_testing/testdata/reference/GCF\_000283155.1\_CerSimSim1.0\_genomic.Sc9M7eS\_2\_HRSCAF\_41.genome | |
| Output files | |
| - results/{dataset}/snpEff/GCF\_000283155.1\_CerSimSim1.0\_genomic.Sc9M7eS\_2\_HRSCAF\_41/{sample,[A-Za-z0-9]+}.merged.rmdup.merged.{processed}.snps5.noIndel.QUAL30.dp.AB.repma.biallelic.fmissing{fmiss}.vcf | |
| Container image | |
| docker://quay.io/biocontainers/bedtools:2.29.2--hc088bd4\_0 | |
| Code | |
| ``` 1 2 ``` \| ``` bedtools intersect -a {input.vcf} -b {input.bed} -header -sorted -g {input.genomefile} > {output.filtered} 2> {log} ``` | |

###### Rule repmasked\_bcf2vcf\_snpEff

×

Rule properties

|  |  |
| --- | --- |
| Jobs | 6 |
| Input files | |
| - results/{dataset}/vcf/GCF\_000283155.1\_CerSimSim1.0\_genomic.Sc9M7eS\_2\_HRSCAF\_41/{sample}.merged.rmdup.merged.{processed}.snps5.noIndel.QUAL30.dp.AB.repma.bcf - results/{dataset}/vcf/GCF\_000283155.1\_CerSimSim1.0\_genomic.Sc9M7eS\_2\_HRSCAF\_41/{sample}.merged.rmdup.merged.{processed}.snps5.noIndel.QUAL30.dp.AB.repma.bcf.csi | |
| Output files | |
| - results/{dataset}/snpEff/GCF\_000283155.1\_CerSimSim1.0\_genomic.Sc9M7eS\_2\_HRSCAF\_41/{sample,[A-Za-z0-9]+}.merged.rmdup.merged.{processed}.snps5.noIndel.QUAL30.dp.AB.repma.vcf.gz | |
| Container image | |
| docker://quay.io/biocontainers/bcftools:1.9--h68d8f2e\_9 | |
| Code | |
| ``` 1 2 ``` \| ``` bcftools convert -O z -o {output.vcf} {input.bcf} 2> {log} ``` | |

###### Rule historical\_snpEff\_multiqc

×

Rule properties

|  |  |
| --- | --- |
| Jobs | 1 |
| Input files | |
| Output files | |
| - results/historical/snpEff/GCF\_000283155.1\_CerSimSim1.0\_genomic.Sc9M7eS\_2\_HRSCAF\_41/multiqc/multiqc\_report.html | |
| Container image | |
| docker://quay.io/biocontainers/multiqc:1.9--pyh9f0ad1d\_0 | |
| Code | |
| ``` 1 2 ``` \| ``` multiqc -f {params.indir} -o {params.outdir} 2> {log} ``` | |

###### Rule modern\_snpEff\_multiqc

×

Rule properties

|  |  |
| --- | --- |
| Jobs | 1 |
| Input files | |
| Output files | |
| - results/modern/snpEff/GCF\_000283155.1\_CerSimSim1.0\_genomic.Sc9M7eS\_2\_HRSCAF\_41/multiqc/multiqc\_report.html | |
| Container image | |
| docker://quay.io/biocontainers/multiqc:1.9--pyh9f0ad1d\_0 | |
| Code | |
| ``` 1 2 ``` \| ``` multiqc -f {params.indir} -o {params.outdir} 2> {log} ``` | |

###### Rule plot\_gerp\_hist

×

Rule properties

|  |  |
| --- | --- |
| Jobs | 1 |
| Input files | |
| - results/gerp/GCF\_000283155.1\_CerSimSim1.0\_genomic.Sc9M7eS\_2\_HRSCAF\_41.ancestral.rates.gz | |
| Output files | |
| - results/gerp/GCF\_000283155.1\_CerSimSim1.0\_genomic.Sc9M7eS\_2\_HRSCAF\_41.ancestral.rates.gerp.hist.pdf | |

###### Rule merge\_gerp\_gz

×

Rule properties

|  |  |
| --- | --- |
| Jobs | 1 |
| Input files | |
| - results/gerp/chunks/GCF\_000283155.1\_CerSimSim1.0\_genomic.Sc9M7eS\_2\_HRSCAF\_41/gerp/chunk1.fasta.parsed.rates - results/gerp/chunks/GCF\_000283155.1\_CerSimSim1.0\_genomic.Sc9M7eS\_2\_HRSCAF\_41/gerp/chunk2.fasta.parsed.rates - results/gerp/chunks/GCF\_000283155.1\_CerSimSim1.0\_genomic.Sc9M7eS\_2\_HRSCAF\_41/gerp/chunk3.fasta.parsed.rates | |
| Output files | |
| - results/gerp/GCF\_000283155.1\_CerSimSim1.0\_genomic.Sc9M7eS\_2\_HRSCAF\_41.ancestral.rates.gz | |
| Code | |
| ``` 1 2 ``` \| ``` cat {input.gerp_chunks_merged} \| gzip - > {output.gerp_out} 2> {log} ``` | |

###### Rule merge\_gerp\_per\_chunk

×

Rule properties

|  |  |
| --- | --- |
| Jobs | 3 |
| Input files | |
| - results/gerp/chunks/GCF\_000283155.1\_CerSimSim1.0\_genomic.Sc9M7eS\_2\_HRSCAF\_41/gerp/{chunk}\_gerp\_merged - /proj/sllstore2017093/b2016342/b2016342\_nobackup/lts/genome\_erosion\_pipeline/verena\_testing/testdata/reference/gerp/GCF\_000283155.1\_CerSimSim1.0\_genomic.Sc9M7eS\_2\_HRSCAF\_41/split\_bed\_files/{chunk}.bed | |
| Output files | |
| - results/gerp/chunks/GCF\_000283155.1\_CerSimSim1.0\_genomic.Sc9M7eS\_2\_HRSCAF\_41/gerp/{chunk,chunk[0-9]+}.fasta.parsed.rates | |

###### Rule produce\_contig\_out

×

Rule properties

|  |  |
| --- | --- |
| Jobs | 3 |
| Input files | |
| - results/gerp/chunks/GCF\_000283155.1\_CerSimSim1.0\_genomic.Sc9M7eS\_2\_HRSCAF\_41/gerp/{chunk}\_fasta\_ancestral - results/gerp/chunks/GCF\_000283155.1\_CerSimSim1.0\_genomic.Sc9M7eS\_2\_HRSCAF\_41/gerp/{chunk}\_gerp\_rescaled - /proj/sllstore2017093/b2016342/b2016342\_nobackup/lts/genome\_erosion\_pipeline/verena\_testing/testdata/reference/gerp/GCF\_000283155.1\_CerSimSim1.0\_genomic.Sc9M7eS\_2\_HRSCAF\_41/split\_bed\_files/{chunk}.bed | |
| Output files | |
| - results/gerp/chunks/GCF\_000283155.1\_CerSimSim1.0\_genomic.Sc9M7eS\_2\_HRSCAF\_41/gerp/{chunk,chunk[0-9]+}\_gerp\_merged | |

###### Rule get\_ancestral\_state

×

Rule properties

|  |  |
| --- | --- |
| Jobs | 3 |
| Input files | |
| - results/gerp/chunks/GCF\_000283155.1\_CerSimSim1.0\_genomic.Sc9M7eS\_2\_HRSCAF\_41/fasta/concatenated\_{chunk} - /proj/sllstore2017093/b2016342/b2016342\_nobackup/lts/genome\_erosion\_pipeline/verena\_testing/testdata/reference/gerp/GCF\_000283155.1\_CerSimSim1.0\_genomic.Sc9M7eS\_2\_HRSCAF\_41/split\_bed\_files/{chunk}.bed | |
| Output files | |
| - results/gerp/chunks/GCF\_000283155.1\_CerSimSim1.0\_genomic.Sc9M7eS\_2\_HRSCAF\_41/gerp/{chunk,chunk[0-9]+}\_fasta\_ancestral | |

###### Rule concatenate\_fasta\_per\_contig

×

Rule properties

|  |  |
| --- | --- |
| Jobs | 3 |
| Input files | |
| - results/gerp/chunks/GCF\_000283155.1\_CerSimSim1.0\_genomic.Sc9M7eS\_2\_HRSCAF\_41/fasta/Ovis\_aries\_{chunk} - results/gerp/chunks/GCF\_000283155.1\_CerSimSim1.0\_genomic.Sc9M7eS\_2\_HRSCAF\_41/fasta/Catagonus\_wagneri\_{chunk} - results/gerp/chunks/GCF\_000283155.1\_CerSimSim1.0\_genomic.Sc9M7eS\_2\_HRSCAF\_41/fasta/Lipotes\_vexillifer\_{chunk} - results/gerp/chunks/GCF\_000283155.1\_CerSimSim1.0\_genomic.Sc9M7eS\_2\_HRSCAF\_41/fasta/Cervus\_elaphus\_{chunk} - results/gerp/chunks/GCF\_000283155.1\_CerSimSim1.0\_genomic.Sc9M7eS\_2\_HRSCAF\_41/fasta/Physeter\_catodon\_{chunk} - results/gerp/chunks/GCF\_000283155.1\_CerSimSim1.0\_genomic.Sc9M7eS\_2\_HRSCAF\_41/fasta/Hyaena\_hyaena\_{chunk} - results/gerp/chunks/GCF\_000283155.1\_CerSimSim1.0\_genomic.Sc9M7eS\_2\_HRSCAF\_41/fasta/Paradoxurus\_hermaphroditus\_{chunk} - results/gerp/chunks/GCF\_000283155.1\_CerSimSim1.0\_genomic.Sc9M7eS\_2\_HRSCAF\_41/fasta/Procyon\_lotor\_{chunk} - results/gerp/chunks/GCF\_000283155.1\_CerSimSim1.0\_genomic.Sc9M7eS\_2\_HRSCAF\_41/fasta/Panthera\_leo\_{chunk} - results/gerp/chunks/GCF\_000283155.1\_CerSimSim1.0\_genomic.Sc9M7eS\_2\_HRSCAF\_41/fasta/Enhydra\_lutris\_{chunk} - results/gerp/chunks/GCF\_000283155.1\_CerSimSim1.0\_genomic.Sc9M7eS\_2\_HRSCAF\_41/fasta/Suricata\_suricatta\_{chunk} - results/gerp/chunks/GCF\_000283155.1\_CerSimSim1.0\_genomic.Sc9M7eS\_2\_HRSCAF\_41/fasta/Ailurus\_fulgens\_{chunk} - results/gerp/chunks/GCF\_000283155.1\_CerSimSim1.0\_genomic.Sc9M7eS\_2\_HRSCAF\_41/fasta/Antilocapra\_americana\_{chunk} - results/gerp/chunks/GCF\_000283155.1\_CerSimSim1.0\_genomic.Sc9M7eS\_2\_HRSCAF\_41/fasta/Camelus\_dromedarius\_{chunk} - results/gerp/chunks/GCF\_000283155.1\_CerSimSim1.0\_genomic.Sc9M7eS\_2\_HRSCAF\_41/fasta/Zalophus\_californianus\_{chunk} - results/gerp/chunks/GCF\_000283155.1\_CerSimSim1.0\_genomic.Sc9M7eS\_2\_HRSCAF\_41/fasta/Spilogale\_gracilis\_{chunk} - results/gerp/chunks/GCF\_000283155.1\_CerSimSim1.0\_genomic.Sc9M7eS\_2\_HRSCAF\_41/fasta/Diceros\_bicornis\_{chunk} - results/gerp/chunks/GCF\_000283155.1\_CerSimSim1.0\_genomic.Sc9M7eS\_2\_HRSCAF\_41/fasta/Balaenoptera\_acutorostrata\_{chunk} - results/gerp/chunks/GCF\_000283155.1\_CerSimSim1.0\_genomic.Sc9M7eS\_2\_HRSCAF\_41/fasta/Tapirus\_indicus\_{chunk} - results/gerp/chunks/GCF\_000283155.1\_CerSimSim1.0\_genomic.Sc9M7eS\_2\_HRSCAF\_41/fasta/Ursus\_maritimus\_{chunk} - results/gerp/chunks/GCF\_000283155.1\_CerSimSim1.0\_genomic.Sc9M7eS\_2\_HRSCAF\_41/fasta/Tragulus\_javanicus\_{chunk} - results/gerp/chunks/GCF\_000283155.1\_CerSimSim1.0\_genomic.Sc9M7eS\_2\_HRSCAF\_41/fasta/Manis\_javanica\_{chunk} - results/gerp/chunks/GCF\_000283155.1\_CerSimSim1.0\_genomic.Sc9M7eS\_2\_HRSCAF\_41/fasta/Giraffa\_camelopardalis\_{chunk} - results/gerp/chunks/GCF\_000283155.1\_CerSimSim1.0\_genomic.Sc9M7eS\_2\_HRSCAF\_41/fasta/Leptonychotes\_weddellii\_{chunk} - results/gerp/chunks/GCF\_000283155.1\_CerSimSim1.0\_genomic.Sc9M7eS\_2\_HRSCAF\_41/fasta/Equus\_asinus\_{chunk} - results/gerp/chunks/GCF\_000283155.1\_CerSimSim1.0\_genomic.Sc9M7eS\_2\_HRSCAF\_41/fasta/Mesoplodon\_bidens\_{chunk} - results/gerp/chunks/GCF\_000283155.1\_CerSimSim1.0\_genomic.Sc9M7eS\_2\_HRSCAF\_41/fasta/Tursiops\_truncatus\_{chunk} - results/gerp/chunks/GCF\_000283155.1\_CerSimSim1.0\_genomic.Sc9M7eS\_2\_HRSCAF\_41/fasta/Bubalus\_bubalis\_{chunk} - results/gerp/chunks/GCF\_000283155.1\_CerSimSim1.0\_genomic.Sc9M7eS\_2\_HRSCAF\_41/fasta/Canis\_lupus\_{chunk} - results/gerp/chunks/GCF\_000283155.1\_CerSimSim1.0\_genomic.Sc9M7eS\_2\_HRSCAF\_41/fasta/Hippopotamus\_amphibius\_{chunk} - results/gerp/chunks/GCF\_000283155.1\_CerSimSim1.0\_genomic.Sc9M7eS\_2\_HRSCAF\_41/fasta/GCF\_000283155.1\_CerSimSim1.0\_genomic.Sc9M7eS\_2\_HRSCAF\_41\_{chunk} - /proj/sllstore2017093/b2016342/b2016342\_nobackup/lts/genome\_erosion\_pipeline/verena\_testing/testdata/reference/gerp/GCF\_000283155.1\_CerSimSim1.0\_genomic.Sc9M7eS\_2\_HRSCAF\_41/split\_bed\_files/{chunk}.bed | |
| Output files | |
| - results/gerp/chunks/GCF\_000283155.1\_CerSimSim1.0\_genomic.Sc9M7eS\_2\_HRSCAF\_41/fasta/concatenated\_{chunk,chunk[0-9]+} | |

###### Rule bam2fasta

×

Rule properties

|  |  |
| --- | --- |
| Jobs | 90 |
| Input files | |
| - results/gerp/alignment/GCF\_000283155.1\_CerSimSim1.0\_genomic.Sc9M7eS\_2\_HRSCAF\_41/{gerpref}.bam - results/gerp/alignment/GCF\_000283155.1\_CerSimSim1.0\_genomic.Sc9M7eS\_2\_HRSCAF\_41/{gerpref}.bam.bai - results/gerp/alignment/GCF\_000283155.1\_CerSimSim1.0\_genomic.Sc9M7eS\_2\_HRSCAF\_41/stats/multiqc/multiqc\_report.html - /proj/sllstore2017093/b2016342/b2016342\_nobackup/lts/genome\_erosion\_pipeline/verena\_testing/testdata/reference/gerp/GCF\_000283155.1\_CerSimSim1.0\_genomic.Sc9M7eS\_2\_HRSCAF\_41/split\_bed\_files/{chunk}.bed | |
| Output files | |
| - results/gerp/chunks/GCF\_000283155.1\_CerSimSim1.0\_genomic.Sc9M7eS\_2\_HRSCAF\_41/fasta/{gerpref}\_{chunk,chunk[0-9]+} | |
| Container image | |
| docker://biocontainers/samtools:v1.9-4-deb\_cv1 | |
| Code | |
| ``` 1 2 3 4 5 6 7 8 9 10 11 ``` \| ``` if [ ! -d {output.fasta_dir} ]; then mkdir -p {output.fasta_dir} fi for contig in $(awk -F'	' '{{print $1}}' {input.chunk_bed}) # run the analysis per contig do samtools mpileup -aa -r $contig {input.bam} \| python3 workflow/scripts/filter_mpile.py > {output.fasta_dir}/{params.gerpref}_${{contig}}.mpile 2> {log} && python3 workflow/scripts/sequence_to_fastafile.py {output.fasta_dir}/{params.gerpref}_${{contig}}.mpile $contig {params.gerpref} 2>> {log} && echo "BAM file converted to fasta for" $contig >> {log} done ``` | |

###### Rule align2target

×

Rule properties

|  |  |
| --- | --- |
| Jobs | 30 |
| Input files | |
| - /proj/sllstore2017093/b2016342/b2016342\_nobackup/lts/genome\_erosion\_pipeline/verena\_testing/testdata/reference/GCF\_000283155.1\_CerSimSim1.0\_genomic.Sc9M7eS\_2\_HRSCAF\_41.fasta - /proj/sllstore2017093/b2016342/b2016342\_nobackup/lts/genome\_erosion\_pipeline/verena\_testing/testdata/reference/GCF\_000283155.1\_CerSimSim1.0\_genomic.Sc9M7eS\_2\_HRSCAF\_41.fasta.amb - /proj/sllstore2017093/b2016342/b2016342\_nobackup/lts/genome\_erosion\_pipeline/verena\_testing/testdata/reference/GCF\_000283155.1\_CerSimSim1.0\_genomic.Sc9M7eS\_2\_HRSCAF\_41.fasta.ann - /proj/sllstore2017093/b2016342/b2016342\_nobackup/lts/genome\_erosion\_pipeline/verena\_testing/testdata/reference/GCF\_000283155.1\_CerSimSim1.0\_genomic.Sc9M7eS\_2\_HRSCAF\_41.fasta.bwt - /proj/sllstore2017093/b2016342/b2016342\_nobackup/lts/genome\_erosion\_pipeline/verena\_testing/testdata/reference/GCF\_000283155.1\_CerSimSim1.0\_genomic.Sc9M7eS\_2\_HRSCAF\_41.fasta.pac - /proj/sllstore2017093/b2016342/b2016342\_nobackup/lts/genome\_erosion\_pipeline/verena\_testing/testdata/reference/GCF\_000283155.1\_CerSimSim1.0\_genomic.Sc9M7eS\_2\_HRSCAF\_41.fasta.sa - results/gerp/fastq\_files/{gerpref}.fq.gz - results/gerp/fastq\_files/stats/multiqc/multiqc\_report.html | |
| Output files | |
| - results/gerp/alignment/GCF\_000283155.1\_CerSimSim1.0\_genomic.Sc9M7eS\_2\_HRSCAF\_41/{gerpref}.bam | |
| Container image | |
| docker://nbisweden/generode-bwa:latest | |
| Code | |
| ``` 1 2 ``` \| ``` bwa mem {params.extra} -t {threads} {input.target} {input.fastq} \| samtools view -@ {threads} -h -q 1 -F 4 -F 256 \| grep -v XA:Z \| grep -v SA:Z \| samtools view -@ {threads} -b - \| samtools sort -@ {threads} - > {output.bam} 2> {log} ``` | |

###### Rule outgroups2fastq

×

Rule properties

|  |  |
| --- | --- |
| Jobs | 30 |
| Input files | |
| - /proj/sllstore2017093/b2016342/b2016342\_nobackup/lts/genome\_erosion\_pipeline/verena\_testing/testdata/gerp/outgroup\_Sc9M7eS\_2\_HRSCAF\_41/all\_scaffolds/{gerpref}.fa.gz | |
| Output files | |
| - results/gerp/fastq\_files/{gerpref}.fq.gz | |
| Code | |
| ``` 1 2 ``` \| ``` python3 workflow/scripts/fa2fq.py {input.fasta} {output.fastq} 2> {log} ``` | |

###### Rule outgroup\_fastqc\_multiqc

×

Rule properties

|  |  |
| --- | --- |
| Jobs | 1 |
| Input files | |
| - results/gerp/fastq\_files/stats/Ovis\_aries\_fastqc.html - results/gerp/fastq\_files/stats/Catagonus\_wagneri\_fastqc.html - results/gerp/fastq\_files/stats/Lipotes\_vexillifer\_fastqc.html - results/gerp/fastq\_files/stats/Cervus\_elaphus\_fastqc.html - results/gerp/fastq\_files/stats/Physeter\_catodon\_fastqc.html - results/gerp/fastq\_files/stats/Hyaena\_hyaena\_fastqc.html - results/gerp/fastq\_files/stats/Paradoxurus\_hermaphroditus\_fastqc.html - results/gerp/fastq\_files/stats/Procyon\_lotor\_fastqc.html - results/gerp/fastq\_files/stats/Panthera\_leo\_fastqc.html - results/gerp/fastq\_files/stats/Enhydra\_lutris\_fastqc.html - results/gerp/fastq\_files/stats/Suricata\_suricatta\_fastqc.html - results/gerp/fastq\_files/stats/Ailurus\_fulgens\_fastqc.html - results/gerp/fastq\_files/stats/Antilocapra\_americana\_fastqc.html - results/gerp/fastq\_files/stats/Camelus\_dromedarius\_fastqc.html - results/gerp/fastq\_files/stats/Zalophus\_californianus\_fastqc.html - results/gerp/fastq\_files/stats/Spilogale\_gracilis\_fastqc.html - results/gerp/fastq\_files/stats/Diceros\_bicornis\_fastqc.html - results/gerp/fastq\_files/stats/Balaenoptera\_acutorostrata\_fastqc.html - results/gerp/fastq\_files/stats/Tapirus\_indicus\_fastqc.html - results/gerp/fastq\_files/stats/Ursus\_maritimus\_fastqc.html - results/gerp/fastq\_files/stats/Tragulus\_javanicus\_fastqc.html - results/gerp/fastq\_files/stats/Manis\_javanica\_fastqc.html - results/gerp/fastq\_files/stats/Giraffa\_camelopardalis\_fastqc.html - results/gerp/fastq\_files/stats/Leptonychotes\_weddellii\_fastqc.html - results/gerp/fastq\_files/stats/Equus\_asinus\_fastqc.html - results/gerp/fastq\_files/stats/Mesoplodon\_bidens\_fastqc.html - results/gerp/fastq\_files/stats/Tursiops\_truncatus\_fastqc.html - results/gerp/fastq\_files/stats/Bubalus\_bubalis\_fastqc.html - results/gerp/fastq\_files/stats/Canis\_lupus\_fastqc.html - results/gerp/fastq\_files/stats/Hippopotamus\_amphibius\_fastqc.html - results/gerp/fastq\_files/stats/Ovis\_aries\_fastqc.zip - results/gerp/fastq\_files/stats/Catagonus\_wagneri\_fastqc.zip - results/gerp/fastq\_files/stats/Lipotes\_vexillifer\_fastqc.zip - results/gerp/fastq\_files/stats/Cervus\_elaphus\_fastqc.zip - results/gerp/fastq\_files/stats/Physeter\_catodon\_fastqc.zip - results/gerp/fastq\_files/stats/Hyaena\_hyaena\_fastqc.zip - results/gerp/fastq\_files/stats/Paradoxurus\_hermaphroditus\_fastqc.zip - results/gerp/fastq\_files/stats/Procyon\_lotor\_fastqc.zip - results/gerp/fastq\_files/stats/Panthera\_leo\_fastqc.zip - results/gerp/fastq\_files/stats/Enhydra\_lutris\_fastqc.zip - results/gerp/fastq\_files/stats/Suricata\_suricatta\_fastqc.zip - results/gerp/fastq\_files/stats/Ailurus\_fulgens\_fastqc.zip - results/gerp/fastq\_files/stats/Antilocapra\_americana\_fastqc.zip - results/gerp/fastq\_files/stats/Camelus\_dromedarius\_fastqc.zip - results/gerp/fastq\_files/stats/Zalophus\_californianus\_fastqc.zip - results/gerp/fastq\_files/stats/Spilogale\_gracilis\_fastqc.zip - results/gerp/fastq\_files/stats/Diceros\_bicornis\_fastqc.zip - results/gerp/fastq\_files/stats/Balaenoptera\_acutorostrata\_fastqc.zip - results/gerp/fastq\_files/stats/Tapirus\_indicus\_fastqc.zip - results/gerp/fastq\_files/stats/Ursus\_maritimus\_fastqc.zip - results/gerp/fastq\_files/stats/Tragulus\_javanicus\_fastqc.zip - results/gerp/fastq\_files/stats/Manis\_javanica\_fastqc.zip - results/gerp/fastq\_files/stats/Giraffa\_camelopardalis\_fastqc.zip - results/gerp/fastq\_files/stats/Leptonychotes\_weddellii\_fastqc.zip - results/gerp/fastq\_files/stats/Equus\_asinus\_fastqc.zip - results/gerp/fastq\_files/stats/Mesoplodon\_bidens\_fastqc.zip - results/gerp/fastq\_files/stats/Tursiops\_truncatus\_fastqc.zip - results/gerp/fastq\_files/stats/Bubalus\_bubalis\_fastqc.zip - results/gerp/fastq\_files/stats/Canis\_lupus\_fastqc.zip - results/gerp/fastq\_files/stats/Hippopotamus\_amphibius\_fastqc.zip | |
| Output files | |
| - results/gerp/fastq\_files/stats/multiqc/multiqc\_report.html | |
| Container image | |
| docker://quay.io/biocontainers/multiqc:1.9--pyh9f0ad1d\_0 | |
| Code | |
| ``` 1 2 ``` \| ``` multiqc -f {params.indir} -o {params.outdir} 2> {log} ``` | |

###### Rule outgroup\_fastqc

×

Rule properties

|  |  |
| --- | --- |
| Jobs | 30 |
| Input files | |
| - results/gerp/fastq\_files/{gerpref}.fq.gz | |
| Output files | |
| - results/gerp/fastq\_files/stats/{gerpref}\_fastqc.html - results/gerp/fastq\_files/stats/{gerpref}\_fastqc.zip - results/gerp/fastq\_files/stats/{gerpref}\_fastqc | |
| Container image | |
| docker://biocontainers/fastqc:v0.11.9\_cv7 | |
| Code | |
| ``` 1 2 ``` \| ``` fastqc -o {params.dir} -t {threads} --extract {input.fastq} 2> {log} ``` | |

###### Rule index\_gerp\_bams

×

Rule properties

|  |  |
| --- | --- |
| Jobs | 30 |
| Input files | |
| - results/gerp/alignment/GCF\_000283155.1\_CerSimSim1.0\_genomic.Sc9M7eS\_2\_HRSCAF\_41/{gerpref}.bam | |
| Output files | |
| - results/gerp/alignment/GCF\_000283155.1\_CerSimSim1.0\_genomic.Sc9M7eS\_2\_HRSCAF\_41/{gerpref}.bam.bai | |
| Container image | |
| docker://biocontainers/samtools:v1.9-4-deb\_cv1 | |
| Code | |
| ``` 1 2 ``` \| ``` samtools index {input.bam} {output.index} 2> {log} ``` | |

###### Rule gerp\_bam\_multiqc

×

Rule properties

|  |  |
| --- | --- |
| Jobs | 1 |
| Input files | |
| - results/gerp/alignment/GCF\_000283155.1\_CerSimSim1.0\_genomic.Sc9M7eS\_2\_HRSCAF\_41/stats/Ovis\_aries.bam.stats.txt - results/gerp/alignment/GCF\_000283155.1\_CerSimSim1.0\_genomic.Sc9M7eS\_2\_HRSCAF\_41/stats/Catagonus\_wagneri.bam.stats.txt - results/gerp/alignment/GCF\_000283155.1\_CerSimSim1.0\_genomic.Sc9M7eS\_2\_HRSCAF\_41/stats/Lipotes\_vexillifer.bam.stats.txt - results/gerp/alignment/GCF\_000283155.1\_CerSimSim1.0\_genomic.Sc9M7eS\_2\_HRSCAF\_41/stats/Cervus\_elaphus.bam.stats.txt - results/gerp/alignment/GCF\_000283155.1\_CerSimSim1.0\_genomic.Sc9M7eS\_2\_HRSCAF\_41/stats/Physeter\_catodon.bam.stats.txt - results/gerp/alignment/GCF\_000283155.1\_CerSimSim1.0\_genomic.Sc9M7eS\_2\_HRSCAF\_41/stats/Hyaena\_hyaena.bam.stats.txt - results/gerp/alignment/GCF\_000283155.1\_CerSimSim1.0\_genomic.Sc9M7eS\_2\_HRSCAF\_41/stats/Paradoxurus\_hermaphroditus.bam.stats.txt - results/gerp/alignment/GCF\_000283155.1\_CerSimSim1.0\_genomic.Sc9M7eS\_2\_HRSCAF\_41/stats/Procyon\_lotor.bam.stats.txt - results/gerp/alignment/GCF\_000283155.1\_CerSimSim1.0\_genomic.Sc9M7eS\_2\_HRSCAF\_41/stats/Panthera\_leo.bam.stats.txt - results/gerp/alignment/GCF\_000283155.1\_CerSimSim1.0\_genomic.Sc9M7eS\_2\_HRSCAF\_41/stats/Enhydra\_lutris.bam.stats.txt - results/gerp/alignment/GCF\_000283155.1\_CerSimSim1.0\_genomic.Sc9M7eS\_2\_HRSCAF\_41/stats/Suricata\_suricatta.bam.stats.txt - results/gerp/alignment/GCF\_000283155.1\_CerSimSim1.0\_genomic.Sc9M7eS\_2\_HRSCAF\_41/stats/Ailurus\_fulgens.bam.stats.txt - results/gerp/alignment/GCF\_000283155.1\_CerSimSim1.0\_genomic.Sc9M7eS\_2\_HRSCAF\_41/stats/Antilocapra\_americana.bam.stats.txt - results/gerp/alignment/GCF\_000283155.1\_CerSimSim1.0\_genomic.Sc9M7eS\_2\_HRSCAF\_41/stats/Camelus\_dromedarius.bam.stats.txt - results/gerp/alignment/GCF\_000283155.1\_CerSimSim1.0\_genomic.Sc9M7eS\_2\_HRSCAF\_41/stats/Zalophus\_californianus.bam.stats.txt - results/gerp/alignment/GCF\_000283155.1\_CerSimSim1.0\_genomic.Sc9M7eS\_2\_HRSCAF\_41/stats/Spilogale\_gracilis.bam.stats.txt - results/gerp/alignment/GCF\_000283155.1\_CerSimSim1.0\_genomic.Sc9M7eS\_2\_HRSCAF\_41/stats/Diceros\_bicornis.bam.stats.txt - results/gerp/alignment/GCF\_000283155.1\_CerSimSim1.0\_genomic.Sc9M7eS\_2\_HRSCAF\_41/stats/Balaenoptera\_acutorostrata.bam.stats.txt - results/gerp/alignment/GCF\_000283155.1\_CerSimSim1.0\_genomic.Sc9M7eS\_2\_HRSCAF\_41/stats/Tapirus\_indicus.bam.stats.txt - results/gerp/alignment/GCF\_000283155.1\_CerSimSim1.0\_genomic.Sc9M7eS\_2\_HRSCAF\_41/stats/Ursus\_maritimus.bam.stats.txt - results/gerp/alignment/GCF\_000283155.1\_CerSimSim1.0\_genomic.Sc9M7eS\_2\_HRSCAF\_41/stats/Tragulus\_javanicus.bam.stats.txt - results/gerp/alignment/GCF\_000283155.1\_CerSimSim1.0\_genomic.Sc9M7eS\_2\_HRSCAF\_41/stats/Manis\_javanica.bam.stats.txt - results/gerp/alignment/GCF\_000283155.1\_CerSimSim1.0\_genomic.Sc9M7eS\_2\_HRSCAF\_41/stats/Giraffa\_camelopardalis.bam.stats.txt - results/gerp/alignment/GCF\_000283155.1\_CerSimSim1.0\_genomic.Sc9M7eS\_2\_HRSCAF\_41/stats/Leptonychotes\_weddellii.bam.stats.txt - results/gerp/alignment/GCF\_000283155.1\_CerSimSim1.0\_genomic.Sc9M7eS\_2\_HRSCAF\_41/stats/Equus\_asinus.bam.stats.txt - results/gerp/alignment/GCF\_000283155.1\_CerSimSim1.0\_genomic.Sc9M7eS\_2\_HRSCAF\_41/stats/Mesoplodon\_bidens.bam.stats.txt - results/gerp/alignment/GCF\_000283155.1\_CerSimSim1.0\_genomic.Sc9M7eS\_2\_HRSCAF\_41/stats/Tursiops\_truncatus.bam.stats.txt - results/gerp/alignment/GCF\_000283155.1\_CerSimSim1.0\_genomic.Sc9M7eS\_2\_HRSCAF\_41/stats/Bubalus\_bubalis.bam.stats.txt - results/gerp/alignment/GCF\_000283155.1\_CerSimSim1.0\_genomic.Sc9M7eS\_2\_HRSCAF\_41/stats/Canis\_lupus.bam.stats.txt - results/gerp/alignment/GCF\_000283155.1\_CerSimSim1.0\_genomic.Sc9M7eS\_2\_HRSCAF\_41/stats/Hippopotamus\_amphibius.bam.stats.txt | |
| Output files | |
| - results/gerp/alignment/GCF\_000283155.1\_CerSimSim1.0\_genomic.Sc9M7eS\_2\_HRSCAF\_41/stats/multiqc/multiqc\_report.html | |
| Container image | |
| docker://quay.io/biocontainers/multiqc:1.9--pyh9f0ad1d\_0 | |
| Code | |
| ``` 1 2 ``` \| ``` multiqc -f {params.indir} -o {params.outdir} 2> {log} ``` | |

###### Rule gerp\_bam\_stats

×

Rule properties

|  |  |
| --- | --- |
| Jobs | 30 |
| Input files | |
| - results/gerp/alignment/GCF\_000283155.1\_CerSimSim1.0\_genomic.Sc9M7eS\_2\_HRSCAF\_41/{gerpref}.bam - results/gerp/alignment/GCF\_000283155.1\_CerSimSim1.0\_genomic.Sc9M7eS\_2\_HRSCAF\_41/{gerpref}.bam.bai | |
| Output files | |
| - results/gerp/alignment/GCF\_000283155.1\_CerSimSim1.0\_genomic.Sc9M7eS\_2\_HRSCAF\_41/stats/{gerpref}.bam.stats.txt | |
| Container image | |
| docker://biocontainers/samtools:v1.9-4-deb\_cv1 | |
| Code | |
| ``` 1 2 ``` \| ``` samtools flagstat {input.bam} > {output.stats} 2> {log} ``` | |

###### Rule split\_ref\_contigs

×

Rule properties

|  |  |
| --- | --- |
| Jobs | 3 |
| Input files | |
| - /proj/sllstore2017093/b2016342/b2016342\_nobackup/lts/genome\_erosion\_pipeline/verena\_testing/testdata/reference/GCF\_000283155.1\_CerSimSim1.0\_genomic.Sc9M7eS\_2\_HRSCAF\_41.fasta - /proj/sllstore2017093/b2016342/b2016342\_nobackup/lts/genome\_erosion\_pipeline/verena\_testing/testdata/reference/GCF\_000283155.1\_CerSimSim1.0\_genomic.Sc9M7eS\_2\_HRSCAF\_41.fasta.fai - /proj/sllstore2017093/b2016342/b2016342\_nobackup/lts/genome\_erosion\_pipeline/verena\_testing/testdata/reference/gerp/GCF\_000283155.1\_CerSimSim1.0\_genomic.Sc9M7eS\_2\_HRSCAF\_41/split\_bed\_files/{chunk}.bed - results/gerp/alignment/GCF\_000283155.1\_CerSimSim1.0\_genomic.Sc9M7eS\_2\_HRSCAF\_41/stats/multiqc/multiqc\_report.html | |
| Output files | |
| - results/gerp/chunks/GCF\_000283155.1\_CerSimSim1.0\_genomic.Sc9M7eS\_2\_HRSCAF\_41/fasta/GCF\_000283155.1\_CerSimSim1.0\_genomic.Sc9M7eS\_2\_HRSCAF\_41\_{chunk,chunk[0-9]+} | |
| Container image | |
| docker://quay.io/biocontainers/seqtk:1.3--hed695b0\_2 | |
| Code | |
| ``` 1 2 3 4 5 6 7 8 9 10 11 ``` \| ``` if [ ! -d {output.fasta_dir} ]; then mkdir -p {output.fasta_dir}; fi for contig in $(awk -F'	' '{{print $1}}' {input.chunk_bed}) # run the analysis per contig do echo $contig > {output.fasta_dir}/${{contig}}.lst && seqtk subseq {input.ref} {output.fasta_dir}/${{contig}}.lst \| sed "s/$contig/{params.gerpref}/g" \| seqtk seq > {output.fasta_dir}/{params.gerpref}_${{contig}}.fasta 2> {log} && echo $contig "extracted from reference" >> {log} done ``` | |

###### Rule rescale\_gerp

×

Rule properties

|  |  |
| --- | --- |
| Jobs | 3 |
| Input files | |
| - results/gerp/chunks/GCF\_000283155.1\_CerSimSim1.0\_genomic.Sc9M7eS\_2\_HRSCAF\_41/gerp/{chunk}\_gerp\_coords - /proj/sllstore2017093/b2016342/b2016342\_nobackup/lts/genome\_erosion\_pipeline/verena\_testing/testdata/reference/gerp/GCF\_000283155.1\_CerSimSim1.0\_genomic.Sc9M7eS\_2\_HRSCAF\_41/split\_bed\_files/{chunk}.bed | |
| Output files | |
| - results/gerp/chunks/GCF\_000283155.1\_CerSimSim1.0\_genomic.Sc9M7eS\_2\_HRSCAF\_41/gerp/{chunk,chunk[0-9]+}\_gerp\_rescaled | |
| Code | |
| ``` 1 2 3 4 5 6 7 8 9 ``` \| ``` if [ ! -d {output.gerp_rescaled_dir} ]; then mkdir -p {output.gerp_rescaled_dir}; fi for contig in $(awk -F'	' '{{print $1}}' {input.chunk_bed}) # run the analysis per contig do awk -F'	' '{{ if($1 ~ /[0-9]+/ && $1 != 0) {{print $1/1000}} else {{print $1}} }}' OFS='	' {input.gerp_coords_dir}/${{contig}}.fasta.rates.parsed > {output.gerp_rescaled_dir}/${{contig}}.fasta.rates.parsed.rescaled 2>> {log} && echo "GERP scores rescaled for" $contig >> {log} done ``` | |

###### Rule gerp2coords

×

Rule properties

|  |  |
| --- | --- |
| Jobs | 3 |
| Input files | |
| - results/gerp/chunks/GCF\_000283155.1\_CerSimSim1.0\_genomic.Sc9M7eS\_2\_HRSCAF\_41/fasta/concatenated\_{chunk} - results/gerp/chunks/GCF\_000283155.1\_CerSimSim1.0\_genomic.Sc9M7eS\_2\_HRSCAF\_41/gerp/{chunk}\_gerp\_raw - /proj/sllstore2017093/b2016342/b2016342\_nobackup/lts/genome\_erosion\_pipeline/verena\_testing/testdata/reference/gerp/GCF\_000283155.1\_CerSimSim1.0\_genomic.Sc9M7eS\_2\_HRSCAF\_41/split\_bed\_files/{chunk}.bed | |
| Output files | |
| - results/gerp/chunks/GCF\_000283155.1\_CerSimSim1.0\_genomic.Sc9M7eS\_2\_HRSCAF\_41/gerp/{chunk,chunk[0-9]+}\_gerp\_coords | |

###### Rule compute\_gerp

×

Rule properties

|  |  |
| --- | --- |
| Jobs | 3 |
| Input files | |
| - results/gerp/chunks/GCF\_000283155.1\_CerSimSim1.0\_genomic.Sc9M7eS\_2\_HRSCAF\_41/fasta/concatenated\_{chunk} - /proj/sllstore2017093/b2016342/b2016342\_nobackup/lts/genome\_erosion\_pipeline/verena\_testing/testdata/reference/gerp/GCF\_000283155.1\_CerSimSim1.0\_genomic.Sc9M7eS\_2\_HRSCAF\_41/split\_bed\_files/{chunk}.bed - /proj/sllstore2017093/b2016342/b2016342\_nobackup/lts/genome\_erosion\_pipeline/verena\_testing/testdata/scripts/testdataset\_paper/Sc9M7eS\_2\_HRSCAF\_41/gerp/gerp\_30\_outgroups\_plus\_whi\_rhino\_names.nwk | |
| Output files | |
| - results/gerp/chunks/GCF\_000283155.1\_CerSimSim1.0\_genomic.Sc9M7eS\_2\_HRSCAF\_41/gerp/{chunk,chunk[0-9]+}\_gerp\_raw | |
| Container image | |
| docker://quay.io/biocontainers/gerp:2.1--hfc679d8\_0 | |
| Code | |
| ``` 1 2 3 4 5 6 7 8 9 10 ``` \| ``` if [ ! -d {output.gerp_dir} ]; then mkdir -p {output.gerp_dir}; fi for contig in $(awk -F'	' '{{print $1}}' {input.chunk_bed}) # run the analysis per contig do gerpcol -v -f {input.concatenated_fasta_dir}/${{contig}}.fasta -t {input.tree} -a -e {params.name} 2> {log} && mv {input.concatenated_fasta_dir}/${{contig}}.fasta.rates {output.gerp_dir} 2>> {log} && echo "Computed GERP++ scores for" $contig >> {log} done ``` | |

###### Rule historical\_biallelic\_missing\_filtered\_vcf\_gerp\_multiqc

×

Rule properties

|  |  |
| --- | --- |
| Jobs | 1 |
| Input files | |
| Output files | |
| - results/gerp/historical/GCF\_000283155.1\_CerSimSim1.0\_genomic.Sc9M7eS\_2\_HRSCAF\_41/vcf/stats/multiqc/multiqc\_report.html | |
| Container image | |
| docker://quay.io/biocontainers/multiqc:1.9--pyh9f0ad1d\_0 | |
| Code | |
| ``` 1 2 ``` \| ``` multiqc -f {params.indir} -o {params.outdir} 2> {log} ``` | |

###### Rule biallelic\_missing\_filtered\_vcf\_gerp\_stats

×

Rule properties

|  |  |
| --- | --- |
| Jobs | 6 |
| Input files | |
| - results/gerp/{dataset}/GCF\_000283155.1\_CerSimSim1.0\_genomic.Sc9M7eS\_2\_HRSCAF\_41/vcf/{sample}.merged.rmdup.merged.{processed}.snps5.noIndel.QUAL30.dp.AB.repma.biallelic.fmissing{fmiss}.vcf | |
| Output files | |
| - results/gerp/{dataset}/GCF\_000283155.1\_CerSimSim1.0\_genomic.Sc9M7eS\_2\_HRSCAF\_41/vcf/stats/{sample,[A-Za-z0-9]+}.merged.rmdup.merged.{processed}.snps5.noIndel.QUAL30.dp.AB.repma.biallelic.fmissing{fmiss}.vcf.stats.txt | |
| Container image | |
| docker://quay.io/biocontainers/bcftools:1.9--h68d8f2e\_9 | |
| Code | |
| ``` 1 2 ``` \| ``` bcftools stats {input.filtered} > {output.stats} 2> {log} ``` | |

###### Rule filter\_biallelic\_missing\_vcf\_gerp

×

Rule properties

|  |  |
| --- | --- |
| Jobs | 6 |
| Input files | |
| - results/gerp/chunks/GCF\_000283155.1\_CerSimSim1.0\_genomic.Sc9M7eS\_2\_HRSCAF\_41/{dataset}/vcf/{sample}.merged.rmdup.merged.{processed}.snps5.noIndel.QUAL30.dp.AB.repma.vcf.gz - results/all/vcf/GCF\_000283155.1\_CerSimSim1.0\_genomic.Sc9M7eS\_2\_HRSCAF\_41.all.merged.biallelic.fmissing{fmiss}.bed - /proj/sllstore2017093/b2016342/b2016342\_nobackup/lts/genome\_erosion\_pipeline/verena\_testing/testdata/reference/GCF\_000283155.1\_CerSimSim1.0\_genomic.Sc9M7eS\_2\_HRSCAF\_41.genome | |
| Output files | |
| - results/gerp/{dataset}/GCF\_000283155.1\_CerSimSim1.0\_genomic.Sc9M7eS\_2\_HRSCAF\_41/vcf/{sample,[A-Za-z0-9]+}.merged.rmdup.merged.{processed}.snps5.noIndel.QUAL30.dp.AB.repma.biallelic.fmissing{fmiss}.vcf | |
| Container image | |
| docker://quay.io/biocontainers/bedtools:2.29.2--hc088bd4\_0 | |
| Code | |
| ``` 1 2 ``` \| ``` bedtools intersect -a {input.vcf} -b {input.bed} -header -sorted -g {input.genomefile} > {output.filtered} 2> {log} ``` | |

###### Rule repmasked\_bcf2vcf\_gerp

×

Rule properties

|  |  |
| --- | --- |
| Jobs | 6 |
| Input files | |
| - results/{dataset}/vcf/GCF\_000283155.1\_CerSimSim1.0\_genomic.Sc9M7eS\_2\_HRSCAF\_41/{sample}.merged.rmdup.merged.{processed}.snps5.noIndel.QUAL30.dp.AB.repma.bcf - results/{dataset}/vcf/GCF\_000283155.1\_CerSimSim1.0\_genomic.Sc9M7eS\_2\_HRSCAF\_41/{sample}.merged.rmdup.merged.{processed}.snps5.noIndel.QUAL30.dp.AB.repma.bcf.csi | |
| Output files | |
| - results/gerp/chunks/GCF\_000283155.1\_CerSimSim1.0\_genomic.Sc9M7eS\_2\_HRSCAF\_41/{dataset}/vcf/{sample,[A-Za-z0-9]+}.merged.rmdup.merged.{processed}.snps5.noIndel.QUAL30.dp.AB.repma.vcf.gz | |
| Container image | |
| docker://quay.io/biocontainers/bcftools:1.9--h68d8f2e\_9 | |
| Code | |
| ``` 1 2 ``` \| ``` bcftools convert -O z -o {output.vcf} {input.bcf} 2> {log} ``` | |

###### Rule modern\_biallelic\_missing\_filtered\_vcf\_gerp\_multiqc

×

Rule properties

|  |  |
| --- | --- |
| Jobs | 1 |
| Input files | |
| Output files | |
| - results/gerp/modern/GCF\_000283155.1\_CerSimSim1.0\_genomic.Sc9M7eS\_2\_HRSCAF\_41/vcf/stats/multiqc/multiqc\_report.html | |
| Container image | |
| docker://quay.io/biocontainers/multiqc:1.9--pyh9f0ad1d\_0 | |
| Code | |
| ``` 1 2 ``` \| ``` multiqc -f {params.indir} -o {params.outdir} 2> {log} ``` | |

###### Rule relative\_mutational\_load\_plot

×

Rule properties

|  |  |
| --- | --- |
| Jobs | 1 |
| Input files | |
| Output files | |
| - results/gerp/{dataset}/GCF\_000283155.1\_CerSimSim1.0\_genomic.Sc9M7eS\_2\_HRSCAF\_41.{dataset}.fmissing{fmiss}.relative\_mutational\_load.gerp\_{minGERP}\_{maxGERP}\_plot.pdf | |

###### Rule relative\_mutational\_load\_table

×

Rule properties

|  |  |
| --- | --- |
| Jobs | 1 |
| Input files | |
| Output files | |
| - results/gerp/{dataset}/GCF\_000283155.1\_CerSimSim1.0\_genomic.Sc9M7eS\_2\_HRSCAF\_41.{dataset}.fmissing{fmiss}.relative\_mutational\_load.gerp\_{minGERP}\_{maxGERP}\_table.txt | |

###### Rule relative\_mutational\_load\_per\_sample

×

Rule properties

|  |  |
| --- | --- |
| Jobs | 6 |
| Input files | |
| - results/gerp/{dataset}/GCF\_000283155.1\_CerSimSim1.0\_genomic.Sc9M7eS\_2\_HRSCAF\_41/{sample}.merged.rmdup.merged.{processed}.snps5.noIndel.QUAL30.dp.AB.repma.biallelic.fmissing{fmiss}.ancestral.rates.derived.alleles.gz | |
| Output files | |
| - results/gerp/{dataset}/GCF\_000283155.1\_CerSimSim1.0\_genomic.Sc9M7eS\_2\_HRSCAF\_41/{sample,[A-Za-z0-9]+}.merged.rmdup.merged.{processed}.snps5.noIndel.QUAL30.dp.AB.repma.biallelic.fmissing{fmiss}.relative\_mutational\_load.gerp\_{minGERP}\_{maxGERP}\_table.txt | |
| Code | |
| ``` 1 2 ``` \| ``` python3 workflow/scripts/gerp_rel_mut_load_sample.py {input.gerp_out} {params.min_gerp} {params.max_gerp} {output.mut_load} 2> {log} ``` | |

###### Rule merge\_gerp\_alleles\_gz

×

Rule properties

|  |  |
| --- | --- |
| Jobs | 6 |
| Input files | |
| - results/gerp/chunks/GCF\_000283155.1\_CerSimSim1.0\_genomic.Sc9M7eS\_2\_HRSCAF\_41/{dataset}/{sample}.merged.rmdup.merged.{processed}.snps5.noIndel.QUAL30.dp.AB.repma.biallelic.fmissing0.1.chunk1.fasta.parsed.rates.derived\_alleles - results/gerp/chunks/GCF\_000283155.1\_CerSimSim1.0\_genomic.Sc9M7eS\_2\_HRSCAF\_41/{dataset}/{sample}.merged.rmdup.merged.{processed}.snps5.noIndel.QUAL30.dp.AB.repma.biallelic.fmissing0.1.chunk2.fasta.parsed.rates.derived\_alleles - results/gerp/chunks/GCF\_000283155.1\_CerSimSim1.0\_genomic.Sc9M7eS\_2\_HRSCAF\_41/{dataset}/{sample}.merged.rmdup.merged.{processed}.snps5.noIndel.QUAL30.dp.AB.repma.biallelic.fmissing0.1.chunk3.fasta.parsed.rates.derived\_alleles | |
| Output files | |
| - results/gerp/{dataset}/GCF\_000283155.1\_CerSimSim1.0\_genomic.Sc9M7eS\_2\_HRSCAF\_41/{sample,[A-Za-z0-9]+}.merged.rmdup.merged.{processed}.snps5.noIndel.QUAL30.dp.AB.repma.biallelic.fmissing{fmiss}.ancestral.rates.derived.alleles.gz | |
| Code | |
| ``` 1 2 ``` \| ``` awk 'FNR>1 \|\| NR==1' {input.gerp_chunks_merged} \| gzip - > {output.gerp_out} 2> {log} ``` | |

###### Rule merge\_gerp\_alleles\_per\_chunk

×

Rule properties

|  |  |
| --- | --- |
| Jobs | 18 |
| Input files | |
| - results/gerp/chunks/GCF\_000283155.1\_CerSimSim1.0\_genomic.Sc9M7eS\_2\_HRSCAF\_41/{dataset}/{sample}.merged.rmdup.merged.{processed}.snps5.noIndel.QUAL30.dp.AB.repma.biallelic.fmissing{fmiss}.{chunk}\_gerp\_derived\_alleles - /proj/sllstore2017093/b2016342/b2016342\_nobackup/lts/genome\_erosion\_pipeline/verena\_testing/testdata/reference/gerp/GCF\_000283155.1\_CerSimSim1.0\_genomic.Sc9M7eS\_2\_HRSCAF\_41/split\_bed\_files/windows/{chunk}\_10Mwindows.bed | |
| Output files | |
| - results/gerp/chunks/GCF\_000283155.1\_CerSimSim1.0\_genomic.Sc9M7eS\_2\_HRSCAF\_41/{dataset}/{sample,[A-Za-z0-9]+}.merged.rmdup.merged.{processed}.snps5.noIndel.QUAL30.dp.AB.repma.biallelic.fmissing{fmiss}.{chunk,chunk[0-9]+}.fasta.parsed.rates.derived\_alleles | |

###### Rule gerp\_derived\_alleles

×

Rule properties

|  |  |
| --- | --- |
| Jobs | 18 |
| Input files | |
| - results/gerp/chunks/GCF\_000283155.1\_CerSimSim1.0\_genomic.Sc9M7eS\_2\_HRSCAF\_41/gerp/{chunk}.fasta.parsed.rates - results/gerp/chunks/GCF\_000283155.1\_CerSimSim1.0\_genomic.Sc9M7eS\_2\_HRSCAF\_41/gerp/{chunk}\_gerp\_merged - results/gerp/chunks/GCF\_000283155.1\_CerSimSim1.0\_genomic.Sc9M7eS\_2\_HRSCAF\_41/{dataset}/vcf/{sample}.merged.rmdup.merged.{processed}.snps5.noIndel.QUAL30.dp.AB.repma.biallelic.fmissing{fmiss}.{chunk}.vcf.gz - /proj/sllstore2017093/b2016342/b2016342\_nobackup/lts/genome\_erosion\_pipeline/verena\_testing/testdata/reference/gerp/GCF\_000283155.1\_CerSimSim1.0\_genomic.Sc9M7eS\_2\_HRSCAF\_41/split\_bed\_files/windows/{chunk}\_10Mwindows.bed | |
| Output files | |
| - results/gerp/chunks/GCF\_000283155.1\_CerSimSim1.0\_genomic.Sc9M7eS\_2\_HRSCAF\_41/{dataset}/{sample,[A-Za-z0-9]+}.merged.rmdup.merged.{processed}.snps5.noIndel.QUAL30.dp.AB.repma.biallelic.fmissing{fmiss}.{chunk,chunk[0-9]+}\_gerp\_derived\_alleles | |
| Code | |
| ``` 1 2 3 4 5 6 7 ``` \| ``` if [ ! -d {output.gerp_alleles_dir} ]; then mkdir -p {output.gerp_alleles_dir}; fi while IFS="	" read -r contig start end; do python3 workflow/scripts/gerp_derived_alleles.py {input.gerp_merged_dir}/${{contig}}.fasta.parsed.rates {input.vcf} ${{contig}} ${{start}} ${{end}} {output.gerp_alleles_dir}/${{contig}}_${{start}}_${{end}}.fasta.parsed.rates.derived_alleles; done < {input.chunk_win_bed} 2>> {log} ``` | |

###### Rule split\_vcf\_files

×

Rule properties

|  |  |
| --- | --- |
| Jobs | 18 |
| Input files | |
| - results/gerp/{dataset}/GCF\_000283155.1\_CerSimSim1.0\_genomic.Sc9M7eS\_2\_HRSCAF\_41/vcf/{sample}.merged.rmdup.merged.{processed}.snps5.noIndel.QUAL30.dp.AB.repma.biallelic.fmissing{fmiss}.vcf - /proj/sllstore2017093/b2016342/b2016342\_nobackup/lts/genome\_erosion\_pipeline/verena\_testing/testdata/reference/gerp/GCF\_000283155.1\_CerSimSim1.0\_genomic.Sc9M7eS\_2\_HRSCAF\_41/split\_bed\_files/{chunk}.bed - /proj/sllstore2017093/b2016342/b2016342\_nobackup/lts/genome\_erosion\_pipeline/verena\_testing/testdata/reference/GCF\_000283155.1\_CerSimSim1.0\_genomic.Sc9M7eS\_2\_HRSCAF\_41.genome | |
| Output files | |
| - results/gerp/chunks/GCF\_000283155.1\_CerSimSim1.0\_genomic.Sc9M7eS\_2\_HRSCAF\_41/{dataset}/vcf/{sample,[A-Za-z0-9]+}.merged.rmdup.merged.{processed}.snps5.noIndel.QUAL30.dp.AB.repma.biallelic.fmissing{fmiss}.{chunk,chunk[0-9]+}.vcf.gz | |
| Container image | |
| docker://quay.io/biocontainers/bedtools:2.29.2--hc088bd4\_0 | |
| Code | |
| ``` 1 2 ``` \| ``` bedtools intersect -a {input.vcf} -b {input.chunk_bed} -g {input.genomefile} -header \| gzip - > {output.vcf_chunk} 2> {log} ``` | |

###### Rule split\_chunk\_bed\_files

×

Rule properties

|  |  |
| --- | --- |
| Jobs | 3 |
| Input files | |
| - /proj/sllstore2017093/b2016342/b2016342\_nobackup/lts/genome\_erosion\_pipeline/verena\_testing/testdata/reference/gerp/GCF\_000283155.1\_CerSimSim1.0\_genomic.Sc9M7eS\_2\_HRSCAF\_41/split\_bed\_files/{chunk}.bed | |
| Output files | |
| - /proj/sllstore2017093/b2016342/b2016342\_nobackup/lts/genome\_erosion\_pipeline/verena\_testing/testdata/reference/gerp/GCF\_000283155.1\_CerSimSim1.0\_genomic.Sc9M7eS\_2\_HRSCAF\_41/split\_bed\_files/windows/{chunk,chunk[0-9]+}\_10Mwindows.bed | |
| Container image | |
| docker://quay.io/biocontainers/bedtools:2.29.2--hc088bd4\_0 | |
| Code | |
| ``` 1 2 ``` \| ``` bedtools makewindows -b {input.chunk_bed} -w 10000000 > {output.chunk_win_bed} 2> {log} ``` | |
