## Additional file 3 for "GenErode: a bioinformatics pipeline to investigate genome erosion in endangered and extinct species": Additional_file_3_v3.html

Snakemake Report

Loading Snakemake Report...

Please enable Javascript in your browser to see this report.

Loading 2.6 MB. For large reports, this can take a while.

Snakemake Report

- Wed Jan 19 11:52:16 2022 CET
- Snakemake 6.12.1

Loading...

×

Download original

##### Rule bwa\_index\_reference

×

Rule properties

|  |  |
| --- | --- |
| Jobs | 1 |
| Input files | |
| - /proj/sllstore2017093/b2016342/b2016342\_nobackup/lts/genome\_erosion\_pipeline/verena\_testing/testdata/reference/sumatran\_rhino\_22Jul2017\_9M7eS\_haploidified\_headersFixed\_Sc9M7eS\_2\_HRSCAF\_41.fasta | |
| Output files | |
| - /proj/sllstore2017093/b2016342/b2016342\_nobackup/lts/genome\_erosion\_pipeline/verena\_testing/testdata/reference/sumatran\_rhino\_22Jul2017\_9M7eS\_haploidified\_headersFixed\_Sc9M7eS\_2\_HRSCAF\_41.fasta.amb - /proj/sllstore2017093/b2016342/b2016342\_nobackup/lts/genome\_erosion\_pipeline/verena\_testing/testdata/reference/sumatran\_rhino\_22Jul2017\_9M7eS\_haploidified\_headersFixed\_Sc9M7eS\_2\_HRSCAF\_41.fasta.ann - /proj/sllstore2017093/b2016342/b2016342\_nobackup/lts/genome\_erosion\_pipeline/verena\_testing/testdata/reference/sumatran\_rhino\_22Jul2017\_9M7eS\_haploidified\_headersFixed\_Sc9M7eS\_2\_HRSCAF\_41.fasta.bwt - /proj/sllstore2017093/b2016342/b2016342\_nobackup/lts/genome\_erosion\_pipeline/verena\_testing/testdata/reference/sumatran\_rhino\_22Jul2017\_9M7eS\_haploidified\_headersFixed\_Sc9M7eS\_2\_HRSCAF\_41.fasta.pac - /proj/sllstore2017093/b2016342/b2016342\_nobackup/lts/genome\_erosion\_pipeline/verena\_testing/testdata/reference/sumatran\_rhino\_22Jul2017\_9M7eS\_haploidified\_headersFixed\_Sc9M7eS\_2\_HRSCAF\_41.fasta.sa | |
| Container image | |
| docker://biocontainers/bwa:v0.7.17-3-deb\_cv1 | |
| Code | |
| ``` 1 2 ``` \| ``` bwa index -a bwtsw {input.ref} 2> {log} ``` | |

##### Rule samtools\_fasta\_index

×

Rule properties

|  |  |
| --- | --- |
| Jobs | 1 |
| Input files | |
| - /proj/sllstore2017093/b2016342/b2016342\_nobackup/lts/genome\_erosion\_pipeline/verena\_testing/testdata/reference/sumatran\_rhino\_22Jul2017\_9M7eS\_haploidified\_headersFixed\_Sc9M7eS\_2\_HRSCAF\_41.fasta | |
| Output files | |
| - /proj/sllstore2017093/b2016342/b2016342\_nobackup/lts/genome\_erosion\_pipeline/verena\_testing/testdata/reference/sumatran\_rhino\_22Jul2017\_9M7eS\_haploidified\_headersFixed\_Sc9M7eS\_2\_HRSCAF\_41.fasta.fai | |
| Container image | |
| docker://biocontainers/samtools:v1.9-4-deb\_cv1 | |
| Code | |
| ``` 1 2 ``` \| ``` samtools faidx {input.ref} 2> {log} ``` | |

##### Rule picard\_fasta\_dict

×

Rule properties

|  |  |
| --- | --- |
| Jobs | 1 |
| Input files | |
| - /proj/sllstore2017093/b2016342/b2016342\_nobackup/lts/genome\_erosion\_pipeline/verena\_testing/testdata/reference/sumatran\_rhino\_22Jul2017\_9M7eS\_haploidified\_headersFixed\_Sc9M7eS\_2\_HRSCAF\_41.fasta | |
| Output files | |
| - /proj/sllstore2017093/b2016342/b2016342\_nobackup/lts/genome\_erosion\_pipeline/verena\_testing/testdata/reference/sumatran\_rhino\_22Jul2017\_9M7eS\_haploidified\_headersFixed\_Sc9M7eS\_2\_HRSCAF\_41.dict | |
| Container image | |
| docker://quay.io/biocontainers/picard:2.26.6--hdfd78af\_0 | |
| Code | |
| ``` 1 2 ``` \| ``` picard CreateSequenceDictionary -Xmx{params.mem} R={input.ref} O={output.fdict} 2> {log} ``` | |

##### Rule genome\_file

×

Rule properties

|  |  |
| --- | --- |
| Jobs | 1 |
| Input files | |
| - /proj/sllstore2017093/b2016342/b2016342\_nobackup/lts/genome\_erosion\_pipeline/verena\_testing/testdata/reference/sumatran\_rhino\_22Jul2017\_9M7eS\_haploidified\_headersFixed\_Sc9M7eS\_2\_HRSCAF\_41.fasta.fai | |
| Output files | |
| - /proj/sllstore2017093/b2016342/b2016342\_nobackup/lts/genome\_erosion\_pipeline/verena\_testing/testdata/reference/sumatran\_rhino\_22Jul2017\_9M7eS\_haploidified\_headersFixed\_Sc9M7eS\_2\_HRSCAF\_41.genome | |
| Code | |
| ``` 1 2 ``` \| ``` awk -v OFS='	' '{{print $1, $2}}' {input.fai} > {output.genomefile} 2> {log} ``` | |

##### Rule make\_reference\_bed

×

Rule properties

|  |  |
| --- | --- |
| Jobs | 1 |
| Input files | |
| - /proj/sllstore2017093/b2016342/b2016342\_nobackup/lts/genome\_erosion\_pipeline/verena\_testing/testdata/reference/sumatran\_rhino\_22Jul2017\_9M7eS\_haploidified\_headersFixed\_Sc9M7eS\_2\_HRSCAF\_41.fasta.fai | |
| Output files | |
| - /proj/sllstore2017093/b2016342/b2016342\_nobackup/lts/genome\_erosion\_pipeline/verena\_testing/testdata/reference/sumatran\_rhino\_22Jul2017\_9M7eS\_haploidified\_headersFixed\_Sc9M7eS\_2\_HRSCAF\_41.bed | |
| Code | |
| ``` 1 2 ``` \| ``` awk -v OFS='	' '{{print $1, "0", $2}}' {input.fai} > {output.ref_bed} 2> {log} ``` | |

##### Rule make\_repeats\_bed

×

Rule properties

|  |  |
| --- | --- |
| Jobs | 1 |
| Input files | |
| - /proj/sllstore2017093/b2016342/b2016342\_nobackup/lts/genome\_erosion\_pipeline/verena\_testing/testdata/reference/repeatmasker/sumatran\_rhino\_22Jul2017\_9M7eS\_haploidified\_headersFixed\_Sc9M7eS\_2\_HRSCAF\_41/sumatran\_rhino\_22Jul2017\_9M7eS\_haploidified\_headersFixed\_Sc9M7eS\_2\_HRSCAF\_41.upper.fasta.out | |
| Output files | |
| - /proj/sllstore2017093/b2016342/b2016342\_nobackup/lts/genome\_erosion\_pipeline/verena\_testing/testdata/reference/sumatran\_rhino\_22Jul2017\_9M7eS\_haploidified\_headersFixed\_Sc9M7eS\_2\_HRSCAF\_41.repeats.bed | |

##### Rule repeatmasker

×

Rule properties

|  |  |
| --- | --- |
| Jobs | 1 |
| Input files | |
| - /proj/sllstore2017093/b2016342/b2016342\_nobackup/lts/genome\_erosion\_pipeline/verena\_testing/testdata/reference/sumatran\_rhino\_22Jul2017\_9M7eS\_haploidified\_headersFixed\_Sc9M7eS\_2\_HRSCAF\_41.upper.fasta - /proj/sllstore2017093/b2016342/b2016342\_nobackup/lts/genome\_erosion\_pipeline/verena\_testing/testdata/reference/repeatmodeler/sumatran\_rhino\_22Jul2017\_9M7eS\_haploidified\_headersFixed\_Sc9M7eS\_2\_HRSCAF\_41/RM\_raw.out/consensi.fa.classified - workflow/resources/RepeatMasker/Libraries/RepeatMasker.lib - workflow/resources/RepeatMasker/Libraries/RepeatMasker.lib.nhr - workflow/resources/RepeatMasker/Libraries/RepeatMasker.lib.nin - workflow/resources/RepeatMasker/Libraries/RepeatMasker.lib.nsq - workflow/resources/RepeatMasker/Libraries/Artefacts.embl - workflow/resources/RepeatMasker/Libraries/Dfam.embl - workflow/resources/RepeatMasker/Libraries/Dfam.hmm - workflow/resources/RepeatMasker/Libraries/RepeatAnnotationData.pm - workflow/resources/RepeatMasker/Libraries/RepeatPeps.lib.phr - workflow/resources/RepeatMasker/Libraries/RepeatPeps.lib.psq - workflow/resources/RepeatMasker/Libraries/RepeatPeps.lib - workflow/resources/RepeatMasker/Libraries/RepeatPeps.lib.pin - workflow/resources/RepeatMasker/Libraries/RepeatPeps.readme - workflow/resources/RepeatMasker/Libraries/RMRBMeta.embl - workflow/resources/RepeatMasker/Libraries/README.meta - workflow/resources/RepeatMasker/Libraries/taxonomy.dat | |
| Output files | |
| - /proj/sllstore2017093/b2016342/b2016342\_nobackup/lts/genome\_erosion\_pipeline/verena\_testing/testdata/reference/repeatmasker/sumatran\_rhino\_22Jul2017\_9M7eS\_haploidified\_headersFixed\_Sc9M7eS\_2\_HRSCAF\_41/sumatran\_rhino\_22Jul2017\_9M7eS\_haploidified\_headersFixed\_Sc9M7eS\_2\_HRSCAF\_41.upper.fasta.masked - /proj/sllstore2017093/b2016342/b2016342\_nobackup/lts/genome\_erosion\_pipeline/verena\_testing/testdata/reference/repeatmasker/sumatran\_rhino\_22Jul2017\_9M7eS\_haploidified\_headersFixed\_Sc9M7eS\_2\_HRSCAF\_41/sumatran\_rhino\_22Jul2017\_9M7eS\_haploidified\_headersFixed\_Sc9M7eS\_2\_HRSCAF\_41.upper.fasta.align - /proj/sllstore2017093/b2016342/b2016342\_nobackup/lts/genome\_erosion\_pipeline/verena\_testing/testdata/reference/repeatmasker/sumatran\_rhino\_22Jul2017\_9M7eS\_haploidified\_headersFixed\_Sc9M7eS\_2\_HRSCAF\_41/sumatran\_rhino\_22Jul2017\_9M7eS\_haploidified\_headersFixed\_Sc9M7eS\_2\_HRSCAF\_41.upper.fasta.tbl - /proj/sllstore2017093/b2016342/b2016342\_nobackup/lts/genome\_erosion\_pipeline/verena\_testing/testdata/reference/repeatmasker/sumatran\_rhino\_22Jul2017\_9M7eS\_haploidified\_headersFixed\_Sc9M7eS\_2\_HRSCAF\_41/sumatran\_rhino\_22Jul2017\_9M7eS\_haploidified\_headersFixed\_Sc9M7eS\_2\_HRSCAF\_41.upper.fasta.out - /proj/sllstore2017093/b2016342/b2016342\_nobackup/lts/genome\_erosion\_pipeline/verena\_testing/testdata/reference/repeatmasker/sumatran\_rhino\_22Jul2017\_9M7eS\_haploidified\_headersFixed\_Sc9M7eS\_2\_HRSCAF\_41/sumatran\_rhino\_22Jul2017\_9M7eS\_haploidified\_headersFixed\_Sc9M7eS\_2\_HRSCAF\_41.upper.fasta.cat.gz | |
| Container image | |
| docker://quay.io/biocontainers/repeatmasker:4.0.9\_p2--pl526\_2 | |
| Code | |
| ``` 1 2 3 4 5 6 7 8 9 10 ``` \| ``` export REPEATMASKER_LIB_DIR=$PWD/workflow/resources/RepeatMasker/Libraries && cd {params.dir} && RepeatMasker -pa {threads} -a -xsmall -gccalc -dir ./ -lib {params.repmo} {params.ref_upper} 2> {log} && # Check if *.cat file is compressed or uncompressed if [ ! -f {output.rep_cat} ] # check if there are at least 2 files for merging. If there is only one file, copy the sorted bam file. then gzip {params.rep_cat_unzip} fi ``` | |

##### Rule ref\_upper

×

Rule properties

|  |  |
| --- | --- |
| Jobs | 1 |
| Input files | |
| - /proj/sllstore2017093/b2016342/b2016342\_nobackup/lts/genome\_erosion\_pipeline/verena\_testing/testdata/reference/sumatran\_rhino\_22Jul2017\_9M7eS\_haploidified\_headersFixed\_Sc9M7eS\_2\_HRSCAF\_41.fasta | |
| Output files | |
| - /proj/sllstore2017093/b2016342/b2016342\_nobackup/lts/genome\_erosion\_pipeline/verena\_testing/testdata/reference/sumatran\_rhino\_22Jul2017\_9M7eS\_haploidified\_headersFixed\_Sc9M7eS\_2\_HRSCAF\_41.upper.fasta | |
| Code | |
| ``` 1 2 ``` \| ``` awk '{{ if ($0 !~ />/) {{print toupper($0)}} else {{print $0}} }}' {input.ref} > {output.ref_upper} 2> {log} ``` | |

##### Rule repeatclassifier

×

Rule properties

|  |  |
| --- | --- |
| Jobs | 1 |
| Input files | |
| - /proj/sllstore2017093/b2016342/b2016342\_nobackup/lts/genome\_erosion\_pipeline/verena\_testing/testdata/reference/repeatmodeler/sumatran\_rhino\_22Jul2017\_9M7eS\_haploidified\_headersFixed\_Sc9M7eS\_2\_HRSCAF\_41/RM\_raw.out/consensi.fa - /proj/sllstore2017093/b2016342/b2016342\_nobackup/lts/genome\_erosion\_pipeline/verena\_testing/testdata/reference/repeatmodeler/sumatran\_rhino\_22Jul2017\_9M7eS\_haploidified\_headersFixed\_Sc9M7eS\_2\_HRSCAF\_41/RM\_raw.out/families.stk - workflow/resources/RepeatMasker/Libraries/RepeatMasker.lib - workflow/resources/RepeatMasker/Libraries/RepeatMasker.lib.nhr - workflow/resources/RepeatMasker/Libraries/RepeatMasker.lib.nin - workflow/resources/RepeatMasker/Libraries/RepeatMasker.lib.nsq - workflow/resources/RepeatMasker/Libraries/Artefacts.embl - workflow/resources/RepeatMasker/Libraries/Dfam.embl - workflow/resources/RepeatMasker/Libraries/Dfam.hmm - workflow/resources/RepeatMasker/Libraries/RepeatAnnotationData.pm - workflow/resources/RepeatMasker/Libraries/RepeatPeps.lib.phr - workflow/resources/RepeatMasker/Libraries/RepeatPeps.lib.psq - workflow/resources/RepeatMasker/Libraries/RepeatPeps.lib - workflow/resources/RepeatMasker/Libraries/RepeatPeps.lib.pin - workflow/resources/RepeatMasker/Libraries/RepeatPeps.readme - workflow/resources/RepeatMasker/Libraries/RMRBMeta.embl - workflow/resources/RepeatMasker/Libraries/README.meta - workflow/resources/RepeatMasker/Libraries/taxonomy.dat | |
| Output files | |
| - /proj/sllstore2017093/b2016342/b2016342\_nobackup/lts/genome\_erosion\_pipeline/verena\_testing/testdata/reference/repeatmodeler/sumatran\_rhino\_22Jul2017\_9M7eS\_haploidified\_headersFixed\_Sc9M7eS\_2\_HRSCAF\_41/RM\_raw.out/consensi.fa.classified - /proj/sllstore2017093/b2016342/b2016342\_nobackup/lts/genome\_erosion\_pipeline/verena\_testing/testdata/reference/repeatmodeler/sumatran\_rhino\_22Jul2017\_9M7eS\_haploidified\_headersFixed\_Sc9M7eS\_2\_HRSCAF\_41/RM\_raw.out/families-classified.stk | |
| Container image | |
| docker://quay.io/biocontainers/repeatmodeler:2.0.1--pl526\_0 | |
| Code | |
| ``` 1 2 ``` \| ``` RepeatClassifier -repeatmasker_dir {params.repma_dir} -consensi {input.repmo} -stockholm {input.stk} 2> {log} ``` | |

##### Rule repeatmodeler

×

Rule properties

|  |  |
| --- | --- |
| Jobs | 1 |
| Input files | |
| - /proj/sllstore2017093/b2016342/b2016342\_nobackup/lts/genome\_erosion\_pipeline/verena\_testing/testdata/reference/sumatran\_rhino\_22Jul2017\_9M7eS\_haploidified\_headersFixed\_Sc9M7eS\_2\_HRSCAF\_41.upper.fasta | |
| Output files | |
| - /proj/sllstore2017093/b2016342/b2016342\_nobackup/lts/genome\_erosion\_pipeline/verena\_testing/testdata/reference/repeatmodeler/sumatran\_rhino\_22Jul2017\_9M7eS\_haploidified\_headersFixed\_Sc9M7eS\_2\_HRSCAF\_41/RM\_raw.out/consensi.fa - /proj/sllstore2017093/b2016342/b2016342\_nobackup/lts/genome\_erosion\_pipeline/verena\_testing/testdata/reference/repeatmodeler/sumatran\_rhino\_22Jul2017\_9M7eS\_haploidified\_headersFixed\_Sc9M7eS\_2\_HRSCAF\_41/RM\_raw.out/families.stk | |
| Container image | |
| docker://quay.io/biocontainers/repeatmodeler:2.0.1--pl526\_0 | |
| Code | |
| ``` 1 2 3 4 5 6 7 8 9 10 11 12 13 14 15 16 17 18 ``` \| ``` cd {params.dir} # Build repeat database BuildDatabase -engine ncbi -name {params.name} {params.ref_upper} 2> {log} && # Run RepeatModeler RepeatModeler -engine ncbi -pa {threads} -database {params.name} 2>> {log} && # copy the output files to a new directory cp RM_*.*/consensi.fa RM_raw.out/ 2>> {log} && cp RM_*.*/families.stk RM_raw.out/ 2>> {log} # remove temporary file if [ -f {params.abs_tmp} ] then rm {params.abs_tmp} 2>> {log} fi ``` | |

##### Rule embl2fasta

×

Rule properties

|  |  |
| --- | --- |
| Jobs | 1 |
| Input files | |
| - workflow/resources/RepeatMasker/Libraries/Dfam.embl | |
| Output files | |
| - workflow/resources/RepeatMasker/Libraries/RepeatMasker.lib | |

##### Rule cp\_repeatmasker\_libs

×

Rule properties

|  |  |
| --- | --- |
| Jobs | 1 |
| Output files | |
| - workflow/resources/RepeatMasker/Libraries/Artefacts.embl - workflow/resources/RepeatMasker/Libraries/Dfam.embl - workflow/resources/RepeatMasker/Libraries/Dfam.hmm - workflow/resources/RepeatMasker/Libraries/RepeatAnnotationData.pm - workflow/resources/RepeatMasker/Libraries/RepeatPeps.lib.phr - workflow/resources/RepeatMasker/Libraries/RepeatPeps.lib.psq - workflow/resources/RepeatMasker/Libraries/RepeatPeps.lib - workflow/resources/RepeatMasker/Libraries/RepeatPeps.lib.pin - workflow/resources/RepeatMasker/Libraries/RepeatPeps.readme - workflow/resources/RepeatMasker/Libraries/RMRBMeta.embl - workflow/resources/RepeatMasker/Libraries/README.meta - workflow/resources/RepeatMasker/Libraries/taxonomy.dat | |
| Container image | |
| docker://quay.io/biocontainers/repeatmodeler:2.0.1--pl526\_0 | |
| Code | |
| ``` 1 2 ``` \| ``` cp /usr/local/share/RepeatMasker/Libraries/* workflow/resources/RepeatMasker/Libraries/ 2> {log} ``` | |

##### Rule make\_repma\_blast\_db

×

Rule properties

|  |  |
| --- | --- |
| Jobs | 1 |
| Input files | |
| - workflow/resources/RepeatMasker/Libraries/RepeatMasker.lib | |
| Output files | |
| - workflow/resources/RepeatMasker/Libraries/RepeatMasker.lib.nhr - workflow/resources/RepeatMasker/Libraries/RepeatMasker.lib.nin - workflow/resources/RepeatMasker/Libraries/RepeatMasker.lib.nsq | |
| Container image | |
| docker://quay.io/biocontainers/repeatmodeler:2.0.1--pl526\_0 | |
| Code | |
| ``` 1 2 3 ``` \| ``` cd {params.dir} makeblastdb -dbtype nucl -in {params.rm_lib} 2> {log} ``` | |

##### Rule make\_no\_repeats\_bed

×

Rule properties

|  |  |
| --- | --- |
| Jobs | 1 |
| Input files | |
| - /proj/sllstore2017093/b2016342/b2016342\_nobackup/lts/genome\_erosion\_pipeline/verena\_testing/testdata/reference/sumatran\_rhino\_22Jul2017\_9M7eS\_haploidified\_headersFixed\_Sc9M7eS\_2\_HRSCAF\_41.bed - /proj/sllstore2017093/b2016342/b2016342\_nobackup/lts/genome\_erosion\_pipeline/verena\_testing/testdata/reference/sumatran\_rhino\_22Jul2017\_9M7eS\_haploidified\_headersFixed\_Sc9M7eS\_2\_HRSCAF\_41.repeats.sorted.bed | |
| Output files | |
| - /proj/sllstore2017093/b2016342/b2016342\_nobackup/lts/genome\_erosion\_pipeline/verena\_testing/testdata/reference/sumatran\_rhino\_22Jul2017\_9M7eS\_haploidified\_headersFixed\_Sc9M7eS\_2\_HRSCAF\_41.repma.bed - results/sumatran\_rhino\_22Jul2017\_9M7eS\_haploidified\_headersFixed\_Sc9M7eS\_2\_HRSCAF\_41.repma.bed | |
| Container image | |
| docker://quay.io/biocontainers/bedtools:2.29.2--hc088bd4\_0 | |
| Code | |
| ``` 1 2 3 ``` \| ``` bedtools subtract -a {input.ref_bed} -b {input.sorted_rep_bed} > {output.no_rep_bed} 2> {log} && cp {output.no_rep_bed} {output.no_rep_bed_dir} 2>> {log} ``` | |

##### Rule sort\_repeats\_bed

×

Rule properties

|  |  |
| --- | --- |
| Jobs | 1 |
| Input files | |
| - /proj/sllstore2017093/b2016342/b2016342\_nobackup/lts/genome\_erosion\_pipeline/verena\_testing/testdata/reference/sumatran\_rhino\_22Jul2017\_9M7eS\_haploidified\_headersFixed\_Sc9M7eS\_2\_HRSCAF\_41.repeats.bed - /proj/sllstore2017093/b2016342/b2016342\_nobackup/lts/genome\_erosion\_pipeline/verena\_testing/testdata/reference/sumatran\_rhino\_22Jul2017\_9M7eS\_haploidified\_headersFixed\_Sc9M7eS\_2\_HRSCAF\_41.genome | |
| Output files | |
| - /proj/sllstore2017093/b2016342/b2016342\_nobackup/lts/genome\_erosion\_pipeline/verena\_testing/testdata/reference/sumatran\_rhino\_22Jul2017\_9M7eS\_haploidified\_headersFixed\_Sc9M7eS\_2\_HRSCAF\_41.repeats.sorted.bed | |
| Container image | |
| docker://quay.io/biocontainers/bedtools:2.29.2--hc088bd4\_0 | |
| Code | |
| ``` 1 2 ``` \| ``` bedtools sort -g {input.genomefile} -i {input.rep_bed} > {output.sorted_rep_bed} 2> {log} ``` | |

##### Rule historical\_raw\_bam\_multiqc

×

Rule properties

|  |  |
| --- | --- |
| Jobs | 1 |
| Input files | |
| - results/historical/mapping/sumatran\_rhino\_22Jul2017\_9M7eS\_haploidified\_headersFixed\_Sc9M7eS\_2\_HRSCAF\_41/stats/bams\_sorted/JvS008\_08\_L2.sorted.bam.stats.txt - results/historical/mapping/sumatran\_rhino\_22Jul2017\_9M7eS\_haploidified\_headersFixed\_Sc9M7eS\_2\_HRSCAF\_41/stats/bams\_sorted/JvS008\_08\_L6.sorted.bam.stats.txt - results/historical/mapping/sumatran\_rhino\_22Jul2017\_9M7eS\_haploidified\_headersFixed\_Sc9M7eS\_2\_HRSCAF\_41/stats/bams\_sorted/JvS008\_10\_L2.sorted.bam.stats.txt - results/historical/mapping/sumatran\_rhino\_22Jul2017\_9M7eS\_haploidified\_headersFixed\_Sc9M7eS\_2\_HRSCAF\_41/stats/bams\_sorted/JvS008\_10\_L6.sorted.bam.stats.txt - results/historical/mapping/sumatran\_rhino\_22Jul2017\_9M7eS\_haploidified\_headersFixed\_Sc9M7eS\_2\_HRSCAF\_41/stats/bams\_sorted/JvS008\_11\_L2.sorted.bam.stats.txt - results/historical/mapping/sumatran\_rhino\_22Jul2017\_9M7eS\_haploidified\_headersFixed\_Sc9M7eS\_2\_HRSCAF\_41/stats/bams\_sorted/JvS008\_11\_L6.sorted.bam.stats.txt - results/historical/mapping/sumatran\_rhino\_22Jul2017\_9M7eS\_haploidified\_headersFixed\_Sc9M7eS\_2\_HRSCAF\_41/stats/bams\_sorted/JvS009\_09\_L3.sorted.bam.stats.txt - results/historical/mapping/sumatran\_rhino\_22Jul2017\_9M7eS\_haploidified\_headersFixed\_Sc9M7eS\_2\_HRSCAF\_41/stats/bams\_sorted/JvS009\_09\_L7.sorted.bam.stats.txt - results/historical/mapping/sumatran\_rhino\_22Jul2017\_9M7eS\_haploidified\_headersFixed\_Sc9M7eS\_2\_HRSCAF\_41/stats/bams\_sorted/JvS009\_15\_L3.sorted.bam.stats.txt - results/historical/mapping/sumatran\_rhino\_22Jul2017\_9M7eS\_haploidified\_headersFixed\_Sc9M7eS\_2\_HRSCAF\_41/stats/bams\_sorted/JvS009\_15\_L7.sorted.bam.stats.txt - results/historical/mapping/sumatran\_rhino\_22Jul2017\_9M7eS\_haploidified\_headersFixed\_Sc9M7eS\_2\_HRSCAF\_41/stats/bams\_sorted/JvS009\_19\_L3.sorted.bam.stats.txt - results/historical/mapping/sumatran\_rhino\_22Jul2017\_9M7eS\_haploidified\_headersFixed\_Sc9M7eS\_2\_HRSCAF\_41/stats/bams\_sorted/JvS009\_19\_L7.sorted.bam.stats.txt - results/historical/mapping/sumatran\_rhino\_22Jul2017\_9M7eS\_haploidified\_headersFixed\_Sc9M7eS\_2\_HRSCAF\_41/stats/bams\_sorted/JvS022\_01\_L1.sorted.bam.stats.txt - results/historical/mapping/sumatran\_rhino\_22Jul2017\_9M7eS\_haploidified\_headersFixed\_Sc9M7eS\_2\_HRSCAF\_41/stats/bams\_sorted/JvS022\_02\_L1.sorted.bam.stats.txt - results/historical/mapping/sumatran\_rhino\_22Jul2017\_9M7eS\_haploidified\_headersFixed\_Sc9M7eS\_2\_HRSCAF\_41/stats/bams\_sorted/JvS022\_03\_L1.sorted.bam.stats.txt - results/historical/mapping/sumatran\_rhino\_22Jul2017\_9M7eS\_haploidified\_headersFixed\_Sc9M7eS\_2\_HRSCAF\_41/stats/bams\_sorted/JvS022\_04\_L1.sorted.bam.stats.txt - results/historical/mapping/sumatran\_rhino\_22Jul2017\_9M7eS\_haploidified\_headersFixed\_Sc9M7eS\_2\_HRSCAF\_41/stats/bams\_sorted/JvS022\_05\_L1.sorted.bam.stats.txt - results/historical/mapping/sumatran\_rhino\_22Jul2017\_9M7eS\_haploidified\_headersFixed\_Sc9M7eS\_2\_HRSCAF\_41/stats/bams\_sorted/JvS022\_06\_L1.sorted.bam.stats.txt - results/historical/mapping/sumatran\_rhino\_22Jul2017\_9M7eS\_haploidified\_headersFixed\_Sc9M7eS\_2\_HRSCAF\_41/stats/bams\_sorted/JvS022\_07\_L1.sorted.bam.stats.txt - results/historical/mapping/sumatran\_rhino\_22Jul2017\_9M7eS\_haploidified\_headersFixed\_Sc9M7eS\_2\_HRSCAF\_41/stats/bams\_sorted/JvS022\_08\_L1.sorted.bam.stats.txt - results/historical/mapping/sumatran\_rhino\_22Jul2017\_9M7eS\_haploidified\_headersFixed\_Sc9M7eS\_2\_HRSCAF\_41/stats/bams\_sorted/JvS022\_09\_L1.sorted.bam.stats.txt - results/historical/mapping/sumatran\_rhino\_22Jul2017\_9M7eS\_haploidified\_headersFixed\_Sc9M7eS\_2\_HRSCAF\_41/stats/bams\_sorted/JvS022\_10\_L1.sorted.bam.stats.txt - results/historical/mapping/sumatran\_rhino\_22Jul2017\_9M7eS\_haploidified\_headersFixed\_Sc9M7eS\_2\_HRSCAF\_41/stats/bams\_sorted/JvS022\_11\_L1.sorted.bam.stats.txt - results/historical/mapping/sumatran\_rhino\_22Jul2017\_9M7eS\_haploidified\_headersFixed\_Sc9M7eS\_2\_HRSCAF\_41/stats/bams\_sorted/JvS022\_12\_L1.sorted.bam.stats.txt - results/historical/mapping/sumatran\_rhino\_22Jul2017\_9M7eS\_haploidified\_headersFixed\_Sc9M7eS\_2\_HRSCAF\_41/stats/bams\_sorted/JvS022\_74\_L8.sorted.bam.stats.txt - results/historical/mapping/sumatran\_rhino\_22Jul2017\_9M7eS\_haploidified\_headersFixed\_Sc9M7eS\_2\_HRSCAF\_41/stats/bams\_sorted/JvS022\_75\_L8.sorted.bam.stats.txt - results/historical/mapping/sumatran\_rhino\_22Jul2017\_9M7eS\_haploidified\_headersFixed\_Sc9M7eS\_2\_HRSCAF\_41/stats/bams\_sorted/JvS022\_76\_L8.sorted.bam.stats.txt - results/historical/mapping/sumatran\_rhino\_22Jul2017\_9M7eS\_haploidified\_headersFixed\_Sc9M7eS\_2\_HRSCAF\_41/stats/bams\_sorted/JvS022\_77\_L8.sorted.bam.stats.txt - results/historical/mapping/sumatran\_rhino\_22Jul2017\_9M7eS\_haploidified\_headersFixed\_Sc9M7eS\_2\_HRSCAF\_41/stats/bams\_sorted/JvS022\_78\_L8.sorted.bam.stats.txt - results/historical/mapping/sumatran\_rhino\_22Jul2017\_9M7eS\_haploidified\_headersFixed\_Sc9M7eS\_2\_HRSCAF\_41/stats/bams\_sorted/JvS022\_79\_L8.sorted.bam.stats.txt - results/historical/mapping/sumatran\_rhino\_22Jul2017\_9M7eS\_haploidified\_headersFixed\_Sc9M7eS\_2\_HRSCAF\_41/stats/bams\_sorted/JvS022\_80\_L8.sorted.bam.stats.txt - results/historical/mapping/sumatran\_rhino\_22Jul2017\_9M7eS\_haploidified\_headersFixed\_Sc9M7eS\_2\_HRSCAF\_41/stats/bams\_sorted/JvS022\_81\_L8.sorted.bam.stats.txt - results/historical/mapping/sumatran\_rhino\_22Jul2017\_9M7eS\_haploidified\_headersFixed\_Sc9M7eS\_2\_HRSCAF\_41/stats/bams\_sorted/JvS022\_82\_L8.sorted.bam.stats.txt - results/historical/mapping/sumatran\_rhino\_22Jul2017\_9M7eS\_haploidified\_headersFixed\_Sc9M7eS\_2\_HRSCAF\_41/stats/bams\_sorted/JvS022\_83\_L8.sorted.bam.stats.txt - results/historical/mapping/sumatran\_rhino\_22Jul2017\_9M7eS\_haploidified\_headersFixed\_Sc9M7eS\_2\_HRSCAF\_41/stats/bams\_sorted/JvS022\_84\_L8.sorted.bam.stats.txt - results/historical/mapping/sumatran\_rhino\_22Jul2017\_9M7eS\_haploidified\_headersFixed\_Sc9M7eS\_2\_HRSCAF\_41/stats/bams\_sorted/JvS022\_85\_L8.sorted.bam.stats.txt - results/historical/mapping/sumatran\_rhino\_22Jul2017\_9M7eS\_haploidified\_headersFixed\_Sc9M7eS\_2\_HRSCAF\_41/stats/bams\_sorted/JvS008\_08\_L2.sorted.bam.qualimap/qualimapReport.html - results/historical/mapping/sumatran\_rhino\_22Jul2017\_9M7eS\_haploidified\_headersFixed\_Sc9M7eS\_2\_HRSCAF\_41/stats/bams\_sorted/JvS008\_08\_L6.sorted.bam.qualimap/qualimapReport.html - results/historical/mapping/sumatran\_rhino\_22Jul2017\_9M7eS\_haploidified\_headersFixed\_Sc9M7eS\_2\_HRSCAF\_41/stats/bams\_sorted/JvS008\_10\_L2.sorted.bam.qualimap/qualimapReport.html - results/historical/mapping/sumatran\_rhino\_22Jul2017\_9M7eS\_haploidified\_headersFixed\_Sc9M7eS\_2\_HRSCAF\_41/stats/bams\_sorted/JvS008\_10\_L6.sorted.bam.qualimap/qualimapReport.html - results/historical/mapping/sumatran\_rhino\_22Jul2017\_9M7eS\_haploidified\_headersFixed\_Sc9M7eS\_2\_HRSCAF\_41/stats/bams\_sorted/JvS008\_11\_L2.sorted.bam.qualimap/qualimapReport.html - results/historical/mapping/sumatran\_rhino\_22Jul2017\_9M7eS\_haploidified\_headersFixed\_Sc9M7eS\_2\_HRSCAF\_41/stats/bams\_sorted/JvS008\_11\_L6.sorted.bam.qualimap/qualimapReport.html - results/historical/mapping/sumatran\_rhino\_22Jul2017\_9M7eS\_haploidified\_headersFixed\_Sc9M7eS\_2\_HRSCAF\_41/stats/bams\_sorted/JvS009\_09\_L3.sorted.bam.qualimap/qualimapReport.html - results/historical/mapping/sumatran\_rhino\_22Jul2017\_9M7eS\_haploidified\_headersFixed\_Sc9M7eS\_2\_HRSCAF\_41/stats/bams\_sorted/JvS009\_09\_L7.sorted.bam.qualimap/qualimapReport.html - results/historical/mapping/sumatran\_rhino\_22Jul2017\_9M7eS\_haploidified\_headersFixed\_Sc9M7eS\_2\_HRSCAF\_41/stats/bams\_sorted/JvS009\_15\_L3.sorted.bam.qualimap/qualimapReport.html - results/historical/mapping/sumatran\_rhino\_22Jul2017\_9M7eS\_haploidified\_headersFixed\_Sc9M7eS\_2\_HRSCAF\_41/stats/bams\_sorted/JvS009\_15\_L7.sorted.bam.qualimap/qualimapReport.html - results/historical/mapping/sumatran\_rhino\_22Jul2017\_9M7eS\_haploidified\_headersFixed\_Sc9M7eS\_2\_HRSCAF\_41/stats/bams\_sorted/JvS009\_19\_L3.sorted.bam.qualimap/qualimapReport.html - results/historical/mapping/sumatran\_rhino\_22Jul2017\_9M7eS\_haploidified\_headersFixed\_Sc9M7eS\_2\_HRSCAF\_41/stats/bams\_sorted/JvS009\_19\_L7.sorted.bam.qualimap/qualimapReport.html - results/historical/mapping/sumatran\_rhino\_22Jul2017\_9M7eS\_haploidified\_headersFixed\_Sc9M7eS\_2\_HRSCAF\_41/stats/bams\_sorted/JvS022\_01\_L1.sorted.bam.qualimap/qualimapReport.html - results/historical/mapping/sumatran\_rhino\_22Jul2017\_9M7eS\_haploidified\_headersFixed\_Sc9M7eS\_2\_HRSCAF\_41/stats/bams\_sorted/JvS022\_02\_L1.sorted.bam.qualimap/qualimapReport.html - results/historical/mapping/sumatran\_rhino\_22Jul2017\_9M7eS\_haploidified\_headersFixed\_Sc9M7eS\_2\_HRSCAF\_41/stats/bams\_sorted/JvS022\_03\_L1.sorted.bam.qualimap/qualimapReport.html - results/historical/mapping/sumatran\_rhino\_22Jul2017\_9M7eS\_haploidified\_headersFixed\_Sc9M7eS\_2\_HRSCAF\_41/stats/bams\_sorted/JvS022\_04\_L1.sorted.bam.qualimap/qualimapReport.html - results/historical/mapping/sumatran\_rhino\_22Jul2017\_9M7eS\_haploidified\_headersFixed\_Sc9M7eS\_2\_HRSCAF\_41/stats/bams\_sorted/JvS022\_05\_L1.sorted.bam.qualimap/qualimapReport.html - results/historical/mapping/sumatran\_rhino\_22Jul2017\_9M7eS\_haploidified\_headersFixed\_Sc9M7eS\_2\_HRSCAF\_41/stats/bams\_sorted/JvS022\_06\_L1.sorted.bam.qualimap/qualimapReport.html - results/historical/mapping/sumatran\_rhino\_22Jul2017\_9M7eS\_haploidified\_headersFixed\_Sc9M7eS\_2\_HRSCAF\_41/stats/bams\_sorted/JvS022\_07\_L1.sorted.bam.qualimap/qualimapReport.html - results/historical/mapping/sumatran\_rhino\_22Jul2017\_9M7eS\_haploidified\_headersFixed\_Sc9M7eS\_2\_HRSCAF\_41/stats/bams\_sorted/JvS022\_08\_L1.sorted.bam.qualimap/qualimapReport.html - results/historical/mapping/sumatran\_rhino\_22Jul2017\_9M7eS\_haploidified\_headersFixed\_Sc9M7eS\_2\_HRSCAF\_41/stats/bams\_sorted/JvS022\_09\_L1.sorted.bam.qualimap/qualimapReport.html - results/historical/mapping/sumatran\_rhino\_22Jul2017\_9M7eS\_haploidified\_headersFixed\_Sc9M7eS\_2\_HRSCAF\_41/stats/bams\_sorted/JvS022\_10\_L1.sorted.bam.qualimap/qualimapReport.html - results/historical/mapping/sumatran\_rhino\_22Jul2017\_9M7eS\_haploidified\_headersFixed\_Sc9M7eS\_2\_HRSCAF\_41/stats/bams\_sorted/JvS022\_11\_L1.sorted.bam.qualimap/qualimapReport.html - results/historical/mapping/sumatran\_rhino\_22Jul2017\_9M7eS\_haploidified\_headersFixed\_Sc9M7eS\_2\_HRSCAF\_41/stats/bams\_sorted/JvS022\_12\_L1.sorted.bam.qualimap/qualimapReport.html - results/historical/mapping/sumatran\_rhino\_22Jul2017\_9M7eS\_haploidified\_headersFixed\_Sc9M7eS\_2\_HRSCAF\_41/stats/bams\_sorted/JvS022\_74\_L8.sorted.bam.qualimap/qualimapReport.html - results/historical/mapping/sumatran\_rhino\_22Jul2017\_9M7eS\_haploidified\_headersFixed\_Sc9M7eS\_2\_HRSCAF\_41/stats/bams\_sorted/JvS022\_75\_L8.sorted.bam.qualimap/qualimapReport.html - results/historical/mapping/sumatran\_rhino\_22Jul2017\_9M7eS\_haploidified\_headersFixed\_Sc9M7eS\_2\_HRSCAF\_41/stats/bams\_sorted/JvS022\_76\_L8.sorted.bam.qualimap/qualimapReport.html - results/historical/mapping/sumatran\_rhino\_22Jul2017\_9M7eS\_haploidified\_headersFixed\_Sc9M7eS\_2\_HRSCAF\_41/stats/bams\_sorted/JvS022\_77\_L8.sorted.bam.qualimap/qualimapReport.html - results/historical/mapping/sumatran\_rhino\_22Jul2017\_9M7eS\_haploidified\_headersFixed\_Sc9M7eS\_2\_HRSCAF\_41/stats/bams\_sorted/JvS022\_78\_L8.sorted.bam.qualimap/qualimapReport.html - results/historical/mapping/sumatran\_rhino\_22Jul2017\_9M7eS\_haploidified\_headersFixed\_Sc9M7eS\_2\_HRSCAF\_41/stats/bams\_sorted/JvS022\_79\_L8.sorted.bam.qualimap/qualimapReport.html - results/historical/mapping/sumatran\_rhino\_22Jul2017\_9M7eS\_haploidified\_headersFixed\_Sc9M7eS\_2\_HRSCAF\_41/stats/bams\_sorted/JvS022\_80\_L8.sorted.bam.qualimap/qualimapReport.html - results/historical/mapping/sumatran\_rhino\_22Jul2017\_9M7eS\_haploidified\_headersFixed\_Sc9M7eS\_2\_HRSCAF\_41/stats/bams\_sorted/JvS022\_81\_L8.sorted.bam.qualimap/qualimapReport.html - results/historical/mapping/sumatran\_rhino\_22Jul2017\_9M7eS\_haploidified\_headersFixed\_Sc9M7eS\_2\_HRSCAF\_41/stats/bams\_sorted/JvS022\_82\_L8.sorted.bam.qualimap/qualimapReport.html - results/historical/mapping/sumatran\_rhino\_22Jul2017\_9M7eS\_haploidified\_headersFixed\_Sc9M7eS\_2\_HRSCAF\_41/stats/bams\_sorted/JvS022\_83\_L8.sorted.bam.qualimap/qualimapReport.html - results/historical/mapping/sumatran\_rhino\_22Jul2017\_9M7eS\_haploidified\_headersFixed\_Sc9M7eS\_2\_HRSCAF\_41/stats/bams\_sorted/JvS022\_84\_L8.sorted.bam.qualimap/qualimapReport.html - results/historical/mapping/sumatran\_rhino\_22Jul2017\_9M7eS\_haploidified\_headersFixed\_Sc9M7eS\_2\_HRSCAF\_41/stats/bams\_sorted/JvS022\_85\_L8.sorted.bam.qualimap/qualimapReport.html | |
| Output files | |
| - results/historical/mapping/sumatran\_rhino\_22Jul2017\_9M7eS\_haploidified\_headersFixed\_Sc9M7eS\_2\_HRSCAF\_41/stats/bams\_sorted/multiqc/multiqc\_report.html | |
| Container image | |
| docker://quay.io/biocontainers/multiqc:1.9--pyh9f0ad1d\_0 | |
| Code | |
| ``` 1 2 ``` \| ``` multiqc -f {params.indir} -o {params.outdir} 2> {log} ``` | |

##### Rule sorted\_bam\_stats

×

Rule properties

|  |  |
| --- | --- |
| Jobs | 39 |
| Input files | |
| - results/{dataset}/mapping/sumatran\_rhino\_22Jul2017\_9M7eS\_haploidified\_headersFixed\_Sc9M7eS\_2\_HRSCAF\_41/{sample}\_{index}\_{lane}.sorted.bam - results/{dataset}/mapping/sumatran\_rhino\_22Jul2017\_9M7eS\_haploidified\_headersFixed\_Sc9M7eS\_2\_HRSCAF\_41/{sample}\_{index}\_{lane}.sorted.bam.bai | |
| Output files | |
| - results/{dataset}/mapping/sumatran\_rhino\_22Jul2017\_9M7eS\_haploidified\_headersFixed\_Sc9M7eS\_2\_HRSCAF\_41/stats/bams\_sorted/{sample,[A-Za-z0-9]+}\_{index}\_{lane}.sorted.bam.stats.txt | |
| Container image | |
| docker://biocontainers/samtools:v1.9-4-deb\_cv1 | |
| Code | |
| ``` 1 2 ``` \| ``` samtools flagstat {input.bam} > {output.stats} 2> {log} ``` | |

##### Rule sai2bam

×

Rule properties

|  |  |
| --- | --- |
| Jobs | 36 |
| Input files | |
| - /proj/sllstore2017093/b2016342/b2016342\_nobackup/lts/genome\_erosion\_pipeline/verena\_testing/testdata/reference/sumatran\_rhino\_22Jul2017\_9M7eS\_haploidified\_headersFixed\_Sc9M7eS\_2\_HRSCAF\_41.fasta - /proj/sllstore2017093/b2016342/b2016342\_nobackup/lts/genome\_erosion\_pipeline/verena\_testing/testdata/reference/sumatran\_rhino\_22Jul2017\_9M7eS\_haploidified\_headersFixed\_Sc9M7eS\_2\_HRSCAF\_41.fasta.amb - /proj/sllstore2017093/b2016342/b2016342\_nobackup/lts/genome\_erosion\_pipeline/verena\_testing/testdata/reference/sumatran\_rhino\_22Jul2017\_9M7eS\_haploidified\_headersFixed\_Sc9M7eS\_2\_HRSCAF\_41.fasta.ann - /proj/sllstore2017093/b2016342/b2016342\_nobackup/lts/genome\_erosion\_pipeline/verena\_testing/testdata/reference/sumatran\_rhino\_22Jul2017\_9M7eS\_haploidified\_headersFixed\_Sc9M7eS\_2\_HRSCAF\_41.fasta.bwt - /proj/sllstore2017093/b2016342/b2016342\_nobackup/lts/genome\_erosion\_pipeline/verena\_testing/testdata/reference/sumatran\_rhino\_22Jul2017\_9M7eS\_haploidified\_headersFixed\_Sc9M7eS\_2\_HRSCAF\_41.fasta.pac - /proj/sllstore2017093/b2016342/b2016342\_nobackup/lts/genome\_erosion\_pipeline/verena\_testing/testdata/reference/sumatran\_rhino\_22Jul2017\_9M7eS\_haploidified\_headersFixed\_Sc9M7eS\_2\_HRSCAF\_41.fasta.sa - results/historical/trimming/{sample}\_{index}\_{lane}\_trimmed\_merged.fastq.gz - results/historical/mapping/sumatran\_rhino\_22Jul2017\_9M7eS\_haploidified\_headersFixed\_Sc9M7eS\_2\_HRSCAF\_41/{sample}\_{index}\_{lane}.sai - results/historical/mapping/sumatran\_rhino\_22Jul2017\_9M7eS\_haploidified\_headersFixed\_Sc9M7eS\_2\_HRSCAF\_41/{sample}\_{index}\_{lane}.readgroup.txt | |
| Output files | |
| - results/historical/mapping/sumatran\_rhino\_22Jul2017\_9M7eS\_haploidified\_headersFixed\_Sc9M7eS\_2\_HRSCAF\_41/{sample,[A-Za-z0-9]+}\_{index}\_{lane}.sorted.bam | |
| Container image | |
| docker://nbisweden/generode-bwa:latest | |
| Code | |
| ``` 1 2 ``` \| ``` bwa samse -r $(cat {input.rg}) {input.ref} {input.sai} {input.fastq_hist} \| samtools sort -@ {threads} - > {output.bam} 2> {log} ``` | |

##### Rule map\_historical

×

Rule properties

|  |  |
| --- | --- |
| Jobs | 36 |
| Input files | |
| - /proj/sllstore2017093/b2016342/b2016342\_nobackup/lts/genome\_erosion\_pipeline/verena\_testing/testdata/reference/sumatran\_rhino\_22Jul2017\_9M7eS\_haploidified\_headersFixed\_Sc9M7eS\_2\_HRSCAF\_41.fasta - /proj/sllstore2017093/b2016342/b2016342\_nobackup/lts/genome\_erosion\_pipeline/verena\_testing/testdata/reference/sumatran\_rhino\_22Jul2017\_9M7eS\_haploidified\_headersFixed\_Sc9M7eS\_2\_HRSCAF\_41.fasta.amb - /proj/sllstore2017093/b2016342/b2016342\_nobackup/lts/genome\_erosion\_pipeline/verena\_testing/testdata/reference/sumatran\_rhino\_22Jul2017\_9M7eS\_haploidified\_headersFixed\_Sc9M7eS\_2\_HRSCAF\_41.fasta.ann - /proj/sllstore2017093/b2016342/b2016342\_nobackup/lts/genome\_erosion\_pipeline/verena\_testing/testdata/reference/sumatran\_rhino\_22Jul2017\_9M7eS\_haploidified\_headersFixed\_Sc9M7eS\_2\_HRSCAF\_41.fasta.bwt - /proj/sllstore2017093/b2016342/b2016342\_nobackup/lts/genome\_erosion\_pipeline/verena\_testing/testdata/reference/sumatran\_rhino\_22Jul2017\_9M7eS\_haploidified\_headersFixed\_Sc9M7eS\_2\_HRSCAF\_41.fasta.pac - /proj/sllstore2017093/b2016342/b2016342\_nobackup/lts/genome\_erosion\_pipeline/verena\_testing/testdata/reference/sumatran\_rhino\_22Jul2017\_9M7eS\_haploidified\_headersFixed\_Sc9M7eS\_2\_HRSCAF\_41.fasta.sa - results/historical/trimming/{sample}\_{index}\_{lane}\_trimmed\_merged.fastq.gz | |
| Output files | |
| - results/historical/mapping/sumatran\_rhino\_22Jul2017\_9M7eS\_haploidified\_headersFixed\_Sc9M7eS\_2\_HRSCAF\_41/{sample,[A-Za-z0-9]+}\_{index}\_{lane}.sai | |
| Container image | |
| docker://biocontainers/bwa:v0.7.17-3-deb\_cv1 | |
| Code | |
| ``` 1 2 ``` \| ``` bwa aln -l 16500 -n 0.01 -o 2 -t {threads} {input.ref} {input.fastq_hist} > {output.sai} 2> {log} ``` | |

##### Rule readgroup\_ID\_historical

×

Rule properties

|  |  |
| --- | --- |
| Jobs | 36 |
| Input files | |
| - results/historical/mapping/sumatran\_rhino\_22Jul2017\_9M7eS\_haploidified\_headersFixed\_Sc9M7eS\_2\_HRSCAF\_41/{sample}\_{index}\_{lane}.sai | |
| Output files | |
| - results/historical/mapping/sumatran\_rhino\_22Jul2017\_9M7eS\_haploidified\_headersFixed\_Sc9M7eS\_2\_HRSCAF\_41/{sample,[A-Za-z0-9]+}\_{index}\_{lane}.readgroup.txt | |

##### Rule index\_sorted\_bams

×

Rule properties

|  |  |
| --- | --- |
| Jobs | 39 |
| Input files | |
| - results/{dataset}/mapping/sumatran\_rhino\_22Jul2017\_9M7eS\_haploidified\_headersFixed\_Sc9M7eS\_2\_HRSCAF\_41/{sample}\_{index}\_{lane}.sorted.bam | |
| Output files | |
| - results/{dataset}/mapping/sumatran\_rhino\_22Jul2017\_9M7eS\_haploidified\_headersFixed\_Sc9M7eS\_2\_HRSCAF\_41/{sample,[A-Za-z0-9]+}\_{index}\_{lane}.sorted.bam.bai | |
| Container image | |
| docker://biocontainers/samtools:v1.9-4-deb\_cv1 | |
| Code | |
| ``` 1 2 ``` \| ``` samtools index {input.bam} {output.index} 2> {log} ``` | |

##### Rule sorted\_bam\_qualimap

×

Rule properties

|  |  |
| --- | --- |
| Jobs | 39 |
| Input files | |
| - results/{dataset}/mapping/sumatran\_rhino\_22Jul2017\_9M7eS\_haploidified\_headersFixed\_Sc9M7eS\_2\_HRSCAF\_41/{sample}\_{index}\_{lane}.sorted.bam - results/{dataset}/mapping/sumatran\_rhino\_22Jul2017\_9M7eS\_haploidified\_headersFixed\_Sc9M7eS\_2\_HRSCAF\_41/{sample}\_{index}\_{lane}.sorted.bam.bai | |
| Output files | |
| - results/{dataset}/mapping/sumatran\_rhino\_22Jul2017\_9M7eS\_haploidified\_headersFixed\_Sc9M7eS\_2\_HRSCAF\_41/stats/bams\_sorted/{sample,[A-Za-z0-9]+}\_{index}\_{lane}.sorted.bam.qualimap/qualimapReport.html - results/{dataset}/mapping/sumatran\_rhino\_22Jul2017\_9M7eS\_haploidified\_headersFixed\_Sc9M7eS\_2\_HRSCAF\_41/stats/bams\_sorted/{sample,[A-Za-z0-9]+}\_{index}\_{lane}.sorted.bam.qualimap/genome\_results.txt - results/{dataset}/mapping/sumatran\_rhino\_22Jul2017\_9M7eS\_haploidified\_headersFixed\_Sc9M7eS\_2\_HRSCAF\_41/stats/bams\_sorted/{sample,[A-Za-z0-9]+}\_{index}\_{lane}.sorted.bam.qualimap | |
| Container image | |
| docker://quay.io/biocontainers/qualimap:2.2.2d--1 | |
| Code | |
| ``` 1 2 3 4 ``` \| ``` mem=$(((6 * {threads}) - 2)) unset DISPLAY qualimap bamqc -bam {input.bam} --java-mem-size=${{mem}}G -nt {threads} -outdir {params.outdir} -outformat html 2> {log} ``` | |

##### Rule modern\_raw\_bam\_multiqc

×

Rule properties

|  |  |
| --- | --- |
| Jobs | 1 |
| Input files | |
| - results/modern/mapping/sumatran\_rhino\_22Jul2017\_9M7eS\_haploidified\_headersFixed\_Sc9M7eS\_2\_HRSCAF\_41/stats/bams\_sorted/JvS033\_01\_L7.sorted.bam.stats.txt - results/modern/mapping/sumatran\_rhino\_22Jul2017\_9M7eS\_haploidified\_headersFixed\_Sc9M7eS\_2\_HRSCAF\_41/stats/bams\_sorted/JvS034\_02\_L8.sorted.bam.stats.txt - results/modern/mapping/sumatran\_rhino\_22Jul2017\_9M7eS\_haploidified\_headersFixed\_Sc9M7eS\_2\_HRSCAF\_41/stats/bams\_sorted/JvS035\_14\_L7.sorted.bam.stats.txt - results/modern/mapping/sumatran\_rhino\_22Jul2017\_9M7eS\_haploidified\_headersFixed\_Sc9M7eS\_2\_HRSCAF\_41/stats/bams\_sorted/JvS033\_01\_L7.sorted.bam.qualimap/qualimapReport.html - results/modern/mapping/sumatran\_rhino\_22Jul2017\_9M7eS\_haploidified\_headersFixed\_Sc9M7eS\_2\_HRSCAF\_41/stats/bams\_sorted/JvS034\_02\_L8.sorted.bam.qualimap/qualimapReport.html - results/modern/mapping/sumatran\_rhino\_22Jul2017\_9M7eS\_haploidified\_headersFixed\_Sc9M7eS\_2\_HRSCAF\_41/stats/bams\_sorted/JvS035\_14\_L7.sorted.bam.qualimap/qualimapReport.html | |
| Output files | |
| - results/modern/mapping/sumatran\_rhino\_22Jul2017\_9M7eS\_haploidified\_headersFixed\_Sc9M7eS\_2\_HRSCAF\_41/stats/bams\_sorted/multiqc/multiqc\_report.html | |
| Container image | |
| docker://quay.io/biocontainers/multiqc:1.9--pyh9f0ad1d\_0 | |
| Code | |
| ``` 1 2 ``` \| ``` multiqc -f {params.indir} -o {params.outdir} 2> {log} ``` | |

##### Rule map\_modern

×

Rule properties

|  |  |
| --- | --- |
| Jobs | 3 |
| Input files | |
| - /proj/sllstore2017093/b2016342/b2016342\_nobackup/lts/genome\_erosion\_pipeline/verena\_testing/testdata/reference/sumatran\_rhino\_22Jul2017\_9M7eS\_haploidified\_headersFixed\_Sc9M7eS\_2\_HRSCAF\_41.fasta - /proj/sllstore2017093/b2016342/b2016342\_nobackup/lts/genome\_erosion\_pipeline/verena\_testing/testdata/reference/sumatran\_rhino\_22Jul2017\_9M7eS\_haploidified\_headersFixed\_Sc9M7eS\_2\_HRSCAF\_41.fasta.amb - /proj/sllstore2017093/b2016342/b2016342\_nobackup/lts/genome\_erosion\_pipeline/verena\_testing/testdata/reference/sumatran\_rhino\_22Jul2017\_9M7eS\_haploidified\_headersFixed\_Sc9M7eS\_2\_HRSCAF\_41.fasta.ann - /proj/sllstore2017093/b2016342/b2016342\_nobackup/lts/genome\_erosion\_pipeline/verena\_testing/testdata/reference/sumatran\_rhino\_22Jul2017\_9M7eS\_haploidified\_headersFixed\_Sc9M7eS\_2\_HRSCAF\_41.fasta.bwt - /proj/sllstore2017093/b2016342/b2016342\_nobackup/lts/genome\_erosion\_pipeline/verena\_testing/testdata/reference/sumatran\_rhino\_22Jul2017\_9M7eS\_haploidified\_headersFixed\_Sc9M7eS\_2\_HRSCAF\_41.fasta.pac - /proj/sllstore2017093/b2016342/b2016342\_nobackup/lts/genome\_erosion\_pipeline/verena\_testing/testdata/reference/sumatran\_rhino\_22Jul2017\_9M7eS\_haploidified\_headersFixed\_Sc9M7eS\_2\_HRSCAF\_41.fasta.sa - results/modern/trimming/{sample}\_{index}\_{lane}\_R1\_trimmed.fastq.gz - results/modern/trimming/{sample}\_{index}\_{lane}\_R2\_trimmed.fastq.gz - results/modern/mapping/sumatran\_rhino\_22Jul2017\_9M7eS\_haploidified\_headersFixed\_Sc9M7eS\_2\_HRSCAF\_41/{sample}\_{index}\_{lane}.readgroup.txt | |
| Output files | |
| - results/modern/mapping/sumatran\_rhino\_22Jul2017\_9M7eS\_haploidified\_headersFixed\_Sc9M7eS\_2\_HRSCAF\_41/{sample,[A-Za-z0-9]+}\_{index}\_{lane}.sorted.bam | |
| Container image | |
| docker://nbisweden/generode-bwa:latest | |
| Code | |
| ``` 1 2 ``` \| ``` bwa mem -M -t {threads} -R $(cat {input.rg}) {input.ref} {input.fastq_mod_R1} {input.fastq_mod_R2} \| samtools sort -@ {threads} - > {output.bam} 2> {log} ``` | |

##### Rule readgroup\_ID\_modern

×

Rule properties

|  |  |
| --- | --- |
| Jobs | 3 |
| Input files | |
| - /proj/sllstore2017093/b2016342/b2016342\_nobackup/lts/genome\_erosion\_pipeline/verena\_testing/testdata/reference/sumatran\_rhino\_22Jul2017\_9M7eS\_haploidified\_headersFixed\_Sc9M7eS\_2\_HRSCAF\_41.fasta - /proj/sllstore2017093/b2016342/b2016342\_nobackup/lts/genome\_erosion\_pipeline/verena\_testing/testdata/reference/sumatran\_rhino\_22Jul2017\_9M7eS\_haploidified\_headersFixed\_Sc9M7eS\_2\_HRSCAF\_41.fasta.amb - /proj/sllstore2017093/b2016342/b2016342\_nobackup/lts/genome\_erosion\_pipeline/verena\_testing/testdata/reference/sumatran\_rhino\_22Jul2017\_9M7eS\_haploidified\_headersFixed\_Sc9M7eS\_2\_HRSCAF\_41.fasta.ann - /proj/sllstore2017093/b2016342/b2016342\_nobackup/lts/genome\_erosion\_pipeline/verena\_testing/testdata/reference/sumatran\_rhino\_22Jul2017\_9M7eS\_haploidified\_headersFixed\_Sc9M7eS\_2\_HRSCAF\_41.fasta.bwt - /proj/sllstore2017093/b2016342/b2016342\_nobackup/lts/genome\_erosion\_pipeline/verena\_testing/testdata/reference/sumatran\_rhino\_22Jul2017\_9M7eS\_haploidified\_headersFixed\_Sc9M7eS\_2\_HRSCAF\_41.fasta.pac - /proj/sllstore2017093/b2016342/b2016342\_nobackup/lts/genome\_erosion\_pipeline/verena\_testing/testdata/reference/sumatran\_rhino\_22Jul2017\_9M7eS\_haploidified\_headersFixed\_Sc9M7eS\_2\_HRSCAF\_41.fasta.sa - results/modern/trimming/{sample}\_{index}\_{lane}\_R1\_trimmed.fastq.gz - results/modern/trimming/{sample}\_{index}\_{lane}\_R2\_trimmed.fastq.gz | |
| Output files | |
| - results/modern/mapping/sumatran\_rhino\_22Jul2017\_9M7eS\_haploidified\_headersFixed\_Sc9M7eS\_2\_HRSCAF\_41/{sample,[A-Za-z0-9]+}\_{index}\_{lane}.readgroup.txt | |

##### Rule historical\_merged\_index\_bam\_multiqc

×

Rule properties

|  |  |
| --- | --- |
| Jobs | 1 |
| Input files | |
| - results/historical/mapping/sumatran\_rhino\_22Jul2017\_9M7eS\_haploidified\_headersFixed\_Sc9M7eS\_2\_HRSCAF\_41/stats/bams\_merged\_index/JvS008\_08.merged.bam.stats.txt - results/historical/mapping/sumatran\_rhino\_22Jul2017\_9M7eS\_haploidified\_headersFixed\_Sc9M7eS\_2\_HRSCAF\_41/stats/bams\_merged\_index/JvS008\_10.merged.bam.stats.txt - results/historical/mapping/sumatran\_rhino\_22Jul2017\_9M7eS\_haploidified\_headersFixed\_Sc9M7eS\_2\_HRSCAF\_41/stats/bams\_merged\_index/JvS008\_11.merged.bam.stats.txt - results/historical/mapping/sumatran\_rhino\_22Jul2017\_9M7eS\_haploidified\_headersFixed\_Sc9M7eS\_2\_HRSCAF\_41/stats/bams\_merged\_index/JvS009\_09.merged.bam.stats.txt - results/historical/mapping/sumatran\_rhino\_22Jul2017\_9M7eS\_haploidified\_headersFixed\_Sc9M7eS\_2\_HRSCAF\_41/stats/bams\_merged\_index/JvS009\_15.merged.bam.stats.txt - results/historical/mapping/sumatran\_rhino\_22Jul2017\_9M7eS\_haploidified\_headersFixed\_Sc9M7eS\_2\_HRSCAF\_41/stats/bams\_merged\_index/JvS009\_19.merged.bam.stats.txt - results/historical/mapping/sumatran\_rhino\_22Jul2017\_9M7eS\_haploidified\_headersFixed\_Sc9M7eS\_2\_HRSCAF\_41/stats/bams\_merged\_index/JvS022\_01.merged.bam.stats.txt - results/historical/mapping/sumatran\_rhino\_22Jul2017\_9M7eS\_haploidified\_headersFixed\_Sc9M7eS\_2\_HRSCAF\_41/stats/bams\_merged\_index/JvS022\_02.merged.bam.stats.txt - results/historical/mapping/sumatran\_rhino\_22Jul2017\_9M7eS\_haploidified\_headersFixed\_Sc9M7eS\_2\_HRSCAF\_41/stats/bams\_merged\_index/JvS022\_03.merged.bam.stats.txt - results/historical/mapping/sumatran\_rhino\_22Jul2017\_9M7eS\_haploidified\_headersFixed\_Sc9M7eS\_2\_HRSCAF\_41/stats/bams\_merged\_index/JvS022\_04.merged.bam.stats.txt - results/historical/mapping/sumatran\_rhino\_22Jul2017\_9M7eS\_haploidified\_headersFixed\_Sc9M7eS\_2\_HRSCAF\_41/stats/bams\_merged\_index/JvS022\_05.merged.bam.stats.txt - results/historical/mapping/sumatran\_rhino\_22Jul2017\_9M7eS\_haploidified\_headersFixed\_Sc9M7eS\_2\_HRSCAF\_41/stats/bams\_merged\_index/JvS022\_06.merged.bam.stats.txt - results/historical/mapping/sumatran\_rhino\_22Jul2017\_9M7eS\_haploidified\_headersFixed\_Sc9M7eS\_2\_HRSCAF\_41/stats/bams\_merged\_index/JvS022\_07.merged.bam.stats.txt - results/historical/mapping/sumatran\_rhino\_22Jul2017\_9M7eS\_haploidified\_headersFixed\_Sc9M7eS\_2\_HRSCAF\_41/stats/bams\_merged\_index/JvS022\_08.merged.bam.stats.txt - results/historical/mapping/sumatran\_rhino\_22Jul2017\_9M7eS\_haploidified\_headersFixed\_Sc9M7eS\_2\_HRSCAF\_41/stats/bams\_merged\_index/JvS022\_09.merged.bam.stats.txt - results/historical/mapping/sumatran\_rhino\_22Jul2017\_9M7eS\_haploidified\_headersFixed\_Sc9M7eS\_2\_HRSCAF\_41/stats/bams\_merged\_index/JvS022\_10.merged.bam.stats.txt - results/historical/mapping/sumatran\_rhino\_22Jul2017\_9M7eS\_haploidified\_headersFixed\_Sc9M7eS\_2\_HRSCAF\_41/stats/bams\_merged\_index/JvS022\_11.merged.bam.stats.txt - results/historical/mapping/sumatran\_rhino\_22Jul2017\_9M7eS\_haploidified\_headersFixed\_Sc9M7eS\_2\_HRSCAF\_41/stats/bams\_merged\_index/JvS022\_12.merged.bam.stats.txt - results/historical/mapping/sumatran\_rhino\_22Jul2017\_9M7eS\_haploidified\_headersFixed\_Sc9M7eS\_2\_HRSCAF\_41/stats/bams\_merged\_index/JvS022\_74.merged.bam.stats.txt - results/historical/mapping/sumatran\_rhino\_22Jul2017\_9M7eS\_haploidified\_headersFixed\_Sc9M7eS\_2\_HRSCAF\_41/stats/bams\_merged\_index/JvS022\_75.merged.bam.stats.txt - results/historical/mapping/sumatran\_rhino\_22Jul2017\_9M7eS\_haploidified\_headersFixed\_Sc9M7eS\_2\_HRSCAF\_41/stats/bams\_merged\_index/JvS022\_76.merged.bam.stats.txt - results/historical/mapping/sumatran\_rhino\_22Jul2017\_9M7eS\_haploidified\_headersFixed\_Sc9M7eS\_2\_HRSCAF\_41/stats/bams\_merged\_index/JvS022\_77.merged.bam.stats.txt - results/historical/mapping/sumatran\_rhino\_22Jul2017\_9M7eS\_haploidified\_headersFixed\_Sc9M7eS\_2\_HRSCAF\_41/stats/bams\_merged\_index/JvS022\_78.merged.bam.stats.txt - results/historical/mapping/sumatran\_rhino\_22Jul2017\_9M7eS\_haploidified\_headersFixed\_Sc9M7eS\_2\_HRSCAF\_41/stats/bams\_merged\_index/JvS022\_79.merged.bam.stats.txt - results/historical/mapping/sumatran\_rhino\_22Jul2017\_9M7eS\_haploidified\_headersFixed\_Sc9M7eS\_2\_HRSCAF\_41/stats/bams\_merged\_index/JvS022\_80.merged.bam.stats.txt - results/historical/mapping/sumatran\_rhino\_22Jul2017\_9M7eS\_haploidified\_headersFixed\_Sc9M7eS\_2\_HRSCAF\_41/stats/bams\_merged\_index/JvS022\_81.merged.bam.stats.txt - results/historical/mapping/sumatran\_rhino\_22Jul2017\_9M7eS\_haploidified\_headersFixed\_Sc9M7eS\_2\_HRSCAF\_41/stats/bams\_merged\_index/JvS022\_82.merged.bam.stats.txt - results/historical/mapping/sumatran\_rhino\_22Jul2017\_9M7eS\_haploidified\_headersFixed\_Sc9M7eS\_2\_HRSCAF\_41/stats/bams\_merged\_index/JvS022\_83.merged.bam.stats.txt - results/historical/mapping/sumatran\_rhino\_22Jul2017\_9M7eS\_haploidified\_headersFixed\_Sc9M7eS\_2\_HRSCAF\_41/stats/bams\_merged\_index/JvS022\_84.merged.bam.stats.txt - results/historical/mapping/sumatran\_rhino\_22Jul2017\_9M7eS\_haploidified\_headersFixed\_Sc9M7eS\_2\_HRSCAF\_41/stats/bams\_merged\_index/JvS022\_85.merged.bam.stats.txt - results/historical/mapping/sumatran\_rhino\_22Jul2017\_9M7eS\_haploidified\_headersFixed\_Sc9M7eS\_2\_HRSCAF\_41/stats/bams\_merged\_index/JvS008\_08.merged.bam.qualimap/qualimapReport.html - results/historical/mapping/sumatran\_rhino\_22Jul2017\_9M7eS\_haploidified\_headersFixed\_Sc9M7eS\_2\_HRSCAF\_41/stats/bams\_merged\_index/JvS008\_10.merged.bam.qualimap/qualimapReport.html - results/historical/mapping/sumatran\_rhino\_22Jul2017\_9M7eS\_haploidified\_headersFixed\_Sc9M7eS\_2\_HRSCAF\_41/stats/bams\_merged\_index/JvS008\_11.merged.bam.qualimap/qualimapReport.html - results/historical/mapping/sumatran\_rhino\_22Jul2017\_9M7eS\_haploidified\_headersFixed\_Sc9M7eS\_2\_HRSCAF\_41/stats/bams\_merged\_index/JvS009\_09.merged.bam.qualimap/qualimapReport.html - results/historical/mapping/sumatran\_rhino\_22Jul2017\_9M7eS\_haploidified\_headersFixed\_Sc9M7eS\_2\_HRSCAF\_41/stats/bams\_merged\_index/JvS009\_15.merged.bam.qualimap/qualimapReport.html - results/historical/mapping/sumatran\_rhino\_22Jul2017\_9M7eS\_haploidified\_headersFixed\_Sc9M7eS\_2\_HRSCAF\_41/stats/bams\_merged\_index/JvS009\_19.merged.bam.qualimap/qualimapReport.html - results/historical/mapping/sumatran\_rhino\_22Jul2017\_9M7eS\_haploidified\_headersFixed\_Sc9M7eS\_2\_HRSCAF\_41/stats/bams\_merged\_index/JvS022\_01.merged.bam.qualimap/qualimapReport.html - results/historical/mapping/sumatran\_rhino\_22Jul2017\_9M7eS\_haploidified\_headersFixed\_Sc9M7eS\_2\_HRSCAF\_41/stats/bams\_merged\_index/JvS022\_02.merged.bam.qualimap/qualimapReport.html - results/historical/mapping/sumatran\_rhino\_22Jul2017\_9M7eS\_haploidified\_headersFixed\_Sc9M7eS\_2\_HRSCAF\_41/stats/bams\_merged\_index/JvS022\_03.merged.bam.qualimap/qualimapReport.html - results/historical/mapping/sumatran\_rhino\_22Jul2017\_9M7eS\_haploidified\_headersFixed\_Sc9M7eS\_2\_HRSCAF\_41/stats/bams\_merged\_index/JvS022\_04.merged.bam.qualimap/qualimapReport.html - results/historical/mapping/sumatran\_rhino\_22Jul2017\_9M7eS\_haploidified\_headersFixed\_Sc9M7eS\_2\_HRSCAF\_41/stats/bams\_merged\_index/JvS022\_05.merged.bam.qualimap/qualimapReport.html - results/historical/mapping/sumatran\_rhino\_22Jul2017\_9M7eS\_haploidified\_headersFixed\_Sc9M7eS\_2\_HRSCAF\_41/stats/bams\_merged\_index/JvS022\_06.merged.bam.qualimap/qualimapReport.html - results/historical/mapping/sumatran\_rhino\_22Jul2017\_9M7eS\_haploidified\_headersFixed\_Sc9M7eS\_2\_HRSCAF\_41/stats/bams\_merged\_index/JvS022\_07.merged.bam.qualimap/qualimapReport.html - results/historical/mapping/sumatran\_rhino\_22Jul2017\_9M7eS\_haploidified\_headersFixed\_Sc9M7eS\_2\_HRSCAF\_41/stats/bams\_merged\_index/JvS022\_08.merged.bam.qualimap/qualimapReport.html - results/historical/mapping/sumatran\_rhino\_22Jul2017\_9M7eS\_haploidified\_headersFixed\_Sc9M7eS\_2\_HRSCAF\_41/stats/bams\_merged\_index/JvS022\_09.merged.bam.qualimap/qualimapReport.html - results/historical/mapping/sumatran\_rhino\_22Jul2017\_9M7eS\_haploidified\_headersFixed\_Sc9M7eS\_2\_HRSCAF\_41/stats/bams\_merged\_index/JvS022\_10.merged.bam.qualimap/qualimapReport.html - results/historical/mapping/sumatran\_rhino\_22Jul2017\_9M7eS\_haploidified\_headersFixed\_Sc9M7eS\_2\_HRSCAF\_41/stats/bams\_merged\_index/JvS022\_11.merged.bam.qualimap/qualimapReport.html - results/historical/mapping/sumatran\_rhino\_22Jul2017\_9M7eS\_haploidified\_headersFixed\_Sc9M7eS\_2\_HRSCAF\_41/stats/bams\_merged\_index/JvS022\_12.merged.bam.qualimap/qualimapReport.html - results/historical/mapping/sumatran\_rhino\_22Jul2017\_9M7eS\_haploidified\_headersFixed\_Sc9M7eS\_2\_HRSCAF\_41/stats/bams\_merged\_index/JvS022\_74.merged.bam.qualimap/qualimapReport.html - results/historical/mapping/sumatran\_rhino\_22Jul2017\_9M7eS\_haploidified\_headersFixed\_Sc9M7eS\_2\_HRSCAF\_41/stats/bams\_merged\_index/JvS022\_75.merged.bam.qualimap/qualimapReport.html - results/historical/mapping/sumatran\_rhino\_22Jul2017\_9M7eS\_haploidified\_headersFixed\_Sc9M7eS\_2\_HRSCAF\_41/stats/bams\_merged\_index/JvS022\_76.merged.bam.qualimap/qualimapReport.html - results/historical/mapping/sumatran\_rhino\_22Jul2017\_9M7eS\_haploidified\_headersFixed\_Sc9M7eS\_2\_HRSCAF\_41/stats/bams\_merged\_index/JvS022\_77.merged.bam.qualimap/qualimapReport.html - results/historical/mapping/sumatran\_rhino\_22Jul2017\_9M7eS\_haploidified\_headersFixed\_Sc9M7eS\_2\_HRSCAF\_41/stats/bams\_merged\_index/JvS022\_78.merged.bam.qualimap/qualimapReport.html - results/historical/mapping/sumatran\_rhino\_22Jul2017\_9M7eS\_haploidified\_headersFixed\_Sc9M7eS\_2\_HRSCAF\_41/stats/bams\_merged\_index/JvS022\_79.merged.bam.qualimap/qualimapReport.html - results/historical/mapping/sumatran\_rhino\_22Jul2017\_9M7eS\_haploidified\_headersFixed\_Sc9M7eS\_2\_HRSCAF\_41/stats/bams\_merged\_index/JvS022\_80.merged.bam.qualimap/qualimapReport.html - results/historical/mapping/sumatran\_rhino\_22Jul2017\_9M7eS\_haploidified\_headersFixed\_Sc9M7eS\_2\_HRSCAF\_41/stats/bams\_merged\_index/JvS022\_81.merged.bam.qualimap/qualimapReport.html - results/historical/mapping/sumatran\_rhino\_22Jul2017\_9M7eS\_haploidified\_headersFixed\_Sc9M7eS\_2\_HRSCAF\_41/stats/bams\_merged\_index/JvS022\_82.merged.bam.qualimap/qualimapReport.html - results/historical/mapping/sumatran\_rhino\_22Jul2017\_9M7eS\_haploidified\_headersFixed\_Sc9M7eS\_2\_HRSCAF\_41/stats/bams\_merged\_index/JvS022\_83.merged.bam.qualimap/qualimapReport.html - results/historical/mapping/sumatran\_rhino\_22Jul2017\_9M7eS\_haploidified\_headersFixed\_Sc9M7eS\_2\_HRSCAF\_41/stats/bams\_merged\_index/JvS022\_84.merged.bam.qualimap/qualimapReport.html - results/historical/mapping/sumatran\_rhino\_22Jul2017\_9M7eS\_haploidified\_headersFixed\_Sc9M7eS\_2\_HRSCAF\_41/stats/bams\_merged\_index/JvS022\_85.merged.bam.qualimap/qualimapReport.html | |
| Output files | |
| - results/historical/mapping/sumatran\_rhino\_22Jul2017\_9M7eS\_haploidified\_headersFixed\_Sc9M7eS\_2\_HRSCAF\_41/stats/bams\_merged\_index/multiqc/multiqc\_report.html | |
| Container image | |
| docker://quay.io/biocontainers/multiqc:1.9--pyh9f0ad1d\_0 | |
| Code | |
| ``` 1 2 ``` \| ``` multiqc -f {params.indir} -o {params.outdir} 2> {log} ``` | |

##### Rule merged\_index\_bam\_stats

×

Rule properties

|  |  |
| --- | --- |
| Jobs | 33 |
| Input files | |
| - results/{dataset}/mapping/sumatran\_rhino\_22Jul2017\_9M7eS\_haploidified\_headersFixed\_Sc9M7eS\_2\_HRSCAF\_41/{sample}\_{index}.merged.bam - results/{dataset}/mapping/sumatran\_rhino\_22Jul2017\_9M7eS\_haploidified\_headersFixed\_Sc9M7eS\_2\_HRSCAF\_41/{sample}\_{index}.merged.bam.bai | |
| Output files | |
| - results/{dataset}/mapping/sumatran\_rhino\_22Jul2017\_9M7eS\_haploidified\_headersFixed\_Sc9M7eS\_2\_HRSCAF\_41/stats/bams\_merged\_index/{sample,[A-Za-z0-9]+}\_{index}.merged.bam.stats.txt | |
| Container image | |
| docker://biocontainers/samtools:v1.9-4-deb\_cv1 | |
| Code | |
| ``` 1 2 ``` \| ``` samtools flagstat {input.bam} > {output.stats} 2> {log} ``` | |

##### Rule merge\_historical\_bams\_per\_index

×

Rule properties

|  |  |
| --- | --- |
| Jobs | 30 |
| Input files | |
| Output files | |
| - results/historical/mapping/sumatran\_rhino\_22Jul2017\_9M7eS\_haploidified\_headersFixed\_Sc9M7eS\_2\_HRSCAF\_41/{sample,[A-Za-z0-9]+}\_{index}.merged.bam | |
| Container image | |
| docker://biocontainers/samtools:v1.9-4-deb\_cv1 | |
| Code | |
| ``` 1 2 3 4 5 6 7 8 9 ``` \| ``` files=`echo {input} \| awk '{{print NF}}'` if [ $files -gt 1 ] # check if there are at least 2 files for merging. If there is only one file, copy the sorted bam file. then samtools merge {output.merged} {input} 2> {log} else cp {input} {output.merged} && touch {output.merged} 2> {log} echo "Only one file present for merging. Copying the input bam file." >> {log} fi ``` | |

##### Rule index\_merged\_index\_bams

×

Rule properties

|  |  |
| --- | --- |
| Jobs | 33 |
| Input files | |
| - results/{dataset}/mapping/sumatran\_rhino\_22Jul2017\_9M7eS\_haploidified\_headersFixed\_Sc9M7eS\_2\_HRSCAF\_41/{sample}\_{index}.merged.bam | |
| Output files | |
| - results/{dataset}/mapping/sumatran\_rhino\_22Jul2017\_9M7eS\_haploidified\_headersFixed\_Sc9M7eS\_2\_HRSCAF\_41/{sample,[A-Za-z0-9]+}\_{index}.merged.bam.bai | |
| Container image | |
| docker://biocontainers/samtools:v1.9-4-deb\_cv1 | |
| Code | |
| ``` 1 2 ``` \| ``` samtools index {input.bam} {output.index} 2> {log} ``` | |

##### Rule merged\_index\_bam\_qualimap

×

Rule properties

|  |  |
| --- | --- |
| Jobs | 33 |
| Input files | |
| - results/{dataset}/mapping/sumatran\_rhino\_22Jul2017\_9M7eS\_haploidified\_headersFixed\_Sc9M7eS\_2\_HRSCAF\_41/{sample}\_{index}.merged.bam - results/{dataset}/mapping/sumatran\_rhino\_22Jul2017\_9M7eS\_haploidified\_headersFixed\_Sc9M7eS\_2\_HRSCAF\_41/{sample}\_{index}.merged.bam.bai | |
| Output files | |
| - results/{dataset}/mapping/sumatran\_rhino\_22Jul2017\_9M7eS\_haploidified\_headersFixed\_Sc9M7eS\_2\_HRSCAF\_41/stats/bams\_merged\_index/{sample,[A-Za-z0-9]+}\_{index}.merged.bam.qualimap/qualimapReport.html - results/{dataset}/mapping/sumatran\_rhino\_22Jul2017\_9M7eS\_haploidified\_headersFixed\_Sc9M7eS\_2\_HRSCAF\_41/stats/bams\_merged\_index/{sample,[A-Za-z0-9]+}\_{index}.merged.bam.qualimap/genome\_results.txt - results/{dataset}/mapping/sumatran\_rhino\_22Jul2017\_9M7eS\_haploidified\_headersFixed\_Sc9M7eS\_2\_HRSCAF\_41/stats/bams\_merged\_index/{sample,[A-Za-z0-9]+}\_{index}.merged.bam.qualimap | |
| Container image | |
| docker://quay.io/biocontainers/qualimap:2.2.2d--1 | |
| Code | |
| ``` 1 2 3 4 ``` \| ``` mem=$(((6 * {threads}) - 2)) unset DISPLAY qualimap bamqc -bam {input.bam} --java-mem-size=${{mem}}G -nt {threads} -outdir {params.outdir} -outformat html 2> {log} ``` | |

##### Rule historical\_rmdup\_bam\_multiqc

×

Rule properties

|  |  |
| --- | --- |
| Jobs | 1 |
| Input files | |
| - results/historical/mapping/sumatran\_rhino\_22Jul2017\_9M7eS\_haploidified\_headersFixed\_Sc9M7eS\_2\_HRSCAF\_41/stats/bams\_rmdup/JvS008\_08.merged.rmdup.bam.stats.txt - results/historical/mapping/sumatran\_rhino\_22Jul2017\_9M7eS\_haploidified\_headersFixed\_Sc9M7eS\_2\_HRSCAF\_41/stats/bams\_rmdup/JvS008\_10.merged.rmdup.bam.stats.txt - results/historical/mapping/sumatran\_rhino\_22Jul2017\_9M7eS\_haploidified\_headersFixed\_Sc9M7eS\_2\_HRSCAF\_41/stats/bams\_rmdup/JvS008\_11.merged.rmdup.bam.stats.txt - results/historical/mapping/sumatran\_rhino\_22Jul2017\_9M7eS\_haploidified\_headersFixed\_Sc9M7eS\_2\_HRSCAF\_41/stats/bams\_rmdup/JvS009\_09.merged.rmdup.bam.stats.txt - results/historical/mapping/sumatran\_rhino\_22Jul2017\_9M7eS\_haploidified\_headersFixed\_Sc9M7eS\_2\_HRSCAF\_41/stats/bams\_rmdup/JvS009\_15.merged.rmdup.bam.stats.txt - results/historical/mapping/sumatran\_rhino\_22Jul2017\_9M7eS\_haploidified\_headersFixed\_Sc9M7eS\_2\_HRSCAF\_41/stats/bams\_rmdup/JvS009\_19.merged.rmdup.bam.stats.txt - results/historical/mapping/sumatran\_rhino\_22Jul2017\_9M7eS\_haploidified\_headersFixed\_Sc9M7eS\_2\_HRSCAF\_41/stats/bams\_rmdup/JvS022\_01.merged.rmdup.bam.stats.txt - results/historical/mapping/sumatran\_rhino\_22Jul2017\_9M7eS\_haploidified\_headersFixed\_Sc9M7eS\_2\_HRSCAF\_41/stats/bams\_rmdup/JvS022\_02.merged.rmdup.bam.stats.txt - results/historical/mapping/sumatran\_rhino\_22Jul2017\_9M7eS\_haploidified\_headersFixed\_Sc9M7eS\_2\_HRSCAF\_41/stats/bams\_rmdup/JvS022\_03.merged.rmdup.bam.stats.txt - results/historical/mapping/sumatran\_rhino\_22Jul2017\_9M7eS\_haploidified\_headersFixed\_Sc9M7eS\_2\_HRSCAF\_41/stats/bams\_rmdup/JvS022\_04.merged.rmdup.bam.stats.txt - results/historical/mapping/sumatran\_rhino\_22Jul2017\_9M7eS\_haploidified\_headersFixed\_Sc9M7eS\_2\_HRSCAF\_41/stats/bams\_rmdup/JvS022\_05.merged.rmdup.bam.stats.txt - results/historical/mapping/sumatran\_rhino\_22Jul2017\_9M7eS\_haploidified\_headersFixed\_Sc9M7eS\_2\_HRSCAF\_41/stats/bams\_rmdup/JvS022\_06.merged.rmdup.bam.stats.txt - results/historical/mapping/sumatran\_rhino\_22Jul2017\_9M7eS\_haploidified\_headersFixed\_Sc9M7eS\_2\_HRSCAF\_41/stats/bams\_rmdup/JvS022\_07.merged.rmdup.bam.stats.txt - results/historical/mapping/sumatran\_rhino\_22Jul2017\_9M7eS\_haploidified\_headersFixed\_Sc9M7eS\_2\_HRSCAF\_41/stats/bams\_rmdup/JvS022\_08.merged.rmdup.bam.stats.txt - results/historical/mapping/sumatran\_rhino\_22Jul2017\_9M7eS\_haploidified\_headersFixed\_Sc9M7eS\_2\_HRSCAF\_41/stats/bams\_rmdup/JvS022\_09.merged.rmdup.bam.stats.txt - results/historical/mapping/sumatran\_rhino\_22Jul2017\_9M7eS\_haploidified\_headersFixed\_Sc9M7eS\_2\_HRSCAF\_41/stats/bams\_rmdup/JvS022\_10.merged.rmdup.bam.stats.txt - results/historical/mapping/sumatran\_rhino\_22Jul2017\_9M7eS\_haploidified\_headersFixed\_Sc9M7eS\_2\_HRSCAF\_41/stats/bams\_rmdup/JvS022\_11.merged.rmdup.bam.stats.txt - results/historical/mapping/sumatran\_rhino\_22Jul2017\_9M7eS\_haploidified\_headersFixed\_Sc9M7eS\_2\_HRSCAF\_41/stats/bams\_rmdup/JvS022\_12.merged.rmdup.bam.stats.txt - results/historical/mapping/sumatran\_rhino\_22Jul2017\_9M7eS\_haploidified\_headersFixed\_Sc9M7eS\_2\_HRSCAF\_41/stats/bams\_rmdup/JvS022\_74.merged.rmdup.bam.stats.txt - results/historical/mapping/sumatran\_rhino\_22Jul2017\_9M7eS\_haploidified\_headersFixed\_Sc9M7eS\_2\_HRSCAF\_41/stats/bams\_rmdup/JvS022\_75.merged.rmdup.bam.stats.txt - results/historical/mapping/sumatran\_rhino\_22Jul2017\_9M7eS\_haploidified\_headersFixed\_Sc9M7eS\_2\_HRSCAF\_41/stats/bams\_rmdup/JvS022\_76.merged.rmdup.bam.stats.txt - results/historical/mapping/sumatran\_rhino\_22Jul2017\_9M7eS\_haploidified\_headersFixed\_Sc9M7eS\_2\_HRSCAF\_41/stats/bams\_rmdup/JvS022\_77.merged.rmdup.bam.stats.txt - results/historical/mapping/sumatran\_rhino\_22Jul2017\_9M7eS\_haploidified\_headersFixed\_Sc9M7eS\_2\_HRSCAF\_41/stats/bams\_rmdup/JvS022\_78.merged.rmdup.bam.stats.txt - results/historical/mapping/sumatran\_rhino\_22Jul2017\_9M7eS\_haploidified\_headersFixed\_Sc9M7eS\_2\_HRSCAF\_41/stats/bams\_rmdup/JvS022\_79.merged.rmdup.bam.stats.txt - results/historical/mapping/sumatran\_rhino\_22Jul2017\_9M7eS\_haploidified\_headersFixed\_Sc9M7eS\_2\_HRSCAF\_41/stats/bams\_rmdup/JvS022\_80.merged.rmdup.bam.stats.txt - results/historical/mapping/sumatran\_rhino\_22Jul2017\_9M7eS\_haploidified\_headersFixed\_Sc9M7eS\_2\_HRSCAF\_41/stats/bams\_rmdup/JvS022\_81.merged.rmdup.bam.stats.txt - results/historical/mapping/sumatran\_rhino\_22Jul2017\_9M7eS\_haploidified\_headersFixed\_Sc9M7eS\_2\_HRSCAF\_41/stats/bams\_rmdup/JvS022\_82.merged.rmdup.bam.stats.txt - results/historical/mapping/sumatran\_rhino\_22Jul2017\_9M7eS\_haploidified\_headersFixed\_Sc9M7eS\_2\_HRSCAF\_41/stats/bams\_rmdup/JvS022\_83.merged.rmdup.bam.stats.txt - results/historical/mapping/sumatran\_rhino\_22Jul2017\_9M7eS\_haploidified\_headersFixed\_Sc9M7eS\_2\_HRSCAF\_41/stats/bams\_rmdup/JvS022\_84.merged.rmdup.bam.stats.txt - results/historical/mapping/sumatran\_rhino\_22Jul2017\_9M7eS\_haploidified\_headersFixed\_Sc9M7eS\_2\_HRSCAF\_41/stats/bams\_rmdup/JvS022\_85.merged.rmdup.bam.stats.txt - results/historical/mapping/sumatran\_rhino\_22Jul2017\_9M7eS\_haploidified\_headersFixed\_Sc9M7eS\_2\_HRSCAF\_41/stats/bams\_rmdup/JvS008\_08.merged.rmdup.bam.qualimap/qualimapReport.html - results/historical/mapping/sumatran\_rhino\_22Jul2017\_9M7eS\_haploidified\_headersFixed\_Sc9M7eS\_2\_HRSCAF\_41/stats/bams\_rmdup/JvS008\_10.merged.rmdup.bam.qualimap/qualimapReport.html - results/historical/mapping/sumatran\_rhino\_22Jul2017\_9M7eS\_haploidified\_headersFixed\_Sc9M7eS\_2\_HRSCAF\_41/stats/bams\_rmdup/JvS008\_11.merged.rmdup.bam.qualimap/qualimapReport.html - results/historical/mapping/sumatran\_rhino\_22Jul2017\_9M7eS\_haploidified\_headersFixed\_Sc9M7eS\_2\_HRSCAF\_41/stats/bams\_rmdup/JvS009\_09.merged.rmdup.bam.qualimap/qualimapReport.html - results/historical/mapping/sumatran\_rhino\_22Jul2017\_9M7eS\_haploidified\_headersFixed\_Sc9M7eS\_2\_HRSCAF\_41/stats/bams\_rmdup/JvS009\_15.merged.rmdup.bam.qualimap/qualimapReport.html - results/historical/mapping/sumatran\_rhino\_22Jul2017\_9M7eS\_haploidified\_headersFixed\_Sc9M7eS\_2\_HRSCAF\_41/stats/bams\_rmdup/JvS009\_19.merged.rmdup.bam.qualimap/qualimapReport.html - results/historical/mapping/sumatran\_rhino\_22Jul2017\_9M7eS\_haploidified\_headersFixed\_Sc9M7eS\_2\_HRSCAF\_41/stats/bams\_rmdup/JvS022\_01.merged.rmdup.bam.qualimap/qualimapReport.html - results/historical/mapping/sumatran\_rhino\_22Jul2017\_9M7eS\_haploidified\_headersFixed\_Sc9M7eS\_2\_HRSCAF\_41/stats/bams\_rmdup/JvS022\_02.merged.rmdup.bam.qualimap/qualimapReport.html - results/historical/mapping/sumatran\_rhino\_22Jul2017\_9M7eS\_haploidified\_headersFixed\_Sc9M7eS\_2\_HRSCAF\_41/stats/bams\_rmdup/JvS022\_03.merged.rmdup.bam.qualimap/qualimapReport.html - results/historical/mapping/sumatran\_rhino\_22Jul2017\_9M7eS\_haploidified\_headersFixed\_Sc9M7eS\_2\_HRSCAF\_41/stats/bams\_rmdup/JvS022\_04.merged.rmdup.bam.qualimap/qualimapReport.html - results/historical/mapping/sumatran\_rhino\_22Jul2017\_9M7eS\_haploidified\_headersFixed\_Sc9M7eS\_2\_HRSCAF\_41/stats/bams\_rmdup/JvS022\_05.merged.rmdup.bam.qualimap/qualimapReport.html - results/historical/mapping/sumatran\_rhino\_22Jul2017\_9M7eS\_haploidified\_headersFixed\_Sc9M7eS\_2\_HRSCAF\_41/stats/bams\_rmdup/JvS022\_06.merged.rmdup.bam.qualimap/qualimapReport.html - results/historical/mapping/sumatran\_rhino\_22Jul2017\_9M7eS\_haploidified\_headersFixed\_Sc9M7eS\_2\_HRSCAF\_41/stats/bams\_rmdup/JvS022\_07.merged.rmdup.bam.qualimap/qualimapReport.html - results/historical/mapping/sumatran\_rhino\_22Jul2017\_9M7eS\_haploidified\_headersFixed\_Sc9M7eS\_2\_HRSCAF\_41/stats/bams\_rmdup/JvS022\_08.merged.rmdup.bam.qualimap/qualimapReport.html - results/historical/mapping/sumatran\_rhino\_22Jul2017\_9M7eS\_haploidified\_headersFixed\_Sc9M7eS\_2\_HRSCAF\_41/stats/bams\_rmdup/JvS022\_09.merged.rmdup.bam.qualimap/qualimapReport.html - results/historical/mapping/sumatran\_rhino\_22Jul2017\_9M7eS\_haploidified\_headersFixed\_Sc9M7eS\_2\_HRSCAF\_41/stats/bams\_rmdup/JvS022\_10.merged.rmdup.bam.qualimap/qualimapReport.html - results/historical/mapping/sumatran\_rhino\_22Jul2017\_9M7eS\_haploidified\_headersFixed\_Sc9M7eS\_2\_HRSCAF\_41/stats/bams\_rmdup/JvS022\_11.merged.rmdup.bam.qualimap/qualimapReport.html - results/historical/mapping/sumatran\_rhino\_22Jul2017\_9M7eS\_haploidified\_headersFixed\_Sc9M7eS\_2\_HRSCAF\_41/stats/bams\_rmdup/JvS022\_12.merged.rmdup.bam.qualimap/qualimapReport.html - results/historical/mapping/sumatran\_rhino\_22Jul2017\_9M7eS\_haploidified\_headersFixed\_Sc9M7eS\_2\_HRSCAF\_41/stats/bams\_rmdup/JvS022\_74.merged.rmdup.bam.qualimap/qualimapReport.html - results/historical/mapping/sumatran\_rhino\_22Jul2017\_9M7eS\_haploidified\_headersFixed\_Sc9M7eS\_2\_HRSCAF\_41/stats/bams\_rmdup/JvS022\_75.merged.rmdup.bam.qualimap/qualimapReport.html - results/historical/mapping/sumatran\_rhino\_22Jul2017\_9M7eS\_haploidified\_headersFixed\_Sc9M7eS\_2\_HRSCAF\_41/stats/bams\_rmdup/JvS022\_76.merged.rmdup.bam.qualimap/qualimapReport.html - results/historical/mapping/sumatran\_rhino\_22Jul2017\_9M7eS\_haploidified\_headersFixed\_Sc9M7eS\_2\_HRSCAF\_41/stats/bams\_rmdup/JvS022\_77.merged.rmdup.bam.qualimap/qualimapReport.html - results/historical/mapping/sumatran\_rhino\_22Jul2017\_9M7eS\_haploidified\_headersFixed\_Sc9M7eS\_2\_HRSCAF\_41/stats/bams\_rmdup/JvS022\_78.merged.rmdup.bam.qualimap/qualimapReport.html - results/historical/mapping/sumatran\_rhino\_22Jul2017\_9M7eS\_haploidified\_headersFixed\_Sc9M7eS\_2\_HRSCAF\_41/stats/bams\_rmdup/JvS022\_79.merged.rmdup.bam.qualimap/qualimapReport.html - results/historical/mapping/sumatran\_rhino\_22Jul2017\_9M7eS\_haploidified\_headersFixed\_Sc9M7eS\_2\_HRSCAF\_41/stats/bams\_rmdup/JvS022\_80.merged.rmdup.bam.qualimap/qualimapReport.html - results/historical/mapping/sumatran\_rhino\_22Jul2017\_9M7eS\_haploidified\_headersFixed\_Sc9M7eS\_2\_HRSCAF\_41/stats/bams\_rmdup/JvS022\_81.merged.rmdup.bam.qualimap/qualimapReport.html - results/historical/mapping/sumatran\_rhino\_22Jul2017\_9M7eS\_haploidified\_headersFixed\_Sc9M7eS\_2\_HRSCAF\_41/stats/bams\_rmdup/JvS022\_82.merged.rmdup.bam.qualimap/qualimapReport.html - results/historical/mapping/sumatran\_rhino\_22Jul2017\_9M7eS\_haploidified\_headersFixed\_Sc9M7eS\_2\_HRSCAF\_41/stats/bams\_rmdup/JvS022\_83.merged.rmdup.bam.qualimap/qualimapReport.html - results/historical/mapping/sumatran\_rhino\_22Jul2017\_9M7eS\_haploidified\_headersFixed\_Sc9M7eS\_2\_HRSCAF\_41/stats/bams\_rmdup/JvS022\_84.merged.rmdup.bam.qualimap/qualimapReport.html - results/historical/mapping/sumatran\_rhino\_22Jul2017\_9M7eS\_haploidified\_headersFixed\_Sc9M7eS\_2\_HRSCAF\_41/stats/bams\_rmdup/JvS022\_85.merged.rmdup.bam.qualimap/qualimapReport.html | |
| Output files | |
| - results/historical/mapping/sumatran\_rhino\_22Jul2017\_9M7eS\_haploidified\_headersFixed\_Sc9M7eS\_2\_HRSCAF\_41/stats/bams\_rmdup/multiqc/multiqc\_report.html | |
| Container image | |
| docker://quay.io/biocontainers/multiqc:1.9--pyh9f0ad1d\_0 | |
| Code | |
| ``` 1 2 ``` \| ``` multiqc -f {params.indir} -o {params.outdir} 2> {log} ``` | |

##### Rule rmdup\_bam\_stats

×

Rule properties

|  |  |
| --- | --- |
| Jobs | 33 |
| Input files | |
| - results/{dataset}/mapping/sumatran\_rhino\_22Jul2017\_9M7eS\_haploidified\_headersFixed\_Sc9M7eS\_2\_HRSCAF\_41/{sample}\_{index}.merged.rmdup.bam - results/{dataset}/mapping/sumatran\_rhino\_22Jul2017\_9M7eS\_haploidified\_headersFixed\_Sc9M7eS\_2\_HRSCAF\_41/{sample}\_{index}.merged.rmdup.bam.bai | |
| Output files | |
| - results/{dataset}/mapping/sumatran\_rhino\_22Jul2017\_9M7eS\_haploidified\_headersFixed\_Sc9M7eS\_2\_HRSCAF\_41/stats/bams\_rmdup/{sample,[A-Za-z0-9]+}\_{index}.merged.rmdup.bam.stats.txt | |
| Container image | |
| docker://biocontainers/samtools:v1.9-4-deb\_cv1 | |
| Code | |
| ``` 1 2 ``` \| ``` samtools flagstat {input.bam} > {output.stats} 2> {log} ``` | |

##### Rule rmdup\_historical\_bams

×

Rule properties

|  |  |
| --- | --- |
| Jobs | 30 |
| Input files | |
| - results/historical/mapping/sumatran\_rhino\_22Jul2017\_9M7eS\_haploidified\_headersFixed\_Sc9M7eS\_2\_HRSCAF\_41/{sample}\_{index}.merged.bam - results/historical/mapping/sumatran\_rhino\_22Jul2017\_9M7eS\_haploidified\_headersFixed\_Sc9M7eS\_2\_HRSCAF\_41/{sample}\_{index}.merged.bam.bai | |
| Output files | |
| - results/historical/mapping/sumatran\_rhino\_22Jul2017\_9M7eS\_haploidified\_headersFixed\_Sc9M7eS\_2\_HRSCAF\_41/{sample,[A-Za-z0-9]+}\_{index}.merged.rmdup.bam | |
| Container image | |
| docker://biocontainers/samtools:v1.9-4-deb\_cv1 | |
| Code | |
| ``` 1 2 ``` \| ``` samtools view -@ {threads} -h {input.merged} \| python3 workflow/scripts/samremovedup.py \| samtools view -b -o {output.rmdup} 2> {log} ``` | |

##### Rule index\_rmdup\_bams

×

Rule properties

|  |  |
| --- | --- |
| Jobs | 33 |
| Input files | |
| - results/{dataset}/mapping/sumatran\_rhino\_22Jul2017\_9M7eS\_haploidified\_headersFixed\_Sc9M7eS\_2\_HRSCAF\_41/{sample}\_{index}.merged.rmdup.bam | |
| Output files | |
| - results/{dataset}/mapping/sumatran\_rhino\_22Jul2017\_9M7eS\_haploidified\_headersFixed\_Sc9M7eS\_2\_HRSCAF\_41/{sample,[A-Za-z0-9]+}\_{index}.merged.rmdup.bam.bai | |
| Container image | |
| docker://biocontainers/samtools:v1.9-4-deb\_cv1 | |
| Code | |
| ``` 1 2 ``` \| ``` samtools index {input.bam} {output.index} 2> {log} ``` | |

##### Rule rmdup\_bam\_qualimap

×

Rule properties

|  |  |
| --- | --- |
| Jobs | 33 |
| Input files | |
| - results/{dataset}/mapping/sumatran\_rhino\_22Jul2017\_9M7eS\_haploidified\_headersFixed\_Sc9M7eS\_2\_HRSCAF\_41/{sample}\_{index}.merged.rmdup.bam - results/{dataset}/mapping/sumatran\_rhino\_22Jul2017\_9M7eS\_haploidified\_headersFixed\_Sc9M7eS\_2\_HRSCAF\_41/{sample}\_{index}.merged.rmdup.bam.bai | |
| Output files | |
| - results/{dataset}/mapping/sumatran\_rhino\_22Jul2017\_9M7eS\_haploidified\_headersFixed\_Sc9M7eS\_2\_HRSCAF\_41/stats/bams\_rmdup/{sample,[A-Za-z0-9]+}\_{index}.merged.rmdup.bam.qualimap/qualimapReport.html - results/{dataset}/mapping/sumatran\_rhino\_22Jul2017\_9M7eS\_haploidified\_headersFixed\_Sc9M7eS\_2\_HRSCAF\_41/stats/bams\_rmdup/{sample,[A-Za-z0-9]+}\_{index}.merged.rmdup.bam.qualimap/genome\_results.txt - results/{dataset}/mapping/sumatran\_rhino\_22Jul2017\_9M7eS\_haploidified\_headersFixed\_Sc9M7eS\_2\_HRSCAF\_41/stats/bams\_rmdup/{sample,[A-Za-z0-9]+}\_{index}.merged.rmdup.bam.qualimap | |
| Container image | |
| docker://quay.io/biocontainers/qualimap:2.2.2d--1 | |
| Code | |
| ``` 1 2 3 4 ``` \| ``` mem=$(((6 * {threads}) - 2)) unset DISPLAY qualimap bamqc -bam {input.bam} --java-mem-size=${{mem}}G -nt {threads} -outdir {params.outdir} -outformat html 2> {log} ``` | |

##### Rule historical\_merged\_sample\_bam\_multiqc

×

Rule properties

|  |  |
| --- | --- |
| Jobs | 1 |
| Input files | |
| - results/historical/mapping/sumatran\_rhino\_22Jul2017\_9M7eS\_haploidified\_headersFixed\_Sc9M7eS\_2\_HRSCAF\_41/stats/bams\_merged\_sample/JvS008.merged.rmdup.merged.bam.stats.txt - results/historical/mapping/sumatran\_rhino\_22Jul2017\_9M7eS\_haploidified\_headersFixed\_Sc9M7eS\_2\_HRSCAF\_41/stats/bams\_merged\_sample/JvS009.merged.rmdup.merged.bam.stats.txt - results/historical/mapping/sumatran\_rhino\_22Jul2017\_9M7eS\_haploidified\_headersFixed\_Sc9M7eS\_2\_HRSCAF\_41/stats/bams\_merged\_sample/JvS022.merged.rmdup.merged.bam.stats.txt - results/historical/mapping/sumatran\_rhino\_22Jul2017\_9M7eS\_haploidified\_headersFixed\_Sc9M7eS\_2\_HRSCAF\_41/stats/bams\_merged\_sample/JvS008.merged.rmdup.merged.bam.qualimap/qualimapReport.html - results/historical/mapping/sumatran\_rhino\_22Jul2017\_9M7eS\_haploidified\_headersFixed\_Sc9M7eS\_2\_HRSCAF\_41/stats/bams\_merged\_sample/JvS009.merged.rmdup.merged.bam.qualimap/qualimapReport.html - results/historical/mapping/sumatran\_rhino\_22Jul2017\_9M7eS\_haploidified\_headersFixed\_Sc9M7eS\_2\_HRSCAF\_41/stats/bams\_merged\_sample/JvS022.merged.rmdup.merged.bam.qualimap/qualimapReport.html | |
| Output files | |
| - results/historical/mapping/sumatran\_rhino\_22Jul2017\_9M7eS\_haploidified\_headersFixed\_Sc9M7eS\_2\_HRSCAF\_41/stats/bams\_merged\_sample/multiqc/multiqc\_report.html | |
| Container image | |
| docker://quay.io/biocontainers/multiqc:1.9--pyh9f0ad1d\_0 | |
| Code | |
| ``` 1 2 ``` \| ``` multiqc -f {params.indir} -o {params.outdir} 2> {log} ``` | |

##### Rule merged\_sample\_bam\_stats

×

Rule properties

|  |  |
| --- | --- |
| Jobs | 6 |
| Input files | |
| - results/{dataset}/mapping/sumatran\_rhino\_22Jul2017\_9M7eS\_haploidified\_headersFixed\_Sc9M7eS\_2\_HRSCAF\_41/{sample}.merged.rmdup.merged.bam - results/{dataset}/mapping/sumatran\_rhino\_22Jul2017\_9M7eS\_haploidified\_headersFixed\_Sc9M7eS\_2\_HRSCAF\_41/{sample}.merged.rmdup.merged.bam.bai | |
| Output files | |
| - results/{dataset}/mapping/sumatran\_rhino\_22Jul2017\_9M7eS\_haploidified\_headersFixed\_Sc9M7eS\_2\_HRSCAF\_41/stats/bams\_merged\_sample/{sample,[A-Za-z0-9]+}.merged.rmdup.merged.bam.stats.txt | |
| Container image | |
| docker://biocontainers/samtools:v1.9-4-deb\_cv1 | |
| Code | |
| ``` 1 2 ``` \| ``` samtools flagstat {input.bam} > {output.stats} 2> {log} ``` | |

##### Rule merge\_historical\_bams\_per\_sample

×

Rule properties

|  |  |
| --- | --- |
| Jobs | 3 |
| Input files | |
| Output files | |
| - results/historical/mapping/sumatran\_rhino\_22Jul2017\_9M7eS\_haploidified\_headersFixed\_Sc9M7eS\_2\_HRSCAF\_41/{sample,[A-Za-z0-9]+}.merged.rmdup.merged.bam | |
| Container image | |
| docker://biocontainers/samtools:v1.9-4-deb\_cv1 | |
| Code | |
| ``` 1 2 3 4 5 6 7 8 9 ``` \| ``` files=`echo {input} \| awk '{{print NF}}'` if [ $files -gt 1 ] # check if there are at least 2 files for merging. If there is only one file, copy the sorted bam file. then samtools merge {output.merged} {input} 2> {log} else cp {input} {output.merged} && touch {output.merged} 2> {log} echo "Only one file present for merging. Copying the input bam file." >> {log} fi ``` | |

##### Rule index\_merged\_sample\_bams

×

Rule properties

|  |  |
| --- | --- |
| Jobs | 6 |
| Input files | |
| - results/{dataset}/mapping/sumatran\_rhino\_22Jul2017\_9M7eS\_haploidified\_headersFixed\_Sc9M7eS\_2\_HRSCAF\_41/{sample}.merged.rmdup.merged.bam | |
| Output files | |
| - results/{dataset}/mapping/sumatran\_rhino\_22Jul2017\_9M7eS\_haploidified\_headersFixed\_Sc9M7eS\_2\_HRSCAF\_41/{sample,[A-Za-z0-9]+}.merged.rmdup.merged.bam.bai | |
| Container image | |
| docker://biocontainers/samtools:v1.9-4-deb\_cv1 | |
| Code | |
| ``` 1 2 ``` \| ``` samtools index {input.bam} {output.index} 2> {log} ``` | |

##### Rule merged\_sample\_bam\_qualimap

×

Rule properties

|  |  |
| --- | --- |
| Jobs | 6 |
| Input files | |
| - results/{dataset}/mapping/sumatran\_rhino\_22Jul2017\_9M7eS\_haploidified\_headersFixed\_Sc9M7eS\_2\_HRSCAF\_41/{sample}.merged.rmdup.merged.bam - results/{dataset}/mapping/sumatran\_rhino\_22Jul2017\_9M7eS\_haploidified\_headersFixed\_Sc9M7eS\_2\_HRSCAF\_41/{sample}.merged.rmdup.merged.bam.bai | |
| Output files | |
| - results/{dataset}/mapping/sumatran\_rhino\_22Jul2017\_9M7eS\_haploidified\_headersFixed\_Sc9M7eS\_2\_HRSCAF\_41/stats/bams\_merged\_sample/{sample,[A-Za-z0-9]+}.merged.rmdup.merged.bam.qualimap/qualimapReport.html - results/{dataset}/mapping/sumatran\_rhino\_22Jul2017\_9M7eS\_haploidified\_headersFixed\_Sc9M7eS\_2\_HRSCAF\_41/stats/bams\_merged\_sample/{sample,[A-Za-z0-9]+}.merged.rmdup.merged.bam.qualimap/genome\_results.txt - results/{dataset}/mapping/sumatran\_rhino\_22Jul2017\_9M7eS\_haploidified\_headersFixed\_Sc9M7eS\_2\_HRSCAF\_41/stats/bams\_merged\_sample/{sample,[A-Za-z0-9]+}.merged.rmdup.merged.bam.qualimap | |
| Container image | |
| docker://quay.io/biocontainers/qualimap:2.2.2d--1 | |
| Code | |
| ``` 1 2 3 4 ``` \| ``` mem=$(((6 * {threads}) - 2)) unset DISPLAY qualimap bamqc -bam {input.bam} --java-mem-size=${{mem}}G -nt {threads} -outdir {params.outdir} -outformat html 2> {log} ``` | |

##### Rule historical\_realigned\_bam\_multiqc

×

Rule properties

|  |  |
| --- | --- |
| Jobs | 1 |
| Input files | |
| - results/historical/mapping/sumatran\_rhino\_22Jul2017\_9M7eS\_haploidified\_headersFixed\_Sc9M7eS\_2\_HRSCAF\_41/stats/bams\_indels\_realigned/JvS008.merged.rmdup.merged.realn.repma.Q30.bam.dp.hist.pdf - results/historical/mapping/sumatran\_rhino\_22Jul2017\_9M7eS\_haploidified\_headersFixed\_Sc9M7eS\_2\_HRSCAF\_41/stats/bams\_indels\_realigned/JvS009.merged.rmdup.merged.realn.repma.Q30.bam.dp.hist.pdf - results/historical/mapping/sumatran\_rhino\_22Jul2017\_9M7eS\_haploidified\_headersFixed\_Sc9M7eS\_2\_HRSCAF\_41/stats/bams\_indels\_realigned/JvS022.merged.rmdup.merged.realn.repma.Q30.bam.dp.hist.pdf - results/historical/mapping/sumatran\_rhino\_22Jul2017\_9M7eS\_haploidified\_headersFixed\_Sc9M7eS\_2\_HRSCAF\_41/stats/bams\_indels\_realigned/JvS008.merged.rmdup.merged.realn.bam.stats.txt - results/historical/mapping/sumatran\_rhino\_22Jul2017\_9M7eS\_haploidified\_headersFixed\_Sc9M7eS\_2\_HRSCAF\_41/stats/bams\_indels\_realigned/JvS009.merged.rmdup.merged.realn.bam.stats.txt - results/historical/mapping/sumatran\_rhino\_22Jul2017\_9M7eS\_haploidified\_headersFixed\_Sc9M7eS\_2\_HRSCAF\_41/stats/bams\_indels\_realigned/JvS022.merged.rmdup.merged.realn.bam.stats.txt - results/historical/mapping/sumatran\_rhino\_22Jul2017\_9M7eS\_haploidified\_headersFixed\_Sc9M7eS\_2\_HRSCAF\_41/stats/bams\_indels\_realigned/JvS008.merged.rmdup.merged.realn.bam.qualimap/qualimapReport.html - results/historical/mapping/sumatran\_rhino\_22Jul2017\_9M7eS\_haploidified\_headersFixed\_Sc9M7eS\_2\_HRSCAF\_41/stats/bams\_indels\_realigned/JvS009.merged.rmdup.merged.realn.bam.qualimap/qualimapReport.html - results/historical/mapping/sumatran\_rhino\_22Jul2017\_9M7eS\_haploidified\_headersFixed\_Sc9M7eS\_2\_HRSCAF\_41/stats/bams\_indels\_realigned/JvS022.merged.rmdup.merged.realn.bam.qualimap/qualimapReport.html - results/historical/mapping/sumatran\_rhino\_22Jul2017\_9M7eS\_haploidified\_headersFixed\_Sc9M7eS\_2\_HRSCAF\_41/stats/bams\_indels\_realigned/fastqc/JvS008.merged.rmdup.merged.realn\_fastqc.html - results/historical/mapping/sumatran\_rhino\_22Jul2017\_9M7eS\_haploidified\_headersFixed\_Sc9M7eS\_2\_HRSCAF\_41/stats/bams\_indels\_realigned/fastqc/JvS009.merged.rmdup.merged.realn\_fastqc.html - results/historical/mapping/sumatran\_rhino\_22Jul2017\_9M7eS\_haploidified\_headersFixed\_Sc9M7eS\_2\_HRSCAF\_41/stats/bams\_indels\_realigned/fastqc/JvS022.merged.rmdup.merged.realn\_fastqc.html - results/historical/mapping/sumatran\_rhino\_22Jul2017\_9M7eS\_haploidified\_headersFixed\_Sc9M7eS\_2\_HRSCAF\_41/stats/bams\_indels\_realigned/fastqc/JvS008.merged.rmdup.merged.realn\_fastqc.zip - results/historical/mapping/sumatran\_rhino\_22Jul2017\_9M7eS\_haploidified\_headersFixed\_Sc9M7eS\_2\_HRSCAF\_41/stats/bams\_indels\_realigned/fastqc/JvS009.merged.rmdup.merged.realn\_fastqc.zip - results/historical/mapping/sumatran\_rhino\_22Jul2017\_9M7eS\_haploidified\_headersFixed\_Sc9M7eS\_2\_HRSCAF\_41/stats/bams\_indels\_realigned/fastqc/JvS022.merged.rmdup.merged.realn\_fastqc.zip - results/historical/mapping/sumatran\_rhino\_22Jul2017\_9M7eS\_haploidified\_headersFixed\_Sc9M7eS\_2\_HRSCAF\_41/stats/bams\_indels\_realigned/JvS008.merged.rmdup.merged.realn.repma.Q30.bam.dp.hist.pdf - results/historical/mapping/sumatran\_rhino\_22Jul2017\_9M7eS\_haploidified\_headersFixed\_Sc9M7eS\_2\_HRSCAF\_41/stats/bams\_indels\_realigned/JvS009.merged.rmdup.merged.realn.repma.Q30.bam.dp.hist.pdf - results/historical/mapping/sumatran\_rhino\_22Jul2017\_9M7eS\_haploidified\_headersFixed\_Sc9M7eS\_2\_HRSCAF\_41/stats/bams\_indels\_realigned/JvS022.merged.rmdup.merged.realn.repma.Q30.bam.dp.hist.pdf | |
| Output files | |
| - results/historical/mapping/sumatran\_rhino\_22Jul2017\_9M7eS\_haploidified\_headersFixed\_Sc9M7eS\_2\_HRSCAF\_41/stats/bams\_indels\_realigned/multiqc/multiqc\_report.html | |
| Container image | |
| docker://quay.io/biocontainers/multiqc:1.9--pyh9f0ad1d\_0 | |
| Code | |
| ``` 1 2 ``` \| ``` multiqc -f {params.indir} -o {params.outdir} 2> {log} ``` | |

##### Rule plot\_dp\_hist

×

Rule properties

|  |  |
| --- | --- |
| Jobs | 6 |
| Input files | |
| - results/{dataset}/mapping/sumatran\_rhino\_22Jul2017\_9M7eS\_haploidified\_headersFixed\_Sc9M7eS\_2\_HRSCAF\_41/stats/bams\_indels\_realigned/{sample}.merged.rmdup.merged.realn.repma.Q30.bam.dpstats.txt - results/{dataset}/mapping/sumatran\_rhino\_22Jul2017\_9M7eS\_haploidified\_headersFixed\_Sc9M7eS\_2\_HRSCAF\_41/stats/bams\_indels\_realigned/{sample}.merged.rmdup.merged.realn.repma.Q30.bam.dp | |
| Output files | |
| - results/{dataset}/mapping/sumatran\_rhino\_22Jul2017\_9M7eS\_haploidified\_headersFixed\_Sc9M7eS\_2\_HRSCAF\_41/stats/bams\_indels\_realigned/{sample,[A-Za-z0-9]+}.merged.rmdup.merged.realn.repma.Q30.bam.dp.hist.pdf | |

##### Rule realigned\_bam\_depth

×

Rule properties

|  |  |
| --- | --- |
| Jobs | 6 |
| Input files | |
| - results/{dataset}/mapping/sumatran\_rhino\_22Jul2017\_9M7eS\_haploidified\_headersFixed\_Sc9M7eS\_2\_HRSCAF\_41/{sample}.merged.rmdup.merged.realn.bam - /proj/sllstore2017093/b2016342/b2016342\_nobackup/lts/genome\_erosion\_pipeline/verena\_testing/testdata/reference/sumatran\_rhino\_22Jul2017\_9M7eS\_haploidified\_headersFixed\_Sc9M7eS\_2\_HRSCAF\_41.repma.bed | |
| Output files | |
| - results/{dataset}/mapping/sumatran\_rhino\_22Jul2017\_9M7eS\_haploidified\_headersFixed\_Sc9M7eS\_2\_HRSCAF\_41/stats/bams\_indels\_realigned/{sample,[A-Za-z0-9]+}.merged.rmdup.merged.realn.repma.Q30.bam.dp - results/{dataset}/mapping/sumatran\_rhino\_22Jul2017\_9M7eS\_haploidified\_headersFixed\_Sc9M7eS\_2\_HRSCAF\_41/stats/bams\_indels\_realigned/{sample,[A-Za-z0-9]+}.merged.rmdup.merged.realn.repma.Q30.bam.dpstats.txt | |
| Container image | |
| docker://biocontainers/samtools:v1.9-4-deb\_cv1 | |
| Code | |
| ``` 1 2 3 4 5 6 7 8 9 10 ``` \| ``` if [ {params.cov} = "True" ] # include sites with missing data / zero coverage then samtools depth -a -Q 30 -q 30 -b {input.no_rep_bed} {input.bam} > {output.tmp} 2> {log} && awk '{{sum+=$3}} END {{ print sum/NR }}' {output.tmp} \| awk -v min={params.minDP} -v max={params.maxDP} '{{ printf "%.0f %.0f %.0f", $1, $1*min, $1*max }}' > {output.dp} 2>> {log} elif [ {params.cov} = "False" ] # exclude sites with missing data / zero coverage then samtools depth -Q 30 -q 30 -b {input.no_rep_bed} {input.bam} > {output.tmp} 2> {log} && awk '{{sum+=$3}} END {{ print sum/NR }}' {output.tmp} \| awk -v min={params.minDP} -v max={params.maxDP} '{{ printf "%.0f %.0f %.0f", $1, $1*min, $1*max }}' > {output.dp} 2>> {log} fi ``` | |

##### Rule indel\_realigner

×

Rule properties

|  |  |
| --- | --- |
| Jobs | 6 |
| Input files | |
| - /proj/sllstore2017093/b2016342/b2016342\_nobackup/lts/genome\_erosion\_pipeline/verena\_testing/testdata/reference/sumatran\_rhino\_22Jul2017\_9M7eS\_haploidified\_headersFixed\_Sc9M7eS\_2\_HRSCAF\_41.fasta - /proj/sllstore2017093/b2016342/b2016342\_nobackup/lts/genome\_erosion\_pipeline/verena\_testing/testdata/reference/sumatran\_rhino\_22Jul2017\_9M7eS\_haploidified\_headersFixed\_Sc9M7eS\_2\_HRSCAF\_41.dict - /proj/sllstore2017093/b2016342/b2016342\_nobackup/lts/genome\_erosion\_pipeline/verena\_testing/testdata/reference/sumatran\_rhino\_22Jul2017\_9M7eS\_haploidified\_headersFixed\_Sc9M7eS\_2\_HRSCAF\_41.fasta.fai - results/{dataset}/mapping/sumatran\_rhino\_22Jul2017\_9M7eS\_haploidified\_headersFixed\_Sc9M7eS\_2\_HRSCAF\_41/{sample}.merged.rmdup.merged.bam - results/{dataset}/mapping/sumatran\_rhino\_22Jul2017\_9M7eS\_haploidified\_headersFixed\_Sc9M7eS\_2\_HRSCAF\_41/{sample}.merged.rmdup.merged.bam.bai - results/{dataset}/mapping/sumatran\_rhino\_22Jul2017\_9M7eS\_haploidified\_headersFixed\_Sc9M7eS\_2\_HRSCAF\_41/{sample}.merged.rmdup.merged.realn\_targets.list | |
| Output files | |
| - results/{dataset}/mapping/sumatran\_rhino\_22Jul2017\_9M7eS\_haploidified\_headersFixed\_Sc9M7eS\_2\_HRSCAF\_41/{sample,[A-Za-z0-9]+}.merged.rmdup.merged.realn.bam | |
| Container image | |
| docker://broadinstitute/gatk3:3.7-0 | |
| Code | |
| ``` 1 2 ``` \| ``` java -jar /usr/GenomeAnalysisTK.jar -T IndelRealigner -R {input.ref} -I {input.bam} -targetIntervals {input.target_list} -o {output.realigned} 2> {log} ``` | |

##### Rule indel\_realigner\_targets

×

Rule properties

|  |  |
| --- | --- |
| Jobs | 6 |
| Input files | |
| - /proj/sllstore2017093/b2016342/b2016342\_nobackup/lts/genome\_erosion\_pipeline/verena\_testing/testdata/reference/sumatran\_rhino\_22Jul2017\_9M7eS\_haploidified\_headersFixed\_Sc9M7eS\_2\_HRSCAF\_41.fasta - /proj/sllstore2017093/b2016342/b2016342\_nobackup/lts/genome\_erosion\_pipeline/verena\_testing/testdata/reference/sumatran\_rhino\_22Jul2017\_9M7eS\_haploidified\_headersFixed\_Sc9M7eS\_2\_HRSCAF\_41.dict - /proj/sllstore2017093/b2016342/b2016342\_nobackup/lts/genome\_erosion\_pipeline/verena\_testing/testdata/reference/sumatran\_rhino\_22Jul2017\_9M7eS\_haploidified\_headersFixed\_Sc9M7eS\_2\_HRSCAF\_41.fasta.fai - results/{dataset}/mapping/sumatran\_rhino\_22Jul2017\_9M7eS\_haploidified\_headersFixed\_Sc9M7eS\_2\_HRSCAF\_41/{sample}.merged.rmdup.merged.bam - results/{dataset}/mapping/sumatran\_rhino\_22Jul2017\_9M7eS\_haploidified\_headersFixed\_Sc9M7eS\_2\_HRSCAF\_41/{sample}.merged.rmdup.merged.bam.bai | |
| Output files | |
| - results/{dataset}/mapping/sumatran\_rhino\_22Jul2017\_9M7eS\_haploidified\_headersFixed\_Sc9M7eS\_2\_HRSCAF\_41/{sample,[A-Za-z0-9]+}.merged.rmdup.merged.realn\_targets.list | |
| Container image | |
| docker://broadinstitute/gatk3:3.7-0 | |
| Code | |
| ``` 1 2 ``` \| ``` java -jar /usr/GenomeAnalysisTK.jar -T RealignerTargetCreator -R {input.ref} -I {input.bam} -o {output.target_list} -nt {threads} 2> {log} ``` | |

##### Rule realigned\_bam\_stats

×

Rule properties

|  |  |
| --- | --- |
| Jobs | 6 |
| Input files | |
| - results/{dataset}/mapping/sumatran\_rhino\_22Jul2017\_9M7eS\_haploidified\_headersFixed\_Sc9M7eS\_2\_HRSCAF\_41/{sample}.merged.rmdup.merged.realn.bam - results/{dataset}/mapping/sumatran\_rhino\_22Jul2017\_9M7eS\_haploidified\_headersFixed\_Sc9M7eS\_2\_HRSCAF\_41/{sample}.merged.rmdup.merged.realn.bam.bai | |
| Output files | |
| - results/{dataset}/mapping/sumatran\_rhino\_22Jul2017\_9M7eS\_haploidified\_headersFixed\_Sc9M7eS\_2\_HRSCAF\_41/stats/bams\_indels\_realigned/{sample,[A-Za-z0-9]+}.merged.rmdup.merged.realn.bam.stats.txt | |
| Container image | |
| docker://biocontainers/samtools:v1.9-4-deb\_cv1 | |
| Code | |
| ``` 1 2 ``` \| ``` samtools flagstat {input.bam} > {output.stats} 2> {log} ``` | |

##### Rule index\_realigned\_bams

×

Rule properties

|  |  |
| --- | --- |
| Jobs | 6 |
| Input files | |
| - results/{dataset}/mapping/sumatran\_rhino\_22Jul2017\_9M7eS\_haploidified\_headersFixed\_Sc9M7eS\_2\_HRSCAF\_41/{sample}.merged.rmdup.merged.bam | |
| Output files | |
| - results/{dataset}/mapping/sumatran\_rhino\_22Jul2017\_9M7eS\_haploidified\_headersFixed\_Sc9M7eS\_2\_HRSCAF\_41/{sample,[A-Za-z0-9]+}.merged.rmdup.merged.realn.bam.bai | |
| Container image | |
| docker://biocontainers/samtools:v1.9-4-deb\_cv1 | |
| Code | |
| ``` 1 2 ``` \| ``` samtools index {input.bam} {output.index} 2> {log} ``` | |

##### Rule realigned\_bam\_qualimap

×

Rule properties

|  |  |
| --- | --- |
| Jobs | 6 |
| Input files | |
| - results/{dataset}/mapping/sumatran\_rhino\_22Jul2017\_9M7eS\_haploidified\_headersFixed\_Sc9M7eS\_2\_HRSCAF\_41/{sample}.merged.rmdup.merged.realn.bam - results/{dataset}/mapping/sumatran\_rhino\_22Jul2017\_9M7eS\_haploidified\_headersFixed\_Sc9M7eS\_2\_HRSCAF\_41/{sample}.merged.rmdup.merged.realn.bam.bai | |
| Output files | |
| - results/{dataset}/mapping/sumatran\_rhino\_22Jul2017\_9M7eS\_haploidified\_headersFixed\_Sc9M7eS\_2\_HRSCAF\_41/stats/bams\_indels\_realigned/{sample,[A-Za-z0-9]+}.merged.rmdup.merged.realn.bam.qualimap/qualimapReport.html - results/{dataset}/mapping/sumatran\_rhino\_22Jul2017\_9M7eS\_haploidified\_headersFixed\_Sc9M7eS\_2\_HRSCAF\_41/stats/bams\_indels\_realigned/{sample,[A-Za-z0-9]+}.merged.rmdup.merged.realn.bam.qualimap/genome\_results.txt - results/{dataset}/mapping/sumatran\_rhino\_22Jul2017\_9M7eS\_haploidified\_headersFixed\_Sc9M7eS\_2\_HRSCAF\_41/stats/bams\_indels\_realigned/{sample,[A-Za-z0-9]+}.merged.rmdup.merged.realn.bam.qualimap | |
| Container image | |
| docker://quay.io/biocontainers/qualimap:2.2.2d--1 | |
| Code | |
| ``` 1 2 3 4 ``` \| ``` mem=$(((6 * {threads}) - 2)) unset DISPLAY qualimap bamqc -bam {input.bam} --java-mem-size=${{mem}}G -nt {threads} -outdir {params.outdir} -outformat html 2> {log} ``` | |

##### Rule realigned\_bam\_fastqc

×

Rule properties

|  |  |
| --- | --- |
| Jobs | 6 |
| Input files | |
| - results/{dataset}/mapping/sumatran\_rhino\_22Jul2017\_9M7eS\_haploidified\_headersFixed\_Sc9M7eS\_2\_HRSCAF\_41/{sample}.merged.rmdup.merged.realn.bam | |
| Output files | |
| - results/{dataset}/mapping/sumatran\_rhino\_22Jul2017\_9M7eS\_haploidified\_headersFixed\_Sc9M7eS\_2\_HRSCAF\_41/stats/bams\_indels\_realigned/fastqc/{sample,[A-Za-z0-9]+}.merged.rmdup.merged.realn\_fastqc.html - results/{dataset}/mapping/sumatran\_rhino\_22Jul2017\_9M7eS\_haploidified\_headersFixed\_Sc9M7eS\_2\_HRSCAF\_41/stats/bams\_indels\_realigned/fastqc/{sample,[A-Za-z0-9]+}.merged.rmdup.merged.realn\_fastqc.zip - results/{dataset}/mapping/sumatran\_rhino\_22Jul2017\_9M7eS\_haploidified\_headersFixed\_Sc9M7eS\_2\_HRSCAF\_41/stats/bams\_indels\_realigned/fastqc/{sample,[A-Za-z0-9]+}.merged.rmdup.merged.realn\_fastqc | |
| Container image | |
| docker://biocontainers/fastqc:v0.11.9\_cv7 | |
| Code | |
| ``` 1 2 ``` \| ``` fastqc -o {params.dir} -t {threads} --extract {input.bam} 2> {log} ``` | |

##### Rule modern\_merged\_index\_bam\_multiqc

×

Rule properties

|  |  |
| --- | --- |
| Jobs | 1 |
| Input files | |
| - results/modern/mapping/sumatran\_rhino\_22Jul2017\_9M7eS\_haploidified\_headersFixed\_Sc9M7eS\_2\_HRSCAF\_41/stats/bams\_merged\_index/JvS033\_01.merged.bam.stats.txt - results/modern/mapping/sumatran\_rhino\_22Jul2017\_9M7eS\_haploidified\_headersFixed\_Sc9M7eS\_2\_HRSCAF\_41/stats/bams\_merged\_index/JvS034\_02.merged.bam.stats.txt - results/modern/mapping/sumatran\_rhino\_22Jul2017\_9M7eS\_haploidified\_headersFixed\_Sc9M7eS\_2\_HRSCAF\_41/stats/bams\_merged\_index/JvS035\_14.merged.bam.stats.txt - results/modern/mapping/sumatran\_rhino\_22Jul2017\_9M7eS\_haploidified\_headersFixed\_Sc9M7eS\_2\_HRSCAF\_41/stats/bams\_merged\_index/JvS033\_01.merged.bam.qualimap/qualimapReport.html - results/modern/mapping/sumatran\_rhino\_22Jul2017\_9M7eS\_haploidified\_headersFixed\_Sc9M7eS\_2\_HRSCAF\_41/stats/bams\_merged\_index/JvS034\_02.merged.bam.qualimap/qualimapReport.html - results/modern/mapping/sumatran\_rhino\_22Jul2017\_9M7eS\_haploidified\_headersFixed\_Sc9M7eS\_2\_HRSCAF\_41/stats/bams\_merged\_index/JvS035\_14.merged.bam.qualimap/qualimapReport.html | |
| Output files | |
| - results/modern/mapping/sumatran\_rhino\_22Jul2017\_9M7eS\_haploidified\_headersFixed\_Sc9M7eS\_2\_HRSCAF\_41/stats/bams\_merged\_index/multiqc/multiqc\_report.html | |
| Container image | |
| docker://quay.io/biocontainers/multiqc:1.9--pyh9f0ad1d\_0 | |
| Code | |
| ``` 1 2 ``` \| ``` multiqc -f {params.indir} -o {params.outdir} 2> {log} ``` | |

##### Rule merge\_modern\_bams\_per\_index

×

Rule properties

|  |  |
| --- | --- |
| Jobs | 3 |
| Input files | |
| Output files | |
| - results/modern/mapping/sumatran\_rhino\_22Jul2017\_9M7eS\_haploidified\_headersFixed\_Sc9M7eS\_2\_HRSCAF\_41/{sample,[A-Za-z0-9]+}\_{index}.merged.bam | |
| Container image | |
| docker://biocontainers/samtools:v1.9-4-deb\_cv1 | |
| Code | |
| ``` 1 2 3 4 5 6 7 8 9 ``` \| ``` files=`echo {input} \| awk '{{print NF}}'` if [ $files -gt 1 ] # check if there are at least 2 files for merging. If there is only one file, copy the sorted bam file. then samtools merge {output.merged} {input} 2> {log} else cp {input} {output.merged} && touch {output.merged} 2> {log} echo "Only one file present for merging. Copying the input bam file." >> {log} fi ``` | |

##### Rule modern\_rmdup\_bam\_multiqc

×

Rule properties

|  |  |
| --- | --- |
| Jobs | 1 |
| Input files | |
| - results/modern/mapping/sumatran\_rhino\_22Jul2017\_9M7eS\_haploidified\_headersFixed\_Sc9M7eS\_2\_HRSCAF\_41/stats/bams\_rmdup/JvS033\_01.merged.rmdup.bam.stats.txt - results/modern/mapping/sumatran\_rhino\_22Jul2017\_9M7eS\_haploidified\_headersFixed\_Sc9M7eS\_2\_HRSCAF\_41/stats/bams\_rmdup/JvS034\_02.merged.rmdup.bam.stats.txt - results/modern/mapping/sumatran\_rhino\_22Jul2017\_9M7eS\_haploidified\_headersFixed\_Sc9M7eS\_2\_HRSCAF\_41/stats/bams\_rmdup/JvS035\_14.merged.rmdup.bam.stats.txt - results/modern/mapping/sumatran\_rhino\_22Jul2017\_9M7eS\_haploidified\_headersFixed\_Sc9M7eS\_2\_HRSCAF\_41/stats/bams\_rmdup/JvS033\_01.merged.rmdup.bam.qualimap/qualimapReport.html - results/modern/mapping/sumatran\_rhino\_22Jul2017\_9M7eS\_haploidified\_headersFixed\_Sc9M7eS\_2\_HRSCAF\_41/stats/bams\_rmdup/JvS034\_02.merged.rmdup.bam.qualimap/qualimapReport.html - results/modern/mapping/sumatran\_rhino\_22Jul2017\_9M7eS\_haploidified\_headersFixed\_Sc9M7eS\_2\_HRSCAF\_41/stats/bams\_rmdup/JvS035\_14.merged.rmdup.bam.qualimap/qualimapReport.html | |
| Output files | |
| - results/modern/mapping/sumatran\_rhino\_22Jul2017\_9M7eS\_haploidified\_headersFixed\_Sc9M7eS\_2\_HRSCAF\_41/stats/bams\_rmdup/multiqc/multiqc\_report.html | |
| Container image | |
| docker://quay.io/biocontainers/multiqc:1.9--pyh9f0ad1d\_0 | |
| Code | |
| ``` 1 2 ``` \| ``` multiqc -f {params.indir} -o {params.outdir} 2> {log} ``` | |

##### Rule rmdup\_modern\_bams

×

Rule properties

|  |  |
| --- | --- |
| Jobs | 3 |
| Input files | |
| - results/modern/mapping/sumatran\_rhino\_22Jul2017\_9M7eS\_haploidified\_headersFixed\_Sc9M7eS\_2\_HRSCAF\_41/{sample}\_{index}.merged.bam - results/modern/mapping/sumatran\_rhino\_22Jul2017\_9M7eS\_haploidified\_headersFixed\_Sc9M7eS\_2\_HRSCAF\_41/{sample}\_{index}.merged.bam.bai | |
| Output files | |
| - results/modern/mapping/sumatran\_rhino\_22Jul2017\_9M7eS\_haploidified\_headersFixed\_Sc9M7eS\_2\_HRSCAF\_41/{sample,[A-Za-z0-9]+}\_{index}.merged.rmdup.bam - results/modern/mapping/sumatran\_rhino\_22Jul2017\_9M7eS\_haploidified\_headersFixed\_Sc9M7eS\_2\_HRSCAF\_41/{sample,[A-Za-z0-9]+}\_{index}.merged.rmdup\_metrics.txt | |
| Container image | |
| docker://quay.io/biocontainers/picard:2.26.6--hdfd78af\_0 | |
| Code | |
| ``` 1 2 3 ``` \| ``` mem=$(((6 * {threads}) - 2)) picard MarkDuplicates -Xmx${{mem}}g INPUT={input.merged} OUTPUT={output.rmdup} METRICS_FILE={output.metrix} 2> {log} ``` | |

##### Rule modern\_merged\_sample\_bam\_multiqc

×

Rule properties

|  |  |
| --- | --- |
| Jobs | 1 |
| Input files | |
| - results/modern/mapping/sumatran\_rhino\_22Jul2017\_9M7eS\_haploidified\_headersFixed\_Sc9M7eS\_2\_HRSCAF\_41/stats/bams\_merged\_sample/JvS033.merged.rmdup.merged.bam.stats.txt - results/modern/mapping/sumatran\_rhino\_22Jul2017\_9M7eS\_haploidified\_headersFixed\_Sc9M7eS\_2\_HRSCAF\_41/stats/bams\_merged\_sample/JvS034.merged.rmdup.merged.bam.stats.txt - results/modern/mapping/sumatran\_rhino\_22Jul2017\_9M7eS\_haploidified\_headersFixed\_Sc9M7eS\_2\_HRSCAF\_41/stats/bams\_merged\_sample/JvS035.merged.rmdup.merged.bam.stats.txt - results/modern/mapping/sumatran\_rhino\_22Jul2017\_9M7eS\_haploidified\_headersFixed\_Sc9M7eS\_2\_HRSCAF\_41/stats/bams\_merged\_sample/JvS033.merged.rmdup.merged.bam.qualimap/qualimapReport.html - results/modern/mapping/sumatran\_rhino\_22Jul2017\_9M7eS\_haploidified\_headersFixed\_Sc9M7eS\_2\_HRSCAF\_41/stats/bams\_merged\_sample/JvS034.merged.rmdup.merged.bam.qualimap/qualimapReport.html - results/modern/mapping/sumatran\_rhino\_22Jul2017\_9M7eS\_haploidified\_headersFixed\_Sc9M7eS\_2\_HRSCAF\_41/stats/bams\_merged\_sample/JvS035.merged.rmdup.merged.bam.qualimap/qualimapReport.html | |
| Output files | |
| - results/modern/mapping/sumatran\_rhino\_22Jul2017\_9M7eS\_haploidified\_headersFixed\_Sc9M7eS\_2\_HRSCAF\_41/stats/bams\_merged\_sample/multiqc/multiqc\_report.html | |
| Container image | |
| docker://quay.io/biocontainers/multiqc:1.9--pyh9f0ad1d\_0 | |
| Code | |
| ``` 1 2 ``` \| ``` multiqc -f {params.indir} -o {params.outdir} 2> {log} ``` | |

##### Rule merge\_modern\_bams\_per\_sample

×

Rule properties

|  |  |
| --- | --- |
| Jobs | 3 |
| Input files | |
| Output files | |
| - results/modern/mapping/sumatran\_rhino\_22Jul2017\_9M7eS\_haploidified\_headersFixed\_Sc9M7eS\_2\_HRSCAF\_41/{sample,[A-Za-z0-9]+}.merged.rmdup.merged.bam | |
| Container image | |
| docker://biocontainers/samtools:v1.9-4-deb\_cv1 | |
| Code | |
| ``` 1 2 3 4 5 6 7 8 9 ``` \| ``` files=`echo {input} \| awk '{{print NF}}'` if [ $files -gt 1 ] # check if there are at least 2 files for merging. If there is only one file, copy the sorted bam file. then samtools merge {output.merged} {input} 2> {log} else cp {input} {output.merged} && touch {output.merged} 2> {log} echo "Only one file present for merging. Copying the input bam file." >> {log} fi ``` | |

##### Rule modern\_realigned\_bam\_multiqc

×

Rule properties

|  |  |
| --- | --- |
| Jobs | 1 |
| Input files | |
| - results/modern/mapping/sumatran\_rhino\_22Jul2017\_9M7eS\_haploidified\_headersFixed\_Sc9M7eS\_2\_HRSCAF\_41/stats/bams\_indels\_realigned/JvS033.merged.rmdup.merged.realn.repma.Q30.bam.dp.hist.pdf - results/modern/mapping/sumatran\_rhino\_22Jul2017\_9M7eS\_haploidified\_headersFixed\_Sc9M7eS\_2\_HRSCAF\_41/stats/bams\_indels\_realigned/JvS034.merged.rmdup.merged.realn.repma.Q30.bam.dp.hist.pdf - results/modern/mapping/sumatran\_rhino\_22Jul2017\_9M7eS\_haploidified\_headersFixed\_Sc9M7eS\_2\_HRSCAF\_41/stats/bams\_indels\_realigned/JvS035.merged.rmdup.merged.realn.repma.Q30.bam.dp.hist.pdf - results/modern/mapping/sumatran\_rhino\_22Jul2017\_9M7eS\_haploidified\_headersFixed\_Sc9M7eS\_2\_HRSCAF\_41/stats/bams\_indels\_realigned/JvS033.merged.rmdup.merged.realn.bam.stats.txt - results/modern/mapping/sumatran\_rhino\_22Jul2017\_9M7eS\_haploidified\_headersFixed\_Sc9M7eS\_2\_HRSCAF\_41/stats/bams\_indels\_realigned/JvS034.merged.rmdup.merged.realn.bam.stats.txt - results/modern/mapping/sumatran\_rhino\_22Jul2017\_9M7eS\_haploidified\_headersFixed\_Sc9M7eS\_2\_HRSCAF\_41/stats/bams\_indels\_realigned/JvS035.merged.rmdup.merged.realn.bam.stats.txt - results/modern/mapping/sumatran\_rhino\_22Jul2017\_9M7eS\_haploidified\_headersFixed\_Sc9M7eS\_2\_HRSCAF\_41/stats/bams\_indels\_realigned/JvS033.merged.rmdup.merged.realn.bam.qualimap/qualimapReport.html - results/modern/mapping/sumatran\_rhino\_22Jul2017\_9M7eS\_haploidified\_headersFixed\_Sc9M7eS\_2\_HRSCAF\_41/stats/bams\_indels\_realigned/JvS034.merged.rmdup.merged.realn.bam.qualimap/qualimapReport.html - results/modern/mapping/sumatran\_rhino\_22Jul2017\_9M7eS\_haploidified\_headersFixed\_Sc9M7eS\_2\_HRSCAF\_41/stats/bams\_indels\_realigned/JvS035.merged.rmdup.merged.realn.bam.qualimap/qualimapReport.html - results/modern/mapping/sumatran\_rhino\_22Jul2017\_9M7eS\_haploidified\_headersFixed\_Sc9M7eS\_2\_HRSCAF\_41/stats/bams\_indels\_realigned/fastqc/JvS033.merged.rmdup.merged.realn\_fastqc.html - results/modern/mapping/sumatran\_rhino\_22Jul2017\_9M7eS\_haploidified\_headersFixed\_Sc9M7eS\_2\_HRSCAF\_41/stats/bams\_indels\_realigned/fastqc/JvS034.merged.rmdup.merged.realn\_fastqc.html - results/modern/mapping/sumatran\_rhino\_22Jul2017\_9M7eS\_haploidified\_headersFixed\_Sc9M7eS\_2\_HRSCAF\_41/stats/bams\_indels\_realigned/fastqc/JvS035.merged.rmdup.merged.realn\_fastqc.html - results/modern/mapping/sumatran\_rhino\_22Jul2017\_9M7eS\_haploidified\_headersFixed\_Sc9M7eS\_2\_HRSCAF\_41/stats/bams\_indels\_realigned/fastqc/JvS033.merged.rmdup.merged.realn\_fastqc.zip - results/modern/mapping/sumatran\_rhino\_22Jul2017\_9M7eS\_haploidified\_headersFixed\_Sc9M7eS\_2\_HRSCAF\_41/stats/bams\_indels\_realigned/fastqc/JvS034.merged.rmdup.merged.realn\_fastqc.zip - results/modern/mapping/sumatran\_rhino\_22Jul2017\_9M7eS\_haploidified\_headersFixed\_Sc9M7eS\_2\_HRSCAF\_41/stats/bams\_indels\_realigned/fastqc/JvS035.merged.rmdup.merged.realn\_fastqc.zip - results/modern/mapping/sumatran\_rhino\_22Jul2017\_9M7eS\_haploidified\_headersFixed\_Sc9M7eS\_2\_HRSCAF\_41/stats/bams\_indels\_realigned/JvS033.merged.rmdup.merged.realn.repma.Q30.bam.dp.hist.pdf - results/modern/mapping/sumatran\_rhino\_22Jul2017\_9M7eS\_haploidified\_headersFixed\_Sc9M7eS\_2\_HRSCAF\_41/stats/bams\_indels\_realigned/JvS034.merged.rmdup.merged.realn.repma.Q30.bam.dp.hist.pdf - results/modern/mapping/sumatran\_rhino\_22Jul2017\_9M7eS\_haploidified\_headersFixed\_Sc9M7eS\_2\_HRSCAF\_41/stats/bams\_indels\_realigned/JvS035.merged.rmdup.merged.realn.repma.Q30.bam.dp.hist.pdf | |
| Output files | |
| - results/modern/mapping/sumatran\_rhino\_22Jul2017\_9M7eS\_haploidified\_headersFixed\_Sc9M7eS\_2\_HRSCAF\_41/stats/bams\_indels\_realigned/multiqc/multiqc\_report.html | |
| Container image | |
| docker://quay.io/biocontainers/multiqc:1.9--pyh9f0ad1d\_0 | |
| Code | |
| ``` 1 2 ``` \| ``` multiqc -f {params.indir} -o {params.outdir} 2> {log} ``` | |

##### Rule modern\_subsampled\_bam\_multiqc

×

Rule properties

|  |  |
| --- | --- |
| Jobs | 1 |
| Input files | |
| - results/modern/mapping/sumatran\_rhino\_22Jul2017\_9M7eS\_haploidified\_headersFixed\_Sc9M7eS\_2\_HRSCAF\_41/stats/bams\_subsampled/JvS033.merged.rmdup.merged.realn.mapped\_q30.subs\_dp6.bam.stats.txt - results/modern/mapping/sumatran\_rhino\_22Jul2017\_9M7eS\_haploidified\_headersFixed\_Sc9M7eS\_2\_HRSCAF\_41/stats/bams\_subsampled/JvS035.merged.rmdup.merged.realn.mapped\_q30.subs\_dp6.bam.stats.txt - results/modern/mapping/sumatran\_rhino\_22Jul2017\_9M7eS\_haploidified\_headersFixed\_Sc9M7eS\_2\_HRSCAF\_41/stats/bams\_subsampled/JvS034.merged.rmdup.merged.realn.mapped\_q30.subs\_dp6.bam.stats.txt - results/modern/mapping/sumatran\_rhino\_22Jul2017\_9M7eS\_haploidified\_headersFixed\_Sc9M7eS\_2\_HRSCAF\_41/stats/bams\_subsampled/JvS033.merged.rmdup.merged.realn.mapped\_q30.subs\_dp6.bam.qualimap/qualimapReport.html - results/modern/mapping/sumatran\_rhino\_22Jul2017\_9M7eS\_haploidified\_headersFixed\_Sc9M7eS\_2\_HRSCAF\_41/stats/bams\_subsampled/JvS035.merged.rmdup.merged.realn.mapped\_q30.subs\_dp6.bam.qualimap/qualimapReport.html - results/modern/mapping/sumatran\_rhino\_22Jul2017\_9M7eS\_haploidified\_headersFixed\_Sc9M7eS\_2\_HRSCAF\_41/stats/bams\_subsampled/JvS034.merged.rmdup.merged.realn.mapped\_q30.subs\_dp6.bam.qualimap/qualimapReport.html - results/modern/mapping/sumatran\_rhino\_22Jul2017\_9M7eS\_haploidified\_headersFixed\_Sc9M7eS\_2\_HRSCAF\_41/stats/bams\_subsampled/JvS033.merged.rmdup.merged.realn.mapped\_q30.subs\_dp6.repma.Q30.bam.dpstats.txt - results/modern/mapping/sumatran\_rhino\_22Jul2017\_9M7eS\_haploidified\_headersFixed\_Sc9M7eS\_2\_HRSCAF\_41/stats/bams\_subsampled/JvS035.merged.rmdup.merged.realn.mapped\_q30.subs\_dp6.repma.Q30.bam.dpstats.txt - results/modern/mapping/sumatran\_rhino\_22Jul2017\_9M7eS\_haploidified\_headersFixed\_Sc9M7eS\_2\_HRSCAF\_41/stats/bams\_subsampled/JvS034.merged.rmdup.merged.realn.mapped\_q30.subs\_dp6.repma.Q30.bam.dpstats.txt | |
| Output files | |
| - results/modern/mapping/sumatran\_rhino\_22Jul2017\_9M7eS\_haploidified\_headersFixed\_Sc9M7eS\_2\_HRSCAF\_41/stats/bams\_subsampled/multiqc/multiqc\_report.html | |
| Container image | |
| docker://quay.io/biocontainers/multiqc:1.9--pyh9f0ad1d\_0 | |
| Code | |
| ``` 1 2 ``` \| ``` multiqc -f {params.indir} -o {params.outdir} 2> {log} ``` | |

##### Rule subsampled\_bam\_stats

×

Rule properties

|  |  |
| --- | --- |
| Jobs | 3 |
| Input files | |
| - results/{dataset}/mapping/sumatran\_rhino\_22Jul2017\_9M7eS\_haploidified\_headersFixed\_Sc9M7eS\_2\_HRSCAF\_41/{sample}.merged.rmdup.merged.{processed}.mapped\_q30.subs\_dp{DP}.bam - results/{dataset}/mapping/sumatran\_rhino\_22Jul2017\_9M7eS\_haploidified\_headersFixed\_Sc9M7eS\_2\_HRSCAF\_41/{sample}.merged.rmdup.merged.{processed}.mapped\_q30.subs\_dp{DP}.bam.bai | |
| Output files | |
| - results/{dataset}/mapping/sumatran\_rhino\_22Jul2017\_9M7eS\_haploidified\_headersFixed\_Sc9M7eS\_2\_HRSCAF\_41/stats/bams\_subsampled/{sample,[A-Za-z0-9]+}.merged.rmdup.merged.{processed}.mapped\_q30.subs\_dp{DP,[+-]?(\d+(\.\d\*)?\|\.\d+)([eE][+-]?\d+)?}.bam.stats.txt | |
| Container image | |
| docker://biocontainers/samtools:v1.9-4-deb\_cv1 | |
| Code | |
| ``` 1 2 ``` \| ``` samtools flagstat {input.bam} > {output.stats} 2> {log} ``` | |

##### Rule subsample\_bams

×

Rule properties

|  |  |
| --- | --- |
| Jobs | 3 |
| Input files | |
| - results/{dataset}/mapping/sumatran\_rhino\_22Jul2017\_9M7eS\_haploidified\_headersFixed\_Sc9M7eS\_2\_HRSCAF\_41/{sample}.merged.rmdup.merged.{processed}.mapped\_q30.bam - results/{dataset}/mapping/sumatran\_rhino\_22Jul2017\_9M7eS\_haploidified\_headersFixed\_Sc9M7eS\_2\_HRSCAF\_41/stats/bams\_indels\_realigned/{sample}.merged.rmdup.merged.realn.repma.Q30.bam.dpstats.txt | |
| Output files | |
| - results/{dataset}/mapping/sumatran\_rhino\_22Jul2017\_9M7eS\_haploidified\_headersFixed\_Sc9M7eS\_2\_HRSCAF\_41/{sample,[A-Za-z0-9]+}.merged.rmdup.merged.{processed}.mapped\_q30.subs\_dp{DP,[+-]?(\d+(\.\d\*)?\|\.\d+)([eE][+-]?\d+)?}.bam | |
| Container image | |
| docker://biocontainers/samtools:v1.9-4-deb\_cv1 | |
| Code | |
| ``` 1 2 3 4 5 6 7 8 9 10 11 ``` \| ``` depth=`head -n 1 {input.dp} \| cut -d' ' -f 1` frac=`awk -v s={params.DP} -v d=$depth "BEGIN {{print s/d}}"` if [ `awk 'BEGIN {{print ('$frac' <= 1.0)}}'` = 1 ] # awk will return 1 if the statement is true, and 0 if it is false then samtools view -h -b -s $frac -@ {threads} -o {output.subsam} {input.bam} 2> {log} else echo "!!! Warning [genome erosion workflow]: The sample {input.bam} has a lower average depth than the target depth for subsampling. Remove the sample from the subsampling list in the config file or choose a lower target depth. !!!" >> {log} fi ``` | |

##### Rule filter\_bam\_mapped\_mq

×

Rule properties

|  |  |
| --- | --- |
| Jobs | 3 |
| Input files | |
| - results/{dataset}/mapping/sumatran\_rhino\_22Jul2017\_9M7eS\_haploidified\_headersFixed\_Sc9M7eS\_2\_HRSCAF\_41/{sample}.merged.rmdup.merged.{processed}.bam | |
| Output files | |
| - results/{dataset}/mapping/sumatran\_rhino\_22Jul2017\_9M7eS\_haploidified\_headersFixed\_Sc9M7eS\_2\_HRSCAF\_41/{sample,[A-Za-z0-9]+}.merged.rmdup.merged.{processed}.mapped\_q30.bam | |
| Container image | |
| docker://biocontainers/samtools:v1.9-4-deb\_cv1 | |
| Code | |
| ``` 1 2 ``` \| ``` samtools view -h -b -F 4 -q 30 -@ {threads} -o {output.filtered} {input.bam} 2> {log} ``` | |

##### Rule index\_subsampled\_bams

×

Rule properties

|  |  |
| --- | --- |
| Jobs | 3 |
| Input files | |
| - results/{dataset}/mapping/sumatran\_rhino\_22Jul2017\_9M7eS\_haploidified\_headersFixed\_Sc9M7eS\_2\_HRSCAF\_41/{sample}.merged.rmdup.merged.{processed}.mapped\_q30.subs\_dp{DP}.bam | |
| Output files | |
| - results/{dataset}/mapping/sumatran\_rhino\_22Jul2017\_9M7eS\_haploidified\_headersFixed\_Sc9M7eS\_2\_HRSCAF\_41/{sample,[A-Za-z0-9]+}.merged.rmdup.merged.{processed}.mapped\_q30.subs\_dp{DP,[+-]?(\d+(\.\d\*)?\|\.\d+)([eE][+-]?\d+)?}.bam.bai | |
| Container image | |
| docker://biocontainers/samtools:v1.9-4-deb\_cv1 | |
| Code | |
| ``` 1 2 ``` \| ``` samtools index {input.bam} {output.index} 2> {log} ``` | |

##### Rule subsampled\_bam\_qualimap

×

Rule properties

|  |  |
| --- | --- |
| Jobs | 3 |
| Input files | |
| - results/{dataset}/mapping/sumatran\_rhino\_22Jul2017\_9M7eS\_haploidified\_headersFixed\_Sc9M7eS\_2\_HRSCAF\_41/{sample}.merged.rmdup.merged.{processed}.mapped\_q30.subs\_dp{DP}.bam - results/{dataset}/mapping/sumatran\_rhino\_22Jul2017\_9M7eS\_haploidified\_headersFixed\_Sc9M7eS\_2\_HRSCAF\_41/{sample}.merged.rmdup.merged.{processed}.mapped\_q30.subs\_dp{DP}.bam.bai | |
| Output files | |
| - results/{dataset}/mapping/sumatran\_rhino\_22Jul2017\_9M7eS\_haploidified\_headersFixed\_Sc9M7eS\_2\_HRSCAF\_41/stats/bams\_subsampled/{sample,[A-Za-z0-9]+}.merged.rmdup.merged.{processed}.mapped\_q30.subs\_dp{DP,[+-]?(\d+(\.\d\*)?\|\.\d+)([eE][+-]?\d+)?}.bam.qualimap/qualimapReport.html - results/{dataset}/mapping/sumatran\_rhino\_22Jul2017\_9M7eS\_haploidified\_headersFixed\_Sc9M7eS\_2\_HRSCAF\_41/stats/bams\_subsampled/{sample,[A-Za-z0-9]+}.merged.rmdup.merged.{processed}.mapped\_q30.subs\_dp{DP,[+-]?(\d+(\.\d\*)?\|\.\d+)([eE][+-]?\d+)?}.bam.qualimap/genome\_results.txt - results/{dataset}/mapping/sumatran\_rhino\_22Jul2017\_9M7eS\_haploidified\_headersFixed\_Sc9M7eS\_2\_HRSCAF\_41/stats/bams\_subsampled/{sample,[A-Za-z0-9]+}.merged.rmdup.merged.{processed}.mapped\_q30.subs\_dp{DP,[+-]?(\d+(\.\d\*)?\|\.\d+)([eE][+-]?\d+)?}.bam.qualimap | |
| Container image | |
| docker://quay.io/biocontainers/qualimap:2.2.2d--1 | |
| Code | |
| ``` 1 2 3 4 ``` \| ``` mem=$(((6 * {threads}) - 2)) unset DISPLAY qualimap bamqc -bam {input.bam} --java-mem-size=${{mem}}G -nt {threads} -outdir {params.outdir} -outformat html 2> {log} ``` | |

##### Rule subsampled\_bam\_depth

×

Rule properties

|  |  |
| --- | --- |
| Jobs | 3 |
| Input files | |
| - results/{dataset}/mapping/sumatran\_rhino\_22Jul2017\_9M7eS\_haploidified\_headersFixed\_Sc9M7eS\_2\_HRSCAF\_41/{sample}.merged.rmdup.merged.{processed}.mapped\_q30.subs\_dp{DP}.bam - /proj/sllstore2017093/b2016342/b2016342\_nobackup/lts/genome\_erosion\_pipeline/verena\_testing/testdata/reference/sumatran\_rhino\_22Jul2017\_9M7eS\_haploidified\_headersFixed\_Sc9M7eS\_2\_HRSCAF\_41.repma.bed | |
| Output files | |
| - results/{dataset}/mapping/sumatran\_rhino\_22Jul2017\_9M7eS\_haploidified\_headersFixed\_Sc9M7eS\_2\_HRSCAF\_41/stats/bams\_subsampled/{sample,[A-Za-z0-9]+}.merged.rmdup.merged.{processed}.mapped\_q30.subs\_dp{DP,[+-]?(\d+(\.\d\*)?\|\.\d+)([eE][+-]?\d+)?}.repma.Q30.bam.dp - results/{dataset}/mapping/sumatran\_rhino\_22Jul2017\_9M7eS\_haploidified\_headersFixed\_Sc9M7eS\_2\_HRSCAF\_41/stats/bams\_subsampled/{sample,[A-Za-z0-9]+}.merged.rmdup.merged.{processed}.mapped\_q30.subs\_dp{DP,[+-]?(\d+(\.\d\*)?\|\.\d+)([eE][+-]?\d+)?}.repma.Q30.bam.dpstats.txt | |
| Container image | |
| docker://biocontainers/samtools:v1.9-4-deb\_cv1 | |
| Code | |
| ``` 1 2 3 4 5 6 7 8 9 10 ``` \| ``` if [ {params.cov} = "True" ] # include sites with missing data / zero coverage then samtools depth -a -Q 30 -q 30 -b {input.no_rep_bed} {input.bam} > {output.tmp} 2> {log} && awk '{{sum+=$3}} END {{ print sum/NR }}' {output.tmp} \| awk -v min={params.minDP} -v max={params.maxDP} '{{ printf "%.0f %.0f %.0f", $1, $1*min, $1*max }}' > {output.dp} 2>> {log} elif [ {params.cov} = "False" ] # exclude sites with missing data / zero coverage then samtools depth -Q 30 -q 30 -b {input.no_rep_bed} {input.bam} > {output.tmp} 2> {log} && awk '{{sum+=$3}} END {{ print sum/NR }}' {output.tmp} \| awk -v min={params.minDP} -v max={params.maxDP} '{{ printf "%.0f %.0f %.0f", $1, $1*min, $1*max }}' > {output.dp} 2>> {log} fi ``` | |

##### Rule make\_CpG\_reference\_bed

×

Rule properties

|  |  |
| --- | --- |
| Jobs | 1 |
| Input files | |
| - /proj/sllstore2017093/b2016342/b2016342\_nobackup/lts/genome\_erosion\_pipeline/verena\_testing/testdata/reference/sumatran\_rhino\_22Jul2017\_9M7eS\_haploidified\_headersFixed\_Sc9M7eS\_2\_HRSCAF\_41.upper.fasta | |
| Output files | |
| - results/sumatran\_rhino\_22Jul2017\_9M7eS\_haploidified\_headersFixed\_Sc9M7eS\_2\_HRSCAF\_41.CpG\_ref.bed | |
| Code | |
| ``` 1 2 3 ``` \| ``` python workflow/scripts/find_CpG_ref_sites.py {input.ref} {params.refdirbed} 2> {log} && cp {params.refdirbed}.bed {output.bed} 2>> {log} ``` | |

##### Rule make\_noCpG\_bed

×

Rule properties

|  |  |
| --- | --- |
| Jobs | 1 |
| Input files | |
| - /proj/sllstore2017093/b2016342/b2016342\_nobackup/lts/genome\_erosion\_pipeline/verena\_testing/testdata/reference/sumatran\_rhino\_22Jul2017\_9M7eS\_haploidified\_headersFixed\_Sc9M7eS\_2\_HRSCAF\_41.bed - results/sumatran\_rhino\_22Jul2017\_9M7eS\_haploidified\_headersFixed\_Sc9M7eS\_2\_HRSCAF\_41.{CpG\_method}.bed | |
| Output files | |
| - results/sumatran\_rhino\_22Jul2017\_9M7eS\_haploidified\_headersFixed\_Sc9M7eS\_2\_HRSCAF\_41.no{CpG\_method,CpG\_[vcfre]{3,6}}.bed | |
| Container image | |
| docker://quay.io/biocontainers/bedtools:2.29.2--hc088bd4\_0 | |
| Code | |
| ``` 1 2 ``` \| ``` bedtools subtract -a {input.ref_bed} -b {input.CpG_bed} > {output.no_CpG_bed} 2> {log} ``` | |

##### Rule make\_noCpG\_repma\_bed

×

Rule properties

|  |  |
| --- | --- |
| Jobs | 1 |
| Input files | |
| - /proj/sllstore2017093/b2016342/b2016342\_nobackup/lts/genome\_erosion\_pipeline/verena\_testing/testdata/reference/sumatran\_rhino\_22Jul2017\_9M7eS\_haploidified\_headersFixed\_Sc9M7eS\_2\_HRSCAF\_41.bed - results/sumatran\_rhino\_22Jul2017\_9M7eS\_haploidified\_headersFixed\_Sc9M7eS\_2\_HRSCAF\_41.merged.{CpG\_method}.repeats.bed | |
| Output files | |
| - results/sumatran\_rhino\_22Jul2017\_9M7eS\_haploidified\_headersFixed\_Sc9M7eS\_2\_HRSCAF\_41.no{CpG\_method,CpG\_[vcfre]{3,6}}.repma.bed | |
| Container image | |
| docker://quay.io/biocontainers/bedtools:2.29.2--hc088bd4\_0 | |
| Code | |
| ``` 1 2 ``` \| ``` bedtools subtract -a {input.ref_bed} -b {input.merged_bed} > {output.no_CpG_repma_bed} 2> {log} ``` | |

##### Rule merge\_CpG\_repeats\_beds

×

Rule properties

|  |  |
| --- | --- |
| Jobs | 1 |
| Input files | |
| - results/sumatran\_rhino\_22Jul2017\_9M7eS\_haploidified\_headersFixed\_Sc9M7eS\_2\_HRSCAF\_41.{CpG\_method}.bed - /proj/sllstore2017093/b2016342/b2016342\_nobackup/lts/genome\_erosion\_pipeline/verena\_testing/testdata/reference/sumatran\_rhino\_22Jul2017\_9M7eS\_haploidified\_headersFixed\_Sc9M7eS\_2\_HRSCAF\_41.repeats.sorted.bed | |
| Output files | |
| - results/sumatran\_rhino\_22Jul2017\_9M7eS\_haploidified\_headersFixed\_Sc9M7eS\_2\_HRSCAF\_41.concatenated.{CpG\_method,CpG\_[vcfre]{3,6}}.repeats.bed - results/sumatran\_rhino\_22Jul2017\_9M7eS\_haploidified\_headersFixed\_Sc9M7eS\_2\_HRSCAF\_41.merged.{CpG\_method,CpG\_[vcfre]{3,6}}.repeats.bed | |
| Container image | |
| docker://quay.io/biocontainers/bedtools:2.29.2--hc088bd4\_0 | |
| Code | |
| ``` 1 2 3 ``` \| ``` cat {input[0]} {input[1]} \| sort -k1,1 -k2,2n > {output.tmp} 2> {log} && bedtools merge -i {output.tmp} > {output.merged} 2>> {log} ``` | |

##### Rule sort\_CpG\_repeats\_beds

×

Rule properties

|  |  |
| --- | --- |
| Jobs | 1 |
| Input files | |
| - results/sumatran\_rhino\_22Jul2017\_9M7eS\_haploidified\_headersFixed\_Sc9M7eS\_2\_HRSCAF\_41.merged.{CpG\_method}.repeats.bed - /proj/sllstore2017093/b2016342/b2016342\_nobackup/lts/genome\_erosion\_pipeline/verena\_testing/testdata/reference/sumatran\_rhino\_22Jul2017\_9M7eS\_haploidified\_headersFixed\_Sc9M7eS\_2\_HRSCAF\_41.genome | |
| Output files | |
| - results/sumatran\_rhino\_22Jul2017\_9M7eS\_haploidified\_headersFixed\_Sc9M7eS\_2\_HRSCAF\_41.{CpG\_method,CpG\_[vcfre]{3,6}}.repeats.bed | |
| Container image | |
| docker://quay.io/biocontainers/bedtools:2.29.2--hc088bd4\_0 | |
| Code | |
| ``` 1 2 ``` \| ``` bedtools sort -g {input.genomefile} -i {input.merged_bed} > {output.sorted_bed} 2> {log} ``` | |

##### Rule mlRho\_theta\_plot

×

Rule properties

|  |  |
| --- | --- |
| Jobs | 1 |
| Input files | |
| - results/{dataset}/mlRho/sumatran\_rhino\_22Jul2017\_9M7eS\_haploidified\_headersFixed\_Sc9M7eS\_2\_HRSCAF\_41.{dataset}.mlRho\_table.txt | |
| Output files | |
| - results/{dataset}/mlRho/sumatran\_rhino\_22Jul2017\_9M7eS\_haploidified\_headersFixed\_Sc9M7eS\_2\_HRSCAF\_41.{dataset}.mlRho\_theta\_plot.pdf | |

##### Rule mlRho\_table

×

Rule properties

|  |  |
| --- | --- |
| Jobs | 1 |
| Input files | |
| Output files | |
| - results/{dataset}/mlRho/sumatran\_rhino\_22Jul2017\_9M7eS\_haploidified\_headersFixed\_Sc9M7eS\_2\_HRSCAF\_41.{dataset}.mlRho\_table.txt | |

##### Rule mlRho\_all

×

Rule properties

|  |  |
| --- | --- |
| Jobs | 6 |
| Input files | |
| - results/{dataset}/mlRho/sumatran\_rhino\_22Jul2017\_9M7eS\_haploidified\_headersFixed\_Sc9M7eS\_2\_HRSCAF\_41/{sample}.merged.rmdup.merged.{processed}.all.pro | |
| Output files | |
| - results/{dataset}/mlRho/sumatran\_rhino\_22Jul2017\_9M7eS\_haploidified\_headersFixed\_Sc9M7eS\_2\_HRSCAF\_41/{sample,[A-Za-z0-9]+}.merged.rmdup.merged.{processed}.all.mlRho.txt - results/{dataset}/mlRho/sumatran\_rhino\_22Jul2017\_9M7eS\_haploidified\_headersFixed\_Sc9M7eS\_2\_HRSCAF\_41/{sample,[A-Za-z0-9]+}.merged.rmdup.merged.{processed}.all\_profileDb.con - results/{dataset}/mlRho/sumatran\_rhino\_22Jul2017\_9M7eS\_haploidified\_headersFixed\_Sc9M7eS\_2\_HRSCAF\_41/{sample,[A-Za-z0-9]+}.merged.rmdup.merged.{processed}.all\_profileDb.lik - results/{dataset}/mlRho/sumatran\_rhino\_22Jul2017\_9M7eS\_haploidified\_headersFixed\_Sc9M7eS\_2\_HRSCAF\_41/{sample,[A-Za-z0-9]+}.merged.rmdup.merged.{processed}.all\_profileDb.pos - results/{dataset}/mlRho/sumatran\_rhino\_22Jul2017\_9M7eS\_haploidified\_headersFixed\_Sc9M7eS\_2\_HRSCAF\_41/{sample,[A-Za-z0-9]+}.merged.rmdup.merged.{processed}.all\_profileDb.sum | |
| Container image | |
| docker://nbisweden/generode-mlrho | |
| Code | |
| ``` 1 2 3 4 5 6 7 8 9 10 11 12 13 14 ``` \| ``` minDP=`head -n 1 {input.dp} \| cut -d' ' -f 2` # check minimum depth threshold if awk "BEGIN{{exit ! ($minDP < 3)}}" then minDP=3 fi # Further format the pro file formatPro -c $minDP -n {params.db} {input.pro} 2> {log} && # run mlRho mlRho -M 0 -I -n {params.db} > {output.mlRho} 2>> {log} ``` | |

##### Rule bam2pro\_all

×

Rule properties

|  |  |
| --- | --- |
| Jobs | 6 |
| Input files | |
| Output files | |
| - results/{dataset}/mlRho/sumatran\_rhino\_22Jul2017\_9M7eS\_haploidified\_headersFixed\_Sc9M7eS\_2\_HRSCAF\_41/{sample,[A-Za-z0-9]+}.merged.rmdup.merged.{processed}.all.pro | |
| Container image | |
| docker://nbisweden/generode-mlrho | |
| Code | |
| ``` 1 2 3 4 5 6 7 8 9 10 11 ``` \| ``` minDP=`head -n 1 {input.dp} \| cut -d' ' -f 2` maxDP=`head -n 1 {input.dp} \| cut -d' ' -f 3` # check minimum depth threshold if awk "BEGIN{{exit ! ($minDP < 3)}}" then minDP=3 fi samtools mpileup -q 30 -Q 30 -B -l {input.bed} {input.bam[0]} \| awk -v minDP="$minDP" -v maxDP="$maxDP" '$4 >=minDP && $4 <=maxDP' \| sam2pro -c 5 > {output.pro} 2> {log} ``` | |

##### Rule historical\_sorted\_vcf\_multiqc

×

Rule properties

|  |  |
| --- | --- |
| Jobs | 1 |
| Input files | |
| - results/historical/vcf/sumatran\_rhino\_22Jul2017\_9M7eS\_haploidified\_headersFixed\_Sc9M7eS\_2\_HRSCAF\_41/stats/vcf\_sorted/JvS008.merged.rmdup.merged.realn.Q30.sorted.vcf.stats.txt - results/historical/vcf/sumatran\_rhino\_22Jul2017\_9M7eS\_haploidified\_headersFixed\_Sc9M7eS\_2\_HRSCAF\_41/stats/vcf\_sorted/JvS022.merged.rmdup.merged.realn.Q30.sorted.vcf.stats.txt - results/historical/vcf/sumatran\_rhino\_22Jul2017\_9M7eS\_haploidified\_headersFixed\_Sc9M7eS\_2\_HRSCAF\_41/stats/vcf\_sorted/JvS009.merged.rmdup.merged.realn.Q30.sorted.vcf.stats.txt | |
| Output files | |
| - results/historical/vcf/sumatran\_rhino\_22Jul2017\_9M7eS\_haploidified\_headersFixed\_Sc9M7eS\_2\_HRSCAF\_41/stats/vcf\_sorted/multiqc/multiqc\_report.html | |
| Container image | |
| docker://quay.io/biocontainers/multiqc:1.9--pyh9f0ad1d\_0 | |
| Code | |
| ``` 1 2 ``` \| ``` multiqc -f {params.indir} -o {params.outdir} 2> {log} ``` | |

##### Rule sorted\_vcf\_stats

×

Rule properties

|  |  |
| --- | --- |
| Jobs | 6 |
| Input files | |
| - results/{dataset}/vcf/sumatran\_rhino\_22Jul2017\_9M7eS\_haploidified\_headersFixed\_Sc9M7eS\_2\_HRSCAF\_41/{sample}.merged.rmdup.merged.{processed}.Q30.sorted.bcf - results/{dataset}/vcf/sumatran\_rhino\_22Jul2017\_9M7eS\_haploidified\_headersFixed\_Sc9M7eS\_2\_HRSCAF\_41/{sample}.merged.rmdup.merged.{processed}.Q30.sorted.bcf.csi | |
| Output files | |
| - results/{dataset}/vcf/sumatran\_rhino\_22Jul2017\_9M7eS\_haploidified\_headersFixed\_Sc9M7eS\_2\_HRSCAF\_41/stats/vcf\_sorted/{sample,[A-Za-z0-9]+}.merged.rmdup.merged.{processed}.Q30.sorted.vcf.stats.txt | |
| Container image | |
| docker://quay.io/biocontainers/bcftools:1.9--h68d8f2e\_9 | |
| Code | |
| ``` 1 2 ``` \| ``` bcftools stats {input.sort} > {output.stats} 2> {log} ``` | |

##### Rule sort\_vcfs

×

Rule properties

|  |  |
| --- | --- |
| Jobs | 6 |
| Input files | |
| - results/{dataset}/vcf/sumatran\_rhino\_22Jul2017\_9M7eS\_haploidified\_headersFixed\_Sc9M7eS\_2\_HRSCAF\_41/{sample}.merged.rmdup.merged.{processed}.Q30.bcf | |
| Output files | |
| - results/{dataset}/vcf/sumatran\_rhino\_22Jul2017\_9M7eS\_haploidified\_headersFixed\_Sc9M7eS\_2\_HRSCAF\_41/{sample,[A-Za-z0-9]+}.merged.rmdup.merged.{processed}.Q30.sorted.bcf | |
| Container image | |
| docker://quay.io/biocontainers/bcftools:1.9--h68d8f2e\_9 | |
| Code | |
| ``` 1 2 ``` \| ``` bcftools sort -O b -T {params.tmpdir} -o {output.sort} {input.bcf} 2> {log} ``` | |

##### Rule variant\_calling

×

Rule properties

|  |  |
| --- | --- |
| Jobs | 6 |
| Input files | |
| - /proj/sllstore2017093/b2016342/b2016342\_nobackup/lts/genome\_erosion\_pipeline/verena\_testing/testdata/reference/sumatran\_rhino\_22Jul2017\_9M7eS\_haploidified\_headersFixed\_Sc9M7eS\_2\_HRSCAF\_41.fasta - results/{dataset}/mapping/sumatran\_rhino\_22Jul2017\_9M7eS\_haploidified\_headersFixed\_Sc9M7eS\_2\_HRSCAF\_41/{sample}.merged.rmdup.merged.{processed}.bam | |
| Output files | |
| - results/{dataset}/vcf/sumatran\_rhino\_22Jul2017\_9M7eS\_haploidified\_headersFixed\_Sc9M7eS\_2\_HRSCAF\_41/{sample,[A-Za-z0-9]+}.merged.rmdup.merged.{processed}.Q30.bcf | |
| Container image | |
| docker://quay.io/biocontainers/bcftools:1.9--h68d8f2e\_9 | |
| Code | |
| ``` 1 2 ``` \| ``` bcftools mpileup -Ou -Q 30 -q 30 -B -f {input.ref} {input.bam} \| bcftools call -c -M -O b --threads {threads} -o {output.bcf} 2> {log} ``` | |

##### Rule index\_sorted\_vcfs

×

Rule properties

|  |  |
| --- | --- |
| Jobs | 6 |
| Input files | |
| - results/{dataset}/vcf/sumatran\_rhino\_22Jul2017\_9M7eS\_haploidified\_headersFixed\_Sc9M7eS\_2\_HRSCAF\_41/{sample}.merged.rmdup.merged.{processed}.Q30.sorted.bcf | |
| Output files | |
| - results/{dataset}/vcf/sumatran\_rhino\_22Jul2017\_9M7eS\_haploidified\_headersFixed\_Sc9M7eS\_2\_HRSCAF\_41/{sample,[A-Za-z0-9]+}.merged.rmdup.merged.{processed}.Q30.sorted.bcf.csi | |
| Container image | |
| docker://quay.io/biocontainers/bcftools:1.9--h68d8f2e\_9 | |
| Code | |
| ``` 1 2 ``` \| ``` bcftools index -o {output.index} {input.sort} 2> {log} ``` | |

##### Rule modern\_sorted\_vcf\_multiqc

×

Rule properties

|  |  |
| --- | --- |
| Jobs | 1 |
| Input files | |
| - results/modern/vcf/sumatran\_rhino\_22Jul2017\_9M7eS\_haploidified\_headersFixed\_Sc9M7eS\_2\_HRSCAF\_41/stats/vcf\_sorted/JvS033.merged.rmdup.merged.realn.mapped\_q30.subs\_dp6.Q30.sorted.vcf.stats.txt - results/modern/vcf/sumatran\_rhino\_22Jul2017\_9M7eS\_haploidified\_headersFixed\_Sc9M7eS\_2\_HRSCAF\_41/stats/vcf\_sorted/JvS035.merged.rmdup.merged.realn.mapped\_q30.subs\_dp6.Q30.sorted.vcf.stats.txt - results/modern/vcf/sumatran\_rhino\_22Jul2017\_9M7eS\_haploidified\_headersFixed\_Sc9M7eS\_2\_HRSCAF\_41/stats/vcf\_sorted/JvS034.merged.rmdup.merged.realn.mapped\_q30.subs\_dp6.Q30.sorted.vcf.stats.txt | |
| Output files | |
| - results/modern/vcf/sumatran\_rhino\_22Jul2017\_9M7eS\_haploidified\_headersFixed\_Sc9M7eS\_2\_HRSCAF\_41/stats/vcf\_sorted/multiqc/multiqc\_report.html | |
| Container image | |
| docker://quay.io/biocontainers/multiqc:1.9--pyh9f0ad1d\_0 | |
| Code | |
| ``` 1 2 ``` \| ``` multiqc -f {params.indir} -o {params.outdir} 2> {log} ``` | |

##### Rule historical\_CpG\_filtered\_vcf\_multiqc

×

Rule properties

|  |  |
| --- | --- |
| Jobs | 1 |
| Input files | |
| Output files | |
| - results/historical/vcf/sumatran\_rhino\_22Jul2017\_9M7eS\_haploidified\_headersFixed\_Sc9M7eS\_2\_HRSCAF\_41/stats/vcf\_CpG\_filtered/multiqc/multiqc\_report.html | |
| Container image | |
| docker://quay.io/biocontainers/multiqc:1.9--pyh9f0ad1d\_0 | |
| Code | |
| ``` 1 2 ``` \| ``` multiqc -f {params.indir} -o {params.outdir} 2> {log} ``` | |

##### Rule CpG\_filtered\_vcf\_stats

×

Rule properties

|  |  |
| --- | --- |
| Jobs | 6 |
| Input files | |
| - results/{dataset}/vcf/sumatran\_rhino\_22Jul2017\_9M7eS\_haploidified\_headersFixed\_Sc9M7eS\_2\_HRSCAF\_41/{sample}.merged.rmdup.merged.{processed}.Q30.sorted.no{CpG\_method}.bcf - results/{dataset}/vcf/sumatran\_rhino\_22Jul2017\_9M7eS\_haploidified\_headersFixed\_Sc9M7eS\_2\_HRSCAF\_41/{sample}.merged.rmdup.merged.{processed}.Q30.sorted.no{CpG\_method}.bcf.csi | |
| Output files | |
| - results/{dataset}/vcf/sumatran\_rhino\_22Jul2017\_9M7eS\_haploidified\_headersFixed\_Sc9M7eS\_2\_HRSCAF\_41/stats/vcf\_CpG\_filtered/{sample,[A-Za-z0-9]+}.merged.rmdup.merged.{processed}.Q30.sorted.no{CpG\_method,CpG\_[vcfre]{3,6}}.bcf.stats.txt | |
| Container image | |
| docker://quay.io/biocontainers/bcftools:1.9--h68d8f2e\_9 | |
| Code | |
| ``` 1 2 ``` \| ``` bcftools stats {input.bcf} > {output.stats} 2> {log} ``` | |

##### Rule CpG\_vcf2bcf

×

Rule properties

|  |  |
| --- | --- |
| Jobs | 6 |
| Input files | |
| - results/{dataset}/vcf/sumatran\_rhino\_22Jul2017\_9M7eS\_haploidified\_headersFixed\_Sc9M7eS\_2\_HRSCAF\_41/{sample}.merged.rmdup.merged.{processed}.Q30.sorted.no{CpG\_method}.vcf | |
| Output files | |
| - results/{dataset}/vcf/sumatran\_rhino\_22Jul2017\_9M7eS\_haploidified\_headersFixed\_Sc9M7eS\_2\_HRSCAF\_41/{sample,[A-Za-z0-9]+}.merged.rmdup.merged.{processed}.Q30.sorted.no{CpG\_method,CpG\_[vcfre]{3,6}}.bcf | |
| Container image | |
| docker://quay.io/biocontainers/bcftools:1.9--h68d8f2e\_9 | |
| Code | |
| ``` 1 2 ``` \| ``` bcftools convert -O b -o {output.bcf} {input.filtered} 2> {log} ``` | |

##### Rule remove\_CpG\_vcf

×

Rule properties

|  |  |
| --- | --- |
| Jobs | 6 |
| Input files | |
| - results/{dataset}/vcf/sumatran\_rhino\_22Jul2017\_9M7eS\_haploidified\_headersFixed\_Sc9M7eS\_2\_HRSCAF\_41/{sample}.merged.rmdup.merged.{processed}.Q30.sorted.CpG\_rm.vcf.gz - results/sumatran\_rhino\_22Jul2017\_9M7eS\_haploidified\_headersFixed\_Sc9M7eS\_2\_HRSCAF\_41.no{CpG\_method}.bed - /proj/sllstore2017093/b2016342/b2016342\_nobackup/lts/genome\_erosion\_pipeline/verena\_testing/testdata/reference/sumatran\_rhino\_22Jul2017\_9M7eS\_haploidified\_headersFixed\_Sc9M7eS\_2\_HRSCAF\_41.genome | |
| Output files | |
| - results/{dataset}/vcf/sumatran\_rhino\_22Jul2017\_9M7eS\_haploidified\_headersFixed\_Sc9M7eS\_2\_HRSCAF\_41/{sample,[A-Za-z0-9]+}.merged.rmdup.merged.{processed}.Q30.sorted.no{CpG\_method,CpG\_[vcfre]{3,6}}.vcf | |
| Container image | |
| docker://quay.io/biocontainers/bedtools:2.29.2--hc088bd4\_0 | |
| Code | |
| ``` 1 2 ``` \| ``` bedtools intersect -a {input.vcf} -b {input.bed} -header -sorted -g {input.genomefile} > {output.filtered} 2> {log} ``` | |

##### Rule sorted\_bcf2vcf\_CpG\_removal

×

Rule properties

|  |  |
| --- | --- |
| Jobs | 6 |
| Input files | |
| - results/{dataset}/vcf/sumatran\_rhino\_22Jul2017\_9M7eS\_haploidified\_headersFixed\_Sc9M7eS\_2\_HRSCAF\_41/{sample}.merged.rmdup.merged.{processed}.Q30.sorted.bcf | |
| Output files | |
| - results/{dataset}/vcf/sumatran\_rhino\_22Jul2017\_9M7eS\_haploidified\_headersFixed\_Sc9M7eS\_2\_HRSCAF\_41/{sample,[A-Za-z0-9]+}.merged.rmdup.merged.{processed}.Q30.sorted.CpG\_rm.vcf.gz | |
| Container image | |
| docker://quay.io/biocontainers/bcftools:1.9--h68d8f2e\_9 | |
| Code | |
| ``` 1 2 ``` \| ``` bcftools convert -O z -o {output.vcf} {input.bcf} 2> {log} ``` | |

##### Rule index\_CpG\_bcf

×

Rule properties

|  |  |
| --- | --- |
| Jobs | 6 |
| Input files | |
| - results/{dataset}/vcf/sumatran\_rhino\_22Jul2017\_9M7eS\_haploidified\_headersFixed\_Sc9M7eS\_2\_HRSCAF\_41/{sample}.merged.rmdup.merged.{processed}.Q30.sorted.no{CpG\_method}.bcf | |
| Output files | |
| - results/{dataset}/vcf/sumatran\_rhino\_22Jul2017\_9M7eS\_haploidified\_headersFixed\_Sc9M7eS\_2\_HRSCAF\_41/{sample,[A-Za-z0-9]+}.merged.rmdup.merged.{processed}.Q30.sorted.no{CpG\_method,CpG\_[vcfre]{3,6}}.bcf.csi | |
| Container image | |
| docker://quay.io/biocontainers/bcftools:1.9--h68d8f2e\_9 | |
| Code | |
| ``` 1 2 ``` \| ``` bcftools index -o {output.index} {input.bcf} 2> {log} ``` | |

##### Rule modern\_CpG\_filtered\_vcf\_multiqc

×

Rule properties

|  |  |
| --- | --- |
| Jobs | 1 |
| Input files | |
| Output files | |
| - results/modern/vcf/sumatran\_rhino\_22Jul2017\_9M7eS\_haploidified\_headersFixed\_Sc9M7eS\_2\_HRSCAF\_41/stats/vcf\_CpG\_filtered/multiqc/multiqc\_report.html | |
| Container image | |
| docker://quay.io/biocontainers/multiqc:1.9--pyh9f0ad1d\_0 | |
| Code | |
| ``` 1 2 ``` \| ``` multiqc -f {params.indir} -o {params.outdir} 2> {log} ``` | |

##### Rule historical\_quality\_filtered\_vcf\_multiqc

×

Rule properties

|  |  |
| --- | --- |
| Jobs | 1 |
| Input files | |
| Output files | |
| - results/historical/vcf/sumatran\_rhino\_22Jul2017\_9M7eS\_haploidified\_headersFixed\_Sc9M7eS\_2\_HRSCAF\_41/stats/vcf\_qual\_filtered/multiqc/multiqc\_report.html | |
| Container image | |
| docker://quay.io/biocontainers/multiqc:1.9--pyh9f0ad1d\_0 | |
| Code | |
| ``` 1 2 ``` \| ``` multiqc -f {params.indir} -o {params.outdir} 2> {log} ``` | |

##### Rule filtered\_vcf\_stats

×

Rule properties

|  |  |
| --- | --- |
| Jobs | 6 |
| Input files | |
| - results/{dataset}/vcf/sumatran\_rhino\_22Jul2017\_9M7eS\_haploidified\_headersFixed\_Sc9M7eS\_2\_HRSCAF\_41/{sample}.merged.rmdup.merged.{processed}.snps5.noIndel.QUAL30.dp.AB.bcf - results/{dataset}/vcf/sumatran\_rhino\_22Jul2017\_9M7eS\_haploidified\_headersFixed\_Sc9M7eS\_2\_HRSCAF\_41/{sample}.merged.rmdup.merged.{processed}.snps5.noIndel.QUAL30.dp.AB.bcf.csi | |
| Output files | |
| - results/{dataset}/vcf/sumatran\_rhino\_22Jul2017\_9M7eS\_haploidified\_headersFixed\_Sc9M7eS\_2\_HRSCAF\_41/stats/vcf\_qual\_filtered/{sample,[A-Za-z0-9]+}.merged.rmdup.merged.{processed}.snps5.noIndel.QUAL30.dp.AB.bcf.stats.txt | |
| Container image | |
| docker://quay.io/biocontainers/bcftools:1.9--h68d8f2e\_9 | |
| Code | |
| ``` 1 2 ``` \| ``` bcftools stats {input.bcf} > {output.stats} 2> {log} ``` | |

##### Rule filter\_vcfs\_allelic\_balance

×

Rule properties

|  |  |
| --- | --- |
| Jobs | 6 |
| Input files | |
| - results/{dataset}/vcf/sumatran\_rhino\_22Jul2017\_9M7eS\_haploidified\_headersFixed\_Sc9M7eS\_2\_HRSCAF\_41/{sample}.merged.rmdup.merged.{processed}.snps5.noIndel.QUAL30.dp.bcf | |
| Output files | |
| - results/{dataset}/vcf/sumatran\_rhino\_22Jul2017\_9M7eS\_haploidified\_headersFixed\_Sc9M7eS\_2\_HRSCAF\_41/{sample,[A-Za-z0-9]+}.merged.rmdup.merged.{processed}.snps5.noIndel.QUAL30.dp.AB.bcf | |
| Container image | |
| docker://quay.io/biocontainers/bcftools:1.9--h68d8f2e\_9 | |
| Code | |
| ``` 1 2 ``` \| ``` bcftools view -e 'GT="0/1" & (DP4[2]+DP4[3])/(DP4[0]+DP4[1]+DP4[2]+DP4[3]) < 0.2' {input.bcf} \| bcftools view -e 'GT="0/1" & (DP4[2]+DP4[3])/(DP4[0]+DP4[1]+DP4[2]+DP4[3]) > 0.8' -Ob > {output.filtered} 2> {log} ``` | |

##### Rule filter\_vcfs\_qual\_dp

×

Rule properties

|  |  |
| --- | --- |
| Jobs | 6 |
| Input files | |
| - results/{dataset}/vcf/sumatran\_rhino\_22Jul2017\_9M7eS\_haploidified\_headersFixed\_Sc9M7eS\_2\_HRSCAF\_41/{sample}.merged.rmdup.merged.{processed}.snps5.bcf | |
| Output files | |
| - results/{dataset}/vcf/sumatran\_rhino\_22Jul2017\_9M7eS\_haploidified\_headersFixed\_Sc9M7eS\_2\_HRSCAF\_41/{sample,[A-Za-z0-9]+}.merged.rmdup.merged.{processed}.snps5.noIndel.QUAL30.dp.bcf | |
| Container image | |
| docker://quay.io/biocontainers/bcftools:1.9--h68d8f2e\_9 | |
| Code | |
| ``` 1 2 3 4 5 6 7 8 9 10 11 ``` \| ``` minDP=`head -n 1 {input.dp} \| cut -d' ' -f 2` maxDP=`head -n 1 {input.dp} \| cut -d' ' -f 3` # check minimum depth threshold if awk "BEGIN{{exit ! ($minDP < 3)}}" then minDP=3 fi bcftools filter -i "(DP4[0]+DP4[1]+DP4[2]+DP4[3])>$minDP & (DP4[0]+DP4[1]+DP4[2]+DP4[3])<$maxDP & QUAL>=30 & INDEL=0" -O b --threads {threads} -o {output.filtered} {input.bcf} 2> {log} ``` | |

##### Rule remove\_snps\_near\_indels

×

Rule properties

|  |  |
| --- | --- |
| Jobs | 6 |
| Input files | |
| - results/{dataset}/vcf/sumatran\_rhino\_22Jul2017\_9M7eS\_haploidified\_headersFixed\_Sc9M7eS\_2\_HRSCAF\_41/{sample}.merged.rmdup.merged.{processed}.bcf | |
| Output files | |
| - results/{dataset}/vcf/sumatran\_rhino\_22Jul2017\_9M7eS\_haploidified\_headersFixed\_Sc9M7eS\_2\_HRSCAF\_41/{sample,[A-Za-z0-9]+}.merged.rmdup.merged.{processed}.snps5.bcf | |
| Container image | |
| docker://quay.io/biocontainers/bcftools:1.9--h68d8f2e\_9 | |
| Code | |
| ``` 1 2 ``` \| ``` bcftools filter -g 5 -O b --threads {threads} -o {output.snps} {input.bcf} 2> {log} ``` | |

##### Rule index\_filtered\_vcfs

×

Rule properties

|  |  |
| --- | --- |
| Jobs | 6 |
| Input files | |
| - results/{dataset}/vcf/sumatran\_rhino\_22Jul2017\_9M7eS\_haploidified\_headersFixed\_Sc9M7eS\_2\_HRSCAF\_41/{sample}.merged.rmdup.merged.{processed}.snps5.noIndel.QUAL30.dp.AB.bcf | |
| Output files | |
| - results/{dataset}/vcf/sumatran\_rhino\_22Jul2017\_9M7eS\_haploidified\_headersFixed\_Sc9M7eS\_2\_HRSCAF\_41/{sample,[A-Za-z0-9]+}.merged.rmdup.merged.{processed}.snps5.noIndel.QUAL30.dp.AB.bcf.csi | |
| Container image | |
| docker://quay.io/biocontainers/bcftools:1.9--h68d8f2e\_9 | |
| Code | |
| ``` 1 2 ``` \| ``` bcftools index -o {output.index} {input.bcf} 2> {log} ``` | |

##### Rule historical\_repmasked\_vcf\_multiqc

×

Rule properties

|  |  |
| --- | --- |
| Jobs | 1 |
| Input files | |
| Output files | |
| - results/historical/vcf/sumatran\_rhino\_22Jul2017\_9M7eS\_haploidified\_headersFixed\_Sc9M7eS\_2\_HRSCAF\_41/stats/vcf\_repmasked/multiqc/multiqc\_report.html | |
| Container image | |
| docker://quay.io/biocontainers/multiqc:1.9--pyh9f0ad1d\_0 | |
| Code | |
| ``` 1 2 ``` \| ``` multiqc -f {params.indir} -o {params.outdir} 2> {log} ``` | |

##### Rule repmasked\_vcf\_stats

×

Rule properties

|  |  |
| --- | --- |
| Jobs | 6 |
| Input files | |
| - results/{dataset}/vcf/sumatran\_rhino\_22Jul2017\_9M7eS\_haploidified\_headersFixed\_Sc9M7eS\_2\_HRSCAF\_41/{sample}.merged.rmdup.merged.{processed}.snps5.noIndel.QUAL30.dp.AB.repma.bcf - results/{dataset}/vcf/sumatran\_rhino\_22Jul2017\_9M7eS\_haploidified\_headersFixed\_Sc9M7eS\_2\_HRSCAF\_41/{sample}.merged.rmdup.merged.{processed}.snps5.noIndel.QUAL30.dp.AB.repma.bcf.csi | |
| Output files | |
| - results/{dataset}/vcf/sumatran\_rhino\_22Jul2017\_9M7eS\_haploidified\_headersFixed\_Sc9M7eS\_2\_HRSCAF\_41/stats/vcf\_repmasked/{sample,[A-Za-z0-9]+}.merged.rmdup.merged.{processed}.snps5.noIndel.QUAL30.dp.AB.repma.bcf.stats.txt | |
| Container image | |
| docker://quay.io/biocontainers/bcftools:1.9--h68d8f2e\_9 | |
| Code | |
| ``` 1 2 ``` \| ``` bcftools stats {input.bcf} > {output.stats} 2> {log} ``` | |

##### Rule filtered\_vcf2bcf

×

Rule properties

|  |  |
| --- | --- |
| Jobs | 6 |
| Input files | |
| - results/{dataset}/vcf/sumatran\_rhino\_22Jul2017\_9M7eS\_haploidified\_headersFixed\_Sc9M7eS\_2\_HRSCAF\_41/{sample}.merged.rmdup.merged.{processed}.snps5.noIndel.QUAL30.dp.AB.repma.vcf | |
| Output files | |
| - results/{dataset}/vcf/sumatran\_rhino\_22Jul2017\_9M7eS\_haploidified\_headersFixed\_Sc9M7eS\_2\_HRSCAF\_41/{sample,[A-Za-z0-9]+}.merged.rmdup.merged.{processed}.snps5.noIndel.QUAL30.dp.AB.repma.bcf | |
| Container image | |
| docker://quay.io/biocontainers/bcftools:1.9--h68d8f2e\_9 | |
| Code | |
| ``` 1 2 ``` \| ``` bcftools convert -O b -o {output.bcf} {input.filtered} 2> {log} ``` | |

##### Rule remove\_repeats\_vcf

×

Rule properties

|  |  |
| --- | --- |
| Jobs | 6 |
| Input files | |
| - results/{dataset}/vcf/sumatran\_rhino\_22Jul2017\_9M7eS\_haploidified\_headersFixed\_Sc9M7eS\_2\_HRSCAF\_41/{sample}.merged.rmdup.merged.{processed}.snps5.noIndel.QUAL30.dp.AB.vcf.gz - results/sumatran\_rhino\_22Jul2017\_9M7eS\_haploidified\_headersFixed\_Sc9M7eS\_2\_HRSCAF\_41.repma.bed - /proj/sllstore2017093/b2016342/b2016342\_nobackup/lts/genome\_erosion\_pipeline/verena\_testing/testdata/reference/sumatran\_rhino\_22Jul2017\_9M7eS\_haploidified\_headersFixed\_Sc9M7eS\_2\_HRSCAF\_41.genome | |
| Output files | |
| - results/{dataset}/vcf/sumatran\_rhino\_22Jul2017\_9M7eS\_haploidified\_headersFixed\_Sc9M7eS\_2\_HRSCAF\_41/{sample,[A-Za-z0-9]+}.merged.rmdup.merged.{processed}.snps5.noIndel.QUAL30.dp.AB.repma.vcf | |
| Container image | |
| docker://quay.io/biocontainers/bedtools:2.29.2--hc088bd4\_0 | |
| Code | |
| ``` 1 2 ``` \| ``` bedtools intersect -a {input.vcf} -b {input.bed} -header -sorted -g {input.genomefile} > {output.filtered} 2> {log} ``` | |

##### Rule filtered\_bcf2vcf

×

Rule properties

|  |  |
| --- | --- |
| Jobs | 6 |
| Input files | |
| - results/{dataset}/vcf/sumatran\_rhino\_22Jul2017\_9M7eS\_haploidified\_headersFixed\_Sc9M7eS\_2\_HRSCAF\_41/{sample}.merged.rmdup.merged.{processed}.snps5.noIndel.QUAL30.dp.AB.bcf | |
| Output files | |
| - results/{dataset}/vcf/sumatran\_rhino\_22Jul2017\_9M7eS\_haploidified\_headersFixed\_Sc9M7eS\_2\_HRSCAF\_41/{sample,[A-Za-z0-9]+}.merged.rmdup.merged.{processed}.snps5.noIndel.QUAL30.dp.AB.vcf.gz | |
| Container image | |
| docker://quay.io/biocontainers/bcftools:1.9--h68d8f2e\_9 | |
| Code | |
| ``` 1 2 ``` \| ``` bcftools convert -O z -o {output.vcf} {input.bcf} 2> {log} ``` | |

##### Rule index\_repmasked\_vcfs

×

Rule properties

|  |  |
| --- | --- |
| Jobs | 6 |
| Input files | |
| - results/{dataset}/vcf/sumatran\_rhino\_22Jul2017\_9M7eS\_haploidified\_headersFixed\_Sc9M7eS\_2\_HRSCAF\_41/{sample}.merged.rmdup.merged.{processed}.snps5.noIndel.QUAL30.dp.AB.repma.bcf | |
| Output files | |
| - results/{dataset}/vcf/sumatran\_rhino\_22Jul2017\_9M7eS\_haploidified\_headersFixed\_Sc9M7eS\_2\_HRSCAF\_41/{sample,[A-Za-z0-9]+}.merged.rmdup.merged.{processed}.snps5.noIndel.QUAL30.dp.AB.repma.bcf.csi | |
| Container image | |
| docker://quay.io/biocontainers/bcftools:1.9--h68d8f2e\_9 | |
| Code | |
| ``` 1 2 ``` \| ``` bcftools index -o {output.index} {input.bcf} 2> {log} ``` | |

##### Rule modern\_quality\_filtered\_vcf\_multiqc

×

Rule properties

|  |  |
| --- | --- |
| Jobs | 1 |
| Input files | |
| Output files | |
| - results/modern/vcf/sumatran\_rhino\_22Jul2017\_9M7eS\_haploidified\_headersFixed\_Sc9M7eS\_2\_HRSCAF\_41/stats/vcf\_qual\_filtered/multiqc/multiqc\_report.html | |
| Container image | |
| docker://quay.io/biocontainers/multiqc:1.9--pyh9f0ad1d\_0 | |
| Code | |
| ``` 1 2 ``` \| ``` multiqc -f {params.indir} -o {params.outdir} 2> {log} ``` | |

##### Rule modern\_repmasked\_vcf\_multiqc

×

Rule properties

|  |  |
| --- | --- |
| Jobs | 1 |
| Input files | |
| Output files | |
| - results/modern/vcf/sumatran\_rhino\_22Jul2017\_9M7eS\_haploidified\_headersFixed\_Sc9M7eS\_2\_HRSCAF\_41/stats/vcf\_repmasked/multiqc/multiqc\_report.html | |
| Container image | |
| docker://quay.io/biocontainers/multiqc:1.9--pyh9f0ad1d\_0 | |
| Code | |
| ``` 1 2 ``` \| ``` multiqc -f {params.indir} -o {params.outdir} 2> {log} ``` | |

##### Rule missingness\_filtered\_vcf\_multiqc

×

Rule properties

|  |  |
| --- | --- |
| Jobs | 1 |
| Input files | |
| Output files | |
| - results/all/vcf/sumatran\_rhino\_22Jul2017\_9M7eS\_haploidified\_headersFixed\_Sc9M7eS\_2\_HRSCAF\_41/stats/vcf\_merged\_missing/multiqc/multiqc\_report.html | |
| Container image | |
| docker://quay.io/biocontainers/multiqc:1.9--pyh9f0ad1d\_0 | |
| Code | |
| ``` 1 2 ``` \| ``` multiqc -f {params.indir} -o {params.outdir} 2> {log} ``` | |

##### Rule missingness\_filtered\_vcf\_stats

×

Rule properties

|  |  |
| --- | --- |
| Jobs | 3 |
| Input files | |
| - results/{dataset}/vcf/sumatran\_rhino\_22Jul2017\_9M7eS\_haploidified\_headersFixed\_Sc9M7eS\_2\_HRSCAF\_41.{dataset}.merged.biallelic.fmissing{fmiss}.vcf.gz - results/{dataset}/vcf/sumatran\_rhino\_22Jul2017\_9M7eS\_haploidified\_headersFixed\_Sc9M7eS\_2\_HRSCAF\_41.{dataset}.merged.biallelic.fmissing{fmiss}.vcf.gz.csi | |
| Output files | |
| - results/{dataset}/vcf/sumatran\_rhino\_22Jul2017\_9M7eS\_haploidified\_headersFixed\_Sc9M7eS\_2\_HRSCAF\_41/stats/vcf\_merged\_missing/sumatran\_rhino\_22Jul2017\_9M7eS\_haploidified\_headersFixed\_Sc9M7eS\_2\_HRSCAF\_41.{dataset}.merged.biallelic.fmissing{fmiss}.vcf.stats.txt | |
| Container image | |
| docker://quay.io/biocontainers/bcftools:1.9--h68d8f2e\_9 | |
| Code | |
| ``` 1 2 ``` \| ``` bcftools stats {input.merged} > {output.stats} 2> {log} ``` | |

##### Rule filter\_vcf\_missing

×

Rule properties

|  |  |
| --- | --- |
| Jobs | 1 |
| Input files | |
| - results/all/vcf/sumatran\_rhino\_22Jul2017\_9M7eS\_haploidified\_headersFixed\_Sc9M7eS\_2\_HRSCAF\_41.all.merged.biallelic.bcf - results/all/vcf/sumatran\_rhino\_22Jul2017\_9M7eS\_haploidified\_headersFixed\_Sc9M7eS\_2\_HRSCAF\_41.all.merged.biallelic.bcf.csi - results/all/vcf/sumatran\_rhino\_22Jul2017\_9M7eS\_haploidified\_headersFixed\_Sc9M7eS\_2\_HRSCAF\_41/stats/vcf\_merged\_biallelic/sumatran\_rhino\_22Jul2017\_9M7eS\_haploidified\_headersFixed\_Sc9M7eS\_2\_HRSCAF\_41.all.merged.biallelic.vcf.stats.txt - results/all/vcf/sumatran\_rhino\_22Jul2017\_9M7eS\_haploidified\_headersFixed\_Sc9M7eS\_2\_HRSCAF\_41/stats/vcf\_merged\_biallelic/multiqc/multiqc\_report.html | |
| Output files | |
| - results/all/vcf/sumatran\_rhino\_22Jul2017\_9M7eS\_haploidified\_headersFixed\_Sc9M7eS\_2\_HRSCAF\_41.all.merged.biallelic.fmissing{fmiss}.vcf.gz - results/all/vcf/sumatran\_rhino\_22Jul2017\_9M7eS\_haploidified\_headersFixed\_Sc9M7eS\_2\_HRSCAF\_41.all.merged.biallelic.fmissing{fmiss}.vcf.gz.csi | |
| Container image | |
| docker://quay.io/biocontainers/bcftools:1.9--h68d8f2e\_9 | |
| Code | |
| ``` 1 2 3 ``` \| ``` bcftools view -i 'F_MISSING < {params.fmiss}' -Oz -o {output.vcf} {input.bcf} 2> {log} && bcftools index -f {output.vcf} 2>> {log} ``` | |

##### Rule filter\_vcf\_biallelic

×

Rule properties

|  |  |
| --- | --- |
| Jobs | 1 |
| Input files | |
| - results/all/vcf/sumatran\_rhino\_22Jul2017\_9M7eS\_haploidified\_headersFixed\_Sc9M7eS\_2\_HRSCAF\_41.all.merged.snps.bcf - results/all/vcf/sumatran\_rhino\_22Jul2017\_9M7eS\_haploidified\_headersFixed\_Sc9M7eS\_2\_HRSCAF\_41.all.merged.snps.bcf.csi - results/all/vcf/sumatran\_rhino\_22Jul2017\_9M7eS\_haploidified\_headersFixed\_Sc9M7eS\_2\_HRSCAF\_41/stats/vcf\_merged/sumatran\_rhino\_22Jul2017\_9M7eS\_haploidified\_headersFixed\_Sc9M7eS\_2\_HRSCAF\_41.all.merged.snps.bcf.stats.txt - results/all/vcf/sumatran\_rhino\_22Jul2017\_9M7eS\_haploidified\_headersFixed\_Sc9M7eS\_2\_HRSCAF\_41/stats/vcf\_merged/multiqc/multiqc\_report.html | |
| Output files | |
| - results/all/vcf/sumatran\_rhino\_22Jul2017\_9M7eS\_haploidified\_headersFixed\_Sc9M7eS\_2\_HRSCAF\_41.all.merged.biallelic.bcf - results/all/vcf/sumatran\_rhino\_22Jul2017\_9M7eS\_haploidified\_headersFixed\_Sc9M7eS\_2\_HRSCAF\_41.all.merged.biallelic.bcf.csi | |
| Container image | |
| docker://quay.io/biocontainers/bcftools:1.9--h68d8f2e\_9 | |
| Code | |
| ``` 1 2 3 ``` \| ``` bcftools view -m2 -M2 -v snps -Ob -o {output.bcf} {input.bcf} 2> {log} && bcftools index -f {output.bcf} 2>> {log} ``` | |

##### Rule merge\_all\_vcfs

×

Rule properties

|  |  |
| --- | --- |
| Jobs | 1 |
| Input files | |
| Output files | |
| - results/all/vcf/sumatran\_rhino\_22Jul2017\_9M7eS\_haploidified\_headersFixed\_Sc9M7eS\_2\_HRSCAF\_41.all.merged.snps.bcf | |
| Container image | |
| docker://quay.io/biocontainers/bcftools:1.9--h68d8f2e\_9 | |
| Code | |
| ``` 1 2 3 4 5 6 7 8 9 ``` \| ``` files=`echo {input.bcf} \| awk '{{print NF}}'` if [ $files -gt 1 ] # check if there are at least 2 files for merging. If there is only one file, copy the bcf file. then bcftools merge -m snps -O b -o {output.merged} {input.bcf} 2> {log} else cp {input.bcf} {output.merged} && touch {output.merged} 2> {log} echo "Only one file present for merging. Copying the input bcf file." >> {log} fi ``` | |

##### Rule index\_merged\_vcf

×

Rule properties

|  |  |
| --- | --- |
| Jobs | 1 |
| Input files | |
| - results/all/vcf/sumatran\_rhino\_22Jul2017\_9M7eS\_haploidified\_headersFixed\_Sc9M7eS\_2\_HRSCAF\_41.all.merged.snps.bcf | |
| Output files | |
| - results/all/vcf/sumatran\_rhino\_22Jul2017\_9M7eS\_haploidified\_headersFixed\_Sc9M7eS\_2\_HRSCAF\_41.all.merged.snps.bcf.csi | |
| Container image | |
| docker://quay.io/biocontainers/bcftools:1.9--h68d8f2e\_9 | |
| Code | |
| ``` 1 2 ``` \| ``` bcftools index -o {output.index} {input.bcf} 2> {log} ``` | |

##### Rule merged\_vcf\_stats

×

Rule properties

|  |  |
| --- | --- |
| Jobs | 1 |
| Input files | |
| - results/all/vcf/sumatran\_rhino\_22Jul2017\_9M7eS\_haploidified\_headersFixed\_Sc9M7eS\_2\_HRSCAF\_41.all.merged.snps.bcf - results/all/vcf/sumatran\_rhino\_22Jul2017\_9M7eS\_haploidified\_headersFixed\_Sc9M7eS\_2\_HRSCAF\_41.all.merged.snps.bcf.csi | |
| Output files | |
| - results/all/vcf/sumatran\_rhino\_22Jul2017\_9M7eS\_haploidified\_headersFixed\_Sc9M7eS\_2\_HRSCAF\_41/stats/vcf\_merged/sumatran\_rhino\_22Jul2017\_9M7eS\_haploidified\_headersFixed\_Sc9M7eS\_2\_HRSCAF\_41.all.merged.snps.bcf.stats.txt | |
| Container image | |
| docker://quay.io/biocontainers/bcftools:1.9--h68d8f2e\_9 | |
| Code | |
| ``` 1 2 ``` \| ``` bcftools stats {input.merged} > {output.stats} 2> {log} ``` | |

##### Rule merged\_vcf\_multiqc

×

Rule properties

|  |  |
| --- | --- |
| Jobs | 1 |
| Input files | |
| - results/all/vcf/sumatran\_rhino\_22Jul2017\_9M7eS\_haploidified\_headersFixed\_Sc9M7eS\_2\_HRSCAF\_41/stats/vcf\_merged/sumatran\_rhino\_22Jul2017\_9M7eS\_haploidified\_headersFixed\_Sc9M7eS\_2\_HRSCAF\_41.all.merged.snps.bcf.stats.txt | |
| Output files | |
| - results/all/vcf/sumatran\_rhino\_22Jul2017\_9M7eS\_haploidified\_headersFixed\_Sc9M7eS\_2\_HRSCAF\_41/stats/vcf\_merged/multiqc/multiqc\_report.html | |
| Container image | |
| docker://quay.io/biocontainers/multiqc:1.9--pyh9f0ad1d\_0 | |
| Code | |
| ``` 1 2 ``` \| ``` multiqc -f {params.indir} -o {params.outdir} 2> {log} ``` | |

##### Rule biallelic\_filtered\_vcf\_stats

×

Rule properties

|  |  |
| --- | --- |
| Jobs | 1 |
| Input files | |
| - results/all/vcf/sumatran\_rhino\_22Jul2017\_9M7eS\_haploidified\_headersFixed\_Sc9M7eS\_2\_HRSCAF\_41.all.merged.biallelic.bcf - results/all/vcf/sumatran\_rhino\_22Jul2017\_9M7eS\_haploidified\_headersFixed\_Sc9M7eS\_2\_HRSCAF\_41.all.merged.biallelic.bcf.csi | |
| Output files | |
| - results/all/vcf/sumatran\_rhino\_22Jul2017\_9M7eS\_haploidified\_headersFixed\_Sc9M7eS\_2\_HRSCAF\_41/stats/vcf\_merged\_biallelic/sumatran\_rhino\_22Jul2017\_9M7eS\_haploidified\_headersFixed\_Sc9M7eS\_2\_HRSCAF\_41.all.merged.biallelic.vcf.stats.txt | |
| Container image | |
| docker://quay.io/biocontainers/bcftools:1.9--h68d8f2e\_9 | |
| Code | |
| ``` 1 2 ``` \| ``` bcftools stats {input.bcf} > {output.stats} 2> {log} ``` | |

##### Rule biallelic\_filtered\_vcf\_multiqc

×

Rule properties

|  |  |
| --- | --- |
| Jobs | 1 |
| Input files | |
| - results/all/vcf/sumatran\_rhino\_22Jul2017\_9M7eS\_haploidified\_headersFixed\_Sc9M7eS\_2\_HRSCAF\_41/stats/vcf\_merged\_biallelic/sumatran\_rhino\_22Jul2017\_9M7eS\_haploidified\_headersFixed\_Sc9M7eS\_2\_HRSCAF\_41.all.merged.biallelic.vcf.stats.txt | |
| Output files | |
| - results/all/vcf/sumatran\_rhino\_22Jul2017\_9M7eS\_haploidified\_headersFixed\_Sc9M7eS\_2\_HRSCAF\_41/stats/vcf\_merged\_biallelic/multiqc/multiqc\_report.html | |
| Container image | |
| docker://quay.io/biocontainers/multiqc:1.9--pyh9f0ad1d\_0 | |
| Code | |
| ``` 1 2 ``` \| ``` multiqc -f {params.indir} -o {params.outdir} 2> {log} ``` | |

##### Rule extract\_historical\_samples

×

Rule properties

|  |  |
| --- | --- |
| Jobs | 1 |
| Input files | |
| - results/all/vcf/sumatran\_rhino\_22Jul2017\_9M7eS\_haploidified\_headersFixed\_Sc9M7eS\_2\_HRSCAF\_41.all.merged.biallelic.fmissing{fmiss}.vcf.gz - results/all/vcf/sumatran\_rhino\_22Jul2017\_9M7eS\_haploidified\_headersFixed\_Sc9M7eS\_2\_HRSCAF\_41.all.merged.biallelic.fmissing{fmiss}.vcf.gz.csi - results/all/vcf/sumatran\_rhino\_22Jul2017\_9M7eS\_haploidified\_headersFixed\_Sc9M7eS\_2\_HRSCAF\_41/stats/vcf\_merged\_biallelic/sumatran\_rhino\_22Jul2017\_9M7eS\_haploidified\_headersFixed\_Sc9M7eS\_2\_HRSCAF\_41.all.merged.biallelic.vcf.stats.txt - results/all/vcf/sumatran\_rhino\_22Jul2017\_9M7eS\_haploidified\_headersFixed\_Sc9M7eS\_2\_HRSCAF\_41.all.merged.biallelic.fmissing{fmiss}.bed | |
| Output files | |
| - results/historical/vcf/sumatran\_rhino\_22Jul2017\_9M7eS\_haploidified\_headersFixed\_Sc9M7eS\_2\_HRSCAF\_41.historical.merged.biallelic.fmissing{fmiss}.vcf.gz - results/historical/vcf/sumatran\_rhino\_22Jul2017\_9M7eS\_haploidified\_headersFixed\_Sc9M7eS\_2\_HRSCAF\_41.historical.merged.biallelic.fmissing{fmiss}.vcf.gz.csi | |
| Container image | |
| docker://quay.io/biocontainers/bcftools:1.9--h68d8f2e\_9 | |
| Code | |
| ``` 1 2 3 4 5 6 7 8 9 10 11 12 13 14 ``` \| ``` samples_edited=`echo {params.samples} \| sed 's/ /,/g'` samples_len=`echo {params.samples} \| wc -w` # count the number of historical samples all_samples_len=`echo {params.all_samples} \| wc -w` # count the number of all samples if [ $samples_len != $all_samples_len ] then bcftools view -Oz -s $samples_edited -o {output.vcf} {input.vcf} 2> {log} && bcftools index -f {output.vcf} 2>> {log} else cp {input.vcf} {output.vcf} && touch {output.vcf} 2> {log} && bcftools index -f {output.vcf} 2>> {log} echo "Only historical samples present. Copying the input vcf file." >> {log} fi ``` | |

##### Rule filtered\_vcf2bed

×

Rule properties

|  |  |
| --- | --- |
| Jobs | 1 |
| Input files | |
| - results/all/vcf/sumatran\_rhino\_22Jul2017\_9M7eS\_haploidified\_headersFixed\_Sc9M7eS\_2\_HRSCAF\_41.all.merged.biallelic.fmissing{fmiss}.vcf.gz | |
| Output files | |
| - results/all/vcf/sumatran\_rhino\_22Jul2017\_9M7eS\_haploidified\_headersFixed\_Sc9M7eS\_2\_HRSCAF\_41.all.merged.biallelic.fmissing{fmiss}.bed | |
| Container image | |
| docker://quay.io/biocontainers/bedtools:2.29.2--hc088bd4\_0 | |
| Code | |
| ``` 1 2 ``` \| ``` gzip -cd {input.vcf} \| grep -v "^#" \| awk -F'	' '{{print $1, $2-1, $2}}' OFS='	' > {output.bed} 2> {log} ``` | |

##### Rule extract\_modern\_samples

×

Rule properties

|  |  |
| --- | --- |
| Jobs | 1 |
| Input files | |
| - results/all/vcf/sumatran\_rhino\_22Jul2017\_9M7eS\_haploidified\_headersFixed\_Sc9M7eS\_2\_HRSCAF\_41.all.merged.biallelic.fmissing{fmiss}.vcf.gz - results/all/vcf/sumatran\_rhino\_22Jul2017\_9M7eS\_haploidified\_headersFixed\_Sc9M7eS\_2\_HRSCAF\_41.all.merged.biallelic.fmissing{fmiss}.vcf.gz.csi - results/all/vcf/sumatran\_rhino\_22Jul2017\_9M7eS\_haploidified\_headersFixed\_Sc9M7eS\_2\_HRSCAF\_41/stats/vcf\_merged\_biallelic/sumatran\_rhino\_22Jul2017\_9M7eS\_haploidified\_headersFixed\_Sc9M7eS\_2\_HRSCAF\_41.all.merged.biallelic.vcf.stats.txt - results/all/vcf/sumatran\_rhino\_22Jul2017\_9M7eS\_haploidified\_headersFixed\_Sc9M7eS\_2\_HRSCAF\_41.all.merged.biallelic.fmissing{fmiss}.bed | |
| Output files | |
| - results/modern/vcf/sumatran\_rhino\_22Jul2017\_9M7eS\_haploidified\_headersFixed\_Sc9M7eS\_2\_HRSCAF\_41.modern.merged.biallelic.fmissing{fmiss}.vcf.gz - results/modern/vcf/sumatran\_rhino\_22Jul2017\_9M7eS\_haploidified\_headersFixed\_Sc9M7eS\_2\_HRSCAF\_41.modern.merged.biallelic.fmissing{fmiss}.vcf.gz.csi | |
| Container image | |
| docker://quay.io/biocontainers/bcftools:1.9--h68d8f2e\_9 | |
| Code | |
| ``` 1 2 3 4 5 6 7 8 9 10 11 12 13 14 ``` \| ``` samples_edited=`echo {params.samples} \| sed 's/ /,/g'` samples_len=`echo {params.samples} \| wc -w` # count the number of historical samples all_samples_len=`echo {params.all_samples} \| wc -w` # count the number of all samples if [ $samples_len != $all_samples_len ] then bcftools view -Oz -s $samples_edited -o {output.vcf} {input.vcf} 2> {log} && bcftools index -f {output.vcf} 2>> {log} else cp {input.vcf} {output.vcf} && touch {output.vcf} 2> {log} && bcftools index -f {output.vcf} 2>> {log} echo "Only modern samples present. Copying the input vcf file." >> {log} fi ``` | |

##### Rule plot\_pc1\_pc2

×

Rule properties

|  |  |
| --- | --- |
| Jobs | 3 |
| Input files | |
| - results/{dataset}/pca/sumatran\_rhino\_22Jul2017\_9M7eS\_haploidified\_headersFixed\_Sc9M7eS\_2\_HRSCAF\_41.{dataset}.merged.biallelic.fmissing{fmiss}.eigenvec - results/{dataset}/pca/sumatran\_rhino\_22Jul2017\_9M7eS\_haploidified\_headersFixed\_Sc9M7eS\_2\_HRSCAF\_41.{dataset}.merged.biallelic.fmissing{fmiss}.eigenval | |
| Output files | |
| - results/{dataset}/pca/sumatran\_rhino\_22Jul2017\_9M7eS\_haploidified\_headersFixed\_Sc9M7eS\_2\_HRSCAF\_41.{dataset}.merged.biallelic.fmissing{fmiss}.pc1\_pc2.pdf | |

##### Rule plink\_eigenvec

×

Rule properties

|  |  |
| --- | --- |
| Jobs | 3 |
| Input files | |
| - results/{dataset}/pca/sumatran\_rhino\_22Jul2017\_9M7eS\_haploidified\_headersFixed\_Sc9M7eS\_2\_HRSCAF\_41.{dataset}.merged.biallelic.fmissing{fmiss}.bed - results/{dataset}/pca/sumatran\_rhino\_22Jul2017\_9M7eS\_haploidified\_headersFixed\_Sc9M7eS\_2\_HRSCAF\_41.{dataset}.merged.biallelic.fmissing{fmiss}.bim - results/{dataset}/pca/sumatran\_rhino\_22Jul2017\_9M7eS\_haploidified\_headersFixed\_Sc9M7eS\_2\_HRSCAF\_41.{dataset}.merged.biallelic.fmissing{fmiss}.fam - results/{dataset}/pca/sumatran\_rhino\_22Jul2017\_9M7eS\_haploidified\_headersFixed\_Sc9M7eS\_2\_HRSCAF\_41.{dataset}.merged.biallelic.fmissing{fmiss}.nosex | |
| Output files | |
| - results/{dataset}/pca/sumatran\_rhino\_22Jul2017\_9M7eS\_haploidified\_headersFixed\_Sc9M7eS\_2\_HRSCAF\_41.{dataset}.merged.biallelic.fmissing{fmiss}.eigenvec - results/{dataset}/pca/sumatran\_rhino\_22Jul2017\_9M7eS\_haploidified\_headersFixed\_Sc9M7eS\_2\_HRSCAF\_41.{dataset}.merged.biallelic.fmissing{fmiss}.eigenval | |
| Container image | |
| docker://quay.io/biocontainers/plink:1.90b6.12--heea4ae3\_0 | |
| Code | |
| ``` 1 2 3 4 5 6 7 8 9 ``` \| ``` samples=`cat {input.fam} \| wc -l` if [ "$samples" -gt 1 ] then plink --bfile {params.bfile} --allow-extra-chr --pca --out {params.bfile} 2> {log} else touch {output.eigenvec} && touch {output.eigenval} 2> {log} echo "Not enough samples to calculate a PCA." >> {log} fi ``` | |

##### Rule vcf2plink\_pca

×

Rule properties

|  |  |
| --- | --- |
| Jobs | 3 |
| Input files | |
| - results/{dataset}/vcf/sumatran\_rhino\_22Jul2017\_9M7eS\_haploidified\_headersFixed\_Sc9M7eS\_2\_HRSCAF\_41.{dataset}.merged.biallelic.fmissing{fmiss}.vcf.gz - results/{dataset}/vcf/sumatran\_rhino\_22Jul2017\_9M7eS\_haploidified\_headersFixed\_Sc9M7eS\_2\_HRSCAF\_41.{dataset}.merged.biallelic.fmissing{fmiss}.vcf.gz.csi | |
| Output files | |
| - results/{dataset}/pca/sumatran\_rhino\_22Jul2017\_9M7eS\_haploidified\_headersFixed\_Sc9M7eS\_2\_HRSCAF\_41.{dataset}.merged.biallelic.fmissing{fmiss}.bed - results/{dataset}/pca/sumatran\_rhino\_22Jul2017\_9M7eS\_haploidified\_headersFixed\_Sc9M7eS\_2\_HRSCAF\_41.{dataset}.merged.biallelic.fmissing{fmiss}.bim - results/{dataset}/pca/sumatran\_rhino\_22Jul2017\_9M7eS\_haploidified\_headersFixed\_Sc9M7eS\_2\_HRSCAF\_41.{dataset}.merged.biallelic.fmissing{fmiss}.fam - results/{dataset}/pca/sumatran\_rhino\_22Jul2017\_9M7eS\_haploidified\_headersFixed\_Sc9M7eS\_2\_HRSCAF\_41.{dataset}.merged.biallelic.fmissing{fmiss}.nosex | |
| Container image | |
| docker://quay.io/biocontainers/plink:1.90b6.12--heea4ae3\_0 | |
| Code | |
| ``` 1 2 ``` \| ``` plink --vcf {input.vcf} --make-bed --allow-extra-chr --out {params.bfile} 2> {log} ``` | |

##### Rule plot\_pc1\_pc3

×

Rule properties

|  |  |
| --- | --- |
| Jobs | 3 |
| Input files | |
| - results/{dataset}/pca/sumatran\_rhino\_22Jul2017\_9M7eS\_haploidified\_headersFixed\_Sc9M7eS\_2\_HRSCAF\_41.{dataset}.merged.biallelic.fmissing{fmiss}.eigenvec - results/{dataset}/pca/sumatran\_rhino\_22Jul2017\_9M7eS\_haploidified\_headersFixed\_Sc9M7eS\_2\_HRSCAF\_41.{dataset}.merged.biallelic.fmissing{fmiss}.eigenval | |
| Output files | |
| - results/{dataset}/pca/sumatran\_rhino\_22Jul2017\_9M7eS\_haploidified\_headersFixed\_Sc9M7eS\_2\_HRSCAF\_41.{dataset}.merged.biallelic.fmissing{fmiss}.pc1\_pc3.pdf | |

##### Rule FROH\_min\_2Mb\_plot

×

Rule properties

|  |  |
| --- | --- |
| Jobs | 1 |
| Input files | |
| - results/{dataset}/ROH/sumatran\_rhino\_22Jul2017\_9M7eS\_haploidified\_headersFixed\_Sc9M7eS\_2\_HRSCAF\_41.{dataset}.merged.biallelic.fmissing{fmiss}.hwe0.05.homsnp{homsnp}.homkb{homkb}.homwinsnp{homwinsnp}.homwinhet{homwinhet}.homwinmis{homwinmis}.homhet{homhet}.FROH\_min\_2Mb\_table.txt | |
| Output files | |
| - results/{dataset}/ROH/sumatran\_rhino\_22Jul2017\_9M7eS\_haploidified\_headersFixed\_Sc9M7eS\_2\_HRSCAF\_41.{dataset}.merged.biallelic.fmissing{fmiss}.hwe0.05.homsnp{homsnp}.homkb{homkb}.homwinsnp{homwinsnp}.homwinhet{homwinhet}.homwinmis{homwinmis}.homhet{homhet}.FROH\_min\_2Mb\_plot.pdf | |

##### Rule FROH\_min\_2Mb\_table

×

Rule properties

|  |  |
| --- | --- |
| Jobs | 1 |
| Input files | |
| - /proj/sllstore2017093/b2016342/b2016342\_nobackup/lts/genome\_erosion\_pipeline/verena\_testing/testdata/reference/sumatran\_rhino\_22Jul2017\_9M7eS\_haploidified\_headersFixed\_Sc9M7eS\_2\_HRSCAF\_41.genome - results/modern/ROH/sumatran\_rhino\_22Jul2017\_9M7eS\_haploidified\_headersFixed\_Sc9M7eS\_2\_HRSCAF\_41.modern.merged.biallelic.fmissing0.1.hwe0.05.homsnp25.homkb100.homwinsnp250.homwinhet3.homwinmis15.homhet750.hom - results/modern/ROH/sumatran\_rhino\_22Jul2017\_9M7eS\_haploidified\_headersFixed\_Sc9M7eS\_2\_HRSCAF\_41.modern.merged.biallelic.fmissing0.1.hwe0.05.homsnp25.homkb100.homwinsnp250.homwinhet3.homwinmis15.homhet750.hom.indiv - results/historical/ROH/sumatran\_rhino\_22Jul2017\_9M7eS\_haploidified\_headersFixed\_Sc9M7eS\_2\_HRSCAF\_41.historical.merged.biallelic.fmissing0.1.hwe0.05.homsnp25.homkb100.homwinsnp250.homwinhet3.homwinmis15.homhet750.hom - results/historical/ROH/sumatran\_rhino\_22Jul2017\_9M7eS\_haploidified\_headersFixed\_Sc9M7eS\_2\_HRSCAF\_41.historical.merged.biallelic.fmissing0.1.hwe0.05.homsnp25.homkb100.homwinsnp250.homwinhet3.homwinmis15.homhet750.hom.indiv | |
| Output files | |
| - results/{dataset}/ROH/sumatran\_rhino\_22Jul2017\_9M7eS\_haploidified\_headersFixed\_Sc9M7eS\_2\_HRSCAF\_41.{dataset}.merged.biallelic.fmissing{fmiss}.hwe0.05.homsnp{homsnp}.homkb{homkb}.homwinsnp{homwinsnp}.homwinhet{homwinhet}.homwinmis{homwinmis}.homhet{homhet}.FROH\_min\_2Mb\_table.txt | |

##### Rule ROHs

×

Rule properties

|  |  |
| --- | --- |
| Jobs | 2 |
| Input files | |
| - results/{dataset}/ROH/sumatran\_rhino\_22Jul2017\_9M7eS\_haploidified\_headersFixed\_Sc9M7eS\_2\_HRSCAF\_41.{dataset}.merged.biallelic.fmissing{fmiss}.hwe0.05.bed - results/{dataset}/ROH/sumatran\_rhino\_22Jul2017\_9M7eS\_haploidified\_headersFixed\_Sc9M7eS\_2\_HRSCAF\_41.{dataset}.merged.biallelic.fmissing{fmiss}.hwe0.05.bim - results/{dataset}/ROH/sumatran\_rhino\_22Jul2017\_9M7eS\_haploidified\_headersFixed\_Sc9M7eS\_2\_HRSCAF\_41.{dataset}.merged.biallelic.fmissing{fmiss}.hwe0.05.fam - results/{dataset}/ROH/sumatran\_rhino\_22Jul2017\_9M7eS\_haploidified\_headersFixed\_Sc9M7eS\_2\_HRSCAF\_41.{dataset}.merged.biallelic.fmissing{fmiss}.hwe0.05.nosex | |
| Output files | |
| - results/{dataset}/ROH/sumatran\_rhino\_22Jul2017\_9M7eS\_haploidified\_headersFixed\_Sc9M7eS\_2\_HRSCAF\_41.{dataset}.merged.biallelic.fmissing{fmiss}.hwe0.05.homsnp{homsnp}.homkb{homkb}.homwinsnp{homwinsnp}.homwinhet{homwinhet}.homwinmis{homwinmis}.homhet{homhet}.hom - results/{dataset}/ROH/sumatran\_rhino\_22Jul2017\_9M7eS\_haploidified\_headersFixed\_Sc9M7eS\_2\_HRSCAF\_41.{dataset}.merged.biallelic.fmissing{fmiss}.hwe0.05.homsnp{homsnp}.homkb{homkb}.homwinsnp{homwinsnp}.homwinhet{homwinhet}.homwinmis{homwinmis}.homhet{homhet}.hom.indiv - results/{dataset}/ROH/sumatran\_rhino\_22Jul2017\_9M7eS\_haploidified\_headersFixed\_Sc9M7eS\_2\_HRSCAF\_41.{dataset}.merged.biallelic.fmissing{fmiss}.hwe0.05.homsnp{homsnp}.homkb{homkb}.homwinsnp{homwinsnp}.homwinhet{homwinhet}.homwinmis{homwinmis}.homhet{homhet}.hom.summary | |
| Container image | |
| docker://quay.io/biocontainers/plink:1.90b6.12--heea4ae3\_0 | |
| Code | |
| ``` 1 2 ``` \| ``` plink --bfile {params.bfile} --homozyg --homozyg-window-threshold 0.05 --allow-extra-chr --homozyg-snp {params.homsnp} --homozyg-kb {params.homkb} --homozyg-window-snp {params.homwinsnp} --homozyg-window-het {params.homwinhet} --homozyg-window-missing {params.homwinmis} --homozyg-het {params.homhet} --out {params.roh} 2> {log} ``` | |

##### Rule vcf2plink\_hwe

×

Rule properties

|  |  |
| --- | --- |
| Jobs | 2 |
| Input files | |
| - results/{dataset}/ROH/sumatran\_rhino\_22Jul2017\_9M7eS\_haploidified\_headersFixed\_Sc9M7eS\_2\_HRSCAF\_41.{dataset}.merged.biallelic.fmissing{fmiss}.hwe0.05.recode.vcf.gz - results/{dataset}/ROH/sumatran\_rhino\_22Jul2017\_9M7eS\_haploidified\_headersFixed\_Sc9M7eS\_2\_HRSCAF\_41.{dataset}.merged.biallelic.fmissing{fmiss}.hwe0.05.recode.vcf.gz.tbi | |
| Output files | |
| - results/{dataset}/ROH/sumatran\_rhino\_22Jul2017\_9M7eS\_haploidified\_headersFixed\_Sc9M7eS\_2\_HRSCAF\_41.{dataset}.merged.biallelic.fmissing{fmiss}.hwe0.05.bed - results/{dataset}/ROH/sumatran\_rhino\_22Jul2017\_9M7eS\_haploidified\_headersFixed\_Sc9M7eS\_2\_HRSCAF\_41.{dataset}.merged.biallelic.fmissing{fmiss}.hwe0.05.bim - results/{dataset}/ROH/sumatran\_rhino\_22Jul2017\_9M7eS\_haploidified\_headersFixed\_Sc9M7eS\_2\_HRSCAF\_41.{dataset}.merged.biallelic.fmissing{fmiss}.hwe0.05.fam - results/{dataset}/ROH/sumatran\_rhino\_22Jul2017\_9M7eS\_haploidified\_headersFixed\_Sc9M7eS\_2\_HRSCAF\_41.{dataset}.merged.biallelic.fmissing{fmiss}.hwe0.05.nosex | |
| Container image | |
| docker://quay.io/biocontainers/plink:1.90b6.12--heea4ae3\_0 | |
| Code | |
| ``` 1 2 ``` \| ``` plink --vcf {input.vcf} --make-bed --allow-extra-chr --out {params.bfile} 2> {log} ``` | |

##### Rule compress\_roh\_vcf

×

Rule properties

|  |  |
| --- | --- |
| Jobs | 2 |
| Input files | |
| - results/{dataset}/ROH/sumatran\_rhino\_22Jul2017\_9M7eS\_haploidified\_headersFixed\_Sc9M7eS\_2\_HRSCAF\_41.{dataset}.merged.biallelic.fmissing{fmiss}.hwe0.05.recode.vcf | |
| Output files | |
| - results/{dataset}/ROH/sumatran\_rhino\_22Jul2017\_9M7eS\_haploidified\_headersFixed\_Sc9M7eS\_2\_HRSCAF\_41.{dataset}.merged.biallelic.fmissing{fmiss}.hwe0.05.recode.vcf.gz - results/{dataset}/ROH/sumatran\_rhino\_22Jul2017\_9M7eS\_haploidified\_headersFixed\_Sc9M7eS\_2\_HRSCAF\_41.{dataset}.merged.biallelic.fmissing{fmiss}.hwe0.05.recode.vcf.gz.tbi | |
| Container image | |
| docker://quay.io/biocontainers/bcftools:1.9--h68d8f2e\_9 | |
| Code | |
| ``` 1 2 3 ``` \| ``` bcftools view -Oz -o {output.compressed} {input.vcf} 2> {log} && bcftools index -f -t {output.compressed} 2>> {log} ``` | |

##### Rule filter\_vcf\_hwe

×

Rule properties

|  |  |
| --- | --- |
| Jobs | 2 |
| Input files | |
| - results/{dataset}/vcf/sumatran\_rhino\_22Jul2017\_9M7eS\_haploidified\_headersFixed\_Sc9M7eS\_2\_HRSCAF\_41.{dataset}.merged.biallelic.fmissing{fmiss}.vcf.gz - results/{dataset}/vcf/sumatran\_rhino\_22Jul2017\_9M7eS\_haploidified\_headersFixed\_Sc9M7eS\_2\_HRSCAF\_41.{dataset}.merged.biallelic.fmissing{fmiss}.vcf.gz.csi | |
| Output files | |
| - results/{dataset}/ROH/sumatran\_rhino\_22Jul2017\_9M7eS\_haploidified\_headersFixed\_Sc9M7eS\_2\_HRSCAF\_41.{dataset}.merged.biallelic.fmissing{fmiss}.hwe0.05.recode.vcf | |
| Container image | |
| docker://biocontainers/vcftools:v0.1.16-1-deb\_cv1 | |
| Code | |
| ``` 1 2 ``` \| ``` vcftools --gzvcf {input.vcf} --hwe 0.05 --recode --recode-INFO-all --out {params.out} 2> {log} ``` | |
